# A detailed landscape of genomic alterations in malignant peripheral nerve sheath tumor cell lines challenges the current MPNST diagnosis

**DOI:** 10.1101/2022.05.07.491026

**Authors:** Miriam Magallon-Lorenz, Ernest Terribas, Marco Fernández, Gerard Requena, Inma Rosas, Helena Mazuelas, Itziar Uriarte, Alex Negro, Elisabeth Castellanos, Ignacio Blanco, George DeVries, Hiroyuki Kawashima, Eric Legius, Hilde Brems, Viktor Mautner, Lan Kluwe, Nancy Ratner, Margaret Wallace, Juana Fernández Rodriguez, Conxi Lázaro, Jonathan A Fletcher, David Reuss, Meritxell Carrió, Bernat Gel, Eduard Serra

## Abstract

**Background:** Malignant peripheral nerve sheath tumors (MPNSTs) are soft tissue sarcomas that arise from the peripheral nervous system. Half of the tumors develop in the context of the genetic disease Neurofibromatosis type 1 (NF1) and the rest are sporadic sarcomas. MPNSTs have a dismal prognosis due to their aggressiveness and tendency to metastasize, and new treatment options are needed. The diagnosis of MPNSTs can be challenging, especially outside of the NF1 context since specific histological criteria have not been completely established. Genomic analysis may both facilitate differential diagnoses and suggest precision medicine strategies.

**Methods:** We generated a complete genomic resource of a set of widely used human NF1-related and sporadic MPNST cell lines by applying ploidy analysis, whole genome and whole exome sequencing and SNP-array analysis, complemented by methylome-based classification and immunofluorescence of cell identity markers (SOX9, SOX10, S100B).

**Results:** NF1 MPNST cell lines faithfully recapitulated the genomic copy number profile of primary MPNSTs. Structural variants were key players in the complete inactivation of most recurrently altered tumor suppressor genes (TSGs) (*NF1, CDKN2A, SUZ12/EED*), while small variants played a minor role in the NF1 context, both concerning TSG inactivation and due to the absence of gain-of-function mutations. In clear contrast, the sporadic cell lines (STS-26T, HS-Sch-2, HS-PSS) did not recapitulate the copy number profile of primary MPNSTs. They carried different TSG inactivation and exhibited gain-of-function mutations by predicted kinase activation or generation of fusion genes. Mutational frequencies and signatures emerged as promising informative tools for aiding in MPNST differential diagnosis. Due to the multiple genomic differences exhibited, we complemented their characterization using a methylome-based classifier. All NF1-related cell lines were assigned within the MPNST group, while sporadic cell lines clustered either with melanomas or with an uncertain MPNST-like sarcoma group. The staining of cell identity markers reinforced the idea of a potential misdiagnose of the MPNSTs used to derive the sporadic cell lines analyzed.

**Conclusions:** Deep genomic analysis, together with methylome-based sarcoma classification and cell identity marker analysis, challenged the MPNST identity of sporadic cell lines. Results presented here open an opportunity to revise MPNST differential diagnosis and classification.

## Background

Malignant peripheral nerve sheath tumors (MPNSTs) are aggressive soft tissue sarcomas that arise from cells of the peripheral nervous system and account for 3-10% of all malignant soft tissue tumors [1]. Half of these tumors develop in the context of the tumor predisposition syndrome Neurofibromatosis type 1 (NF1) while the other half are sporadic neoplasms [2,3]. The MPNST incidence in the general population is 1 in 100000 [2–4] whereas the lifetime risk of an NF1 individual developing an MPNST is 10-15% [2,5]. Due to its invasive growth and propensity to metastasize, MPNSTs have a poor prognosis and are the leading cause of adult NF1-related mortality [2,5]. Like many soft tissue sarcomas, complete resection with wide margins is essential in MPNST therapy, followed by radiation and/or chemotherapy [6–8].

MPNSTs are usually high-grade malignant spindle cell neoplasms arising in association with large peripheral nerves [9]. Their diagnosis can be challenging, especially outside of individuals with NF1, since MPNSTs are rare tumors and specific histological criteria have not been completely established [10–12]. In the context of NF1, MPNSTs often progress from a pre-existing benign plexiform neurofibroma, commonly through an intermediate discrete nodular tumor termed atypical neurofibroma or ANNUBP [12,13]. Although neurofibromas contain numerous S100B/SOX10-positive Schwann cells and CD34-positive fibroblasts, the expression of both markers is significantly reduced or absent in MPNSTs [12].

MPNSTs contain hyperploid and highly rearranged genomes with a low mutation burden [14–17]. Several tumor suppressor genes (TSGs) are commonly mutated, including *NF1, CDKN2A*, and components of the polycomb repressive complex 2 (PRC2), including *SUZ12* and *EED. TP53* is also frequently lost or mutated. MPNSTs also show recurrently altered chromosomal regions, particularly constituting somatic copy number gains (revised in Serra et al. 2020 [18]). Complete loss of *CDKN2A*, often caused by structural alterations, seems to constitute a bottleneck for MPNST formation [19,20].

Established cell lines are an important tool for gaining insight into cancer biology and treatment. However, there are also different caveats in their use as faithful and useful models, with issues including misidentification and cross-contamination and poor characterization of similarity to their tumor source [21–24]. There is no dedicated registry for MPNST cell lines, but according to Cellosaurus (https://web.expasy.org/cellosaurus/), around forty different MPNST cell lines may have been established by different laboratories, derived from both sporadic and NF1-related MPNSTs. Some of these MPNST cell lines are well distributed among labs [25], [26]or deposited in global repositories (ATCC, RIKEN). These lines have been used as a primary tool for the identification of molecular pathways involved in MPNST pathogenesis [27,28], and served, for instance, for the identification of MEK inhibitors as useful therapeutic agents [29]. Some can be engrafted in mice to generate genuine orthotopic MPNST tumors [30,31]. However, a systematic and comprehensive genomic characterization of these MPNST cell lines is still missing, limiting the use of these cell lines for precision medicine or pharmacogenomic studies.

In this work, we performed a deep genomic characterization of 8 commonly used MPNST cell lines [25,26], identifying heterogeneity regarding the structure of the genome, the inactivation of tumor suppressor genes, the frequency of mutations, the mutational signatures and the presence of gain-of-function mutations, especially among sporadic MPNST cell lines. This characterization challenged the identity of the sporadic MPNST cell lines studied and prompted us to use a methylome sarcoma classifier and to perform immunofluorescence of known cell identity markers. Our results, in addition to providing a valuable resource, uncover the necessity of systematically analyzing MPNSTs, combining pathology with genomic and molecular results, for improved differential diagnosis and classification of these malignancies.

## Methods

### Cell lines

In this study, we used a set of MPNST cell lines that contains some of the most frequently used MPNST cell lines together with a few which can be found in known repositories (ATCC, RIKEN). We studied six NF1-associated cell lines: S462 (RRID:CVCL_1Y70) [32], ST88-14 (RRID:CVCL_8916) [33], NF90-8 (RRID:CVCL_1B47) [34], sNF96.2 (RRID:CVCL_K281) [30], NMS-2 (RRID:CVCL_4662) [35], and T265 (RRID:CVCL_S805)[36], although the latter was discarded as it was found to be misidentified; and three sporadic lines: STS-26T (RRID:CVCL_8917) [37], HS-Sch-2 (RRID:CVCL_8718) [38] and HS-PSS (RRID:CVCL_8717). **Table 1** summarizes clinical information about patients and tumors from whom these cell lines were established. Human foreskin fibroblast (HFF-1, ATCC: SCRC-1041) were used as control cells for ploidy analysis. All cell lines were cultured under standard conditions (37°C and 5% CO2) with High Glucose DMEM with sodium pyruvate (Biowest) supplemented with 10% FBS (Biowest) and 2mM L-glutamine (Life technologies). They were passaged and harvested using trypsin-EDTA (Life technologies).

**Table 1.** General description of the eight MPNST cell lines analyzed.

|  | **S462** | **ST88-14** | **NF90-8** | **sNF96.2** | **NMS-2** | **STS-26T** | **HS-Sch-2** | **HS-PSS** |
| --- | --- | --- | --- | --- | --- | --- | --- | --- |
| **Type of human MPNST** | primary, grade IV | primary | primary, from PNF | primary | primary | metastasis, grade III/III | primary, low grade | UNK |
| **MPNST localization** | thigh | retroperitoneum | left forearm | leg | right thigh | left scapula | left thigh | prostate |
| **NF1/Sp** | NF1 | NF1 | NF1 | NF1 | NF1 | Sp | Sp | Sp |
| **Age of patient** | 19 | 24 | 17 | 27 | 30 | 51 | 54 | UNK |
| **Sex of patient** | F | M | F | M | M | F | F | M |
| **Original Reference** | Frahm-2004 | Fletcher-1991 | Legius-1994 | Perrin-2007 | Imaizumi-1998 | Dahlberg-1993 | Sonobe-2000 | n/a |
| ***NF1* constitutional pathogenic variant** | c.6855C>A<br>p.Y2264X<br>Frahm-2004 | c.1649dupT<br>p.V551Gfs*7<br>Varin-2016 | c.3904_3910del<br>p.D1302Yfs*5<br>Wu-1999 | c.3683delC<br>p.N1229Mfs*11<br>Perrin-2007 | **c.6999+1G>T**<br>**(splice donor)** |  |  |  |
| ***NF1* somatic pathogenic variant** | LOH | LOH | LOH | LOH | **LOH** | **LOH** | **c.270_288del**<br>**p.E91Nfs\*6;**<br>**c.3113+1G>A**<br>**(splice donor)** |  |

MPNST: Malignant peripheral nerve sheath tumor; NF1/Sp: NF1-related/Sporadic MPNST; F: Female; M: Male; Bold annotations: mutations described in this manuscript; LOH: Loss of Heterozygosity; UNK: Unknow

### DNA extraction

Total DNA was extracted from cell lines using the Gentra Puregene Kit (Qiagen). DNA was quantified with Nanodrop 1000 spectrophotometer (Thermo Scientific). For SNP array, whole exome and genome sequencing and methylome experiments, a fluorescence-based quantification of DNA was performed either by using the Quant-iT™ PicoGreen® dsDNA Assay (Life technologies) or a Qubit fluorometer (Life technologies).

### STR profiling

DNA fingerprinting of short tandem repeats (STRs) was conducted for all MPNST cell lines using the AmpFlSTR Identifiler Plus Amplification kit (Life technologies) following the manufacturer’s instructions. This kit is based on the analysis of 16 microsatellites, including the nine STRs used by the ATCC®.

### Calculation of cell ploidy by flow cytometry

About 1-2·10^6^ cells from each cell line were trypsinized, washed with PBS, and fixed in ice-cold 70% ethanol for 2h at −20°C. Then, cells were washed with PBS and resuspended in a citrate-phosphate buffer for at least 30 min, up to 2h. Cells were then washed with PBS-1% FBS and propidium iodide (PI) was added. Cells in PI solution were treated with DNAse-free RNAse A for 30-45 min at 37°C and were ready for flow cytometry analysis. All samples were analyzed on a FACSCanto II flow cytometer (BD Biosciences, San Jose CA) and a total of 10,000 single cells were analyzed for each sample. Aggregated cells were excluded by gating out on a bi-parametric plot with DNA content pulse area versus width. Data was analyzed using FlowJo software (BD Biosciences, San Jose, CA). HFF-1 were used as 2n control cells.

### *NF1* mutational status

Sanger sequencing was used to confirm previously described *NF1* constitutional pathogenic variant of the cell lines S462 [32], ST88-14 [39], NF90-8 [40], sNF96.2 [30]. In this project, we identified the constitutional *NF1* pathogenic variant of NMS-2 cell line, and the *NF1* pathogenic variants present in the HS-Sch-2 cell line by whole-exome sequencing (see below) which were also confirmed by Sanger sequencing. We used specific primers targeting the mutation region in each case and the BigDye Terminator v.3.1 Sequencing Kit (Applied Biosystems). Sequences were generated with the ABI Prism 3100 Genetic Analyzer (Applied Biosystems) and analyzed with CLC Main Workbench 6 software.

### SNP-array analysis

SNP-array data from the different cell lines and tumors was obtained from Magallon-Lorenz et al. (2021) [19] and deposited in Synapse (https://www.synapse.org/, syn22392179) [19]. In short, the analysis was performed using Illumina BeadChips (Human660W-Quad, OmniExpress v1.0 and OmniExpress 1.2) at the IGTP High Content Genomics Core Facility. Raw data were processed with Illumina Genome Studio to extract B allele frequency (BAF) and log R ratio (LRR). We used GAP [41] to perform copy-number calling.

### Whole-exome sequencing (WES) and whole-genome sequencing (WGS)

WES from the 8 MPNST cell lines was also previously analyzed in Magallón-Lorenz et al. (2021) [19] and deposited in Synapse (https://www.synapse.org/, syn22392179)[19]. In short, the exome was captured using Agilent SureSelect Human All Exon V5 kit (Agilent, Santa Clara, CA, US) and sequenced in a HiSeq instrument (Illumina, San Diego, CA, US) at Centro Nacional de Analisis Genomicos (CNAG, Barcelona, Spain) to a median of 165.5 million 100 bp paired-end reads per sample. Sequencing reads were then mapped with BWA-MEM [42] against GRCh38 genome.

The whole genome of two cell lines (ST88-14 and S462) had already been sequenced for Magallón Lorenz et al. (2021) [19]. The WGS of the other 6 cell lines were produced for this work at BGI (Shenzhen, China). In short, the 6 libraries were prepared following standard DNBseq protocols, sequenced in a BGISEQ-500 to a median of 881 million 150 bp paired-end reads per sample and mapped with bwa mem against the GRCh38 genome.

### Selection of putatively pathogenic somatic variants using WES and WGS

Small nucleotide variants were called with strelka2 [43] and annotated with annovar [44]. We filtered strelka2 results from WGS data to select potentially driver variants affecting protein function as follows: we selected exonic and splicing variants and removed all synonymous variants then, we filtered out variants with a population frequency (AF_popmax) higher than 1%, classified as benign in ClinVar [45], annotated as benign or likely benign in InterVar automated[46], present in 3 or more of the cell lines or classified as pathogenic in more than 5 out of 7 in-silico predictors (SIFT pred [47], Polyphen2 HDIV pred[48], LRT pred, Mutation Taster pred[49], Mutation Assessor pred[50], FATHMM pred[51], CLNSIG[45]) Then, we filtered out those variants with a variant allele frequency (VAF) lower than 0.1 as these variants were deemed as unlikely to be present in the original malignant cell. In addition, we removed non-frameshift deletion or insertion variants present in dbSNP and variants in highly variable genes *(MUC3A, MUC5AC, OR52E5, OR52L1, SMPD1, PRAMEF* and *LILR*). Finally, we filtered out the variants present in dbSNP except for those included in COSMIC somatic mutations (https://ftp.ncbi.nlm.nih.gov/snp/others/rs_COSMIC.vcf.gz) or the International Cancer Genome Consortium (ICGC) (https://ftp.ncbi.nlm.nih.gov/snp/others/snp_icgc.vcf.gz) variant lists. WES data was processed using the same approach and used to validate the variants identified in WGS data.

### Mutational signatures

Raw variants called by Strelka2 in WGS data were also the basis for the mutational signature analysis. Since normal pairs were not available, we applied a series of filters to approximate a somatic callset: we filtered out the variants with a population frequency (AF_popmax) higher than 1%, called in more than one cell line, with a variant allele frequency (VAF) lower than 0.1 and, variants in highly variable genes (*MUC3A, MUC5AC, OR52E5, OR52L1, SMPD1, PRAMEF* and *LILR*). We also filtered out the variants in dbSNP except for those present in COSMIC and ICGC. We used this call set enriched in somatic variants with the mutSignatures [52] R package to estimate the contribution of each of the thirty COSMIC mutational signatures to the mutational profile of each cell line.

### Copy number variants (CNVs) from WGS

We called copy-number alterations from WGS using CNVkit [53] with the recommended settings for WGS data with no matched normal pair (flat reference, difficult region blacklist (https://github.com/Boyle-Lab/Blacklist/blob/master/lists/hg38-blacklist.v2.bed.gz), -no-edge option and 1000 bp bins). To obtain the exact copy number profile of each sample we used the threshold method with sample-specific thresholds defined considering the ploidy of each cell line obtained by flow cytometry. Summarized and per-cell line copy number profiles were plotted using the CopyNumberPlots (10.18129/B9.bioc.CopyNumberPlots) and karyoploteR [54] R packages.

### Structural variants and detection of fusion genes

We used LUMPY [55] via Smoove (https://github.com/brentp/smoove) as a structural variant (SV) caller with parameters for small cohorts and excluding the problematic regions defined in https://github.com/hall-lab/speedseq/blob/master/annotations/exclude.cnvnator_100bp.GRCh38.20170403.bed. We also used CliffHunteR (https://github.com/TranslationalBioinformaticsIGTP/CliffHunteR), an in-house developed sensitivity-oriented R package for breakpoint detection, and a thorough visual inspection using Integrative Genomic Viewer (IGV) [56] to detect breakpoints affecting tumor suppressor genes associated with MPNSTs *(NF1, CDKN2A, SUZ12, EED, TP53, PTEN, RB1*). To discard germline structural variants, we filtered out SVs present in the Database of Genomic Variants (DGV) [57] and the SVs with the same breakpoints in more than two MPNST cell lines. Inter-chromosomal and intra-chromosomal rearrangements were plotted using circos [58]. We defined the genome region affected by an SV as 1Mb upstream and downstream of its breakpoints. To investigate the presence of known fusion genes, we crossed the SV breakpoints detected by LUMPY and CliffHunteR with the fusion genes in COSMIC (https://cancer.sanger.ac.uk/census).

### DNA methylation and Uniform Manifold Approximation and Projection (UMAP) analysis

DNA methylation profiles were generated using the Infinium MethylationEPIC (850k) BeadChip array (Illumina, San Diego, USA) according to the manufacturer’s instructions. The data was processed as previously described[59]. The two-dimensional UMAP [60] embedding was created using the 20,000 most variable CpGs from the DNA methylation profiles of the cell lines and the reference cohorts of soft tissue tumors [59]. The UMAP analysis was performed using the R package umap (version 0.2.7.0) with default parameters except for n_neighbors=8.

### Validation of inter-chromosomal rearrangements

Inter-chromosomal rearrangements detected by LUMPY or CliffHunteR affecting genes commonly altered in MPNST were validated by PCR and Sanger sequencing. PCR primers, annealing temperatures and amplicon lengths are summarized in **Supplementary Document 6**.

### Fusion gene validation

EML4-ALK v5a fusion gene breakpoints were detected by LUMPY in HS-PSS cell line. EML4-ALK fusion gene was validated by RT-PCR and Sanger sequencing. Total RNA from HS-PSS cell line was extracted using the 16 LEV simplyRNA Purification Kit (Promega) following the manufacturer’s instructions in the Maxwell 16 Instrument (Promega). RNA was quantified with a Nanodrop 1000 spectrophotometer (Thermo Scientific). RNA (0.5 μg) was reverse transcribed using the Superscript III reverse transcriptase enzyme (Life technologies) according to the manufacturer’s instructions. PCR primers, annealing temperatures, and amplicon lengths were previously described by Takeuchi et al. (2008) [61].

## Results

### A genomic resource for NF1-associated and sporadic MPNST cell lines

We started with 9 NF1-associated and sporadic MPNST cell lines for a short tandem repeat (STR) authentication analysis and a comprehensive genomic characterization. **Table 1** summarizes information on patients and MPNSTs from which cell lines were established and on their *NF1* mutational status. It also provides a reference to the original description of each cell line. We first performed an interspecies PCR of all cell lines to identify any possible interspecies cross-contamination (**Supplementary Document 1**). Then we performed a human STR authentication analysis to identify any possible cross-contamination or misidentification among cell lines of human origin (**Supplementary Table 1**). All STR profiles matched the STR profiles published in Cellosaurus and ATCC when available. However, in this process we identified the same STR profile for ST88-14 and T265 cell lines (**Supplementary Document 2**) in all ST88-14- and T265-related samples provided by different laboratories. To rule out which cell line was misidentified we analyzed the oldest ST88-14 and T265 stored vials in their original labs and more conclusively, the primary tumor from which ST88-14 cell line was isolated (**Supplementary Document 2**). We identified the ST88-14 cell line as the genuine cell line for that STR profile, *NF1* germline mutation and somatic copy number alteration landscape, and dismissed the use of T265 cell line, which we assume was misidentified at some point after its establishment and expansion.

We performed a comprehensive genomic characterization of the remaining 8 MPNST cell lines including flow cytometry, SNP-array analysis, whole-exome sequencing and whole-genome sequencing techniques. We compiled information about their ploidy, global copy-number profile and loss of heterozygosity (LOH) status, structural rearrangements, single nucleotide variants (SNVs) and mutational signatures, and summarized the mutational status of a set of selected MPNST-related genes. With all these data we elaborated a practical summary sheet for each cell line, containing the most relevant information (**Supplementary Document 3**).

### A plethora of different ploidies

We first intended to characterize the karyotype of each MPNST cell line by spectral karyotyping (SKY) and G-banding staining. Both techniques produced results difficult to summarize consistently (data not shown), possibly due to the high degree of variability when analyzing multiple metaphases from the same cell line and to the highly rearranged nature of MPNST genomes. Therefore, we decided to analyze the ploidy by propidium iodide staining analysis using flow cytometry (**Figure 1a**), which revealed a striking diversity among the 8 MPNST cell lines. Most of them exhibited ploidies higher than 2n, three clearly around 4n, yet two cell lines were 2n or less. In addition, two cell lines, NF90-8 and NMS-2, showed two cell populations with different ploidy, resembling the result of a genome duplication event in the population with higher ploidy (**Figure 1a**). STS-26T cell line was composed of two subpopulations with slightly different ploidies. The mean ploidy of each cell line was also calculated considering this diversity.

**Figure 1:**
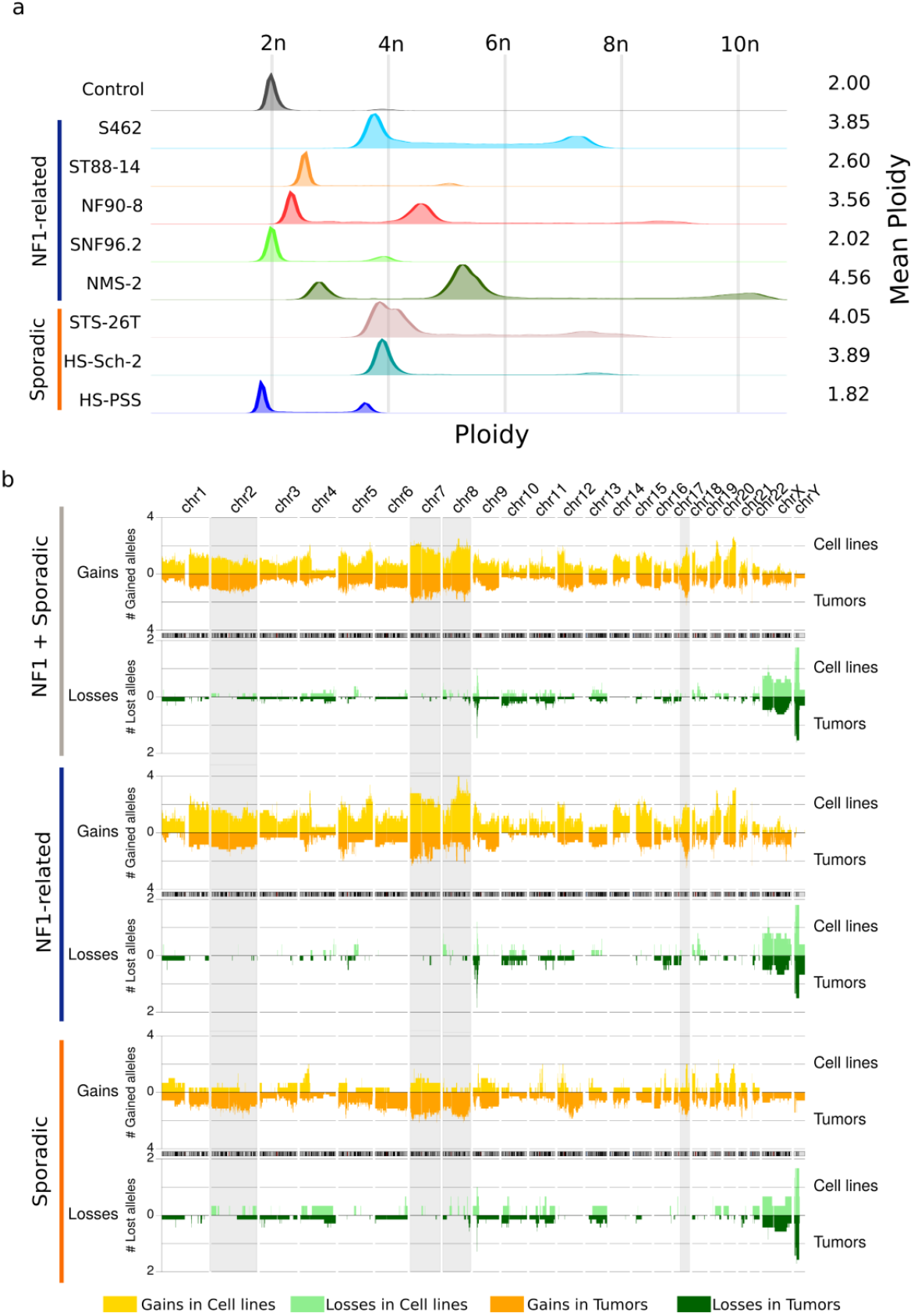
MPNST cell lines have different ploidies and faithfully recapitulate the genomic copy number profile of primary MPNSTs. **(a)** Ploidy status of MPNST cell lines obtained by flow cytometry. Each row represents a cell line, with the mean ploidy shown on the right side. HFF were used as 2n control cells **(b)** Aggregated copy number profiles comparing MPNST cell lines and tumors. Y axis represents the mean number of gained (orange) or lost (green) alleles by sample type and Y axis genomic position. Cell lines are plotted in the upper part of each graph, represented by light colors and tumors in the lower part, represented by dark colors and with inverted Y axis. Three different groups of samples are represented. From top to bottom: all MPNSTs; NF1-associated MPNSTs; and sporadic MPNSTs.

### MPNST cell lines faithfully recapitulate the genomic copy number profile of primary MPNSTs

Next, we performed a global copy number analysis using SNP-array data from the 8 MPNST cell lines. To verify whether these cell lines were capturing the copy number profiles present in primary MPNST tumors, we displayed the copy number profiles of gains and losses separately, comparing cell lines and tumors (**Figure 1b**). Taken together, the copy number profiles of the 8 MPNST cell lines faithfully recapitulated those present in primary MPNSTs. The most recurrently gained genomic regions were chromosomes 7, 8 and 17q, and chromosome 9p was the most recurrently lost genomic region. When we separated NF1-related and sporadic MPNSTs, NF1-related cell lines maintained these profiles and recurrences, but the three sporadic cell lines (**Figure 1b**) differ substantially in many genomic regions (see for example shaded grey regions in **Figure 1b**).

We also performed a copy number analysis from all cell lines based on WGS data. In addition, we obtained a B-allele frequency (BAF)-like profile from variant-allele frequencies (VAF) and a log-R ratio (LRR) from coverage. We plotted them together with the resulting LOH determination and copy number calling from SNP-array comparing both independent sets of data (**Figure 2**; **Supplementary Document 4** for a high-resolution profile). The use of both technologies for generating the same data allowed the validation of the results with all cell lines displaying similar BAF and LRR profiles. It also underscored the difficulty of using copy number callers for analyzing the highly altered MPNST genomes, indicating the necessity of estimating copy number changes in MPNSTs by different means. In addition to its known hyperploid genome, our analysis identified a high degree of generalized LOH (e.g.: almost the entire genome of the sNF96.2 cell line), perhaps pointing to the inactivation of TSGs before the gain of chromosomal regions. Finally, it must be noted that the copy number profile of a given MPNST cell line differed a bit from lab to lab, in terms of a certain number of different genomic alterations. By comparing the genomic profiles of different batches of the same cell lines obtained from distinct labs, we found that despite being quite stable, genomes of MPNST cell lines can accumulate changes in chromosomal regions (**Supplementary Document 5**). Part of this inter-laboratory variability could be removed after growing the cell line as a xenograft in mice [31] when growth selective pressure is applied.

**Figure 2:**
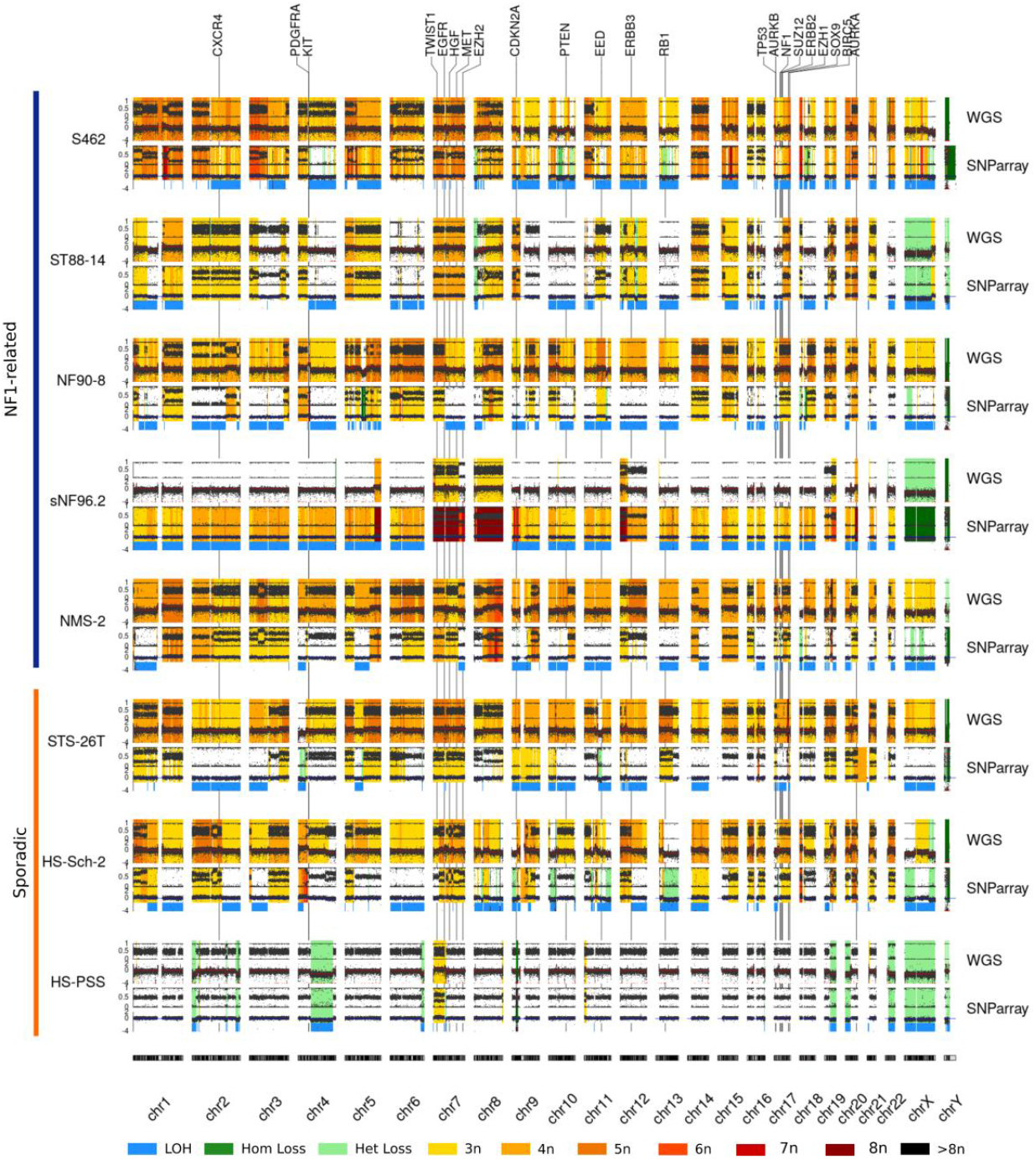
Global view of the copy number profiles of all MPNST cell lines studied. Copy number profiles from SNP-array and WGS data. For both technologies, B-allele frequency (BAF) and Log-R Ratio (LRR) are represented (see M&M for details). Warm colors represent copy number gains, whereas light green indicates a heterozygous loss and dark green a homozygous loss. Loss of heterozygosity (LOH) is highlighted by a thick blue line. The genomic location of MPNST-associated genes is indicated by a vertical black line with the gene symbol at the upper part of the graph. See **Supplementary Document 3** for a high-resolution profile of each cell line, chromosome by chromosome.

### Structural variants as key players for tumor suppressor gene inactivation

Ploidy and copy number profile comprise a partial description of the genomic status of MPNST cell lines. We also took advantage of WGS data to analyze the presence of structural variants, represented by different types of chromosomal rearrangements (**Figure 3**). This analysis revealed that in addition to being hyperploid, MPNST genomes were highly rearranged, adding an extra layer of complexity. Structural changes were spread over all chromosomes, although certain cell lines such as ST88-14 and HS-PSS showed some genomic regions with a high frequency of adjacent rearrangements. Importantly, as previously reported [19], structural changes caused the inactivation of key MPNST TSGs, such as *CDKN2A*, PRC2 genes and *TP53* (**Supplementary Document 6**), that had been previously missed when WGS data was not available. Furthermore, WGS also facilitated the unexpected identification of a translocation resulting in the generation of a fusion gene EML4-ALK variant 5a [61] in the HS-PSS cell line, which was validated by RT-PCR, amplification and sequencing (**Supplementary Document 6**). The significant number of structural variants identified together with the copy number profiles and ploidy exhibited, demonstrates the complex genomic landscape of MPNST cell lines, with highly altered but also fairly stable genomes.

**Figure 3:**
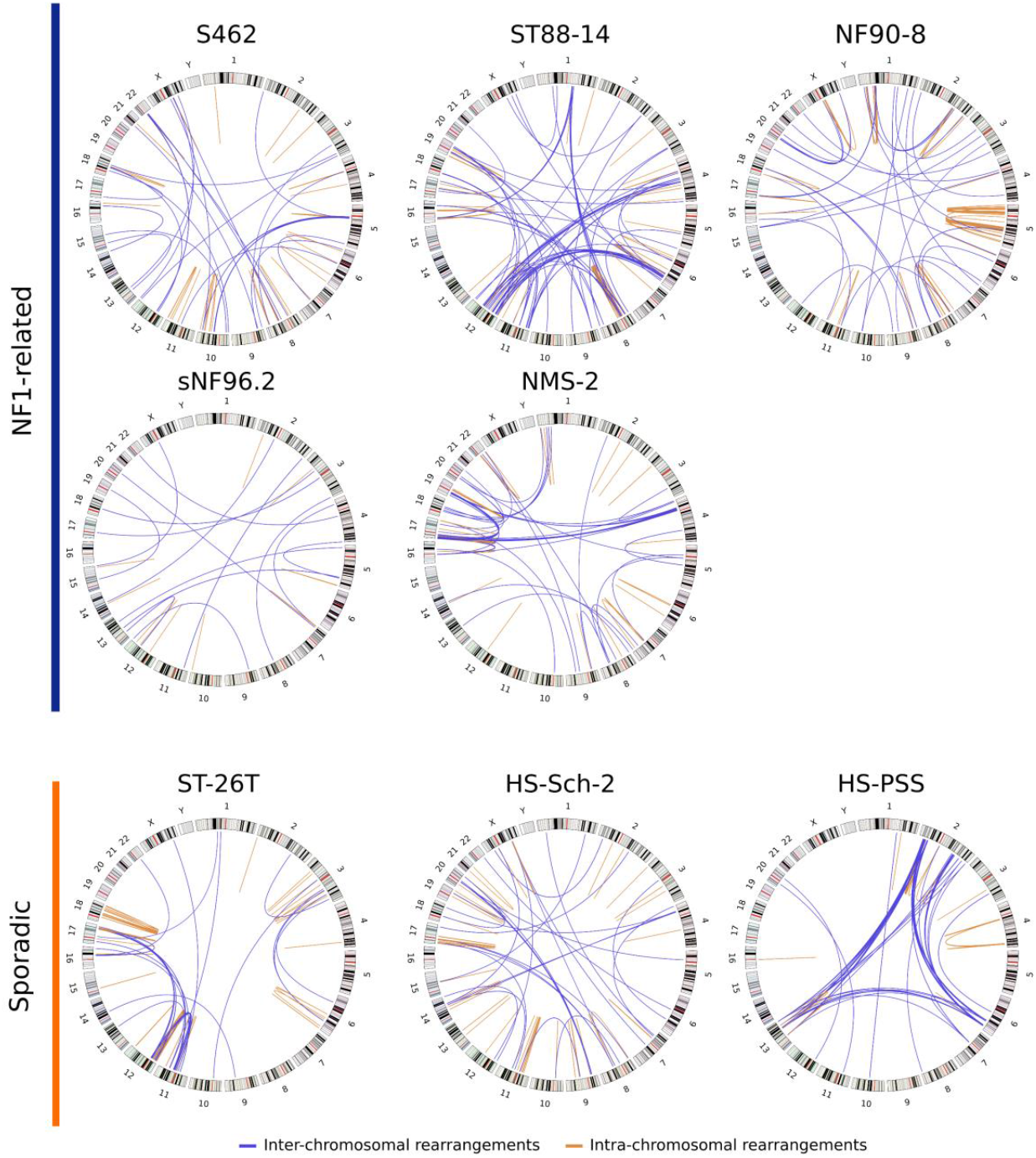
Structural rearrangements as key players for tumor suppressor gene inactivation. Circos plots showing the chromosomal rearrangements identified in the different MPNST cell lines. Blue lines represent inter-chromosomal rearrangements and orange lines intra-chromosomal rearrangements. These rearrangements were obtained using Lumpy and CliffHunteR on WGS data.

### The limited importance of small genomic variants

In contrast to the rich number of gross structural alterations, fine-scale analysis of small variants, including single-nucleotide variants (SNVs) and small indels, uncovered a relatively moderate impact of these alterations in MPNSTs. We used WES and WGS data from all cell lines to call small variants. Since we did not have available non-tumoral tissue counterparts for these cell lines, we filtered the small variants to partially remove germline variants and obtain a call set enriched in somatic variants. The number of variants with a potential impact on protein function in this dataset was modest (**Supplementary Table 2**), particularly for NF1-associated MPNST cell lines, which harbored a mean of 129 SNVs. We then used this quasi-somatic variant dataset to estimate the contribution of the COSMIC mutational signatures [62] in the variant profile of the MPNST cell lines (**Figure 4a**). This analysis did not identify a particular mutational mechanism prevalent in MPNSTs and showed a major contribution of clock-like signatures (signatures 1 and 5), comparable with previous observations in other sarcomas [17]. However, the STS-26T cell line in addition to exhibiting a higher number of mutations compared to the other cell lines (about two times the average of the rest of the cell lines), also presented an important contribution from signature 7, predominantly found in skin cancers (see below).

**Figure 4:**
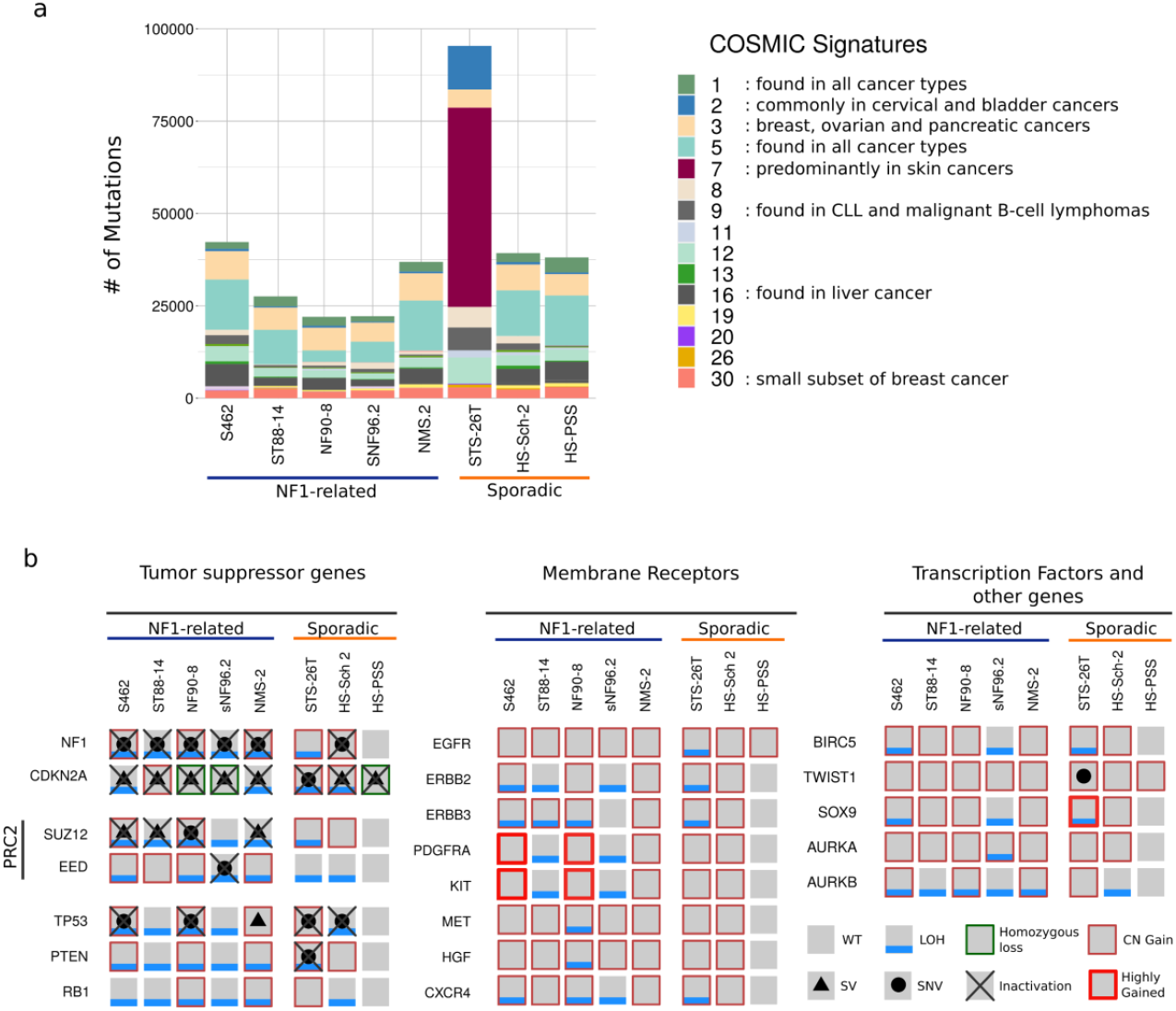
Small genomic variants have limited importance in altering MPNST-associated genes. **(a)** Mutation number and estimated contribution of COSMIC mutational signatures in MPNST cell lines. Each bar represents the total number of somatically-enriched single nucleotide variants (SNVs) found in an MPNST cell line. Each color represents a different COSMIC mutational signature. **(b)** Status of the commonly altered genes in MPNSTs. A grey square represents a wild type (WT) status of the gene; a blue line indicates the presence of loss of heterozygosity (LOH); a black dot a small nucleotide variant affecting the gene; a black triangle indicates that the gene is affected by a structural variant (SV); copy number gain (CN gain) is represented by a thin red square; and a thick red square denotes a highly gained region affecting the gene. A homozygous loss of a gene is specified by a thin dark green square and the complete inactivation of a gene is represented by a black cross. Note that all NF1-related MPNST cell lines have the complete inactivation of *NF1, CDKN2A* and the *PRC2* complex. In addition, *CDKN2A* is also inactivated in sporadic cell lines.

The functional impact of small variants in oncogenes and TSGs was also moderate. We identified some MPNST-related genes inactivated by pathogenic SNVs (**Figure 4b; Supplementary Table 3**). In addition to germline *NF1* mutations, somatic mutations also affected *NF1*, as well as other genes including *TP53*, PRC2 genes, and *PTEN*. Remarkably, we did not identify gain-of-function mutations in oncogenes, except a *BRAF* V600E mutation in the STS-26T cell line. In contrast, we identified gains in genomic regions containing receptors, especially a highly gained region containing *PDGFRA* and *KIT* in two NF1-related cell lines (S462 and NF90-8) (**Figure 4b**). The most frequently inactivated gene in our set of cell lines was *CDKN2A*, a known bottleneck for MPNST development [19,20]. The fact that this gene was inactivated by a point mutation only in one cell line, exemplifies the relatively low functional impact of small variants compared to structural variants in MPNST initiation.

### The combination of genome, methylome and expression marker analysis, represent useful tools for a better differential diagnosis and classification of MPNSTs

This complete description of the genomic status of MPNST cell lines uncovered a fair degree of diversity among them, prompting us to question the MPNST identity of the sporadic cell lines and to perform a further characterization. Since sarcomas comprise a morphologically heterogeneous class of tumors and their diagnosis has been hampered by a high misclassification rate, we decided to perform methylome analysis of all cell lines in comparison to established reference cohorts of different peripheral nerve sheath and soft tissue tumors [59] to clarify whether there was a problem in the diagnosis of MPNSTs or on the identification and classification of different MPNST types. **Figure 5a** shows a dimension reduction using Uniform Manifold Approximation and Projection (UMAP) analysis plot. While all NF1-related MPNST cell lines were located within the MPNST cluster, the sporadic cell lines lay within the melanoma cluster (STS-26T and HS-Sch-2 cell lines) or within the not fully characterized group of MPNST-like sarcomas (HS-PSS cell line). With these results, we decided to further characterize the 8 cell lines by performing immunostaining for 3 markers: SOX9, SOX10 and S100B. SOX9 is normally expressed in MPNSTs [28] and also in melanomas [63]. SOX10 and S100B are markers that define the neural crest-Schwann cell differentiation axis but also the neural crest-melanocytic path. Both markers are frequently reduced or absent in MPNSTs according to the WHO classification [64], lost in transitions of ANNUBP towards a low-grade MPNST [12] and also significantly downregulated/absent using expression analysis[28]; and data not shown). In contrast, SOX10 and S100B are frequently expressed in melanoma [65]. All cell lines stained positive for SOX9 (**Figure 5b**). All NF1-related cell lines stained negative for SOX10 and S100B, as did the STS-26T sporadic cell line. In contrast, HS-PSS and HS-Sch-2 sporadic cell lines stained positive for SOX10 and, in addition, HS-Sch-2 also stained positive for S100B (**Figure 5b**), moving them away from a classic MPNST identity.

**Figure 5:**
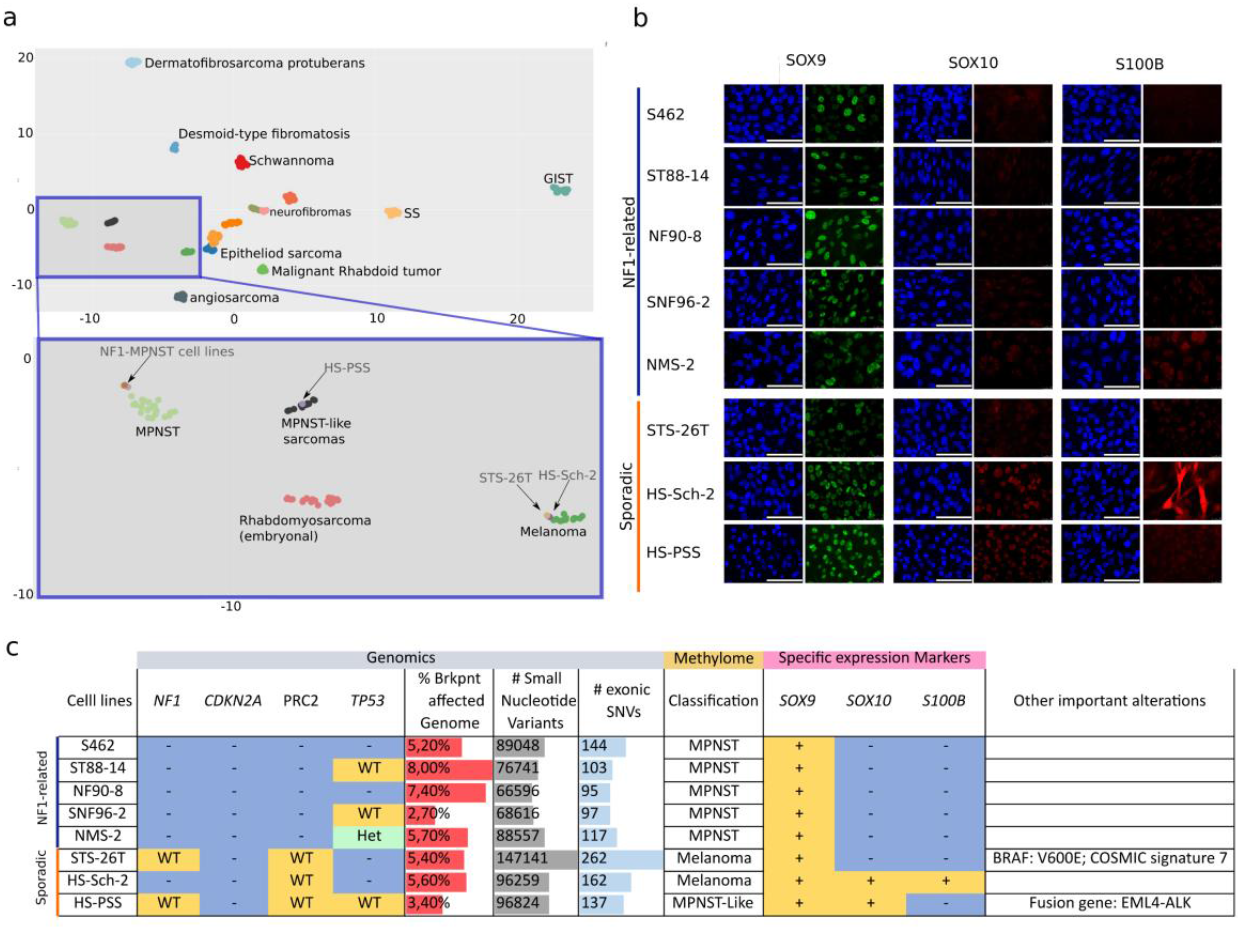
Genome, methylome and marker analysis constitute useful additional tools for a better differential diagnosis and classification of MPNSTs. **(a)** UMAP analysis of the methylome profile of the MPNST cell lines in comparison to different soft tissue tumor types [59]. The upper part of the plot provides a global view of the classification. Each dot represents a sample and every color a different sarcoma type. The lower part of the graph is an inset magnification of a specific part containing all the MPNST cell lines analyzed. All NF1-related MPNST cell lines group within the MPNST methylation group (light green). HS-PSS cell line clustered with MPNST-like sarcomas group (black) and STS-26T and HS-Sch-2 clustered together with the melanoma group (dark green). **(b)** Immunofluorescence images showing the expression of SOX9, SOX10 and S100B markers in the different cell lines. DAPI was used to stain nuclei. Scale bars: 100μm **(c)** Summary table of the genomic, methylome and marker expression status of all MPNST cell lines. Genomics contains the status of *NF1, CDKN2A*, PRC2, and *TP53* genes (blue, complete gene inactivation (-); yellow wild type (WT); light green heterozygous deletion (Het)); percentage (%) of breakpoints affecting the genome +/− 1mb; the number (#) of small nucleotide variants per sample; the # of exonic small nucleotide variants with a potential impact on protein function. Methylome contains the methylome-based classification of each cell line. Expression markers contain the expression of SOX9, SOX10 and S100B identity markers (yellow for expression (+); blue for absence (-). The “other important information” column summarizes additional relevant information identified in each cell line.

Altogether, the fine landscape of genomic alterations, methylome-based classification, marker expression and particular informative gain-of-function mutations, captured a fair degree of variability among MPNST cell lines (**Figure 5c**) uncovering the probable misidentification of some of the tumors from which the sporadic cell lines were isolated and also the need of a complete and systematic characterization of additional MPNST tumors and cell lines to better understand whether MPNSTs constitute a homogeneous group of tumors or there exist different types.

## Discussion

We performed a deep genomic characterization of 9 of the most distributed and used MPNST cell lines [25] including cell lines banked in repositories (ATCC, RIKEN). Before this analysis, we performed authentication assays which resulted in discarding the T265 cell line since it exhibited the same STR profile as the ST88-14 and its matched primary MPNST (**Supplementary Document 2**).

Despite the diverse ploidies exhibited by the MPNST cell lines, NF1-related cell lines faithfully reproduced the copy number profiles present in primary MPNST tumors, something that was not true for sporadic cell lines. These results reinforce the idea that despite the high degree of genomic alterations, MPNSTs contain quite stable genomes, as shown by comparing primary tumors with derived orthotopic PDX [31]. Our use of WGS was crucial for a more complete detection of genomic alterations present in MPNSTs, due to the significant number of structural variants present [19]. This was especially important for detecting the inactivation of MPNST-related tumor suppressor genes (TSGs), particularly for *CDKN2A* and the PRC2-related genes (*SUZ12/EED*). The use of WGS also allowed the identification of fusion genes, like EML4-ALK, generated by an inversion affecting both genes in the HS-PSS sporadic cell line. The presence of fusion genes is common in other types of soft tissue sarcomas that otherwise contain genomes with few genomic alterations. However, fusion genes are not common in the karyotypically complex MPNSTs [66]. In concordance with this idea, and supporting a non-MPNST identity, in addition to the EML4-ALK fusion gene, HS-PSS cell line contained a ploidy close to 2n, containing only a few copy number alterations and structural changes (**Figure 1a**, **Figure 2, Figure3, and Supplementary Document 4**).

In contrast to the importance of somatic copy number alterations and structural rearrangements, MPNST cell lines exhibited a modest frequency of mutations, with moderate functional impact, mainly involving the inactivation of *TP53*. Notably, the frequency and the type of mutation signatures exhibited provided an important differential indicator. While all cell lines exhibited a similar number of mutations and similar mutational signatures, the STS-26T cell line contained a much higher number of mutations and an important contribution from mutational signature 7, predominantly found in skin cancers.

Genomic alterations or mutations constituting a gain-of-function were not common in NF1-related MPNST cell lines. In fact, we identified only two cell lines (S462 and NF90-8) with a highly gained region in chromosome 4 containing the *PDGFRA* and *KIT* receptors, consistent with previous reports [67,68] but not, for instance, kinase-activating mutations or translocations, like those involving NTRK genes[69]. In contrast, we identified the *BRAF* V600E mutation in the STS-26T cell line and the already mentioned EML4-ALK fusion gene in the HS-PSS cell line.

Our deep genomic characterization (ploidy, copy number profile, structural variants, mutation frequencies and signatures, presence of gain-of-function mutations, altered MPNST-related genes) questioned the MPNST identity of the analyzed sporadic cell lines. Methylome-based classification [59] and immunofluorescence of cell identity markers (SOX9, SOX10, S100B) complemented genomic analysis. STS-26T cell line contained a functional PRC2 complex and the *BRAF* V600E mutation. It also exhibited a much higher mutation frequency than the other cell lines and an important COSMIC signature 7, predominantly present in skin cancers as previously described by Hayward et al. (2017) [70]. Finally, a methylome classifier unequivocally classified it as a melanoma. Taking together our compiled evidence, in our opinion, the original diagnosis of a “malignant schwannoma” [37] if made today would probably be “melanoma”. In this regard, it would be interesting to further analyze MPNSTs with *BRAF* V600E mutations described in the literature [71–73] using additional tools like mutation frequencies and signatures, methylome classifier and cell identity marker expression. The HS-Sch-2 cell line showed a WT status for PRC2 genes but harbored the complete inactivation of *NF1* and *CDKN2A*. It was classified as a melanoma by the methylome classifier and expressed the markers SOX10 and S100B. The expression of these two markers is lost in transitions from atypical neurofibromas to MPNSTs and is commonly significantly reduced or absent in MPNSTs [12]. The HS-Sch-2 cell line also stained negative for the melanoma marker Melan-A (data not shown). The combination of positive staining for SOX10 and S100B and negative staining for Melan-A is characteristic of desmoplastic melanoma [74], which also commonly exhibits the complete inactivation of *NF1* and *CDKN2A* [75,76], neurotropism and nerve infiltration [77,78], the latter described in the original publication of this cell line [38]. Finally, HS-PSS also showed a WT status for PRC2 genes, was assigned to the provisional and not fully characterized “MPNST-like sarcoma” methylation group, was positive for SOX10, contained an almost unaltered genome proximal to 2n but harbored a translocation generating the fusion gene EML4-ALK. This fusion gene is associated with a type of sarcoma termed epithelioid inflammatory myofibroblastic sarcoma [79,80] which also contain a spindle cell component, being the most probable identity of the tumor from which HS-PSS was derived.

## Conclusions

In summary, the new genomic and epigenomic characterization of MPNST cell lines provided in this work uncovered the misidentification of the commonly used NF1-related T265 MPNST cell line and, in addition, compiled multiple pieces of evidence to question the identity of the three sporadic MPNST cell lines analyzed, proposing alternative identities for all of them: a melanoma for STS26T; a desmoplastic melanoma for the HS-Sch-2 cell line; and an epithelioid inflammatory myofibroblastic sarcoma for the HS-PSS. These results may imply the need of determining the impact of their use in previous and probably current works being performed, considering the new information provided. It also alerts us, as a scientific community, that we need to improve the characterization and control of the cell lines and tissues we use in our research. But above all, it provides an opportunity to look ahead and improve our understanding of what is an MPNST and which types might exist. In this regard, a systematic combination by different laboratories of a histological characterization together with these new ways of analyzing genomes and epigenomes, opens the door to revising the manner we perform differential diagnostics of MPNSTs and related tumors.

Our results, in addition to generating a valuable resource for the study of new therapeutic strategies for MPNSTs, uncover the need to systematically analyze MPNSTs, combining pathology with genomic and molecular techniques. Genomic analysis such as copy number profiles, structural variants, mutation frequencies and signatures, presence of gain-of-function mutations, and the inactivation of specific TSGs, together with methylome-based sarcoma classification and cell identity marker analysis, emerge as valuable tools for a better differential diagnosis and classification of MPNSTs.

## List of abbreviations

MPNST: Malignant peripheral nerve sheath tumor
NF1: Neurofibromatosis type 1
STR: Short tandem repeats
TSG: Tumor suppressor genes
PRC2: Polycomb repressive complex 2
LOH: Loss of heterozygosity
VAF: Variant allele frequency
BAF: B-allele frequency
LRR: Log-R ratio
SNV: Single nucleotide variant
SV: Structural variant
CNV: Copy number variant

## Data availability

SNP-array, WES and WGS data is publicly available at Synapse (https://www.synapse.org/, syn22392479) and is part of the NF Data Portal (https://nf.synapse.org/).

## Acknowledgements

We would like to thank Aurora Sánchez from the Molecular Genetics and Biochemistry Service, CDB, Hospital Clínic (Barcelona) and Marta Salido, Mar Rodríguez-Rivera and Blanca Espinet from the Molecular Cytogenetics Laboratory, IMIM-Hospital del Mar (Barcelona), for their initial effort in the MPNST cell line cytogenetic characterization using G-banding and SKY. We thank the IGTP core facilities and their staff for their contribution and technical support: High Performance Computing; Translational Genomics Core Facility; High Content Genomics and Bioinformatics Core Facility. We would like to thank the constant support of the NF lay associations: Asociación de Afectados de Neurofibromatosis and ACNefi.

## Author contributions

Conception and design of the work: MM-L, ET, BG and ES. Project supervision: BG, ES. Bioinformatic analysis and visualization: MM-L, BG. Experimental work acquisition, analysis and interpretation: ET, MM-L, MC, MAF, GR, IR, HM, IU, AN, JF-R, BG, ES. Additional resources: JF, DR, CL. Methylome analysis and sarcoma classifier: DR. Writing original draft: MM-L, ET, BG and ES. Writing, reviewing, editing and scientific input: all authors. All authors also approved this version of the manuscript.

## Ethics declarations

This work has been approved by the Germans Trias i Pujol Hospital (HUGTiP) Ethics Committee.

