## Supplemental Documents for "A detailed landscape of genomic alterations in malignant peripheral nerve sheath tumor cell lines challenges the current MPNST diagnosis"

### Supplementary document 1: Interspecies PCR.

In order to find in our MPNST cell line set any possible cross-contamination of cell lines from the most used laboratory species mouse and rat we performed an interspecies PCR assay based on the amplification of the mitochondrial DNA gene cytochrome c oxidase subunit I (*COX1*) that discriminated among human, mouse and rat DNA. **Figure 1** shows the results of our interspecies PCR: no cross-contamination from either mouse or rat cell line was detected in our MPNST cell line set.

Specific primers for *COX1* gene for the three species were obtained from Parodi *et al*. [1]. Moreover, *COX1* gene sequences of human, mouse and rat were used for the design of a fourth primer pair to obtain a shared amplicon of this gene among the three species. **Table 1** shows a full list of all the primers used. Each interspecies PCR reaction included 100 ng of DNA, mixed with 5 uL of GoTaq® Flexi Buffer (Promega), 1.5 mM of $MgCl_2$ (Promega), 0.2 mM of each dNTP (vWR), 0.4 μM of each primer (Life technologies), and 0.75 U of GoTaq® G2 Flexi DNA Polymerase (Promega), in a total volume of 25 uL. PCR experiments were performed in a thermocycler (Applied Biosystems). Conditions for amplification were as follows: 95°C for 5 min; 30 cycles of 95°C for 30 s, 58°C for 30 s, and 72°C for 30 s; and a final step at 72°C for 5 min. A negative control without DNA template and three positive controls with DNA from human 293T cell line and rodent cell lines C2C12 (mouse) and PC12 (rat) were also included. Finally, 5 uL of each amplification product were run on a 1% agarose gel, stained with SybrSafe (Life technologies), visualized in a transilluminator, and photographed.

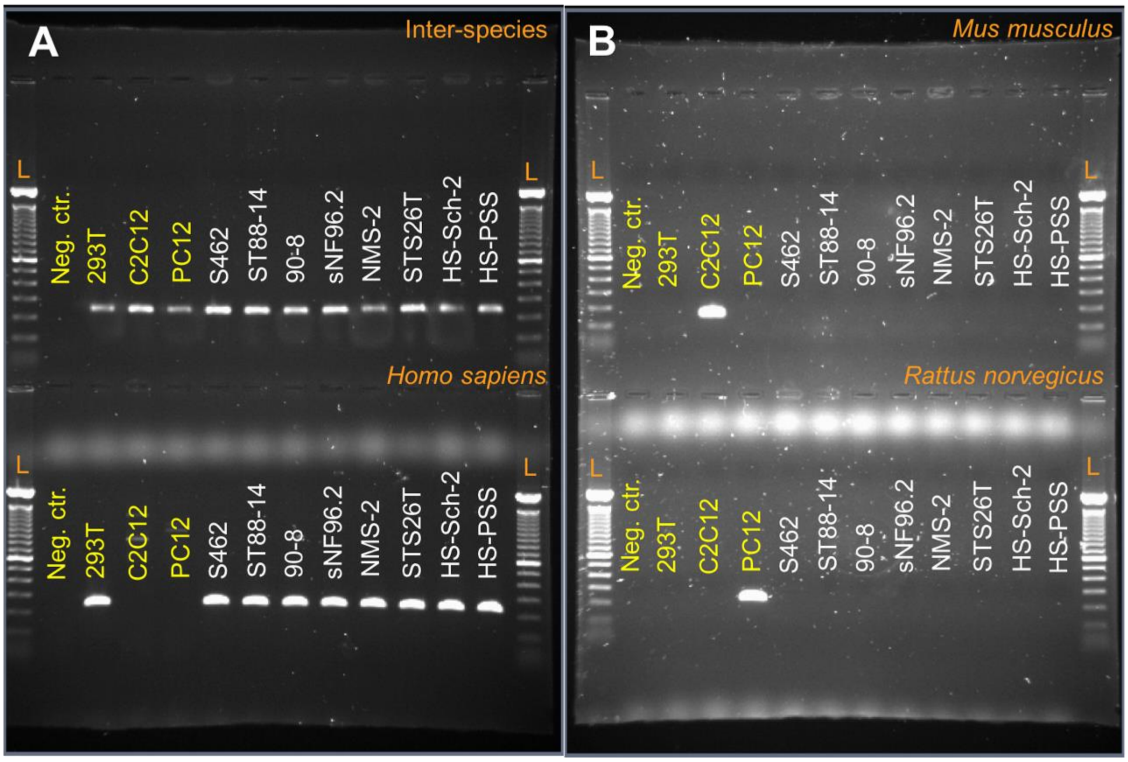


***Figure 1****. Agarose gel electrophoresis showing PCR amplicons from all four cytochrome c oxidase subunit I PCR experiments. A. Top: inter-species PCR showed 233-bp amplicons in all DNA samples. Bottom: Homo sapiens PCR showed 228-bp amplicons in human cell lines (293T and MPNST cell lines). B. Top: Mus musculus PCR showed a 150-bp amplicon only in the murine C2C12 cell line. Bottom: Rattus norvegicus PCR showed a 196-bp amplicon only in the PC12 cell line. Neg. ctr.: negative control; L: DNA ladder.*

| Species | Primer name | Sequence 5'-3' | Amplicon length |
|---|---|---|---|
| Interspecies | IC-COX1-F | GATGCWTACACCACATGAAA | 233 bp |
| | IC-COX1-R | GGGTTCGAWTCCTTCCTT | |
| Human | Hs-COX1-F | TTCGGCGCATGAGCTGGAGTCC | 228 bp |
| | Hs-COX1-R | TATGCGGGGAAACGCCATATCG | |
| Mouse | Mm-Cox1-F | ATTACAGCCGTACTGCTCCTAT | 150 bp |
| | Mm-Cox1-R | CCCAAAGAATCAGAACAGATGC | |
| Rat | Rn-Cox1-F | CGGCCACCCAGAAGTGTACATC | 196 bp |
| | Rn-Cox1-R | GGCTCGGGTGTCTACATCTAGG | |

***Table 1****. Primer sequences and amplicon sizes of interspecies, human, mouse and rat PCR experiments. W = A + T.*

#### *References*

1. Parodi B, Aresu O, Bini D, Lorenzini R, Schena F, Visconti P, et al. Species identification and confirmation of human and animal cell lines: A PCR-based method. Biotechniques. 2002;32:432–40.

### Supplementary document 2: Misidentification of the T265 MPNST cell line.

We obtained the MPNST cell lines used in this paper either from dedicated bioresources (RIKEN and ATCC) or from different collaborating laboratories. In some cases, the same cell line was sourced from distinct groups. That is the case of ST88-14, which was obtained from two labs. (**Figure 1**).

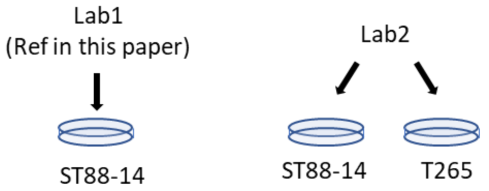


***Figure 1***. *MPNST cell lines were obtained from different labs.*

We first analyzed the small tandem repeat (STR) profile, to authenticate each cell line. The two cell lines from Lab2, ST88-14, and T265, showed the exact same STR profile. We compared them with the STR profile of the ST88-14 cell line we received from Lab 1, and they were all the same (**Table 1**). ST88-14 from Lab1 was finally used as the reference ST88-14 cell line (Ref) for this work.

*Table 1*. *Small Tandem Repeat profile of ST88-14 and T265 cell lines*

| Microsatellite | Chr. location | ST88-14 (Ref) Lab 1 | ST88-14 Lab2 | T265 Lab2 |
|---|---|---|---|---|
| D8S1179 | 8 | 14 | 14 | 14 |
| D21S11 | 21q11.2-q21 | 29, 32.3 | 29, 32.3 | 29, 32.3 |
| D7S820 | 7q11.21-22 | 8 | 8 | 8 |
| CSF1PO | 5q33.3-34 | 9, 12 | 9, 12 | 9, 12 |
| D3S1358 | 3p | 15, 18 | 15, 18 | 15, 18 |
| TH01 | 11p15.5 | 9 | 9 | 9 |
| D13S317 | 13q22-31 | 12 | 12 | 12 |
| D16S539 | 16q24-qter | 13 | 13 | 13 |
| D2S1338 | 2q35-37.1 | 17, 23 | 17, 23 | 17, 23 |
| D19S433 | 19q12-13.1 | 13, 14 | 13, 14 | 13, 14 |
| vWA | 12p12-pter | 16 | 16 | 16 |
| TPOX | 2p23-2per | 11, 12 | 11, 12 | 11, 12 |
| D18S51 | 18q21.3 | 12 | 12 | 12 |
| AMEL | Xp22.1-22.3 Yp11.2 | X, Y | X, Y | X, Y |
| D5S818 | 5q21-31 | 12, 13 | 12, 13 | 12, 13 |
| FGA | 4q28 | 21 | 21 | 21 |

Both cell lines sharing the exact same STR profile pointed to a probable misidentification of one of the cell lines, so we decided to analyze either the original tumor from which the cell line was established, or the oldest possible cell culture passage preserved in the labs that generated these cell lines. In this regard, for the ST88-14 cell line [1] we were able to test DNA from the primary tumor (here labeled as ST88-14_PT), DNA from an early ST88-14 culture passage, passage 69 (ST88-14_CLp69) (Lab3). On the other hand, for the T265 cell line [2] we were able to test DNA from the oldest cryopreserved T265 cell line vial stored in Lab4 where the cell line was established (**Figure 2**). From all these samples we performed both an STR profile (**Table 2**) and an SNP-array analysis (**Figure 3**).

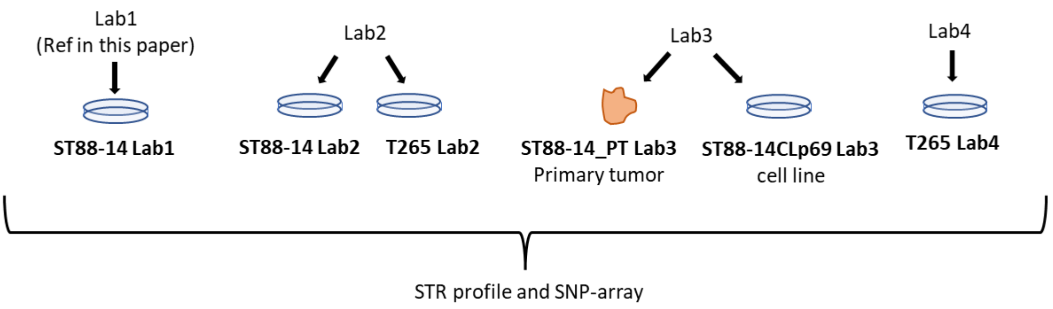


***Figure 2***. *Scheme explaining the origin of the different samples and the analysis performed*

*Table 2. STR profiles of all samples are represented in Figure 2, PT: primary tumor, CL: cell line; *: small peak.*

| Microsatellite | Chr. location | ST88-14<br>Lab 1 (Ref) | ST88-14_PT<br>Lab 3 | ST88-14_CLp69<br>Lab 3 | T265_Lab4<br>Lab4 |
|---|---|---|---|---|---|
| D8S1179 | 8 | 14 | 14 | 14 | 14 |
| D21S11 | 21q11.2-q21 | 29, 32.3 | 29, 32.3 | 29, 32.3 | 29, 32.3 |
| D7S820 | 7q11.21-22 | 8 | 8 | 8 | 8 |
| CSF1PO | 5q33.3-34 | 9, 12 | 9, 12 | 9, 12 | 9, 12 |
| D3S1358 | 3p | 15, 18 | 15, 18* | 15, 18 | 15, 18 |
| TH01 | 11p15.5 | 9 | 9 | 9 | 9 |
| D13S317 | 13q22-31 | 12 | 12 | 12 | 12 |
| D16S539 | 16q24-qter | 13 | 13 | 13 | 13 |
| D2S1338 | 2q35-37.1 | 17, 23 | 17, 23 | 17, 23 | 17, 23 |
| D19S433 | 19q12-13.1 | 13, 14 | 13, 14 | 13, 14 | 13, 14 |
| vWA | 12p12-pter | 16 | 16 | 16 | 16 |
| TPOX | 2p23-2per | 11, 12 | 11, 12 | 11, 12 | 11, 12 |
| D18S51 | 18q21.3 | 12 | 12 | 12 | 12 |
| AMEL | Xp22.1-22.3<br>Yp11.2 | X, Y | X, Y | X, Y | X, Y |
| D5S818 | 5q21-31 | 12, 13 | 12, 13 | 12, 13 | 12, 13 |
| FGA | 4q28 | 21 | 21 | 21 | 21 |

The STR analysis **(Table 2)** demonstrated that all tested cell lines shared the same STR profile as the primary tumor from which the cell line ST88-14 was derived, and thus, both cell lines that arrived at our laboratory (ST88-14 and T265) were actually the same ST88-14 cell line. We cannot rule out the possibility that there exists a genuine T265 cell line in some laboratory. However, the oldest available vial of T265 shared the STR profile of ST88-14, suggesting that the misidentification happened early after the cell line establishment.

In addition to the STR profile, we performed an SNP-array analysis of all available samples (Figure 3 and Figure 4).

In **Figure 3** we can observe the SNP-array profile of the samples obtained from Lab 3 compared with the reference ST88-14 used in this paper from Lab 1. All samples share the same genomic profile, with only minor differences between them, consistent with the expected *in-vitro* evolution of the cell culture.

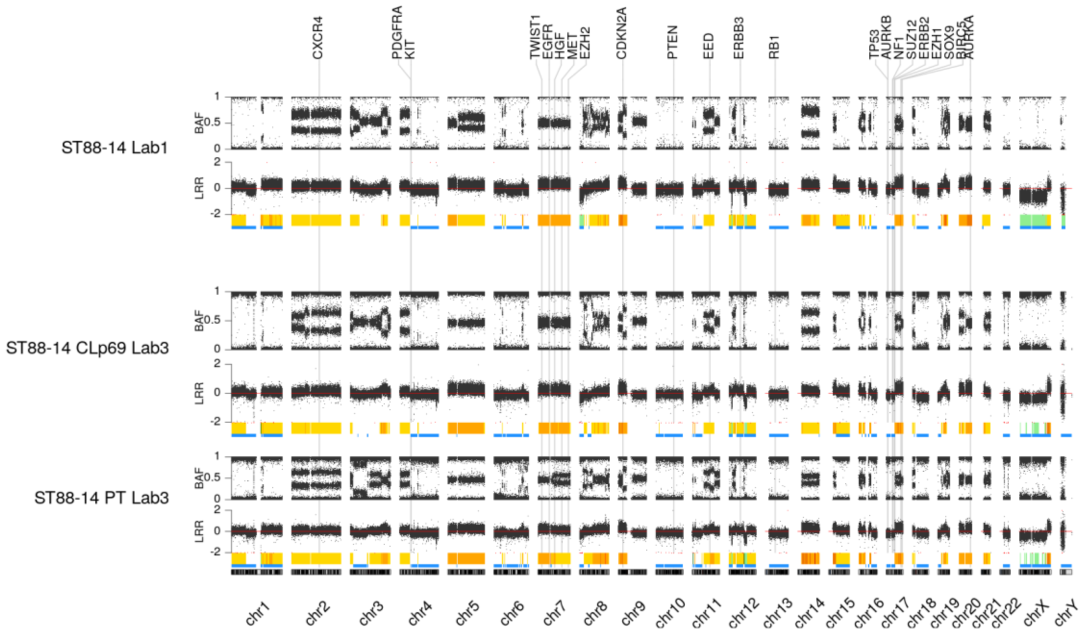


***Figure 3**. SNP-array profile of the samples we obtained from Lab 3 compared with the reference ST88-14 from Lab 1.*

**Figure 4** represents the SNP-array profile of T265 cell lines we obtained from Lab 2 and from Lab 4 compared to our reference ST88-14 cell line. T265 samples again share the genomic profile with the reference ST88-14, although with a few additional differences when compared with ST88-14 samples in **Figure 3**. This could indicate an earlier separation from the original ST88-14 or be the result of culture bottlenecks. In any case, the genomic profiles of both T265 cell line samples show a common origin with the ST88-14 cell line.

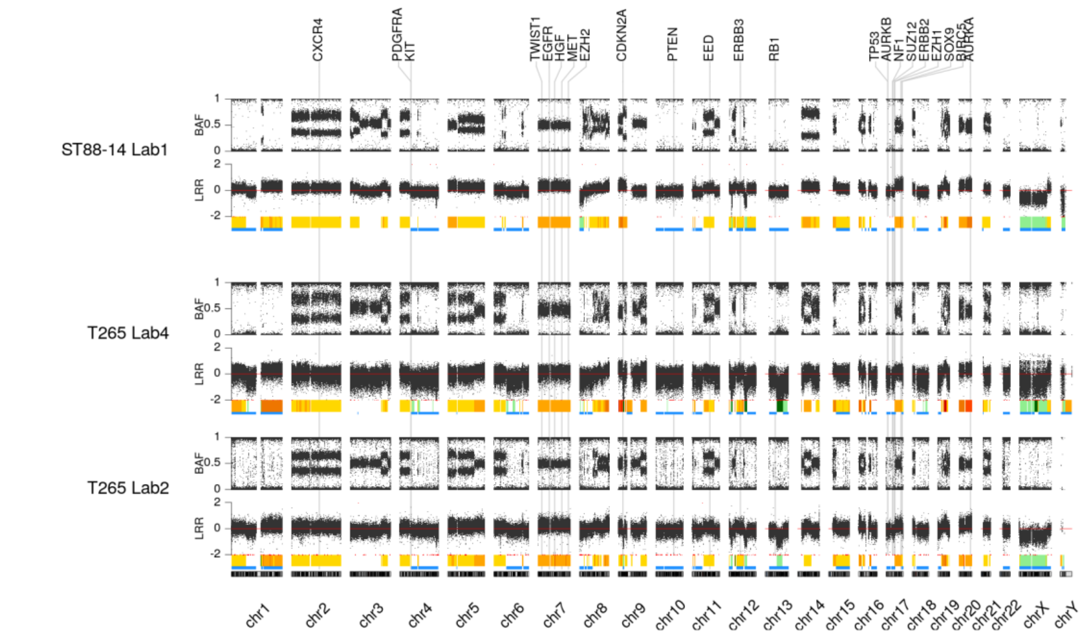


***Figure 4**. SNP-array profile of the T265 cell lines we obtained from Lab 2 and Lab 4compared with the reference ST88-14 from Lab 1.*

All in all, the STR profile and SNP-array analysis of current cell lines and the original primary tumor for ST88-14, confirm that the ST88-14 cell line originated from the expected primary tumor and that the current T265 cell line is, in reality, a misidentified ST88-14 cell line. In addition, the analysis of the oldest available T265 vial confirms that the misidentification happened very early after the establishment of the cell line. It is not clear if a real T265 cell line currently exists.

***References***

1. Fletcher JA, Kozakewich HP, Hoffer FA, Lage JM, Weidner N, Tepper R, et al. Diagnostic Relevance of Clonal Cytogenetic Aberrations in Malignant Soft-Tissue Tumors. New England Journal of Medicine. 1991;324:436–43.

2. Badache A, De Vries GH. Neurofibrosarcoma-derived Schwann cells overexpress platelet-derived growth factor (PDGF) receptors and are induced to proliferate by PDGF BB. Journal of Cellular Physiology. 1998;177:334–42.

# S462

RRID:CVCL_1Y70

NF1

Female, 19yo

#### Phenotype

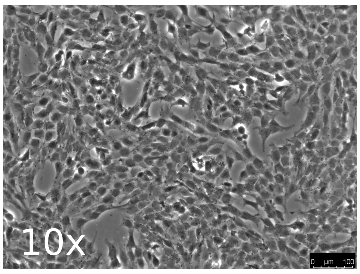


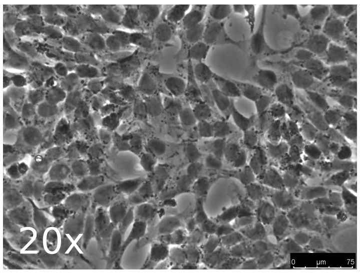


#### Ploidy

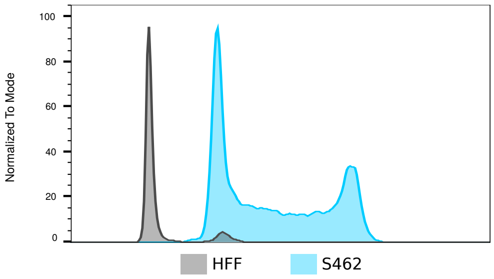


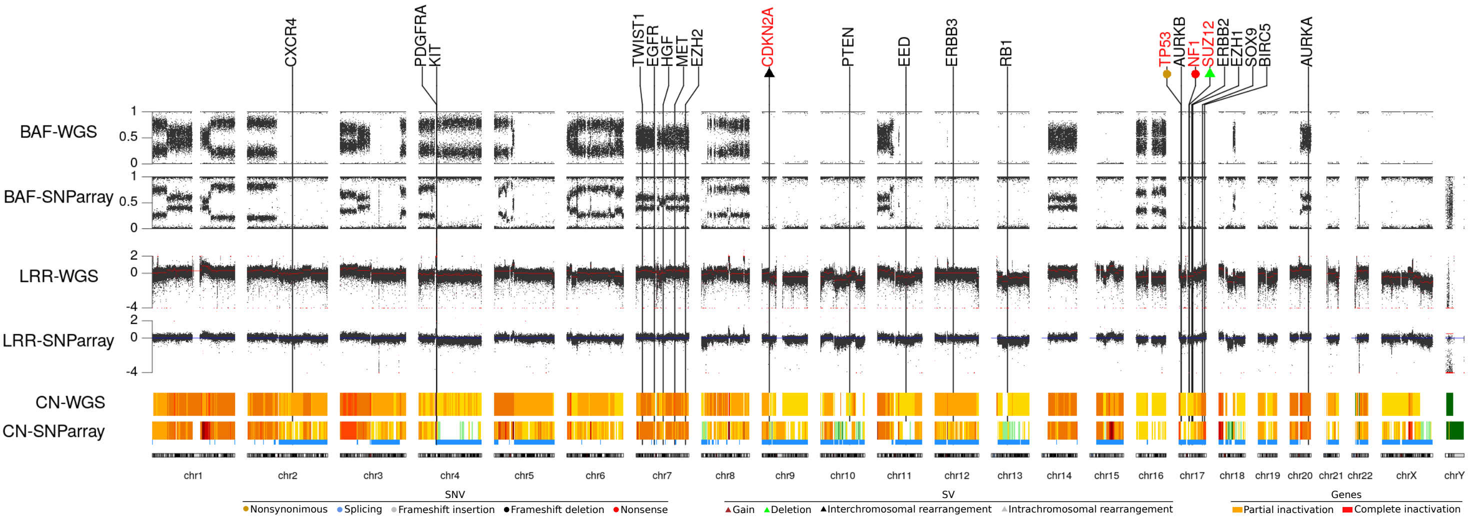


#### Genes

| Tumor Supressor | | Membrane Receptors | Transcription factors & other |
|---|---|---|---|
| NF1 | c.6855C>A;LOH | EGFR | BIRC5 |
| CDKN2A | Partial homozygous deletion | ERBB2 | TWIST1 |
| SUZ12 | Heterozygous deletion;LOH | ERBB3 | SOX9 |
| EED | | PDGFRA | AURKA |
| TP53 | c.329G>C;LOH | KIT | AURKB |
| PTEN | | MET | |
| RB1 | | HGF | |
| | | CXCR4 | |

WT · LOH · Homozygous loss · CN Gain · SV · SNV · Inactivation · Highly Gained

#### STR

| | |
|---|---|
| D8S1179 | 10, 12 |
| D21S11 | 29 |
| D7S820 | 8, 10 |
| CSF1PO | 12 |
| D3S1358 | 14, 17 |
| TH01 | 7, 8 |
| D13S317 | 12 |
| D16S539 | 11, 13 |
| D2S11338 | 23 |
| D19S433 | 14 |
| vWA | 19 |
| TPOX | 8 |
| D18S51 | 16 |
| AMEL | X |
| D5S818 | 12 |
| FGA | 20 |

#### Methylome class

MPNST

#### Marker expression

SOX9: Positive SOX10: Negative S100B: Negative

Additional alterations:

Additional information:

Aliases:

Original publication: Silke Frahm, Victor-F Mautner, Hilde Brems, et. al., Genetic and phenotypic characterization of tumor cells derived from malignant peripheral nerve sheath tumors of neurofibromatosis type 1 patients, Neurobiology of Disease, Volume 16, Issue 1, Pages 85-91, 2004. https://doi.org/10.1016/j.nbd.2004.01.006

# ST88-14

RRID:CVCL_8916

NF1

Male, 24yo

#### Phenotype

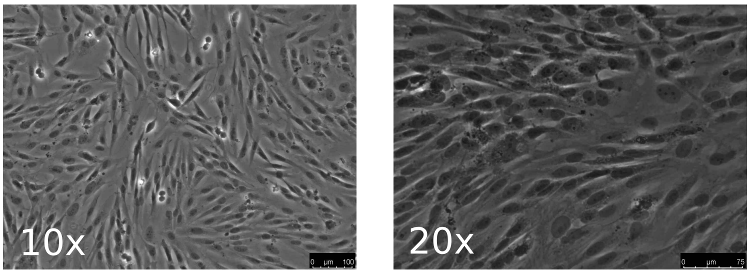


#### Ploidy

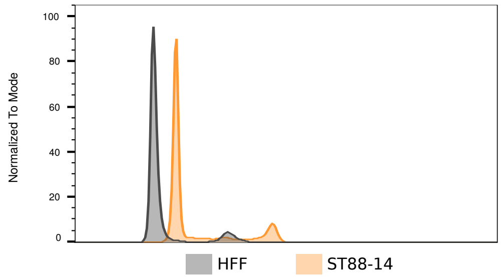


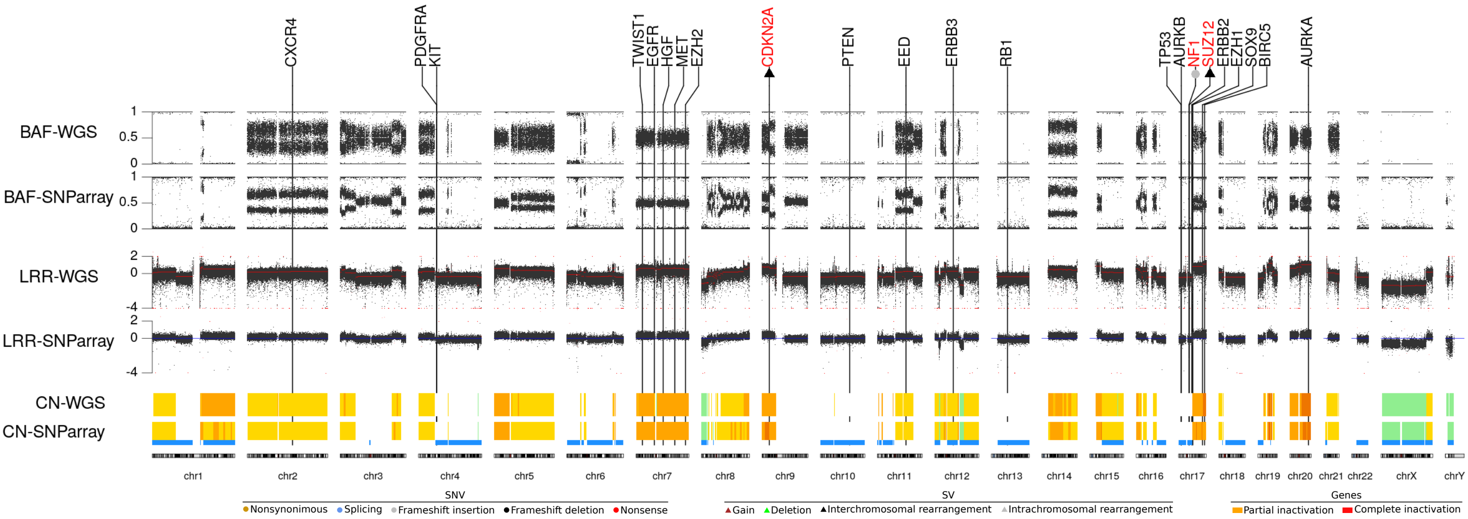


#### Genes

**Tumor Supressor**

- NF1 — c.1649dupT;LOH
- CDKN2A — Complete inactivation by translocations
- SUZ12 — Translocation;LOH
- EED
- TP53
- PTEN
- RB1

**Membrane Receptors**

- EGFR
- ERBB2
- ERBB3
- PDGFRA
- KIT
- MET
- HGF
- CXCR4

**Transcription factors & other**

- BIRC5
- TWIST1
- SOX9
- AURKA
- AURKB

WT, LOH, Homozygous loss, CN Gain, SV, SNV, Inactivation, Highly Gained

#### STR

| | |
|---|---|
| D8S1179 | 14 |
| D21S11 | 29, 32.3 |
| D7S820 | 8 |
| CSF1PO | 9, 12 |
| D3S1358 | 15, 18 |
| TH01 | 9 |
| D13S317 | 12 |
| D16S539 | 13 |
| D2S11338 | 17, 23 |
| D19S433 | 13, 14 |
| vWA | 16 |
| TPOX | 11, 12 |
| D18S51 | 12 |
| AMEL | X, Y |
| D5S818 | 12, 13 |
| FGA | 21 |

#### Methylome class

MPNST

#### Marker expression

SOX9: Positive SOX10: Negative S100B: Negative

Additional alterations:

Additional information:

Aliases: ST88.14; ST 88-14; ST-8814; ST8814; 88-14; NF188-14

Original publication: .

### sNF96.2

RRID:CVCL_K281

NF1
Male, 27yo

#### Phenotype

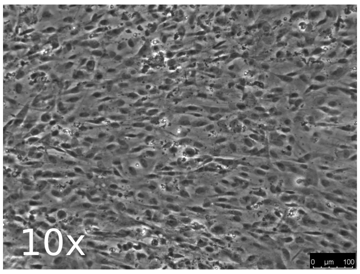


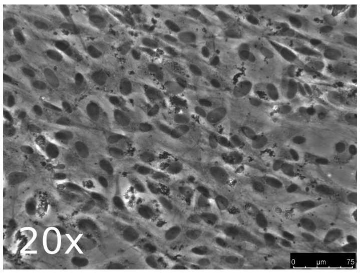


#### Ploidy

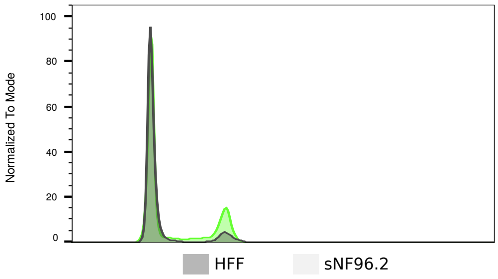


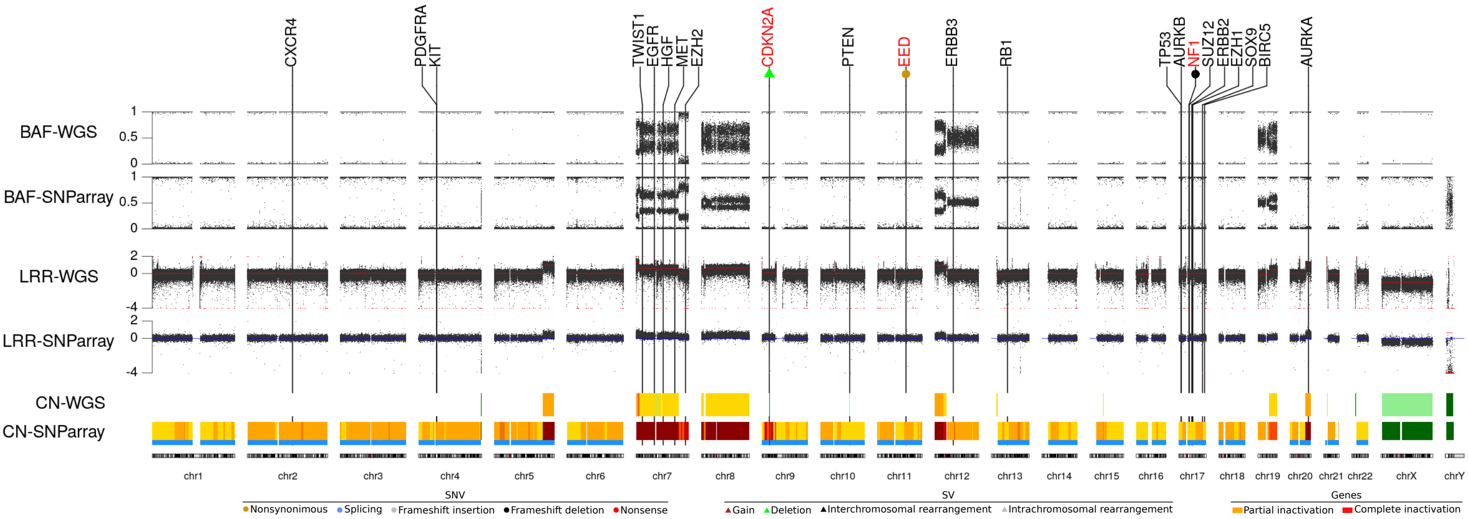


#### Genes

##### Tumor Supressor

- NF1 — c.3683delC;LOH
- CDKN2A — Homozygous deletion
- SUZ12
- EED — c.721A>C;LOH
- TP53
- PTEN
- RB1

##### Membrane Receptors

- EGFR
- ERBB2
- ERBB3
- PDGFRA
- KIT
- MET
- HGF
- CXCR4

##### Transcription factors & other

- BIRC5
- TWIST1
- SOX9
- AURKA
- AURKB

Legend: WT, LOH, Homozygous loss, CN Gain, SV, SNV, Inactivation, Highly Gained

#### STR

| | |
|---|---|
| D8S1179 | 13 |
| D21S11 | 30 |
| D7S820 | 10, 11 |
| CSF1PO | 12 |
| D3S1358 | 16 |
| TH01 | 6 |
| D13S317 | 10 |
| D16S539 | 11 |
| D2S11338 | 17 |
| D19S433 | 13, 14 |
| vWA | 17, 19 |
| TPOX | 11 |
| D18S51 | 16 |
| AMEL | X |
| D5S818 | 11 |
| FGA | 22 |

#### Methylome class

MPNST

#### Marker expression

SOX9: Positive    SOX10: Negative    S100B: Negative

Additional alterations:

Additional information:

Aliases: SNF96.2; sNF96-2

Original publication: Perrin GQ, Li H, Fishbein L, Thomson SA, Hwang MS, Scarborough MT, Yachnis AT, Wallace MR, Mareci TH, Muir D. An orthotopic xenograft model of intraneural NF1 MPNST suggests a potential association between steroid hormones and tumor cell proliferation. Lab Invest. 2007 Nov;87(11):1092-102. https://doi.org/10.1038/labinvest.3700675

# NF90-8

NF1

Female, 17yo

RRID:CVCL_1B47

#### Phenotype

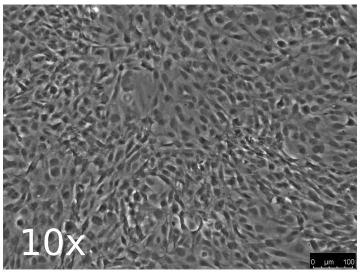


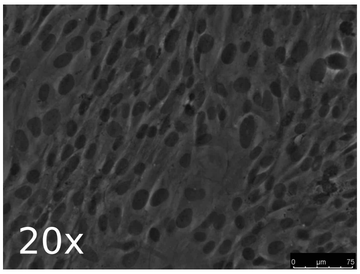


#### Ploidy

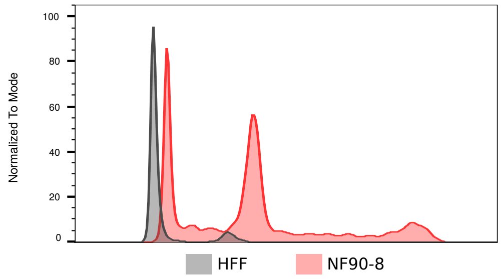


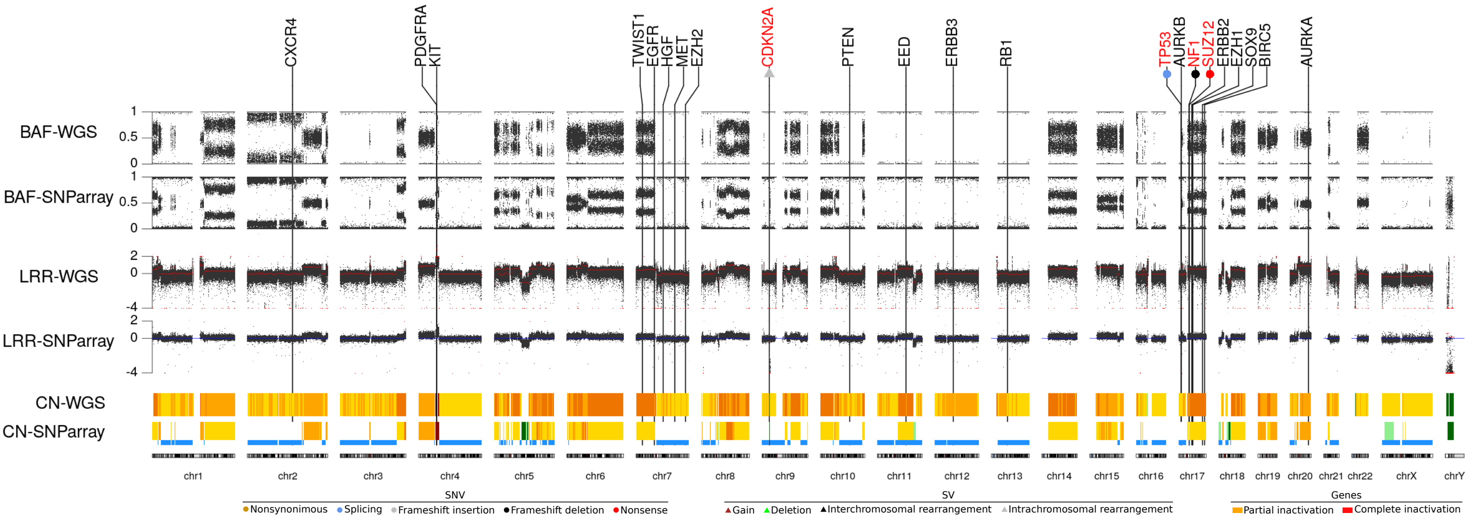


#### Genes

| Tumor Supressor | | Membrane Receptors | Transcription factors & other |
|---|---|---|---|
| NF1 | c.3904_3910del;LOH | EGFR | BIRC5 |
| CDKN2A | Homozygous deletion | ERBB2 | TWIST1 |
| SUZ12 | c.T1290A;LOH | ERBB3 | SOX9 |
| EED | | PDGFRA | AURKA |
| TP53 | c.96+1G>A;LOH | KIT | AURKB |
| PTEN | | MET | |
| RB1 | | HGF | |
| | | CXCR4 | |

WT · LOH · Homozygous loss · CN Gain · SV · SNV · Inactivation · Highly Gained

#### STR

| | |
|---|---|
| D8S1179 | 10, 12 |
| D21S11 | 30 |
| D7S820 | 11 |
| CSF1PO | 12 |
| D3S1358 | 15 |
| TH01 | 6 |
| D13S317 | 8 |
| D16S539 | 9 |
| D2S11338 | 16, 24 |
| D19S433 | 13, 16 |
| vWA | 17 |
| TPOX | 11 |
| D18S51 | 14, 18 |
| AMEL | X |
| D5S818 | 11 |
| FGA | 25 |

#### Methylome class

MPNST

#### Marker expression

SOX9: Positive SOX10: Negative S100B: Negative

Additional alterations:

Additional information:

Aliases: MPNST 90-8; 90-8TL; 90-8-TL; 90-8; NF190-8

Original publication: Legius E, Dierick H, Wu R, Hall BK, Marynen P, Cassiman JJ, Glover TW. TP53 mutations are frequent in malignant NF1 tumors. Genes Chromosomes Cancer. 1994 Aug;10(4):250-5. https://doi.org/10.1002/gcc.2870100405

### NMS-2

RRID:CVCL_4662

NF1

Male, 30yo

#### Phenotype

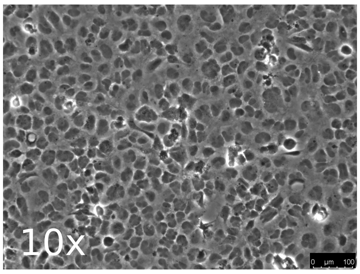


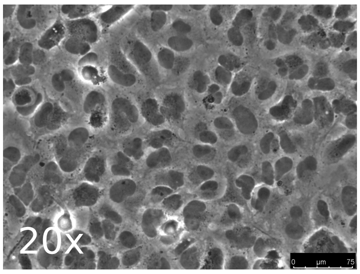


#### Ploidy

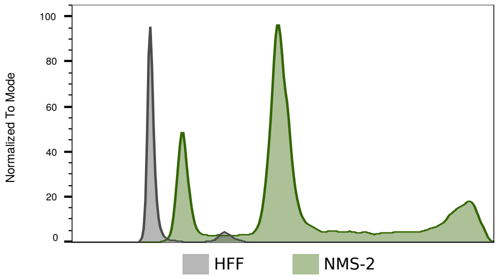


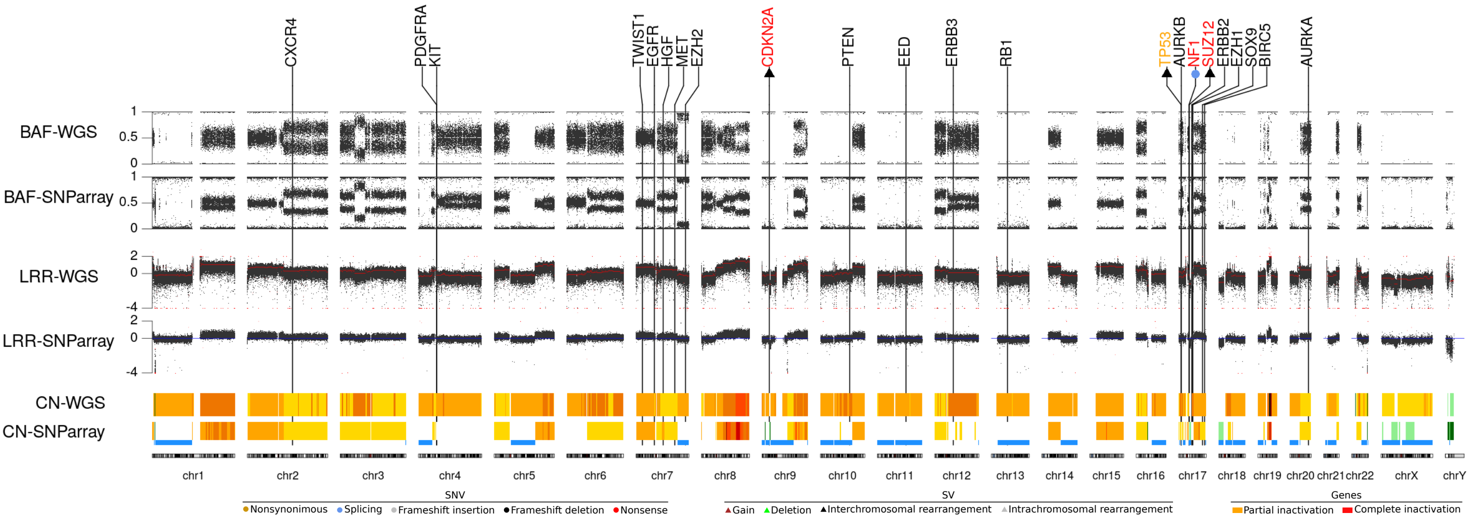


#### Genes

| Tumor Supressor | | Membrane Receptors | Transcription factors & other |
|---|---|---|---|
| NF1 | c.7062+1G>T;LOH | EGFR | BIRC5 |
| CDKN2A | Complete inactivation by translocations | ERBB2 | TWIST1 |
| SUZ12 | Translocation;LOH | ERBB3 | SOX9 |
| EED | | PDGFRA | AURKA |
| TP53 | Translocation | KIT | AURKB |
| PTEN | | MET | |
| RB1 | | HGF | |
| | | CXCR4 | |

WT · LOH · Homozygous loss · CN Gain · SV · SNV · Inactivation · Highly Gained

#### STR

| | |
|---|---|
| D8S1179 | 12, 13 |
| D21S11 | 30 |
| D7S820 | 9, 12 |
| CSF1PO | 12 |
| D3S1358 | 15, 16 |
| TH01 | 6 |
| D13S317 | 12 |
| D16S539 | 9 |
| D2S11338 | 18, 19 |
| D19S433 | 13.2,14.2 |
| vWA | 14, 19 |
| TPOX | 8, 9 |
| D18S51 | 17 |
| AMEL | X, Y |
| D5S818 | 12, 13 |
| FGA | 21, 22 |

#### Methylome class

MPNST

#### Marker expression

SOX9: Positive SOX10: Negative S100B: Negative

Additional alterations:

Additional information:

Aliases:

Original publication: Imaizumi S, Motoyama T, Ogose A, Hotta T, Takahashi HE. Characterization and chemosensitivity of two human malignant peripheral nerve sheath tumour cell lines derived from a patient with neurofibromatosis type 1. Virchows Arch. 1998 Nov;433(5):435-41. https://doi.org/10.1007/s004280050271

### STS-26T

RRID:CVCL_8917

**Sporadic**

Female, 51yo

#### Phenotype

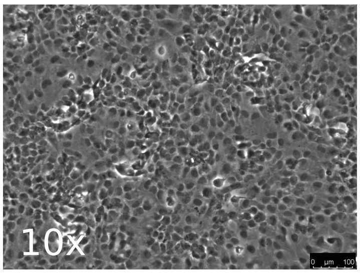


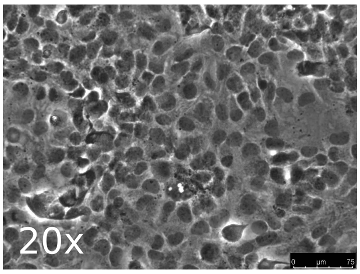


#### Ploidy

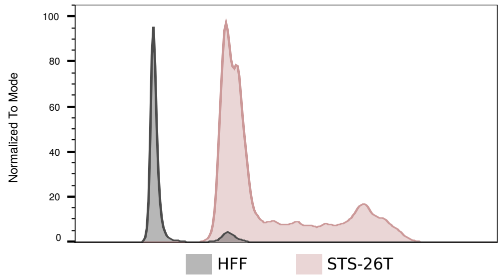


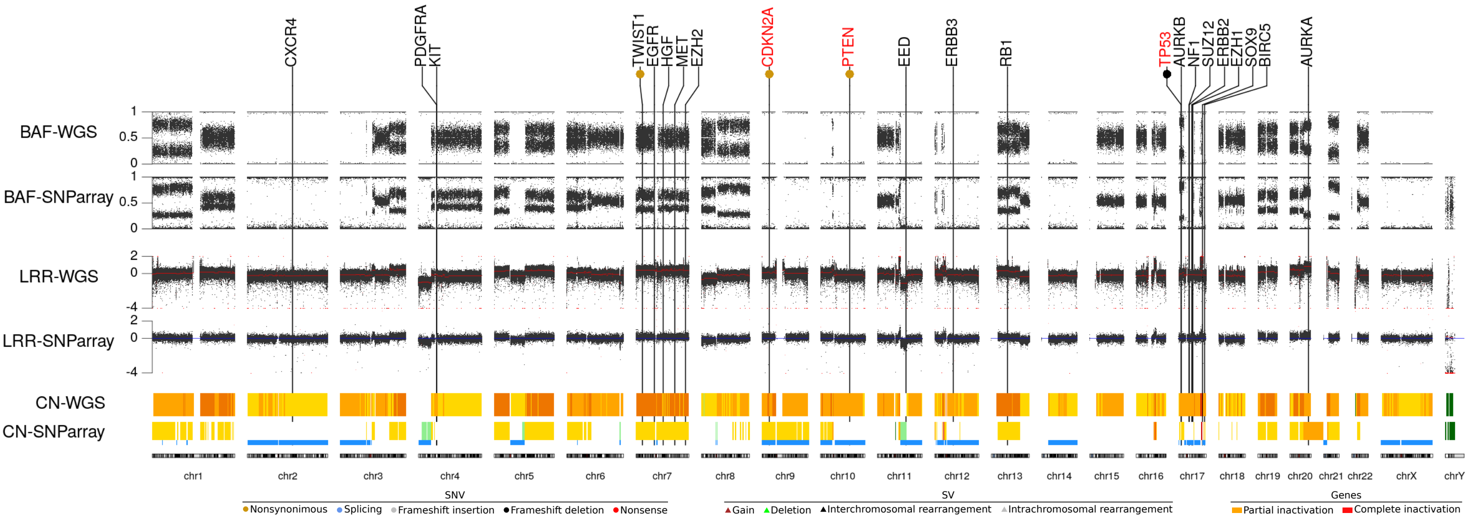


#### Genes

##### Tumor Supressor

- NF1
- CDKN2A c.C341T;LOH
- SUZ12
- EED
- TP53 c.232_241del;LOH
- PTEN c.1021TT>AA;LOH
- RB1

##### Oncogenes

- EGFR
- ERBB2
- ERBB3
- PDGFRA
- KIT
- MET
- HGF
- CXCR4

##### Transcription factors & other

- BIRC5
- TWIST1 c.446T>C
- SOX9
- AURKA
- AURKB

Legend: WT; LOH; Homozygous loss; CN Gain; SV; SNV; Inactivation; Highly Gained

#### STR

| | |
|---|---|
| D8S1179 | 13, 14 |
| D21S11 | 30, 31 |
| D7S820 | 8, 11 |
| CSF1PO | 10, 13 |
| D3S1358 | 14 |
| TH01 | 6, 9.3 |
| D13S317 | 9, 10 |
| D16S539 | 12, 13 |
| D2S11338 | 20 |
| D19S433 | 14 |
| vWA | 17 |
| TPOX | 8 |
| D18S51 | 17, 18 |
| AMEL | X |
| D5S818 | 11, 12 |
| FGA | 22, 23 |

#### Methylome class

Melanoma

#### Marker expression

SOX9: Positive SOX10: Negative S100B: Negative

Additional alterations: BRAF:p.V600E

Additional information: High mutation frequency; Important contribution of COSMIC mutational signature 7 (predominantly seen in skin cancers)

Aliases: STS26T; STS26

Original publication: Dahlberg WK, Little JB, Fletcher JA, Suit HD, Okunieff P. Radiosensitivity in vitro of human soft tissue sarcoma cell lines and skin fibroblasts derived from the same patients. Int J Radiat Biol. 1993 Feb;63(2):191-8. https://doi.org/10.1080/09553009314550251.

### HS-Sch-2

RRID:CVCL_8718

Sporadic

Female, 54yo

#### Phenotype

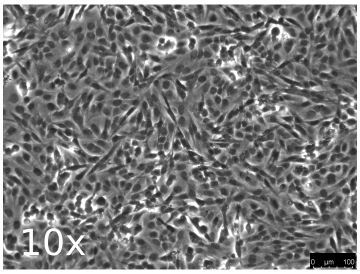


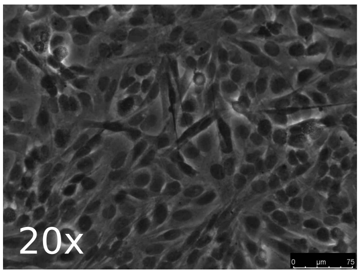


#### Ploidy

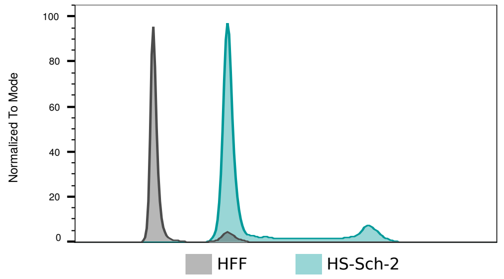


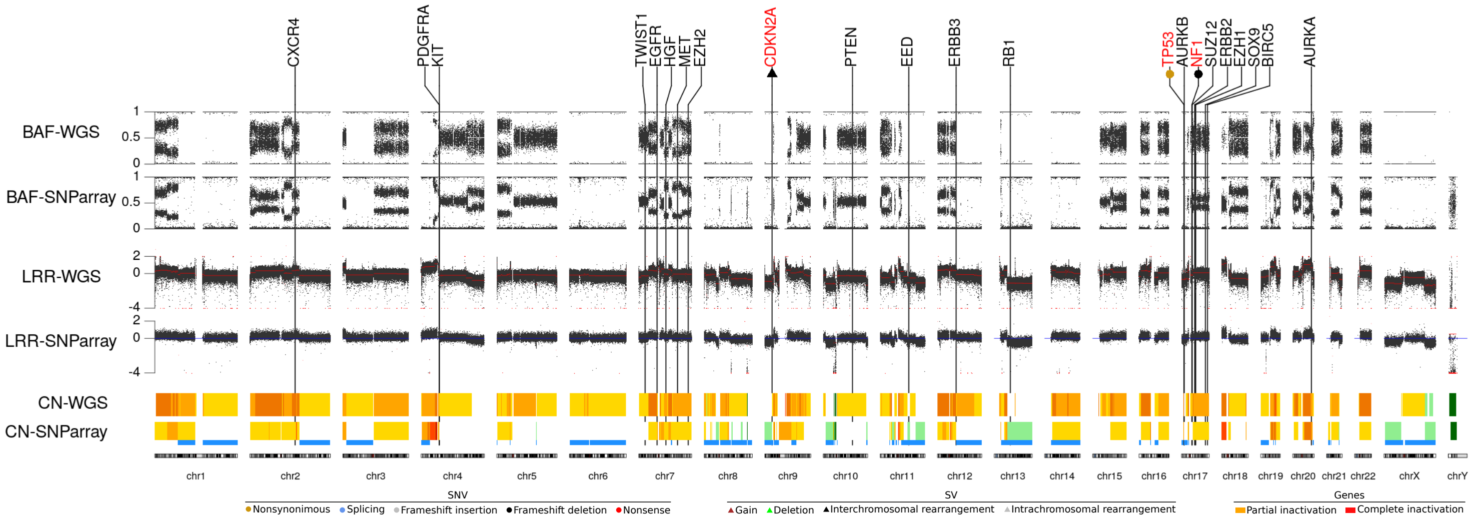


#### Genes

##### Tumor Supressor

- NF1 — c.270_288del;c.3113+1G>A
- CDKN2A — Translocation;LOH
- SUZ12
- EED
- TP53 — c.818G>A;LOH
- PTEN
- RB1

##### Membrane Receptors

- EGFR
- ERBB2
- ERBB3
- PDGFRA
- KIT
- MET
- HGF
- CXCR4

##### Transcription factors & other

- BIRC5
- TWIST1
- SOX9
- AURKA
- AURKB

Legend: WT, LOH, Homozygous loss, CN Gain, SV, SNV, Inactivation, Highly Gained

#### STR

| | |
|---|---|
| D8S1179 | 14 |
| D21S11 | 29, 30 |
| D7S820 | 8, 11 |
| CSF1PO | 11, 13 |
| D3S1358 | 18 |
| TH01 | 6, 7 |
| D13S317 | 11 |
| D16S539 | 11, 12 |
| D2S11338 | 20 |
| D19S433 | 14.2 |
| vWA | 15, 16 |
| TPOX | 8 |
| D18S51 | 13, 14 |
| AMEL | X |
| D5S818 | 13 |
| FGA | 23, 26 |

#### Methylome class

Melanoma

#### Marker expression

SOX9: Positive SOX10: Positive S100B: Positive

Additional alterations:

Additional information:

Aliases: HS-sch-2; HSSCH2

Original publication: Sonobe H, Takeuchi T, Furihata M, Taguchi T, Kawai A, Ohjimi Y, Iwasaki H, Kaneko Y, Ohtsuki Y. A new human malignant peripheral nerve sheath tumour-cell line, HS-sch-2, harbouring p53 point mutation. Int J Oncol. 2000 Aug; 17(2):347-52. https://doi.org/10.3892/ijo.17.2.347

### HS-PSS

RRID:CVCL_8717

Sporadic

Male

#### Phenotype

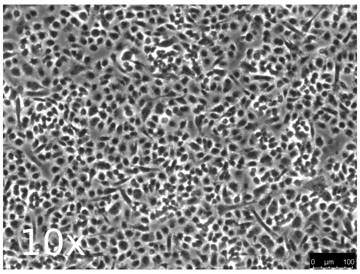


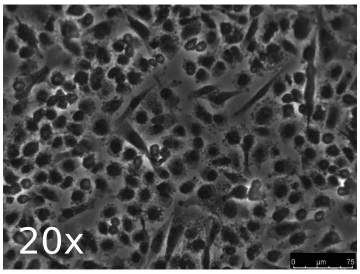


#### Ploidy

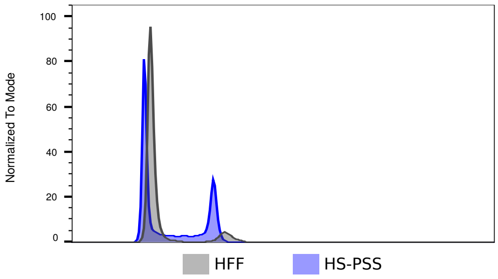


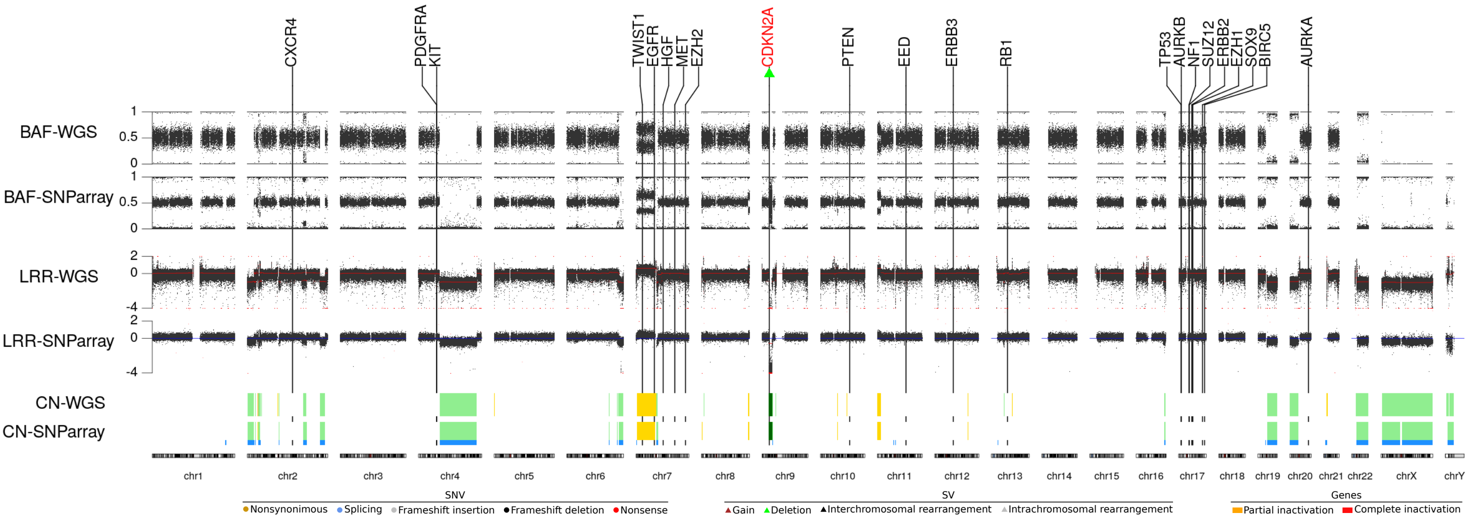


#### Genes

| Tumor Supressor | | Membrane Receptors | | Transcription factors & other | |
|---|---|---|---|---|---|
| NF1 | | EGFR | CN Gain | BIRC5 | |
| CDKN2A | Homozygous deletion | ERBB2 | | TWIST1 | CN Gain |
| SUZ12 | | ERBB3 | | SOX9 | |
| EED | | PDGFRA | | AURKA | |
| TP53 | | KIT | | AURKB | |
| PTEN | | MET | | | |
| RB1 | | HGF | | | |
| | | CXCR4 | | | |

WT · LOH · Homozygous loss · CN Gain · SV · SNV · Inactivation · Highly Gained

#### STR

| | |
|---|---|
| D8S1179 | 14, 15 |
| D21S11 | 28, 32 |
| D7S820 | 10, 12 |
| CSF1PO | 10, 12 |
| D3S1358 | 16 |
| TH01 | 6, 9 |
| D13S317 | 8, 10 |
| D16S539 | 10 |
| D2S11338 | 19 |
| D19S433 | 13 |
| vWA | 14, 15 |
| TPOX | 8 |
| D18S51 | 14, 15 |
| AMEL | X, Y |
| D5S818 | 9, 11 |
| FGA | 22 |

#### Methylome class

MPNST-like

#### Marker expression

SOX9: Positive SOX10: Positive S100B: Negative

Additional alterations: Intrachromosomal rearrangement in chromosome 2 creating an EML4-ALK fusion gene

Additional information:

Aliases: Hs-PSS; HSPSS

Original publication: .

### Supplementary Document 4: Copy number profile for each MPNST cell line, chromosome by chromosome.

S462

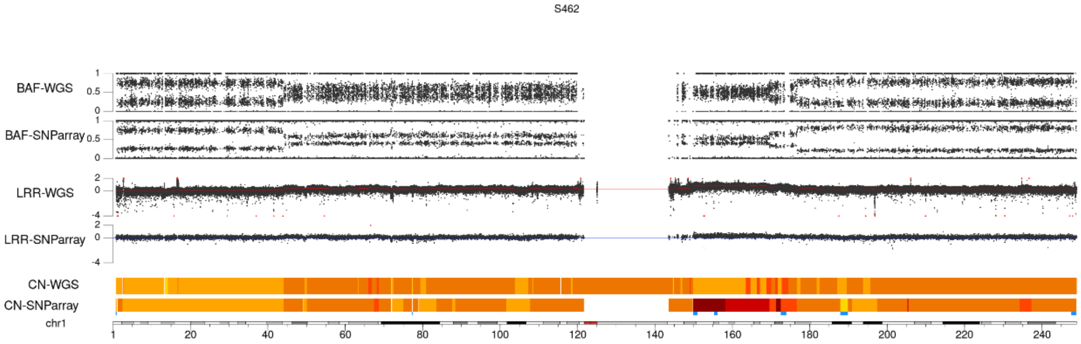


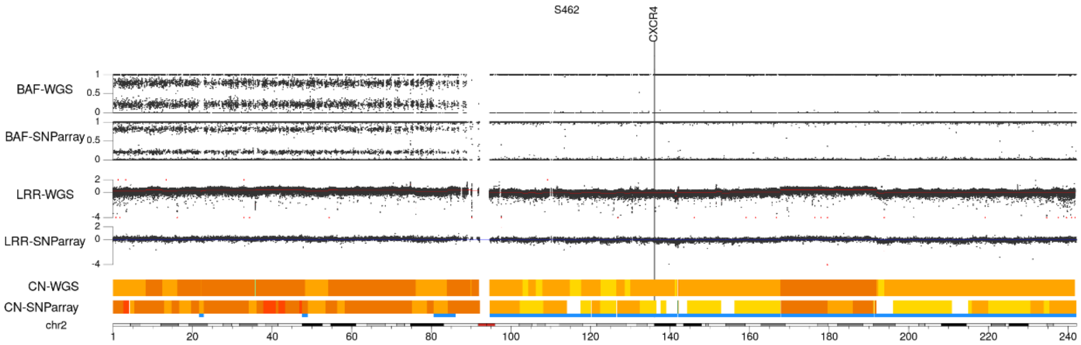


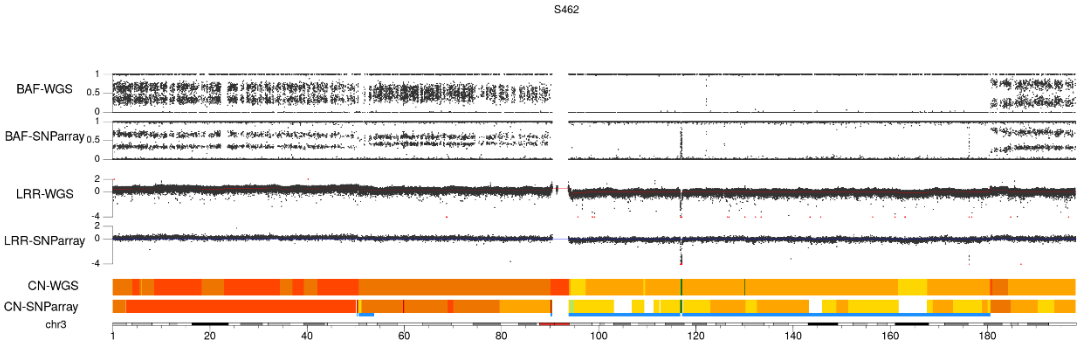


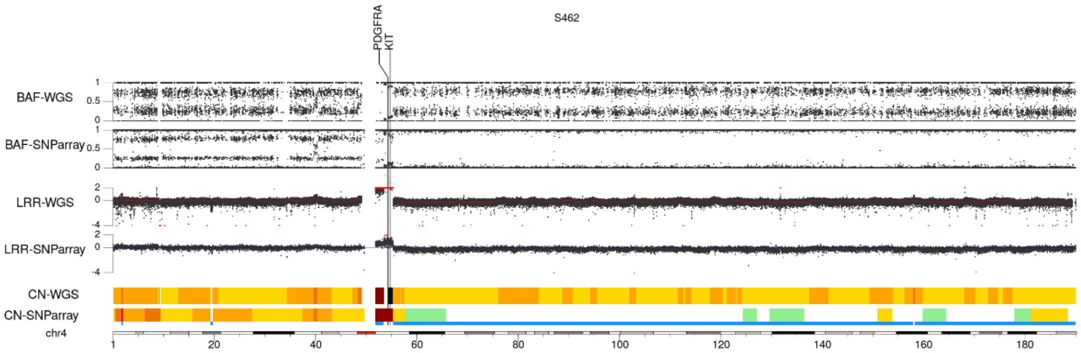

S462

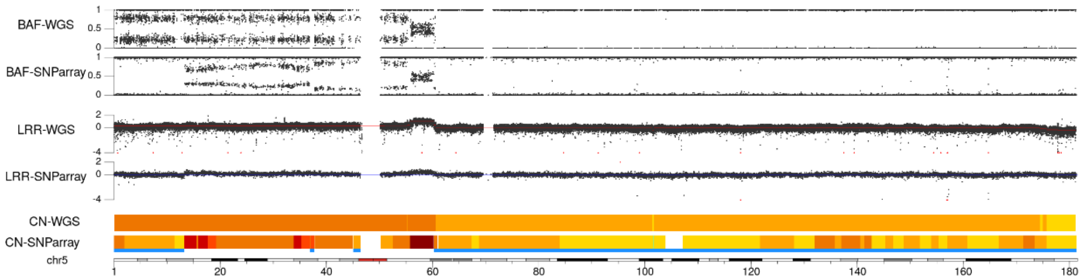


S462

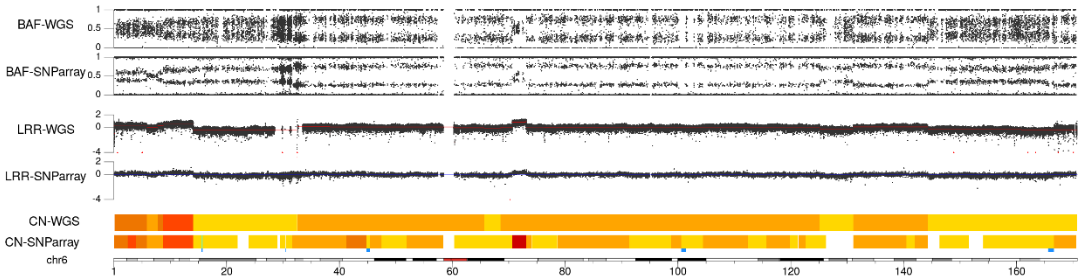


S462

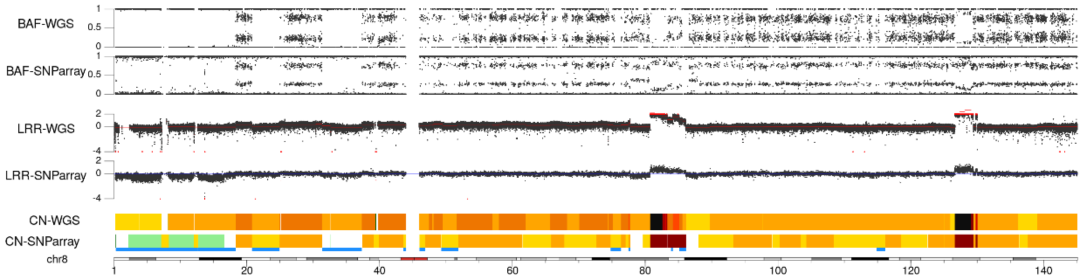


S462

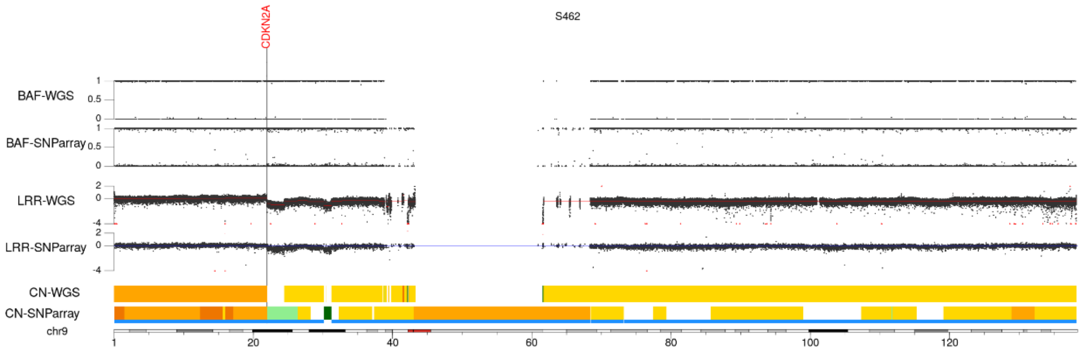


S462

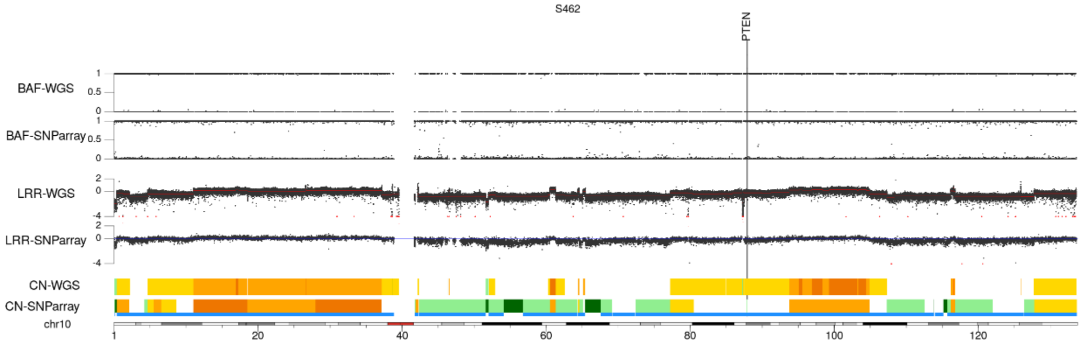

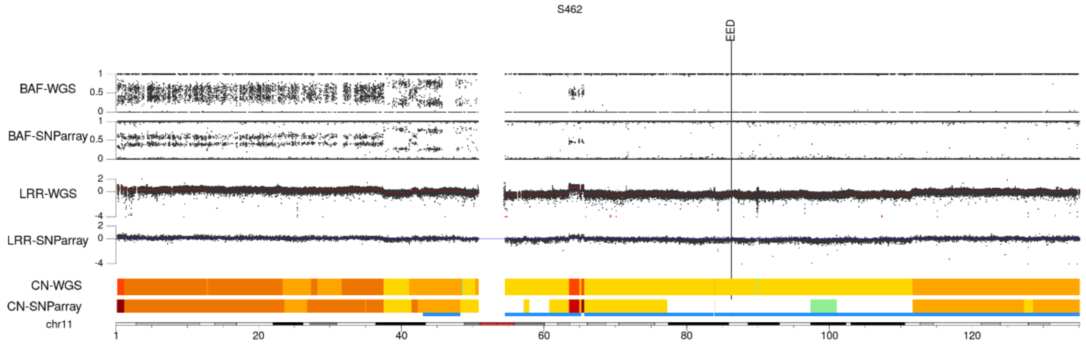
S462
EED
BAF-WGS
BAF-SNParray
LRR-WGS
LRR-SNParray
CN-WGS
CN-SNParray
chr11
1
20
40
60
80
100
120

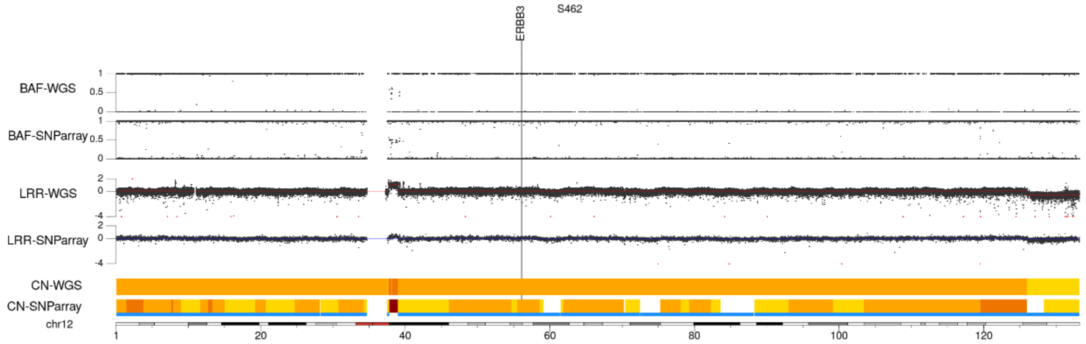
S462
ERBB3
BAF-WGS
BAF-SNParray
LRR-WGS
LRR-SNParray
CN-WGS
CN-SNParray
chr12
1
20
40
60
80
100
120

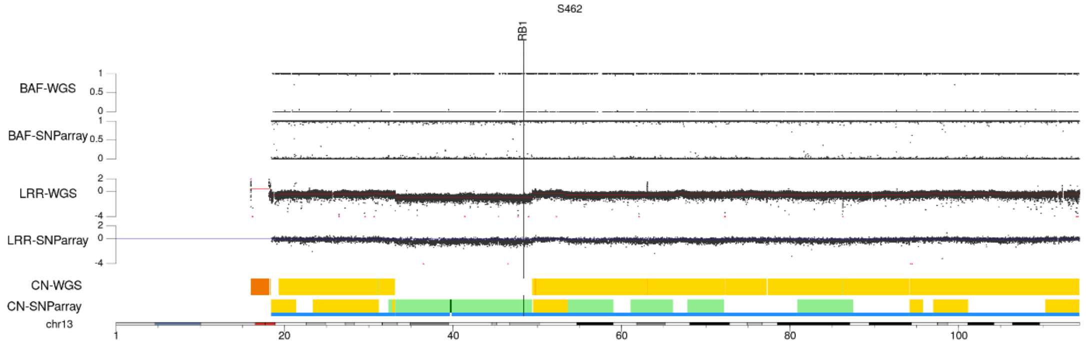
S462
RB1
BAF-WGS
BAF-SNParray
LRR-WGS
LRR-SNParray
CN-WGS
CN-SNParray
chr13
1
20
40
60
80
100

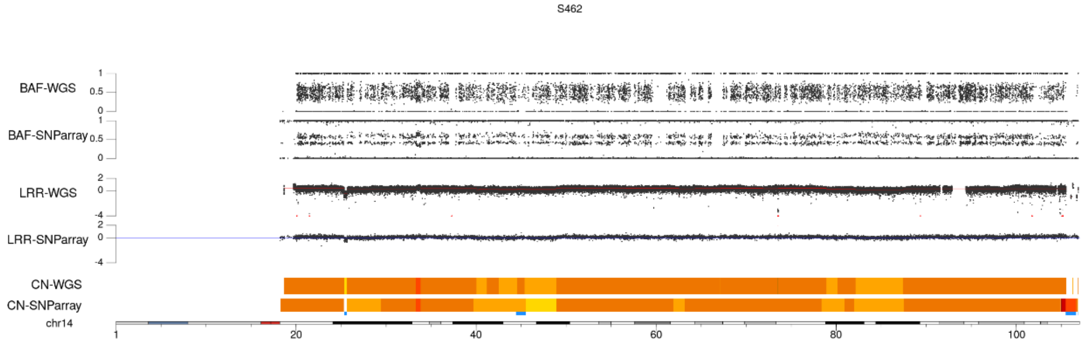
S462
BAF-WGS
BAF-SNParray
LRR-WGS
LRR-SNParray
CN-WGS
CN-SNParray
chr14
1
20
40
60
80
100

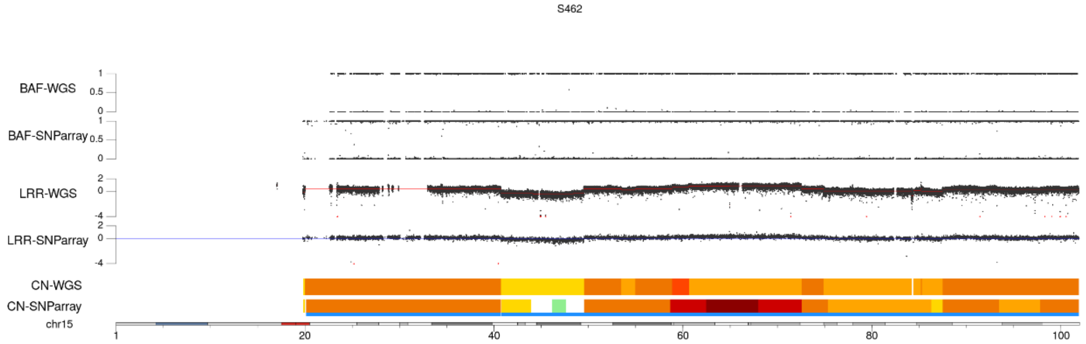
S462
BAF-WGS
BAF-SNParray
LRR-WGS
LRR-SNParray
CN-WGS
CN-SNParray
chr15
1
20
40
60
80
100

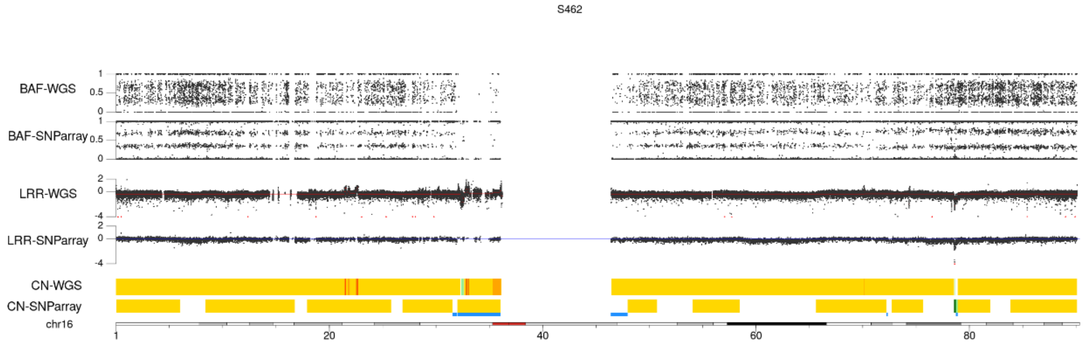
S462
BAF-WGS
BAF-SNParray
LRR-WGS
LRR-SNParray
CN-WGS
CN-SNParray
chr16
1
20
40
60
80

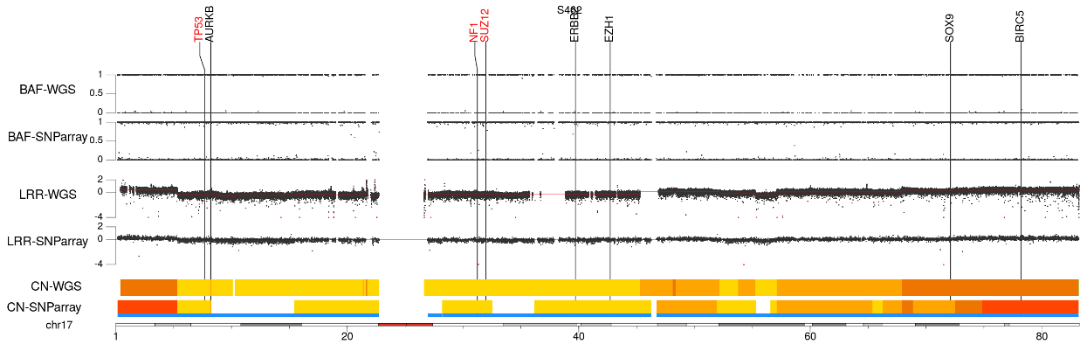
S462
TP53
AURKB
NF1
SUZ12
ERBB2
EZH1
SOX9
BIRC5
BAF-WGS
BAF-SNParray
LRR-WGS
LRR-SNParray
CN-WGS
CN-SNParray
chr17
1
20
40
60
80

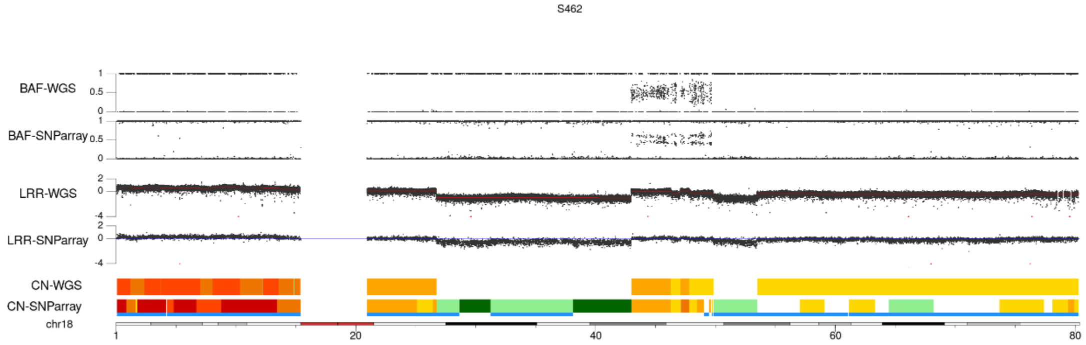
S462
BAF-WGS
BAF-SNParray
LRR-WGS
LRR-SNParray
CN-WGS
CN-SNParray
chr18
1
20
40
60
80

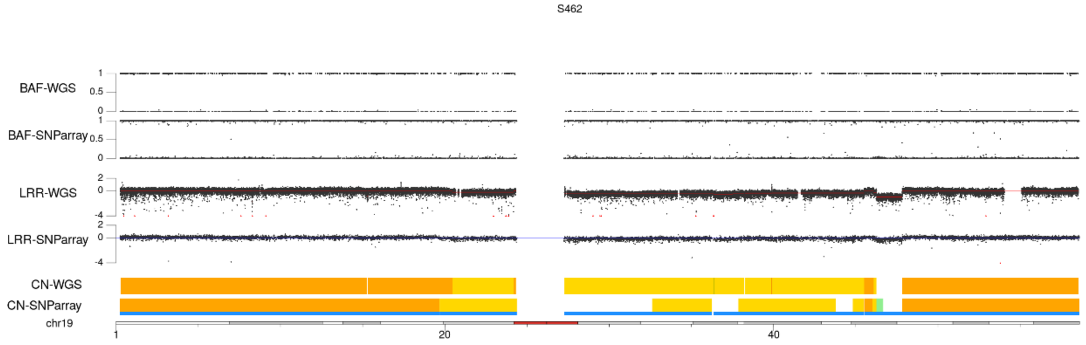
S462
BAF-WGS
BAF-SNParray
LRR-WGS
LRR-SNParray
CN-WGS
CN-SNParray
chr19
1
20
40

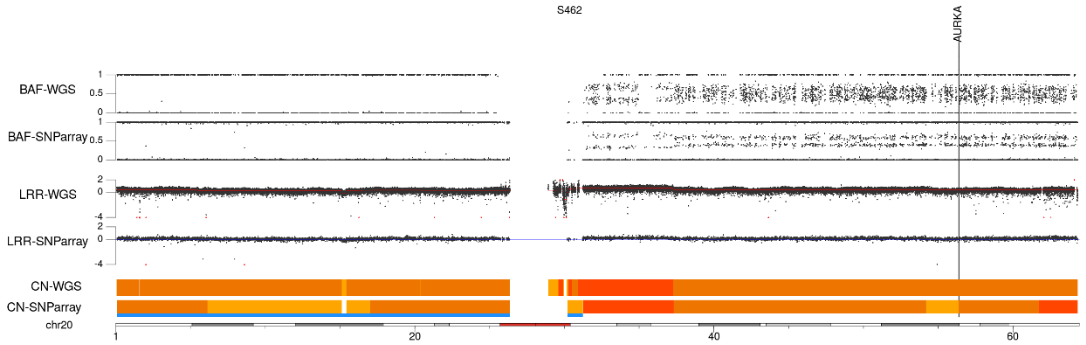
S462
AURKA
BAF-WGS
BAF-SNParray
LRR-WGS
LRR-SNParray
CN-WGS
CN-SNParray
chr20
1
20
40
60

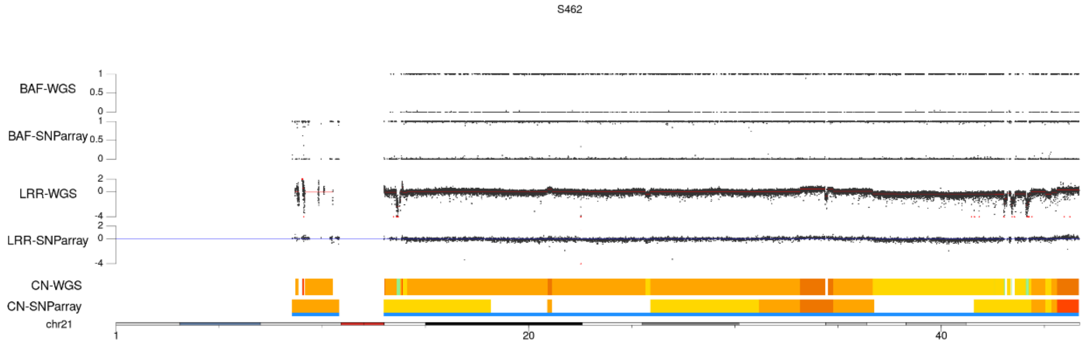
S462
BAF-WGS
BAF-SNParray
LRR-WGS
LRR-SNParray
CN-WGS
CN-SNParray
chr21
1
20
40

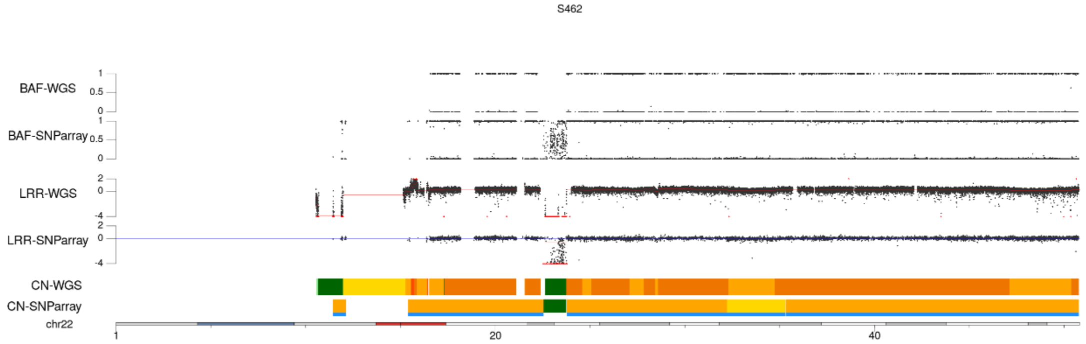
S462
BAF-WGS
BAF-SNParray
LRR-WGS
LRR-SNParray
CN-WGS
CN-SNParray
chr22
1
20
40

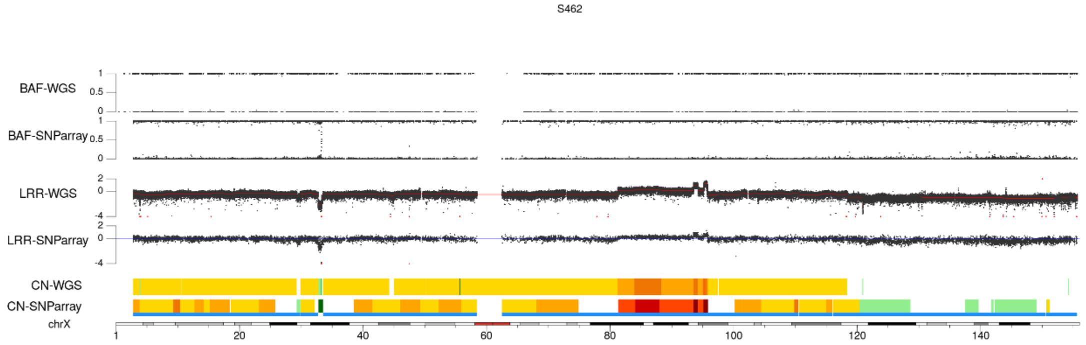
S462
BAF-WGS
BAF-SNParray
LRR-WGS
LRR-SNParray
CN-WGS
CN-SNParray
chrX
1
20
40
60
80
100
120
140

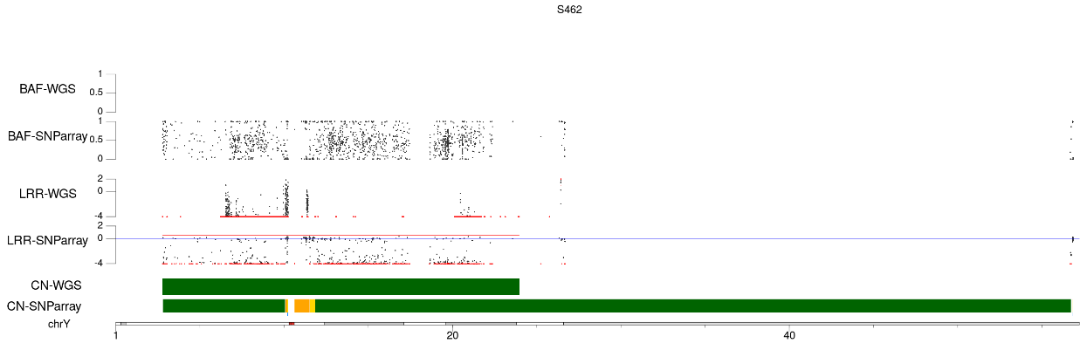
S462
BAF-WGS
BAF-SNParray
LRR-WGS
LRR-SNParray
CN-WGS
CN-SNParray
chrY
1
20
40

# ST88-14

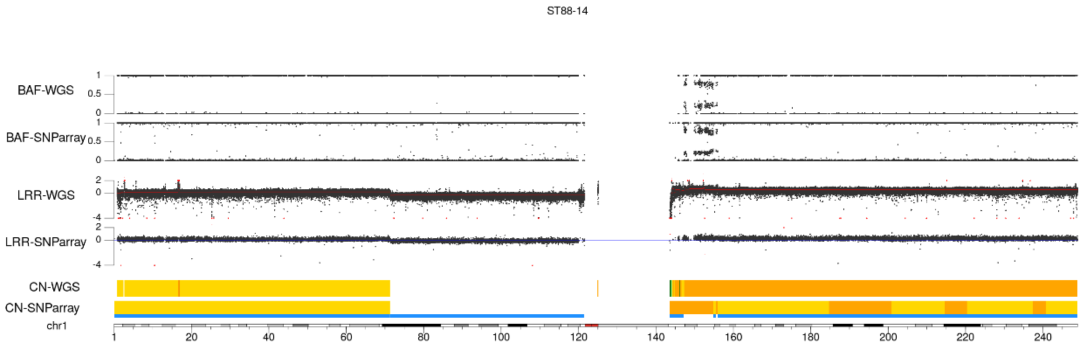
ST88-14
BAF-WGS
BAF-SNParray
LRR-WGS
LRR-SNParray
CN-WGS
CN-SNParray
chr1

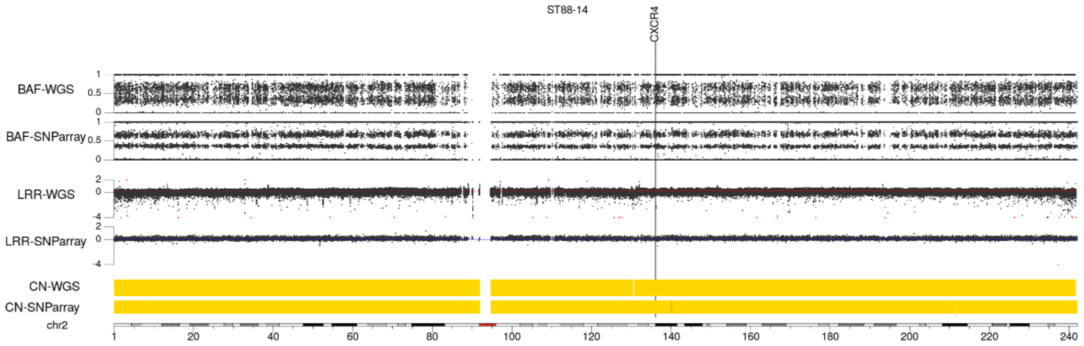
ST88-14
CXCR4
BAF-WGS
BAF-SNParray
LRR-WGS
LRR-SNParray
CN-WGS
CN-SNParray
chr2

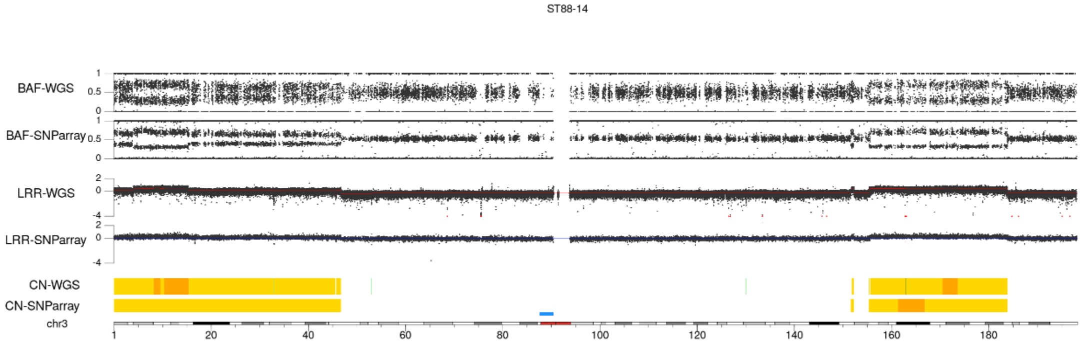
ST88-14
BAF-WGS
BAF-SNParray
LRR-WGS
LRR-SNParray
CN-WGS
CN-SNParray
chr3

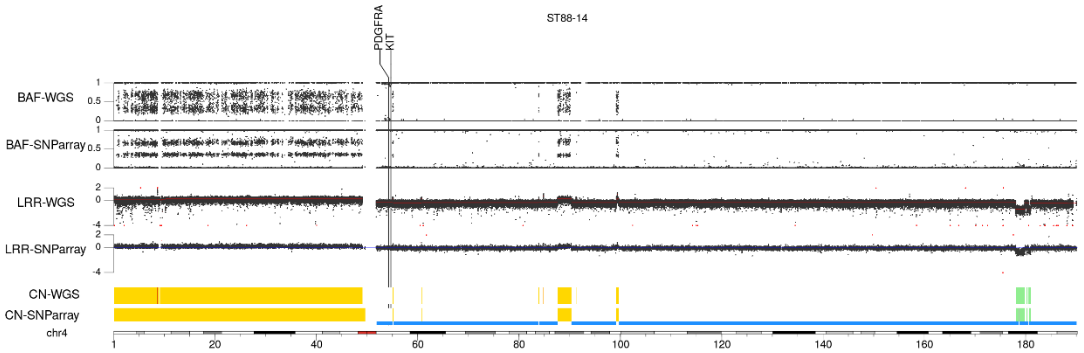
ST88-14
PDGFRA
KIT
BAF-WGS
BAF-SNParray
LRR-WGS
LRR-SNParray
CN-WGS
CN-SNParray
chr4

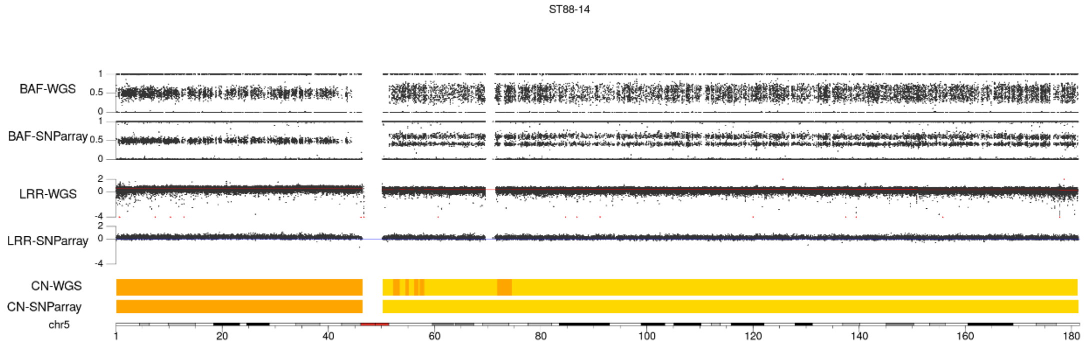
ST88-14
BAF-WGS
BAF-SNParray
LRR-WGS
LRR-SNParray
CN-WGS
CN-SNParray
chr5
1 20 40 60 80 100 120 140 160 180

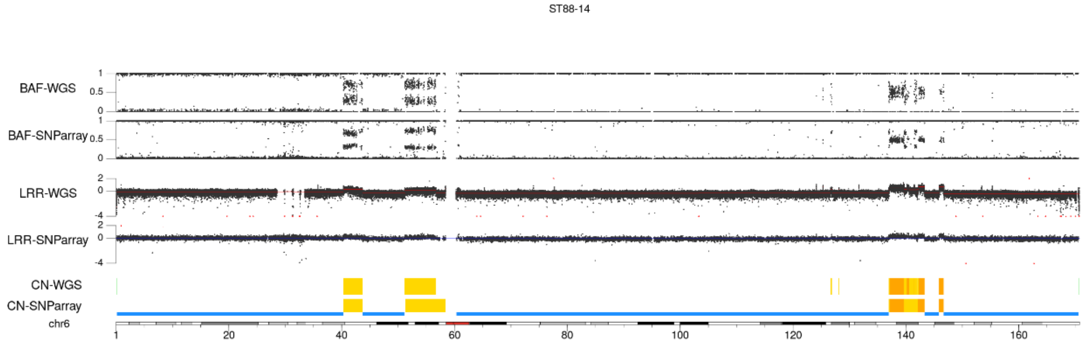
ST88-14
BAF-WGS
BAF-SNParray
LRR-WGS
LRR-SNParray
CN-WGS
CN-SNParray
chr6
1 20 40 60 80 100 120 140 160

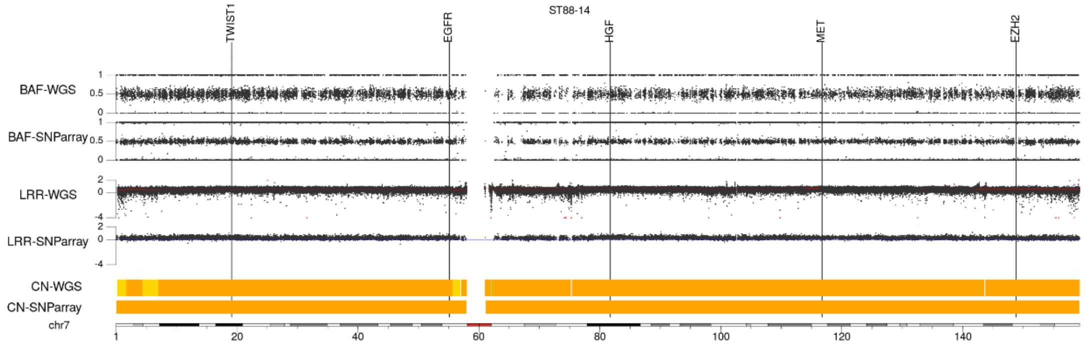
TWIST1
EGFR
HGF
MET
EZH2
ST88-14
BAF-WGS
BAF-SNParray
LRR-WGS
LRR-SNParray
CN-WGS
CN-SNParray
chr7
1 20 40 60 80 100 120 140

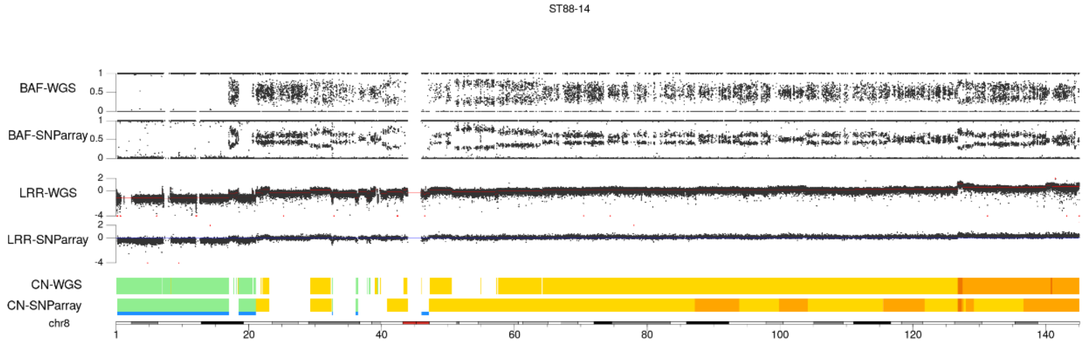
ST88-14
BAF-WGS
BAF-SNParray
LRR-WGS
LRR-SNParray
CN-WGS
CN-SNParray
chr8
1 20 40 60 80 100 120 140

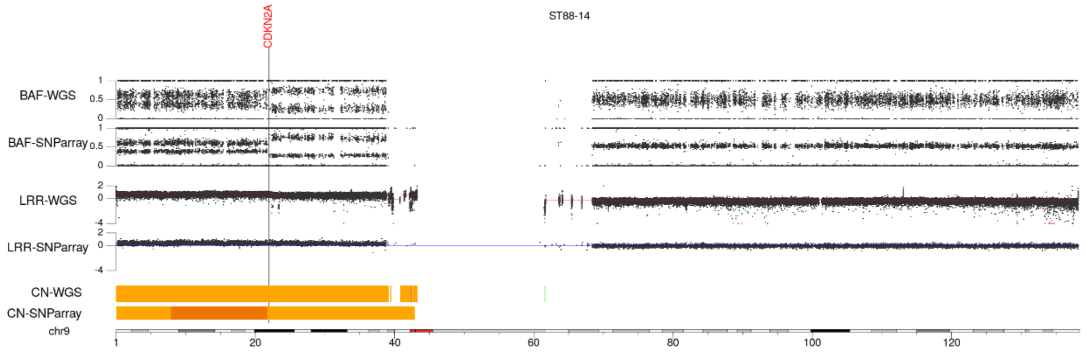
ST88-14
CDKN2A
BAF-WGS
BAF-SNParray
LRR-WGS
LRR-SNParray
CN-WGS
CN-SNParray
chr9
1 20 40 60 80 100 120

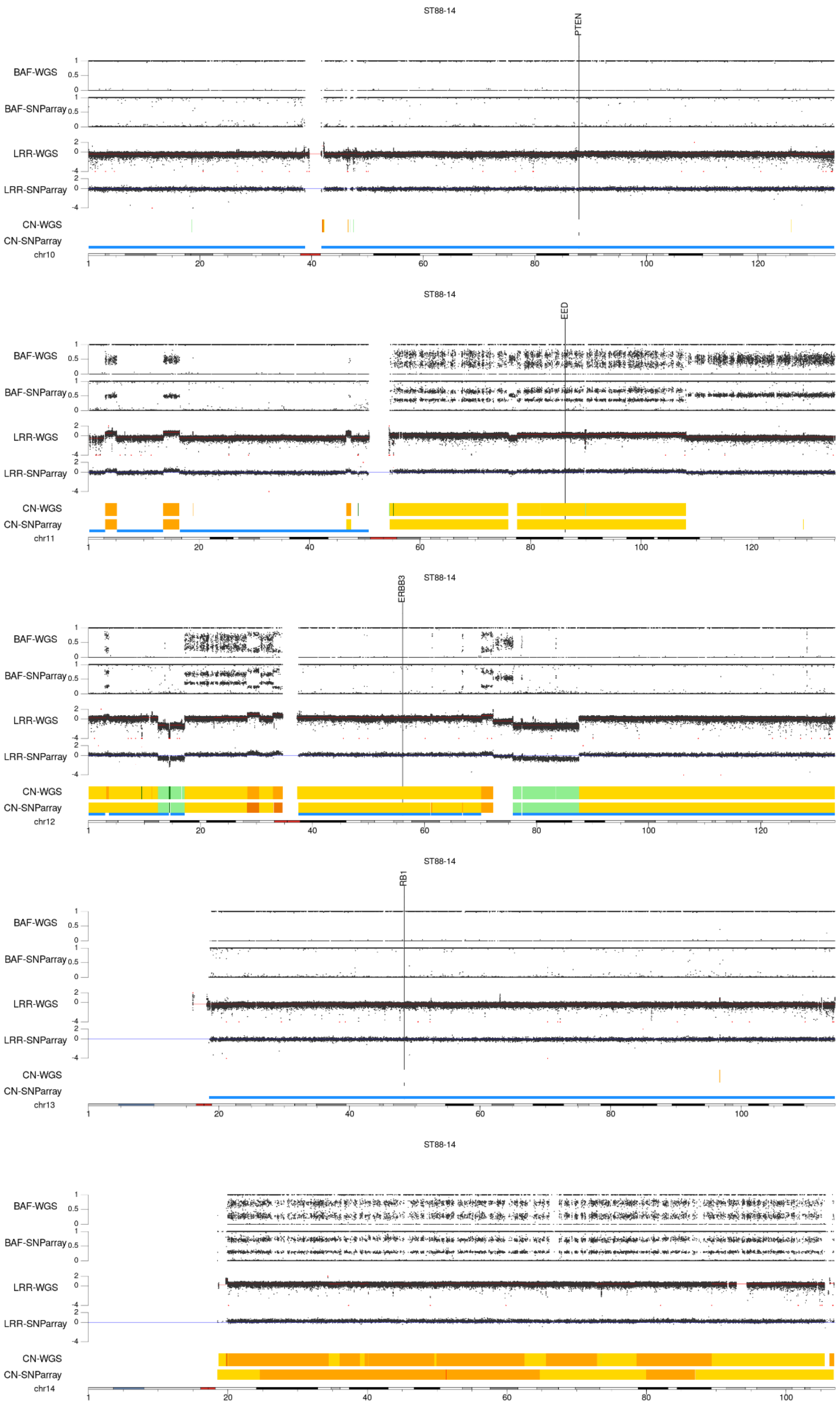
ST88-14
PTEN
BAF-WGS
BAF-SNParray
LRR-WGS
LRR-SNParray
CN-WGS
CN-SNParray
chr10
ST88-14
EED
chr11
ST88-14
ERBB3
chr12
ST88-14
RB1
chr13
ST88-14
chr14

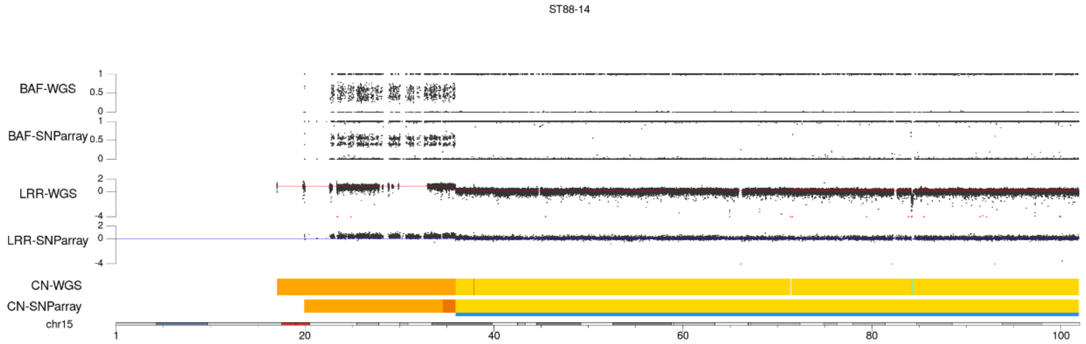
ST88-14
BAF-WGS
BAF-SNParray
LRR-WGS
LRR-SNParray
CN-WGS
CN-SNParray
chr15
1
20
40
60
80
100

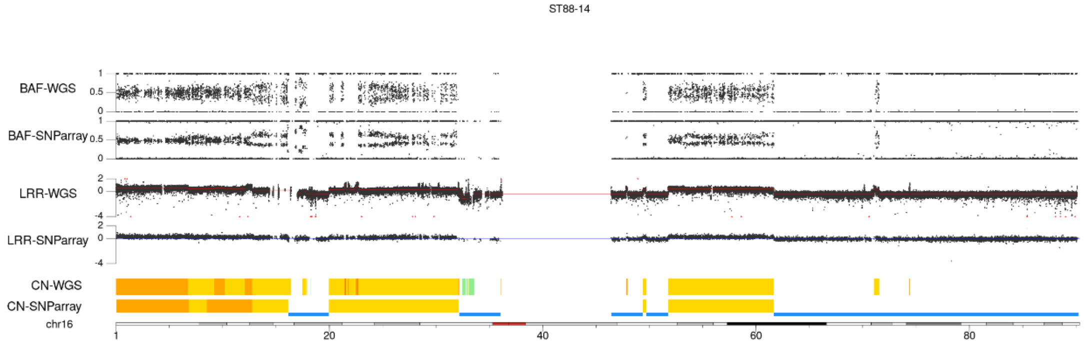
ST88-14
BAF-WGS
BAF-SNParray
LRR-WGS
LRR-SNParray
CN-WGS
CN-SNParray
chr16
1
20
40
60
80

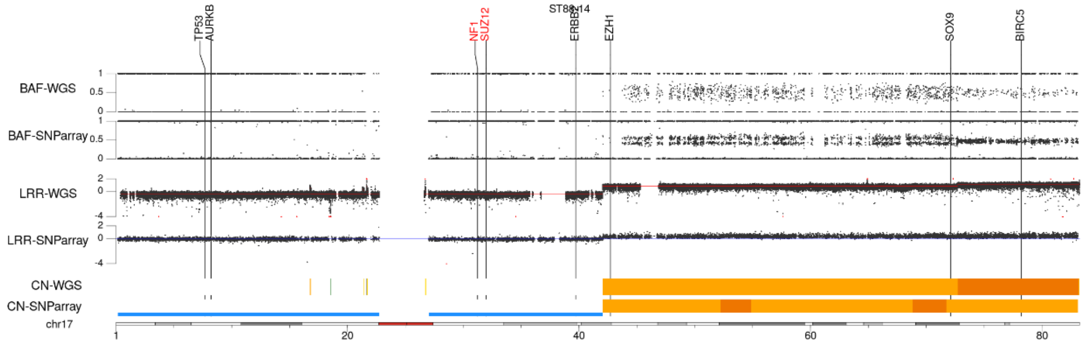
ST88-14
TP53
AURKB
NF1
SUZ12
ERBB2
EZH1
SOX9
BIRC5
BAF-WGS
BAF-SNParray
LRR-WGS
LRR-SNParray
CN-WGS
CN-SNParray
chr17
1
20
40
60
80

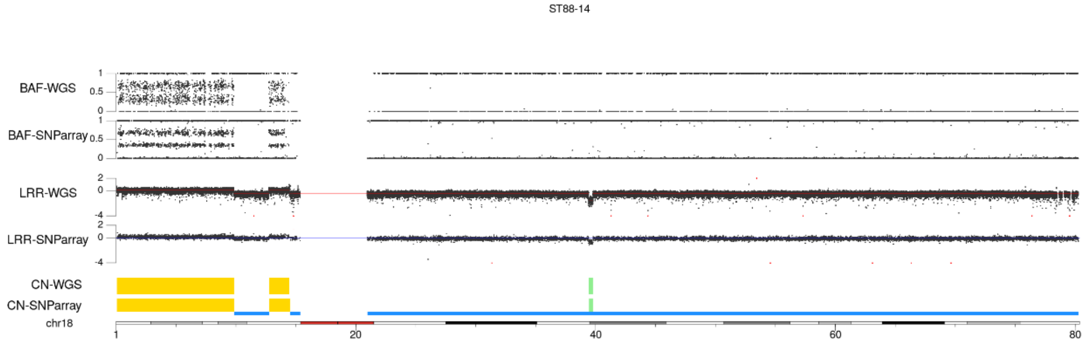
ST88-14
BAF-WGS
BAF-SNParray
LRR-WGS
LRR-SNParray
CN-WGS
CN-SNParray
chr18
1
20
40
60
80

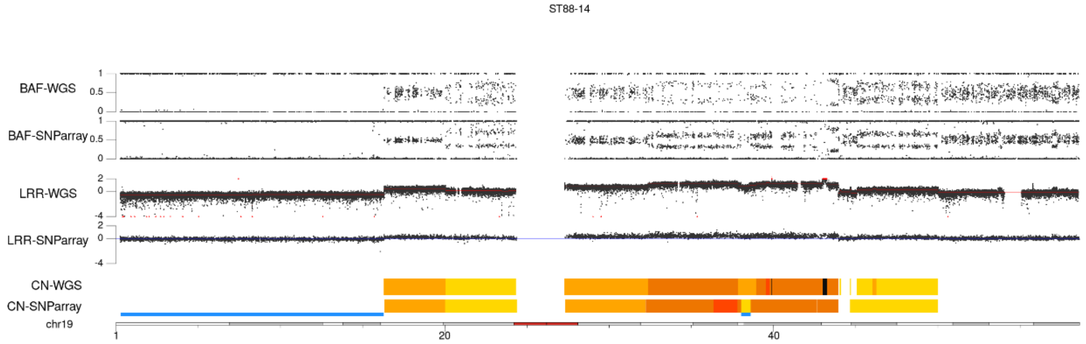
ST88-14
BAF-WGS
BAF-SNParray
LRR-WGS
LRR-SNParray
CN-WGS
CN-SNParray
chr19
1
20
40

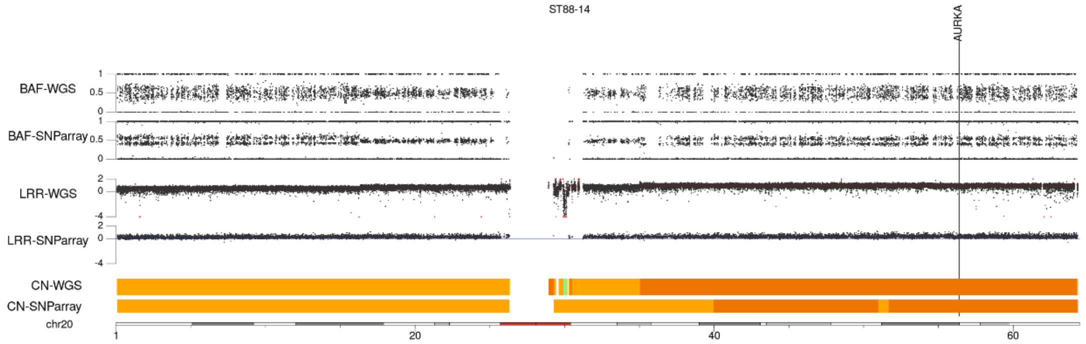
ST88-14
AURKA
BAF-WGS
BAF-SNParray
LRR-WGS
LRR-SNParray
CN-WGS
CN-SNParray
chr20
1
20
40
60

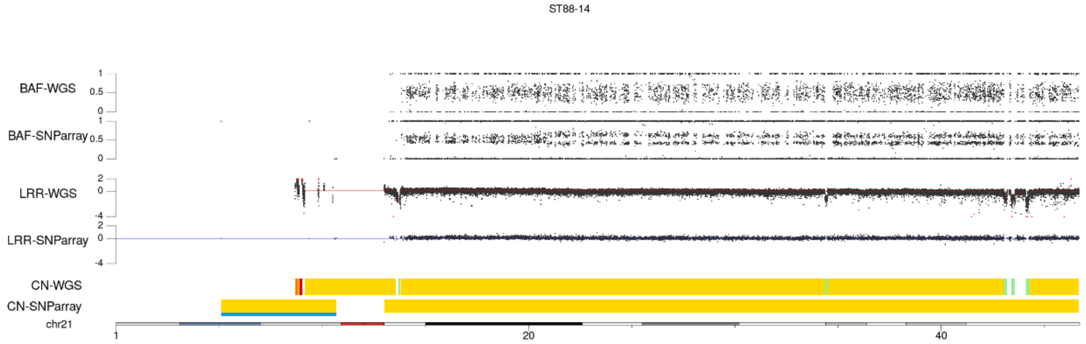
ST88-14
BAF-WGS
BAF-SNParray
LRR-WGS
LRR-SNParray
CN-WGS
CN-SNParray
chr21
1
20
40

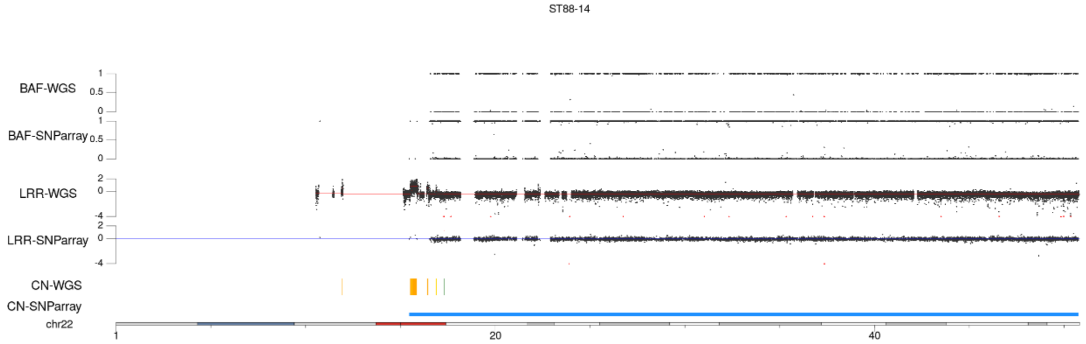
ST88-14
BAF-WGS
BAF-SNParray
LRR-WGS
LRR-SNParray
CN-WGS
CN-SNParray
chr22
1
20
40

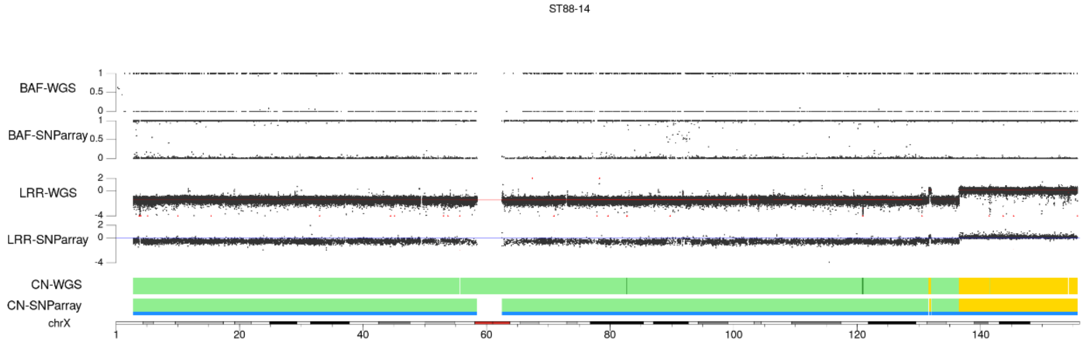
ST88-14
BAF-WGS
BAF-SNParray
LRR-WGS
LRR-SNParray
CN-WGS
CN-SNParray
chrX
1
20
40
60
80
100
120
140

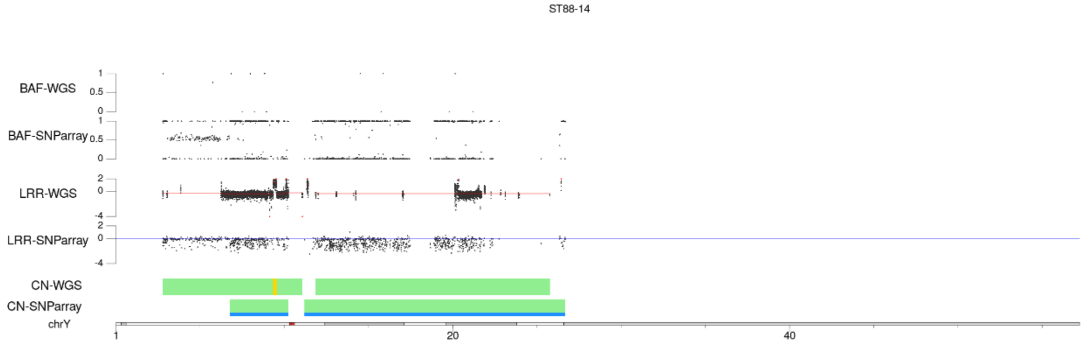
ST88-14
BAF-WGS
BAF-SNParray
LRR-WGS
LRR-SNParray
CN-WGS
CN-SNParray
chrY
1
20
40

# NF90-8

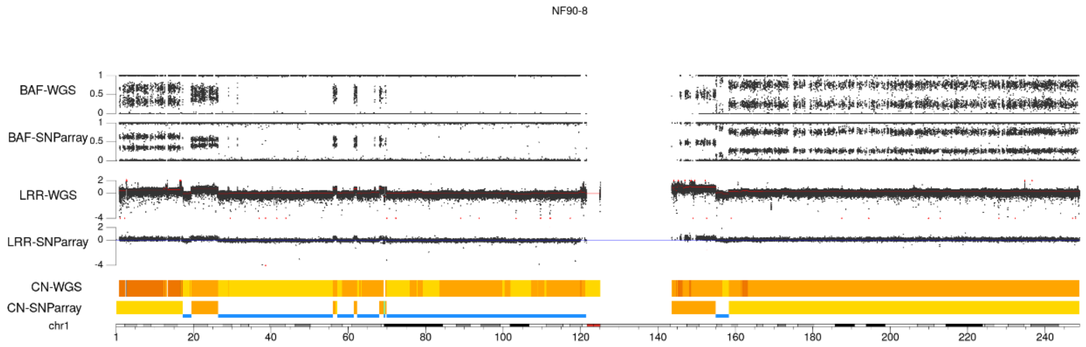
NF90-8
BAF-WGS
BAF-SNParray
LRR-WGS
LRR-SNParray
CN-WGS
CN-SNParray
chr1

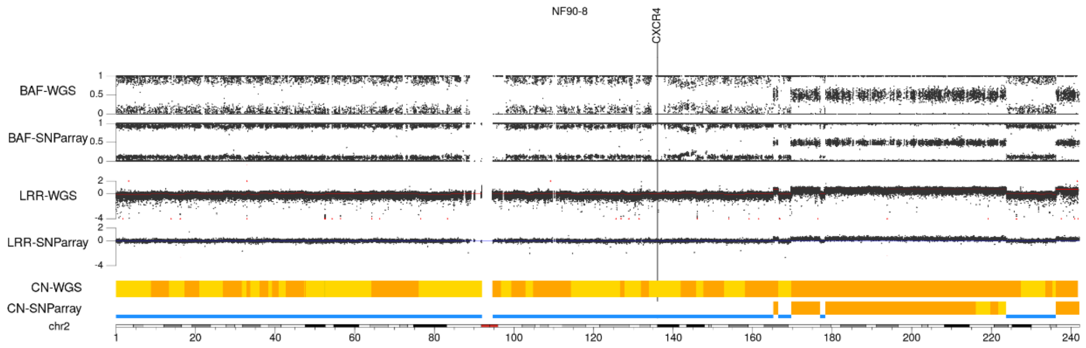
NF90-8
CXCR4
BAF-WGS
BAF-SNParray
LRR-WGS
LRR-SNParray
CN-WGS
CN-SNParray
chr2

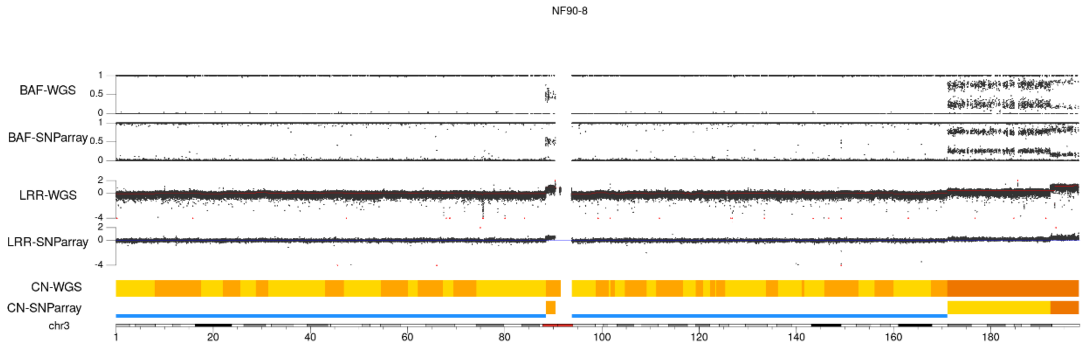
NF90-8
BAF-WGS
BAF-SNParray
LRR-WGS
LRR-SNParray
CN-WGS
CN-SNParray
chr3

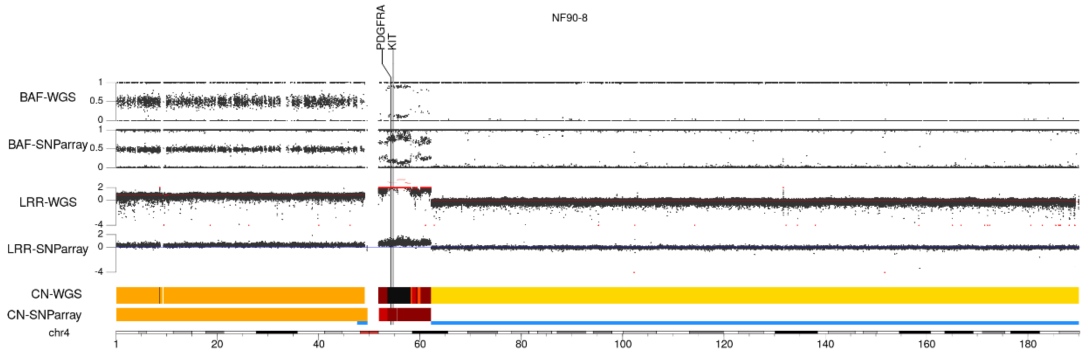
NF90-8
PDGFRA
KIT
BAF-WGS
BAF-SNParray
LRR-WGS
LRR-SNParray
CN-WGS
CN-SNParray
chr4

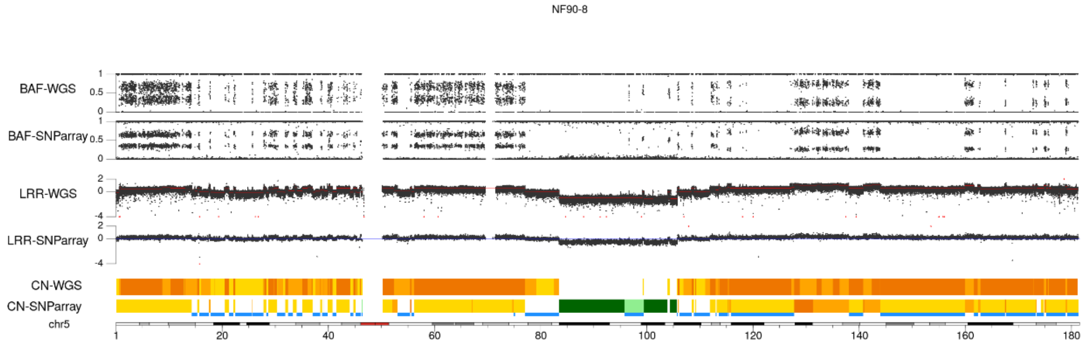
NF90-8
BAF-WGS
BAF-SNParray
LRR-WGS
LRR-SNParray
CN-WGS
CN-SNParray
chr5
1 20 40 60 80 100 120 140 160 180

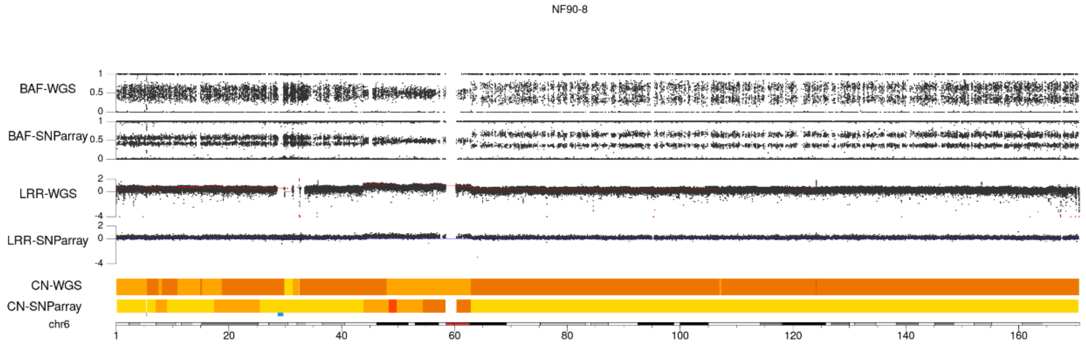
NF90-8
BAF-WGS
BAF-SNParray
LRR-WGS
LRR-SNParray
CN-WGS
CN-SNParray
chr6
1 20 40 60 80 100 120 140 160

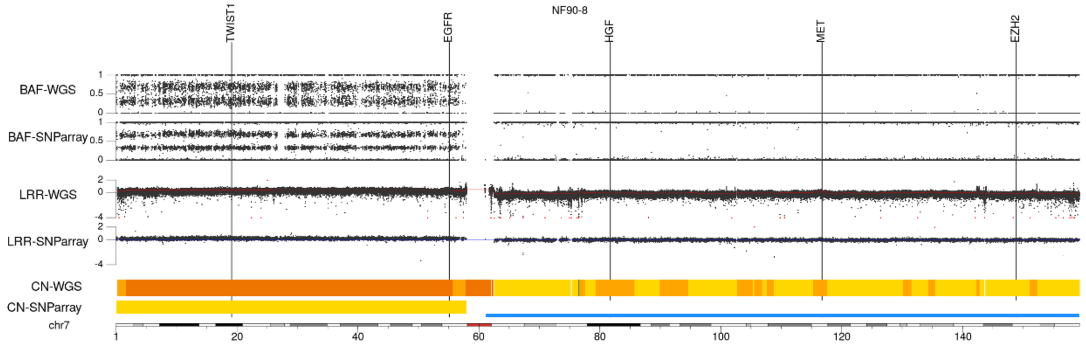
NF90-8
TWIST1
EGFR
HGF
MET
EZH2
BAF-WGS
BAF-SNParray
LRR-WGS
LRR-SNParray
CN-WGS
CN-SNParray
chr7
1 20 40 60 80 100 120 140

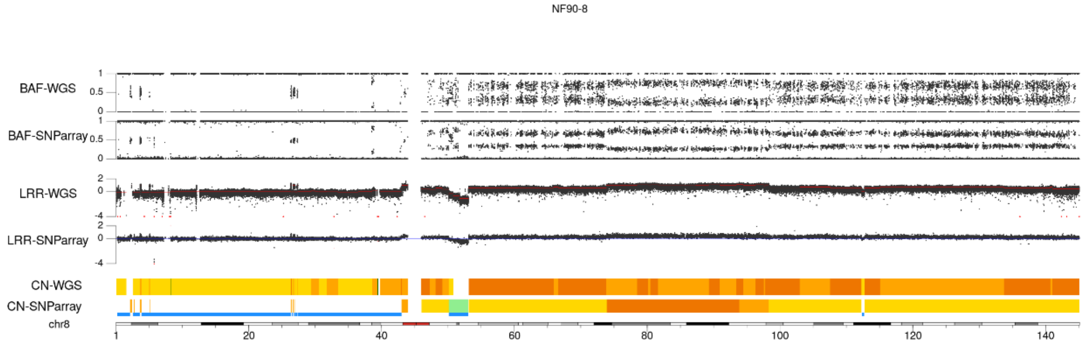
NF90-8
BAF-WGS
BAF-SNParray
LRR-WGS
LRR-SNParray
CN-WGS
CN-SNParray
chr8
1 20 40 60 80 100 120 140

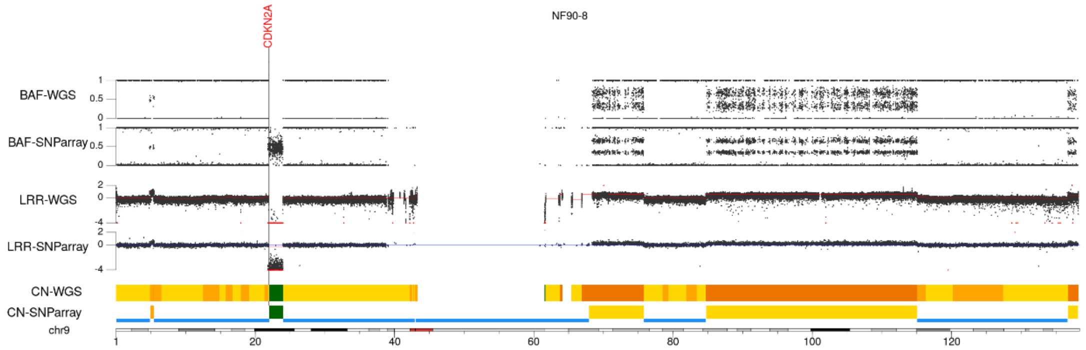
NF90-8
CDKN2A
BAF-WGS
BAF-SNParray
LRR-WGS
LRR-SNParray
CN-WGS
CN-SNParray
chr9
1 20 40 60 80 100 120

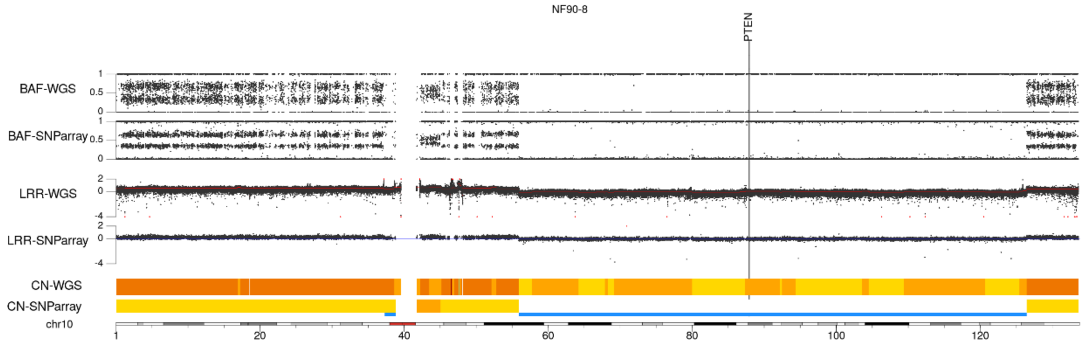
NF90-8
PTEN
BAF-WGS
BAF-SNParray
LRR-WGS
LRR-SNParray
CN-WGS
CN-SNParray
chr10
1 20 40 60 80 100 120

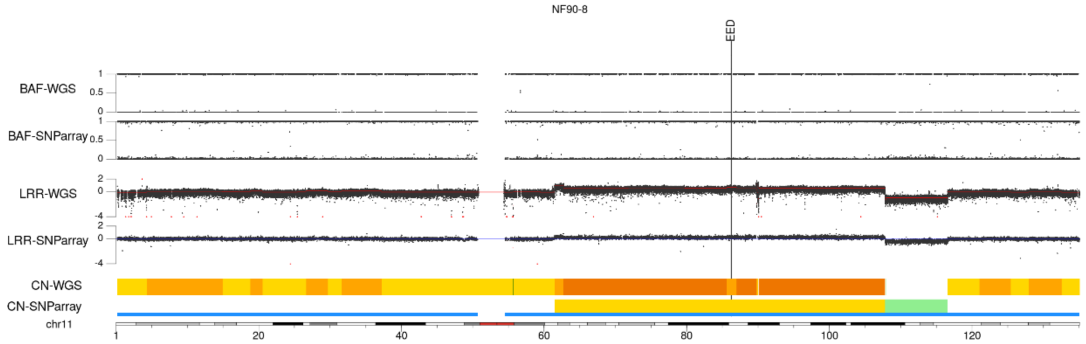
NF90-8
EED
BAF-WGS
BAF-SNParray
LRR-WGS
LRR-SNParray
CN-WGS
CN-SNParray
chr11
1 20 40 60 80 100 120

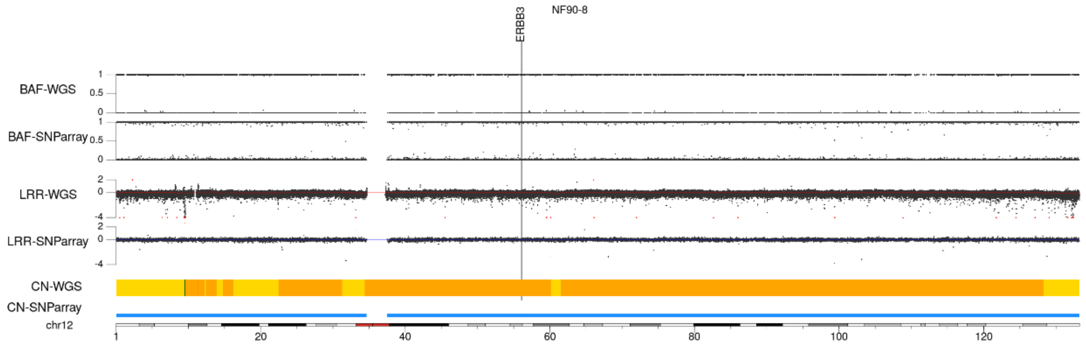
NF90-8
ERBB3
BAF-WGS
BAF-SNParray
LRR-WGS
LRR-SNParray
CN-WGS
CN-SNParray
chr12
1 20 40 60 80 100 120

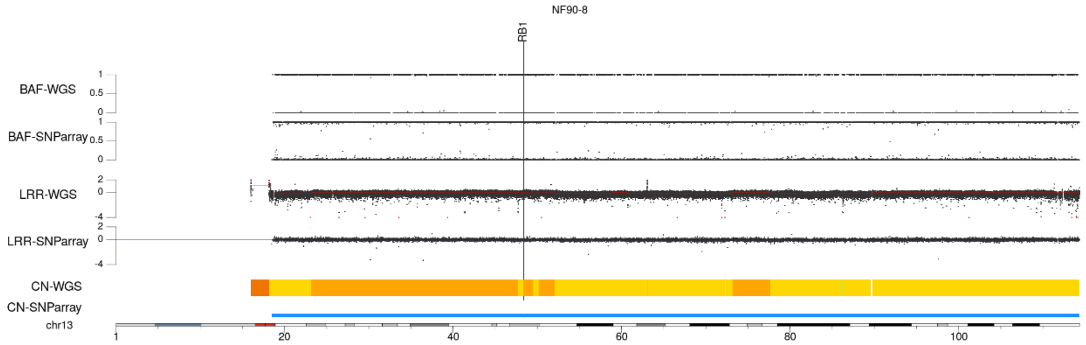
NF90-8
RB1
BAF-WGS
BAF-SNParray
LRR-WGS
LRR-SNParray
CN-WGS
CN-SNParray
chr13
1 20 40 60 80 100

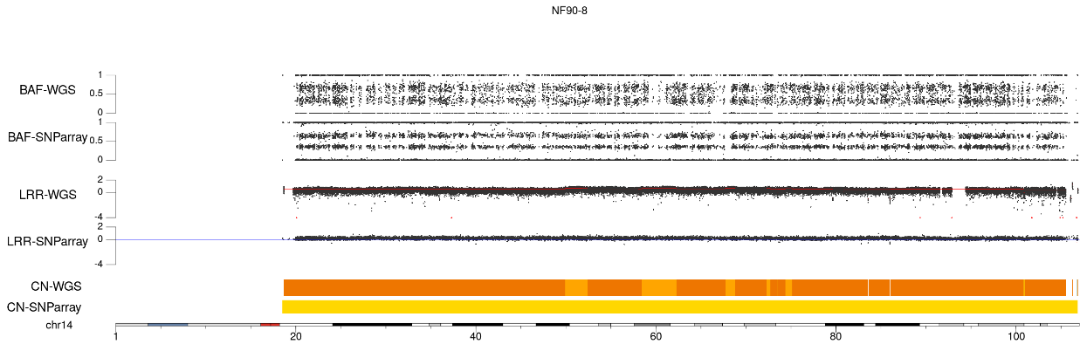
NF90-8
BAF-WGS
BAF-SNParray
LRR-WGS
LRR-SNParray
CN-WGS
CN-SNParray
chr14
1 20 40 60 80 100

NF90-8

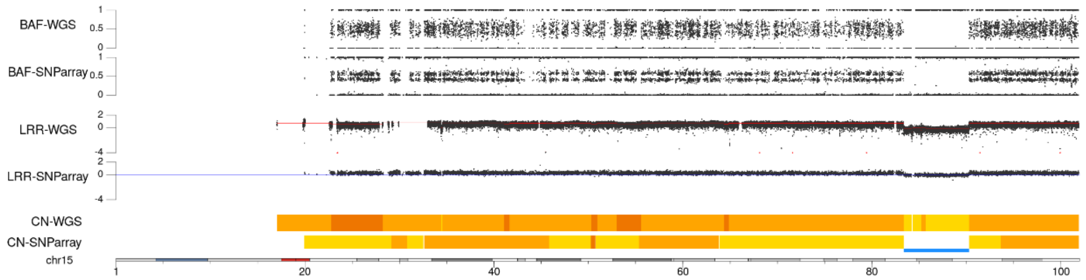


NF90-8

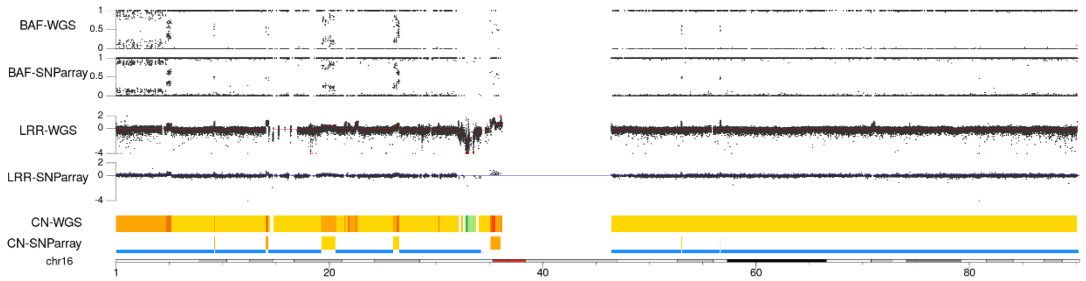


NF90-8

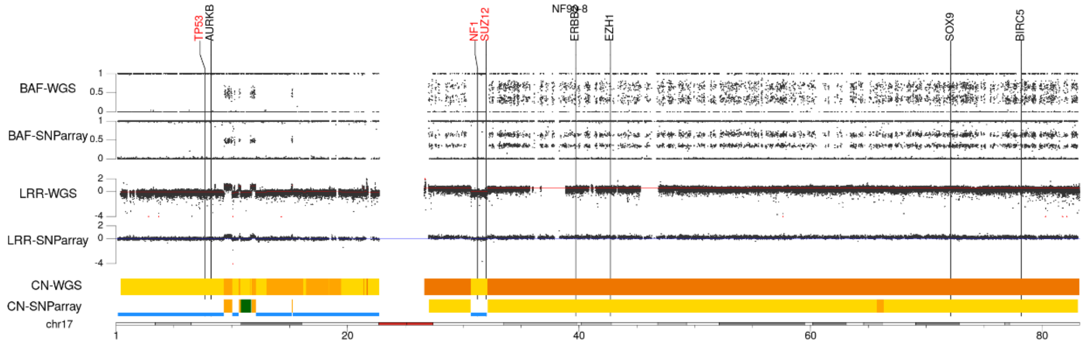


NF90-8

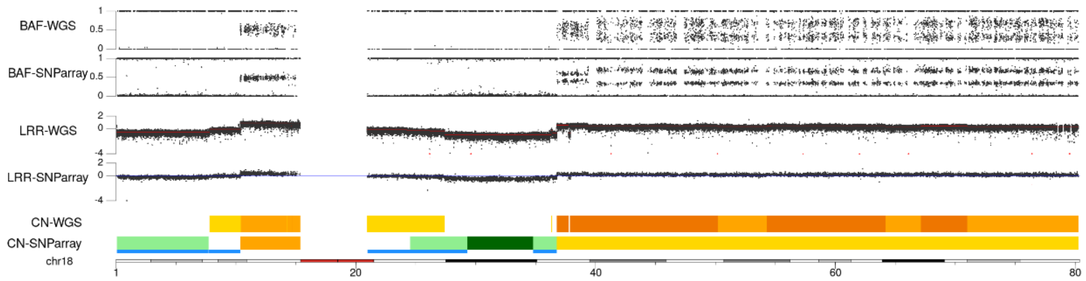


NF90-8

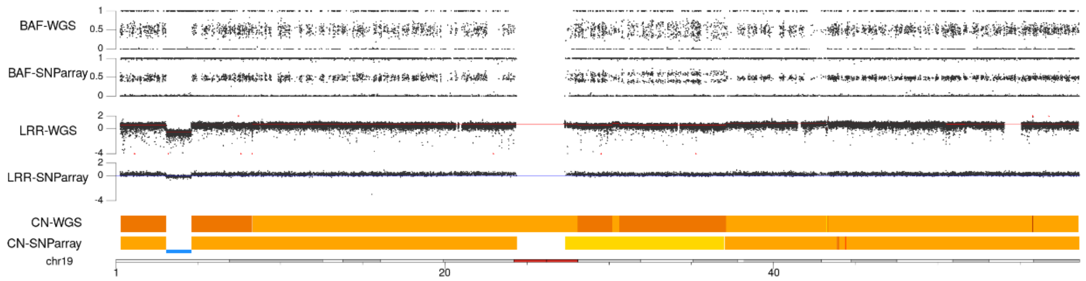

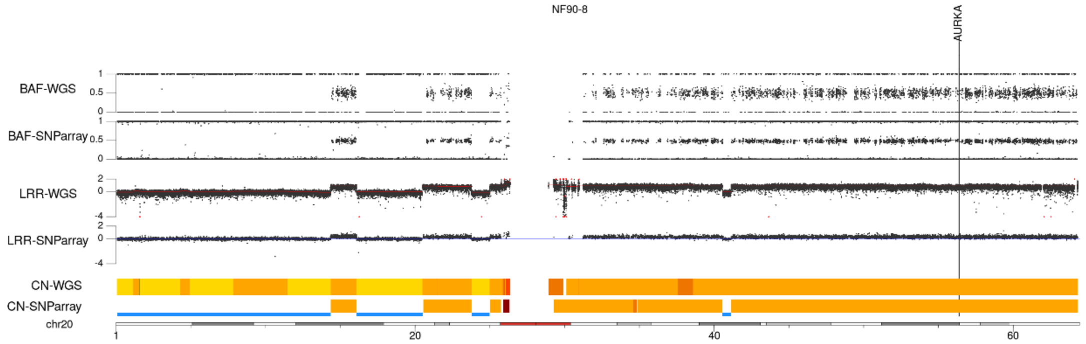
NF90-8
AURKA
BAF-WGS
BAF-SNParray
LRR-WGS
LRR-SNParray
CN-WGS
CN-SNParray
chr20
1
20
40
60

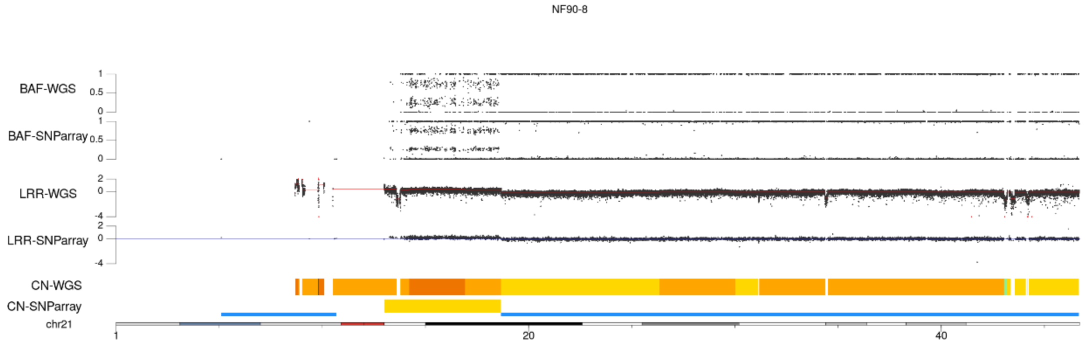
NF90-8
BAF-WGS
BAF-SNParray
LRR-WGS
LRR-SNParray
CN-WGS
CN-SNParray
chr21
1
20
40

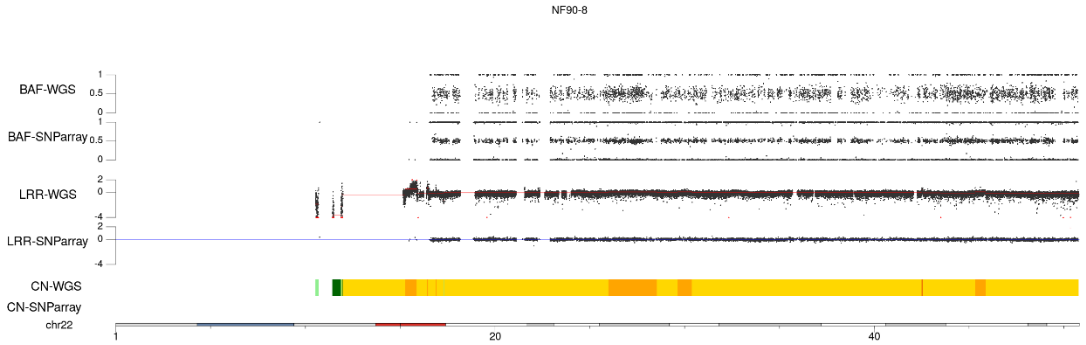
NF90-8
BAF-WGS
BAF-SNParray
LRR-WGS
LRR-SNParray
CN-WGS
CN-SNParray
chr22
1
20
40

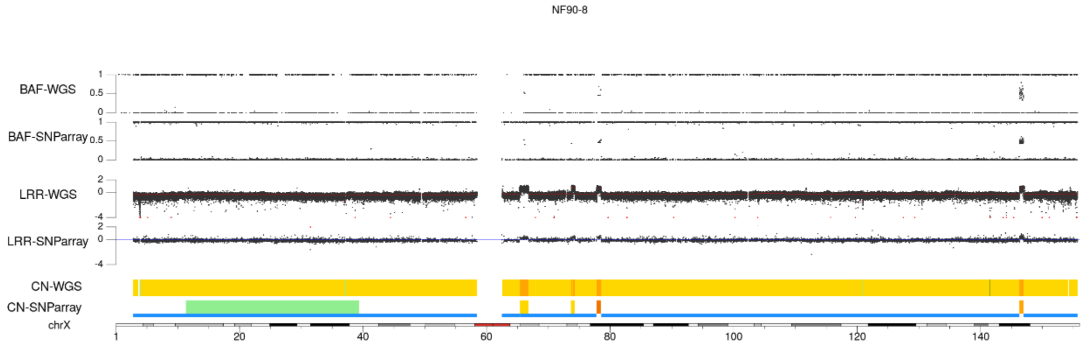
NF90-8
BAF-WGS
BAF-SNParray
LRR-WGS
LRR-SNParray
CN-WGS
CN-SNParray
chrX
1
20
40
60
80
100
120
140

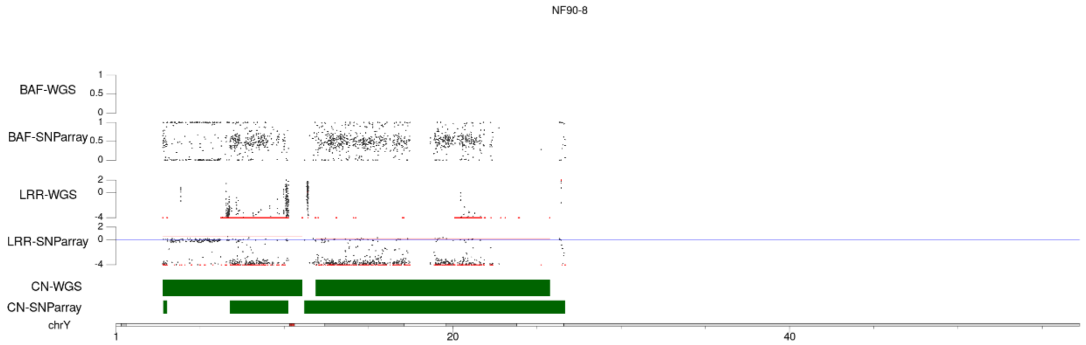
NF90-8
BAF-WGS
BAF-SNParray
LRR-WGS
LRR-SNParray
CN-WGS
CN-SNParray
chrY
1
20
40

### SNF96-2

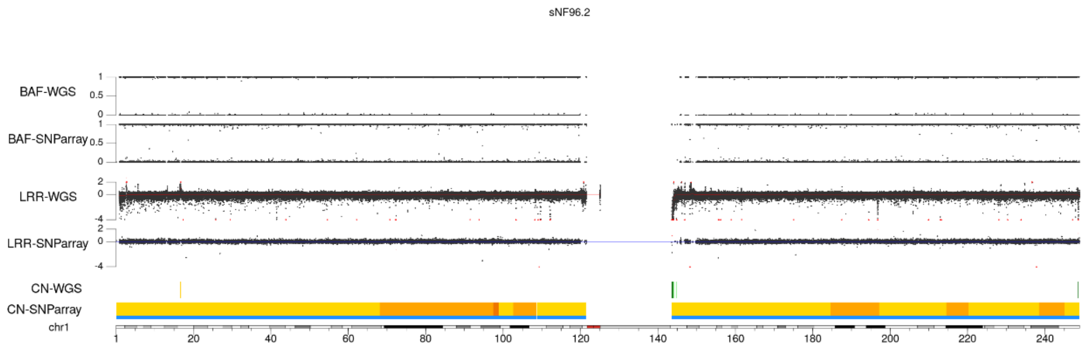
sNF96.2
BAF-WGS
BAF-SNParray
LRR-WGS
LRR-SNParray
CN-WGS
CN-SNParray
chr1

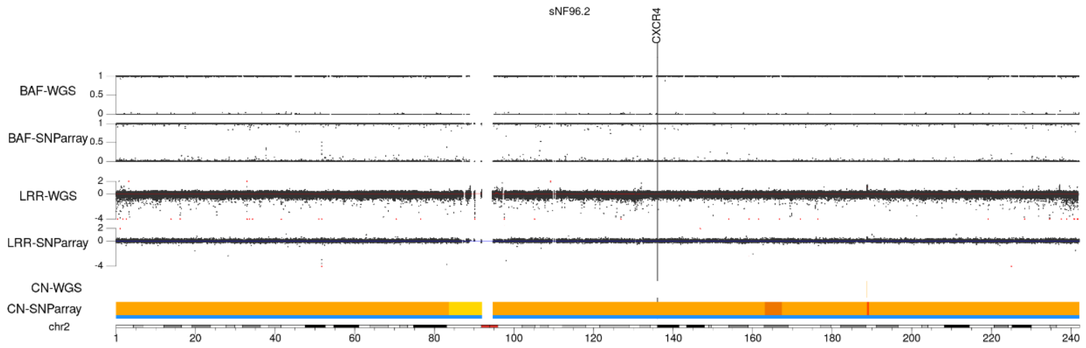
sNF96.2
CXCR4
BAF-WGS
BAF-SNParray
LRR-WGS
LRR-SNParray
CN-WGS
CN-SNParray
chr2

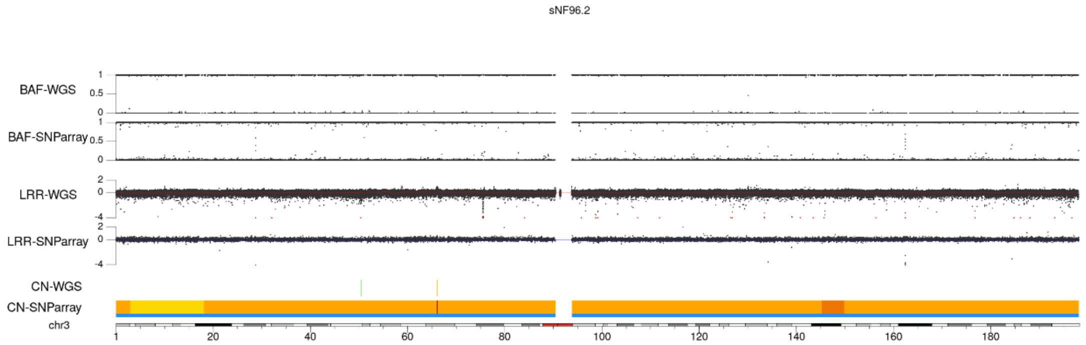
sNF96.2
BAF-WGS
BAF-SNParray
LRR-WGS
LRR-SNParray
CN-WGS
CN-SNParray
chr3

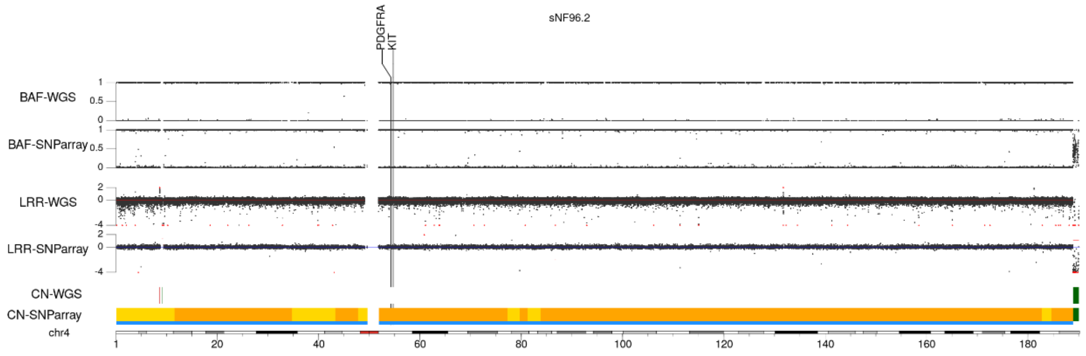
sNF96.2
PDGFRA
KIT
BAF-WGS
BAF-SNParray
LRR-WGS
LRR-SNParray
CN-WGS
CN-SNParray
chr4

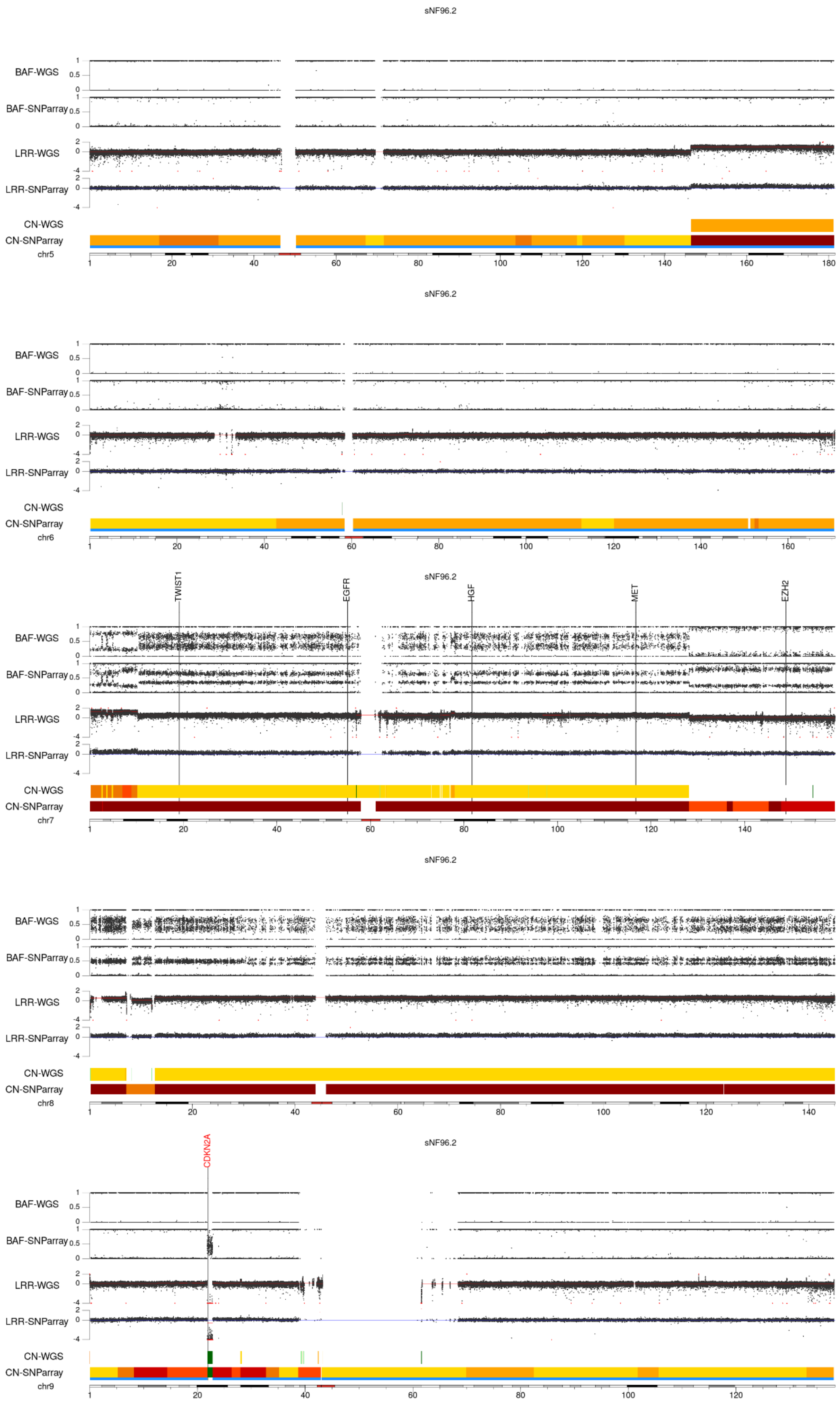
sNF96.2
BAF-WGS
BAF-SNParray
LRR-WGS
LRR-SNParray
CN-WGS
CN-SNParray
chr5
sNF96.2
BAF-WGS
BAF-SNParray
LRR-WGS
LRR-SNParray
CN-WGS
CN-SNParray
chr6
sNF96.2
TWIST1
EGFR
HGF
MET
EZH2
BAF-WGS
BAF-SNParray
LRR-WGS
LRR-SNParray
CN-WGS
CN-SNParray
chr7
sNF96.2
BAF-WGS
BAF-SNParray
LRR-WGS
LRR-SNParray
CN-WGS
CN-SNParray
chr8
sNF96.2
CDKN2A
BAF-WGS
BAF-SNParray
LRR-WGS
LRR-SNParray
CN-WGS
CN-SNParray
chr9

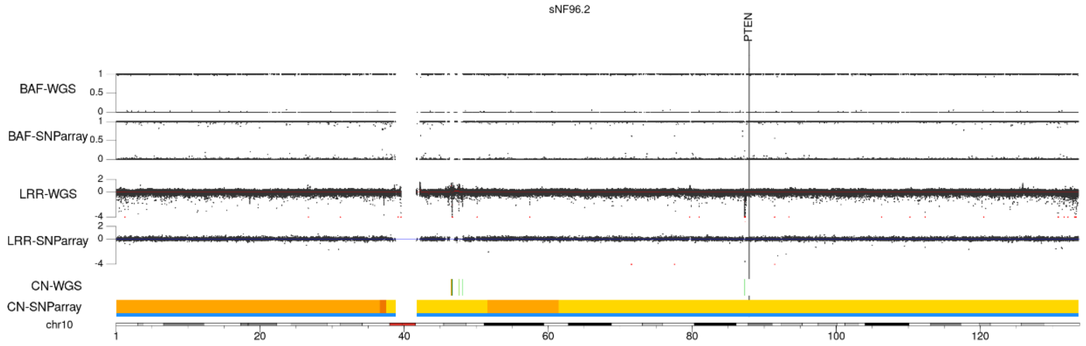
sNF96.2
PTEN
BAF-WGS
BAF-SNParray
LRR-WGS
LRR-SNParray
CN-WGS
CN-SNParray
chr10
1
20
40
60
80
100
120

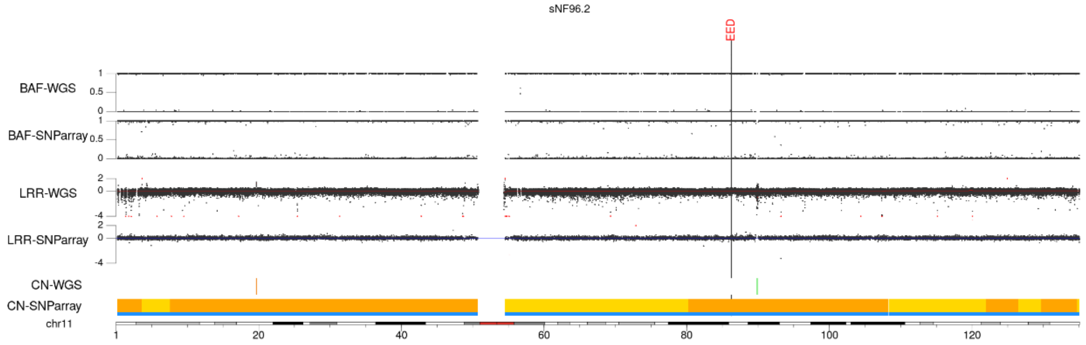
sNF96.2
EED
BAF-WGS
BAF-SNParray
LRR-WGS
LRR-SNParray
CN-WGS
CN-SNParray
chr11
1
20
40
60
80
100
120

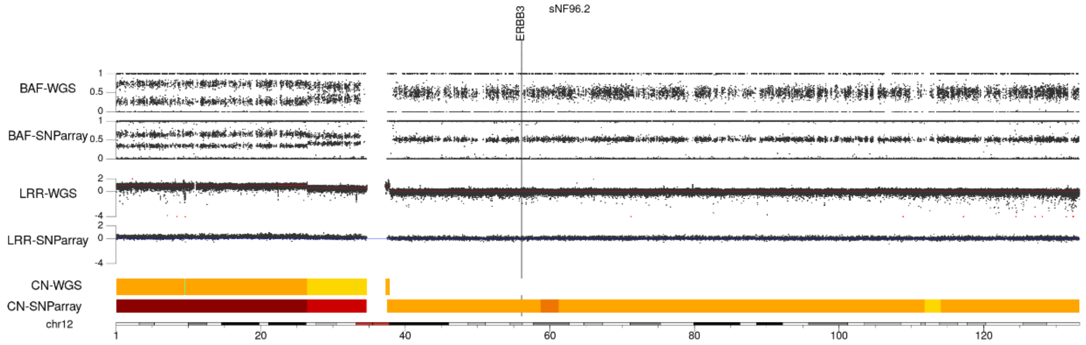
sNF96.2
ERBB3
BAF-WGS
BAF-SNParray
LRR-WGS
LRR-SNParray
CN-WGS
CN-SNParray
chr12
1
20
40
60
80
100
120

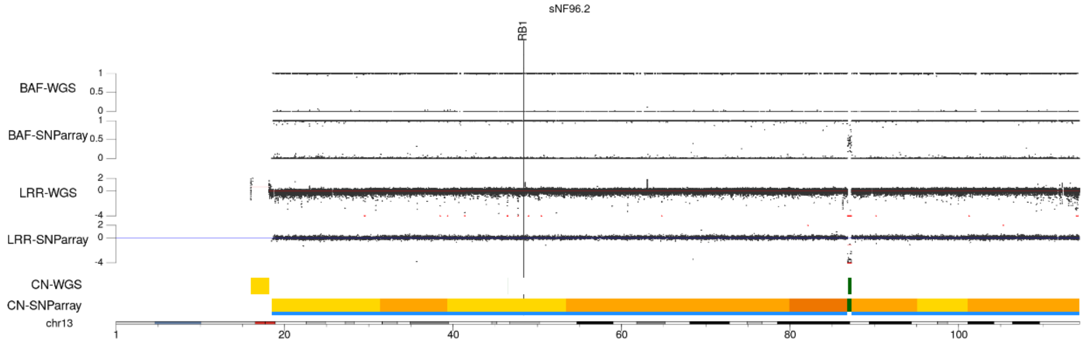
sNF96.2
RB1
BAF-WGS
BAF-SNParray
LRR-WGS
LRR-SNParray
CN-WGS
CN-SNParray
chr13
1
20
40
60
80
100

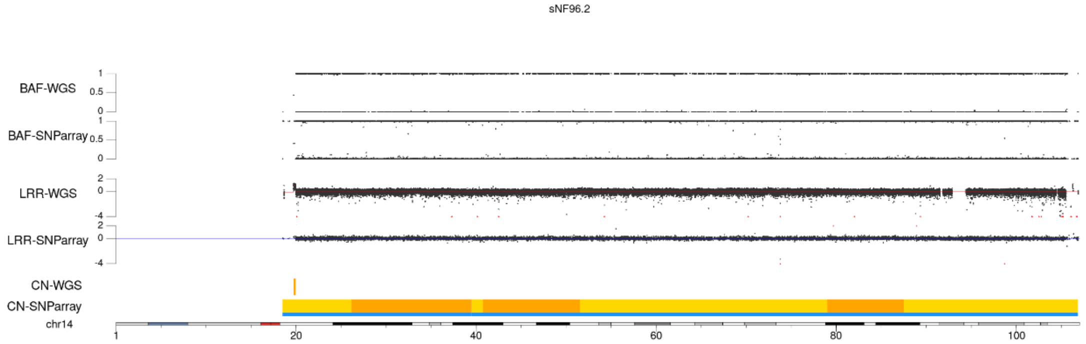
sNF96.2
BAF-WGS
BAF-SNParray
LRR-WGS
LRR-SNParray
CN-WGS
CN-SNParray
chr14
1
20
40
60
80
100

sNF96.2

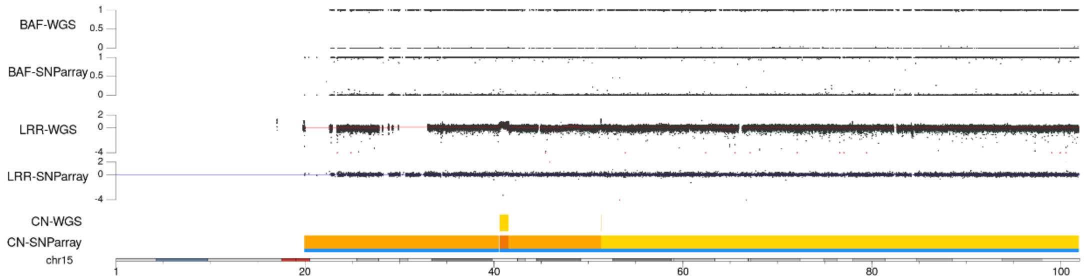


sNF96.2

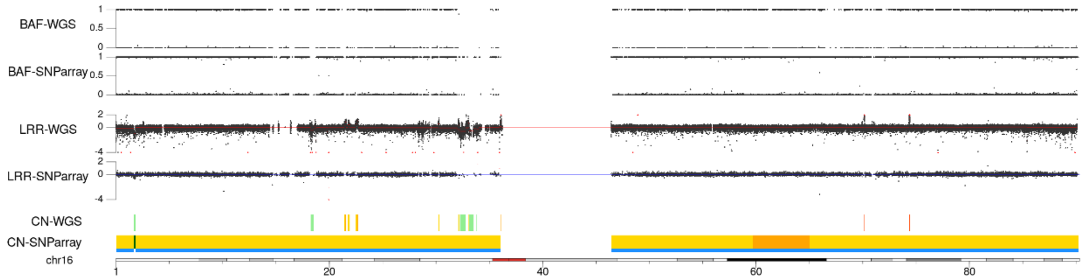


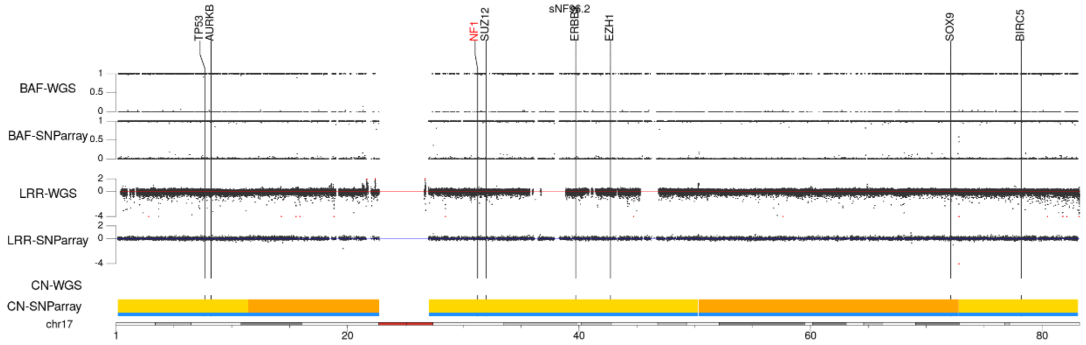


sNF96.2

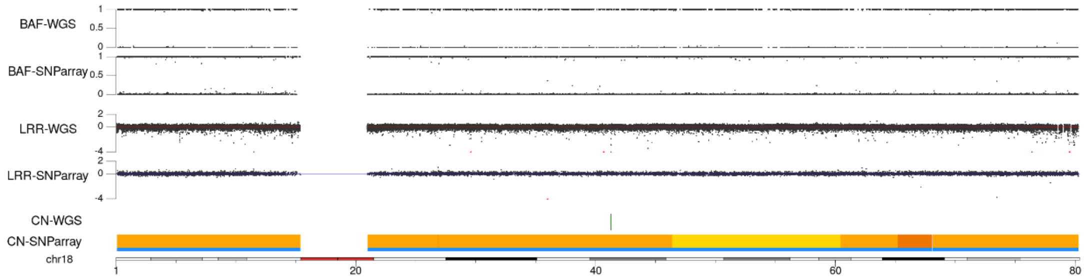


sNF96.2

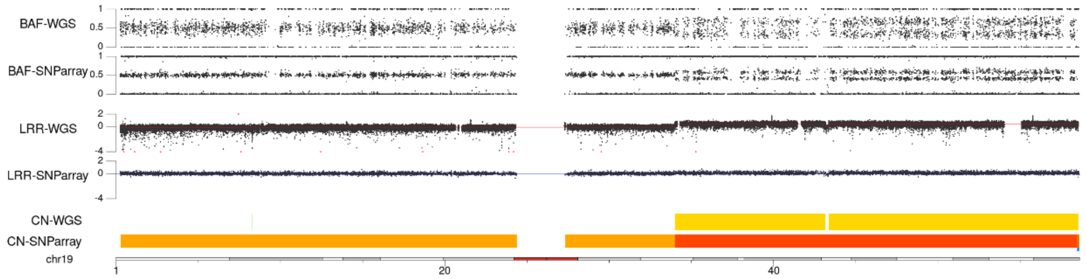

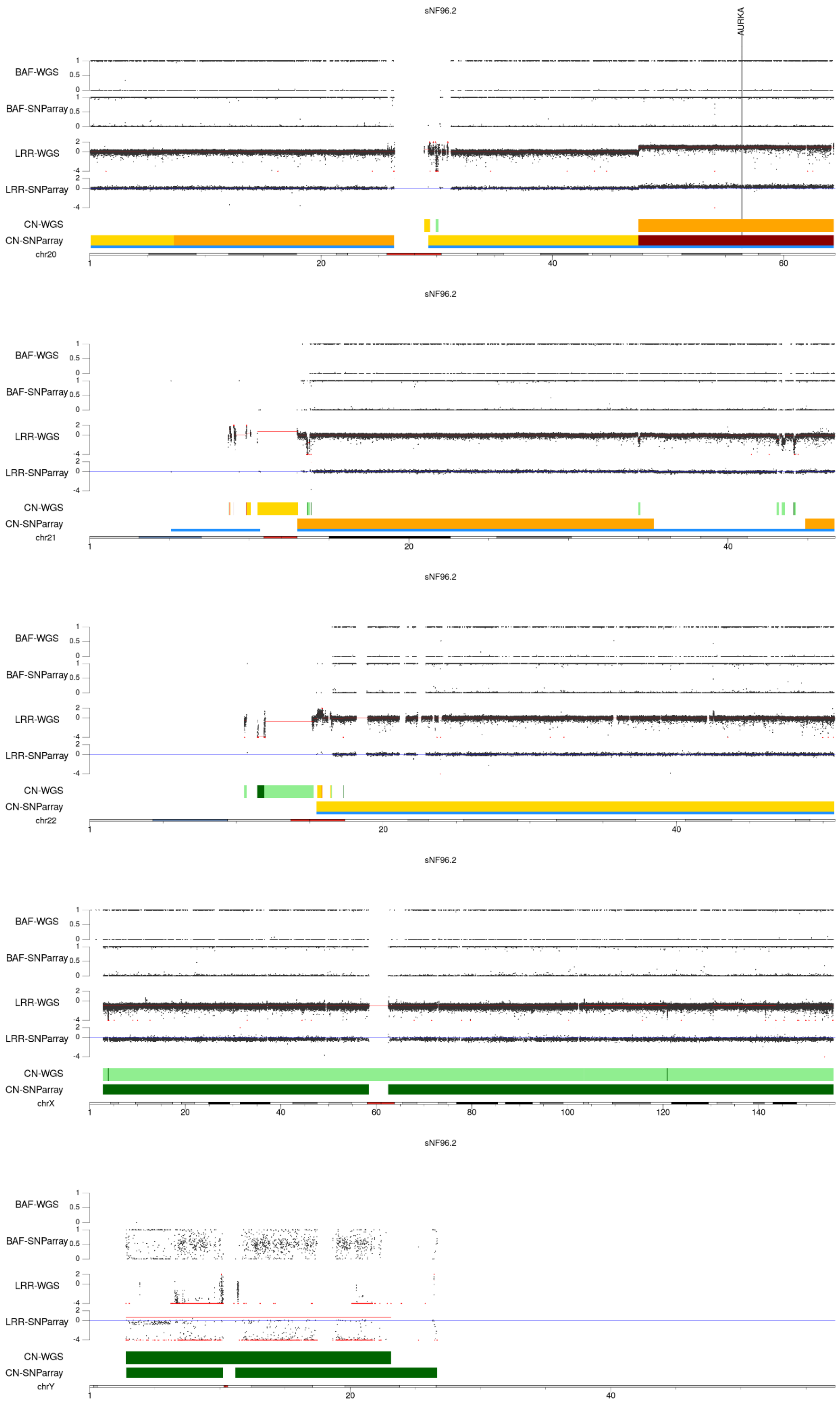
sNF96.2
AURKA
BAF-WGS
BAF-SNParray
LRR-WGS
LRR-SNParray
CN-WGS
CN-SNParray
chr20
chr21
chr22
chrX
chrY

### NMS-2

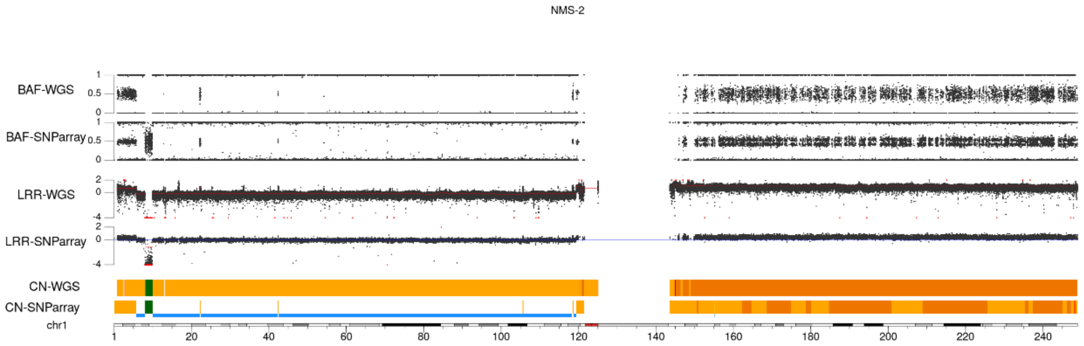
NMS-2
BAF-WGS
BAF-SNParray
LRR-WGS
LRR-SNParray
CN-WGS
CN-SNParray
chr1
1 20 40 60 80 100 120 140 160 180 200 220 240

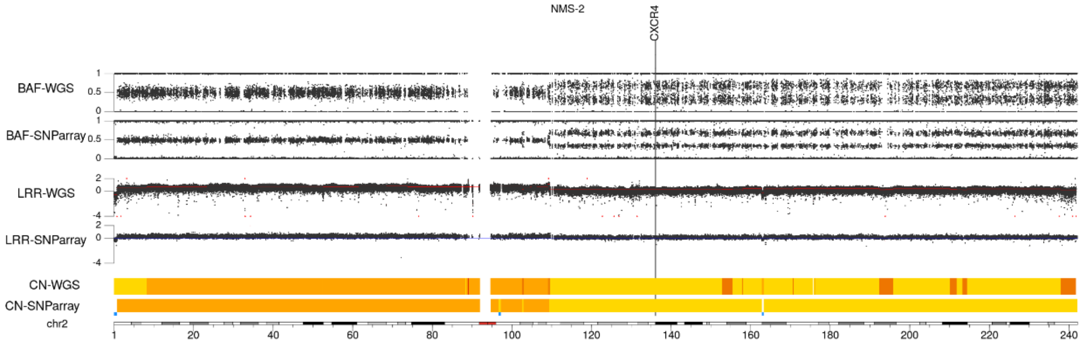
NMS-2
CXCR4
BAF-WGS
BAF-SNParray
LRR-WGS
LRR-SNParray
CN-WGS
CN-SNParray
chr2
1 20 40 60 80 100 120 140 160 180 200 220 240

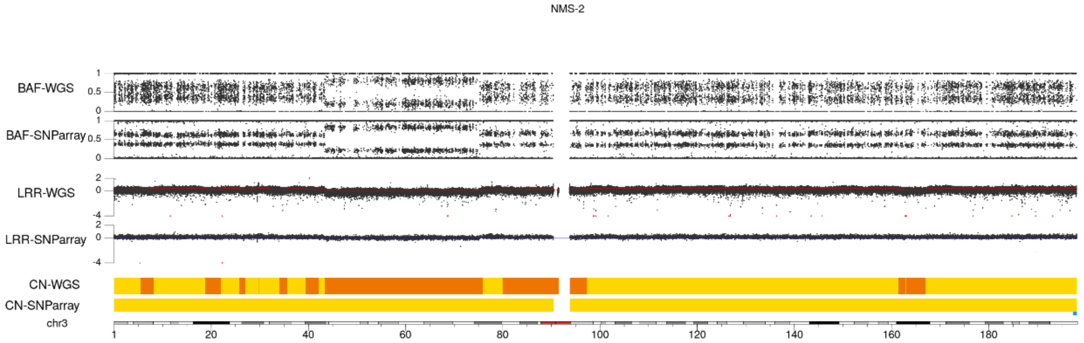
NMS-2
BAF-WGS
BAF-SNParray
LRR-WGS
LRR-SNParray
CN-WGS
CN-SNParray
chr3
1 20 40 60 80 100 120 140 160 180

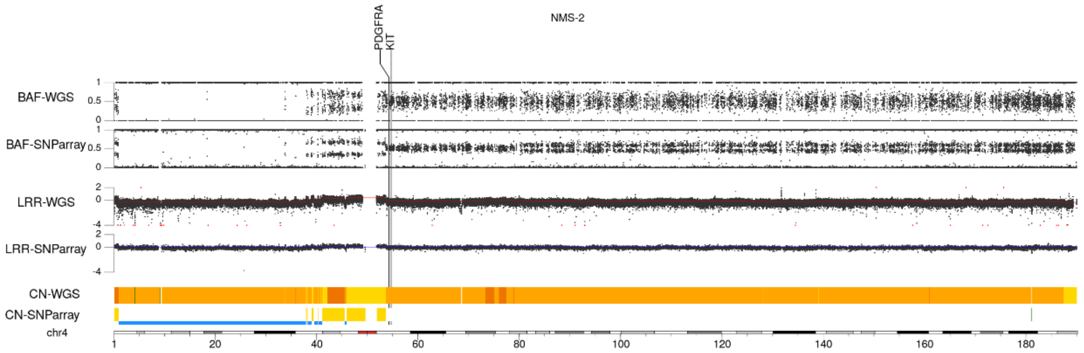
NMS-2
PDGFRA
KIT
BAF-WGS
BAF-SNParray
LRR-WGS
LRR-SNParray
CN-WGS
CN-SNParray
chr4
1 20 40 60 80 100 120 140 160 180

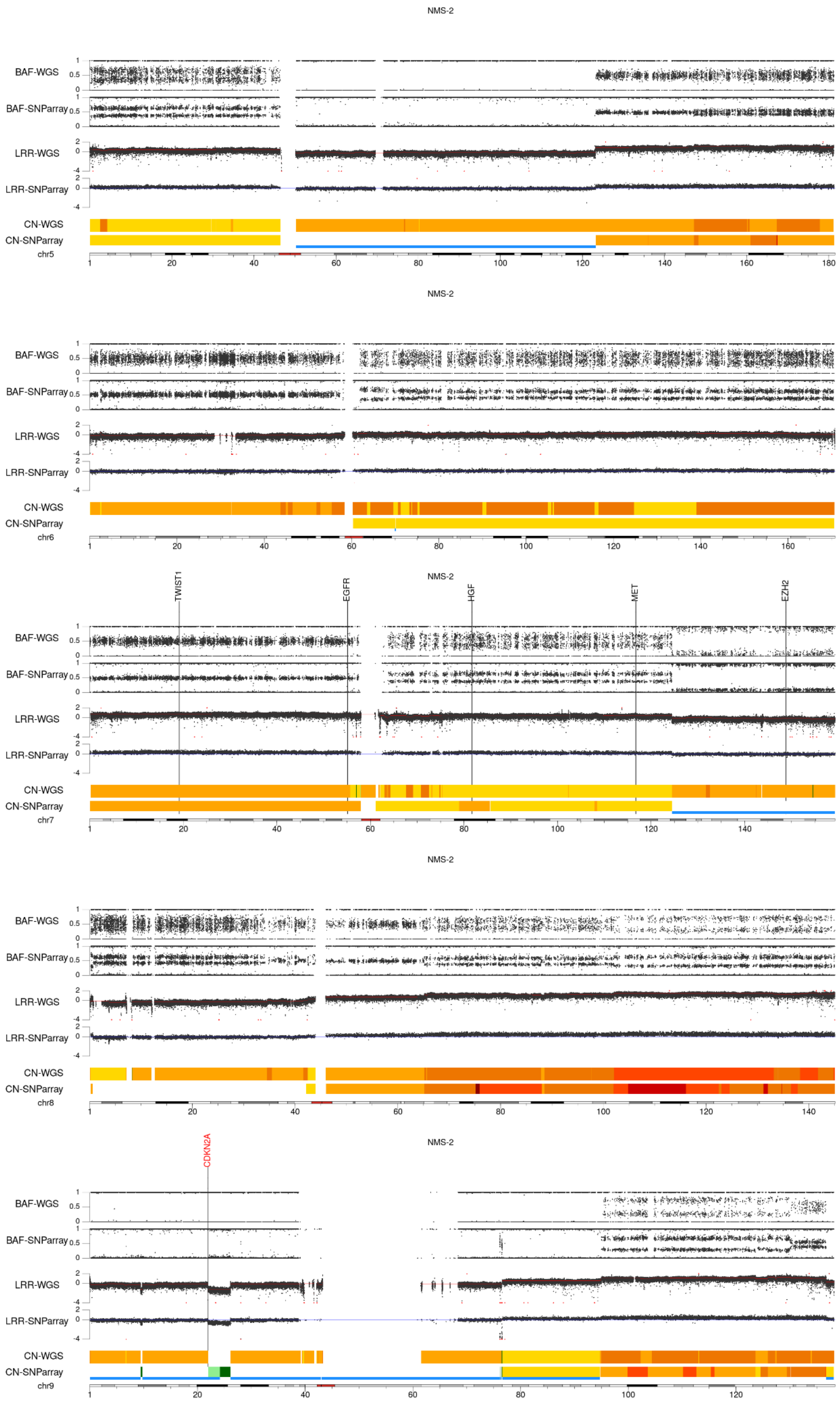
NMS-2
BAF-WGS
BAF-SNParray
LRR-WGS
LRR-SNParray
CN-WGS
CN-SNParray
chr5
NMS-2
BAF-WGS
BAF-SNParray
LRR-WGS
LRR-SNParray
CN-WGS
CN-SNParray
chr6
TWIST1
EGFR
NMS-2
HGF
MET
EZH2
BAF-WGS
BAF-SNParray
LRR-WGS
LRR-SNParray
CN-WGS
CN-SNParray
chr7
NMS-2
BAF-WGS
BAF-SNParray
LRR-WGS
LRR-SNParray
CN-WGS
CN-SNParray
chr8
CDKN2A
NMS-2
BAF-WGS
BAF-SNParray
LRR-WGS
LRR-SNParray
CN-WGS
CN-SNParray
chr9

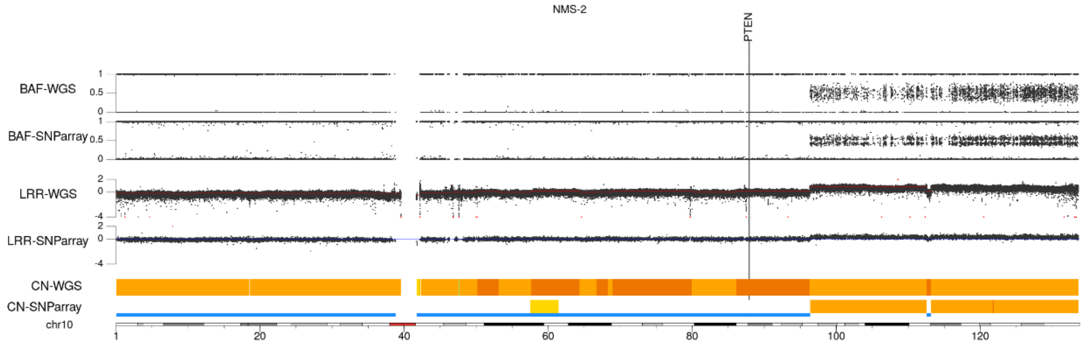
NMS-2
PTEN
BAF-WGS
BAF-SNParray
LRR-WGS
LRR-SNParray
CN-WGS
CN-SNParray
chr10

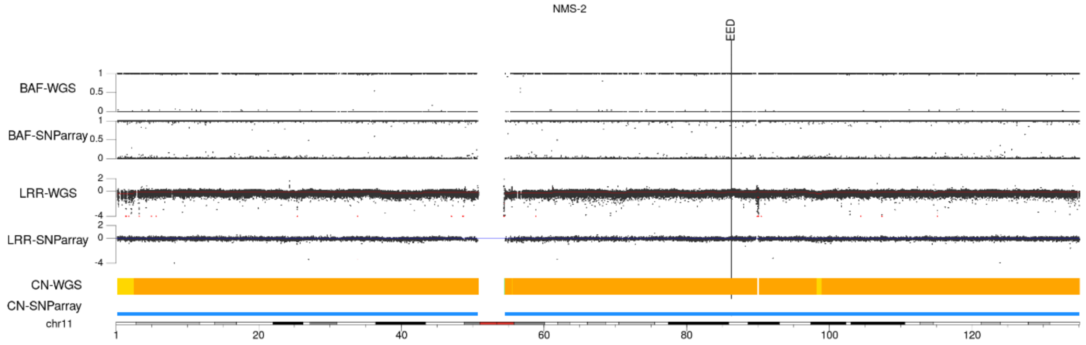
NMS-2
EED
BAF-WGS
BAF-SNParray
LRR-WGS
LRR-SNParray
CN-WGS
CN-SNParray
chr11

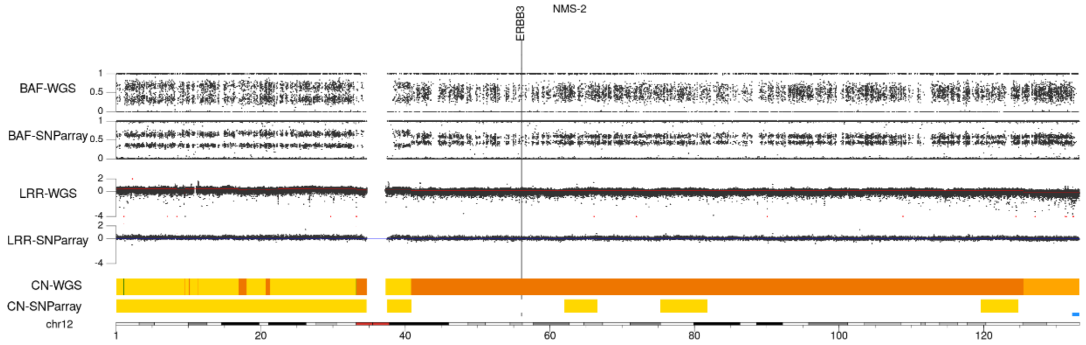
NMS-2
ERBB3
BAF-WGS
BAF-SNParray
LRR-WGS
LRR-SNParray
CN-WGS
CN-SNParray
chr12

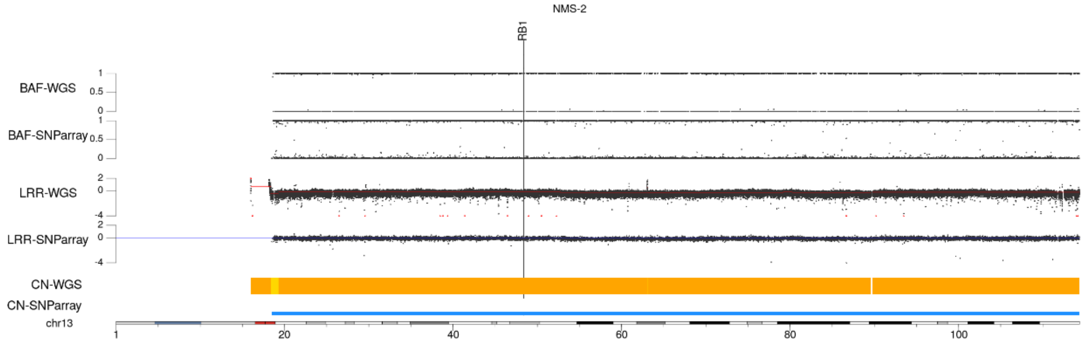
NMS-2
RB1
BAF-WGS
BAF-SNParray
LRR-WGS
LRR-SNParray
CN-WGS
CN-SNParray
chr13

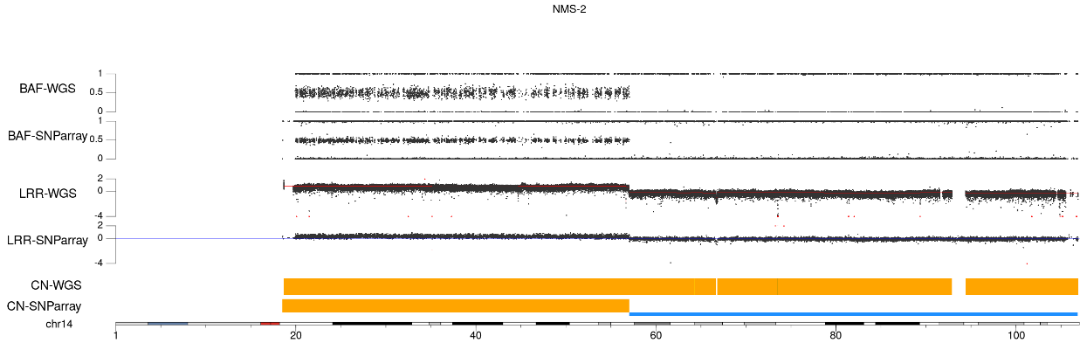
NMS-2
BAF-WGS
BAF-SNParray
LRR-WGS
LRR-SNParray
CN-WGS
CN-SNParray
chr14

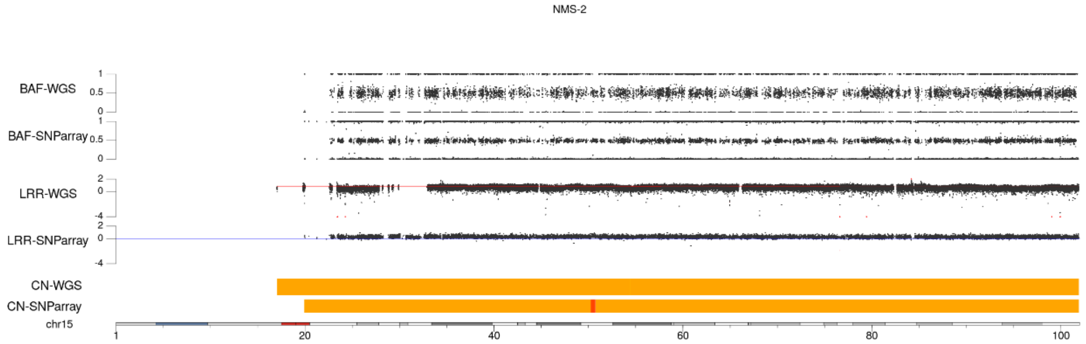
NMS-2
BAF-WGS
BAF-SNParray
LRR-WGS
LRR-SNParray
CN-WGS
CN-SNParray
chr15
1
20
40
60
80
100

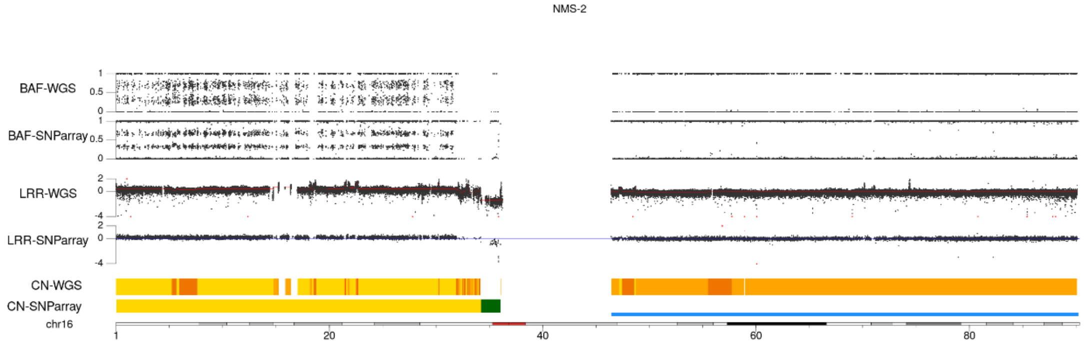
NMS-2
BAF-WGS
BAF-SNParray
LRR-WGS
LRR-SNParray
CN-WGS
CN-SNParray
chr16
1
20
40
60
80

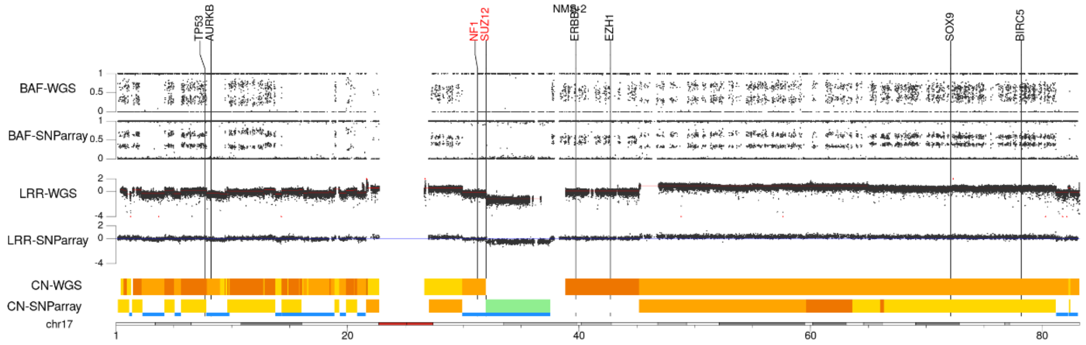
NMS-2
TP53
AURKB
NF1
SUZ12
ERBB2
EZH1
SOX9
BIRC5
BAF-WGS
BAF-SNParray
LRR-WGS
LRR-SNParray
CN-WGS
CN-SNParray
chr17
1
20
40
60
80

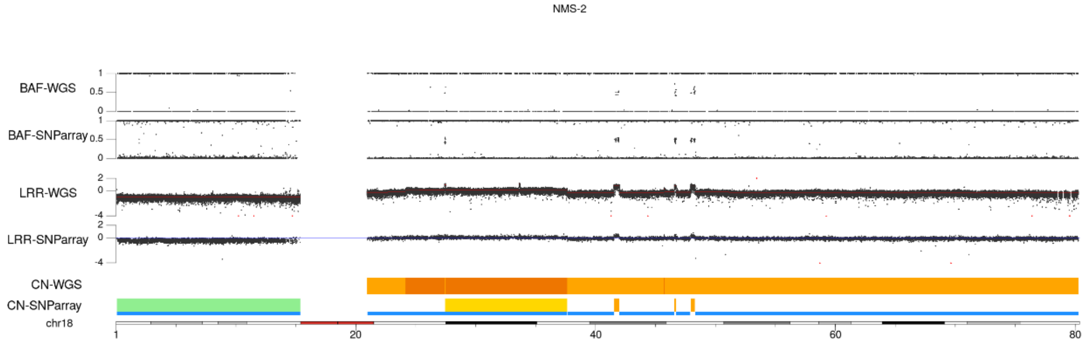
NMS-2
BAF-WGS
BAF-SNParray
LRR-WGS
LRR-SNParray
CN-WGS
CN-SNParray
chr18
1
20
40
60
80

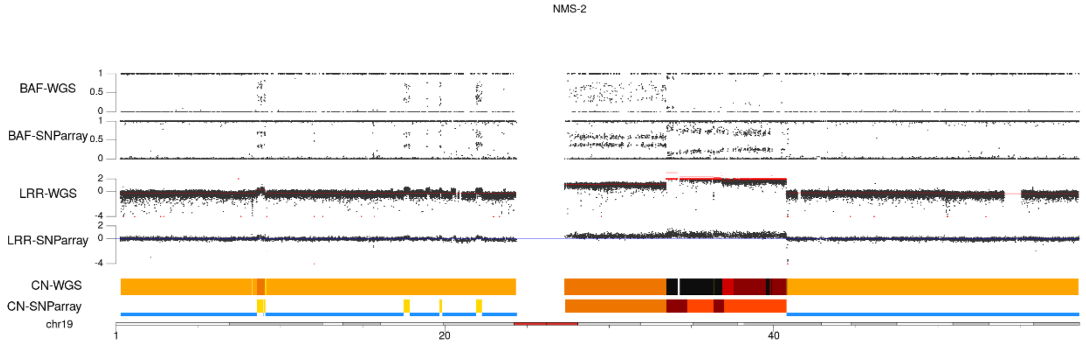
NMS-2
BAF-WGS
BAF-SNParray
LRR-WGS
LRR-SNParray
CN-WGS
CN-SNParray
chr19
1
20
40

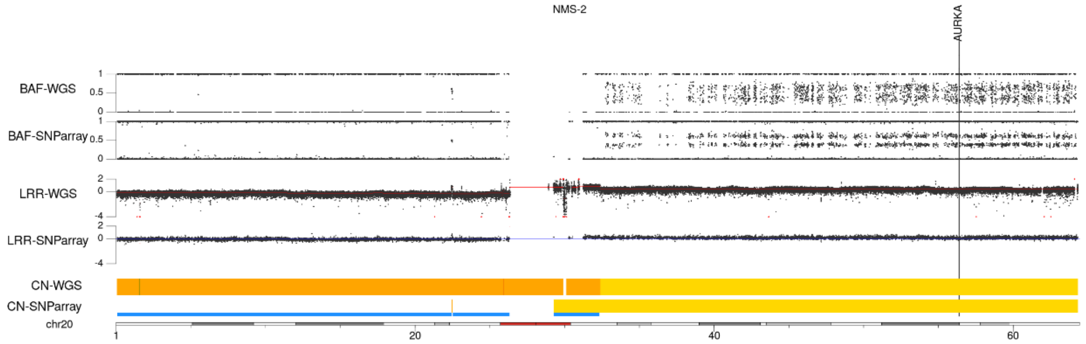
NMS-2
AURKA
BAF-WGS
BAF-SNParray
LRR-WGS
LRR-SNParray
CN-WGS
CN-SNParray
chr20
1
20
40
60

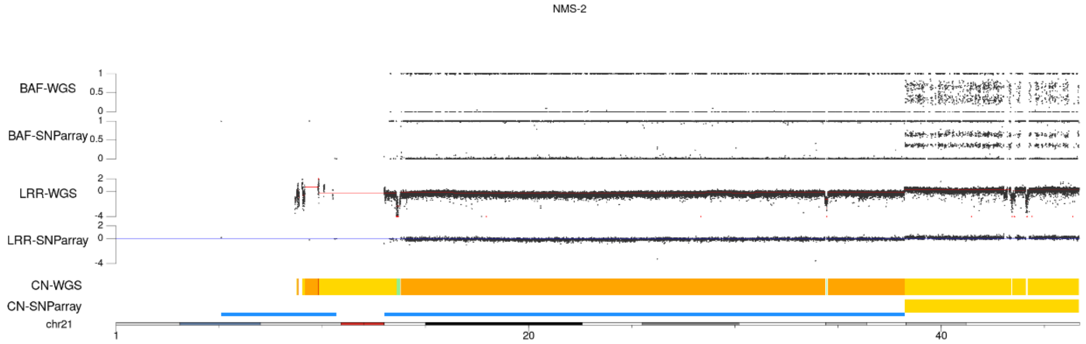
NMS-2
BAF-WGS
BAF-SNParray
LRR-WGS
LRR-SNParray
CN-WGS
CN-SNParray
chr21
1
20
40

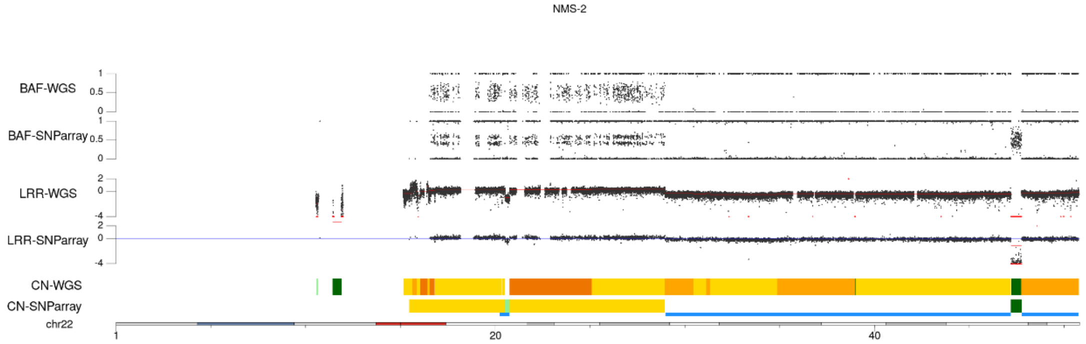
NMS-2
BAF-WGS
BAF-SNParray
LRR-WGS
LRR-SNParray
CN-WGS
CN-SNParray
chr22
1
20
40

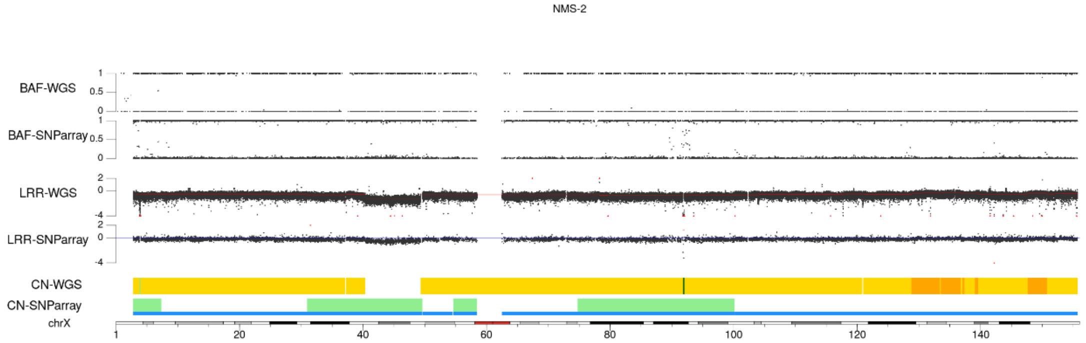
NMS-2
BAF-WGS
BAF-SNParray
LRR-WGS
LRR-SNParray
CN-WGS
CN-SNParray
chrX
1
20
40
60
80
100
120
140

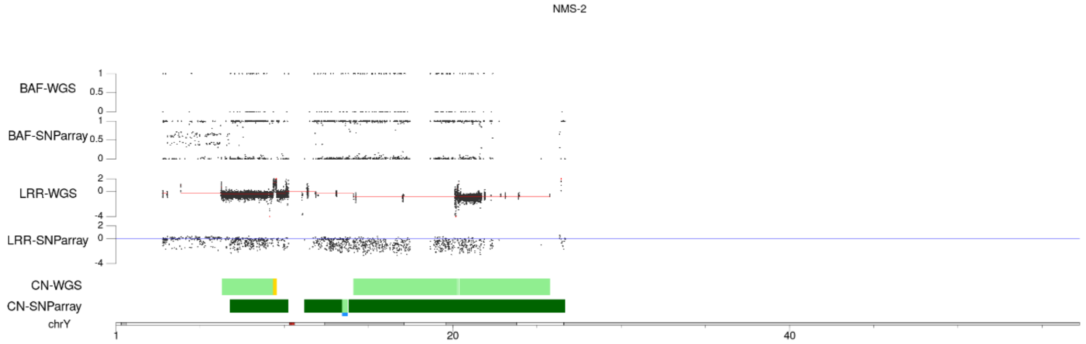
NMS-2
BAF-WGS
BAF-SNParray
LRR-WGS
LRR-SNParray
CN-WGS
CN-SNParray
chrY
1
20
40

#### STS-26T

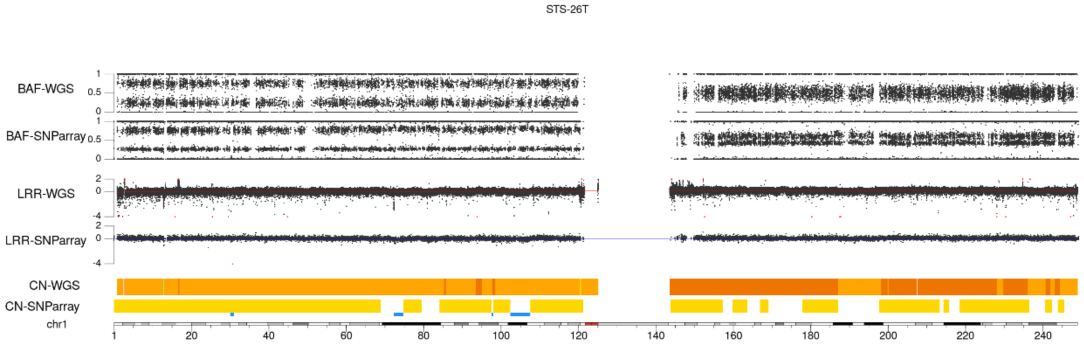
STS-26T
BAF-WGS
BAF-SNParray
LRR-WGS
LRR-SNParray
CN-WGS
CN-SNParray
chr1

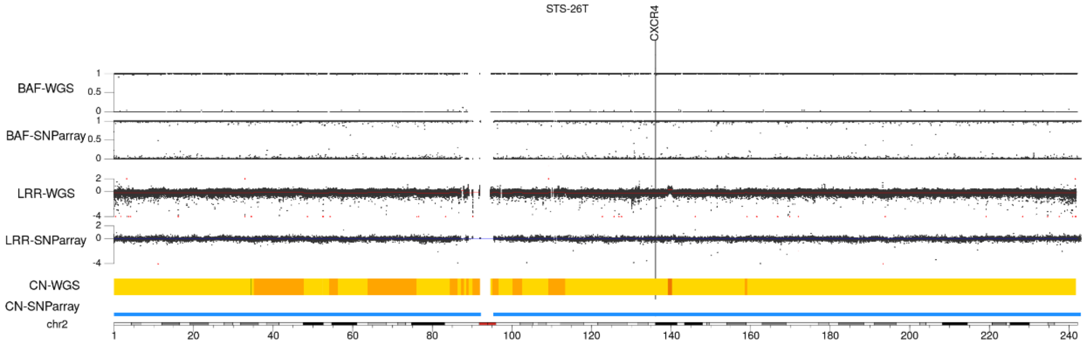
STS-26T
CXCR4
BAF-WGS
BAF-SNParray
LRR-WGS
LRR-SNParray
CN-WGS
CN-SNParray
chr2

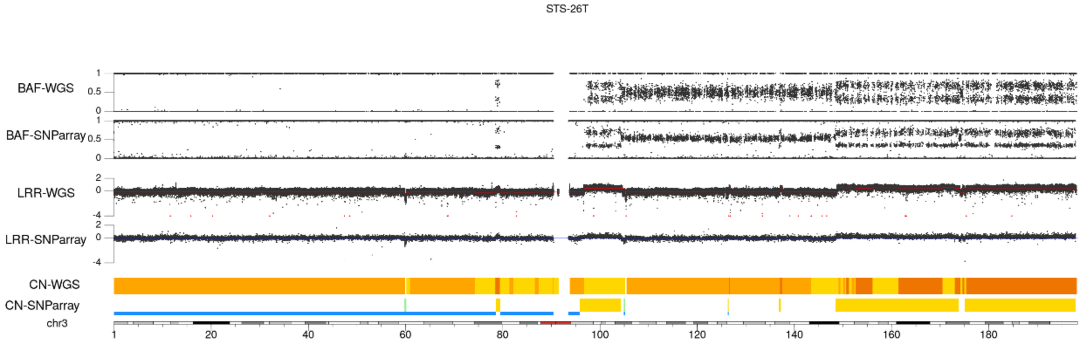
STS-26T
BAF-WGS
BAF-SNParray
LRR-WGS
LRR-SNParray
CN-WGS
CN-SNParray
chr3

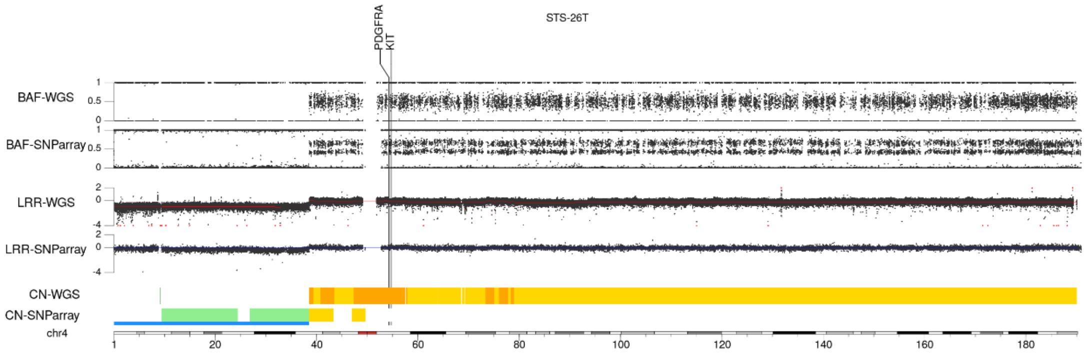
STS-26T
PDGFRA
KIT
BAF-WGS
BAF-SNParray
LRR-WGS
LRR-SNParray
CN-WGS
CN-SNParray
chr4

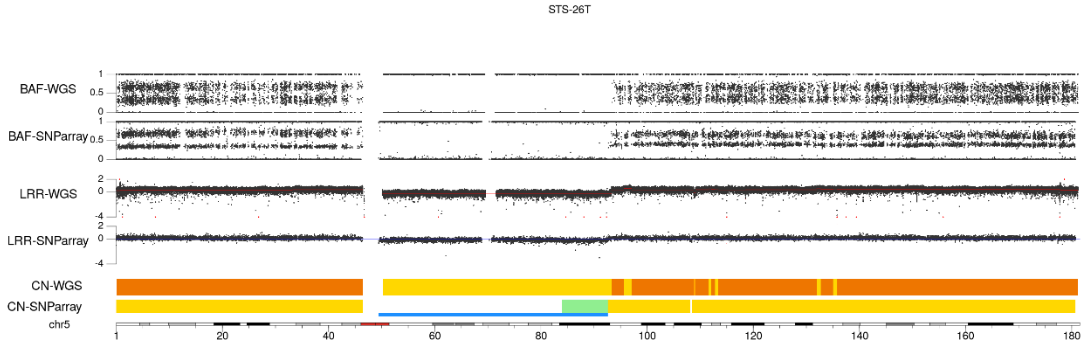
STS-26T
BAF-WGS
BAF-SNParray
LRR-WGS
LRR-SNParray
CN-WGS
CN-SNParray
chr5
1 20 40 60 80 100 120 140 160 180

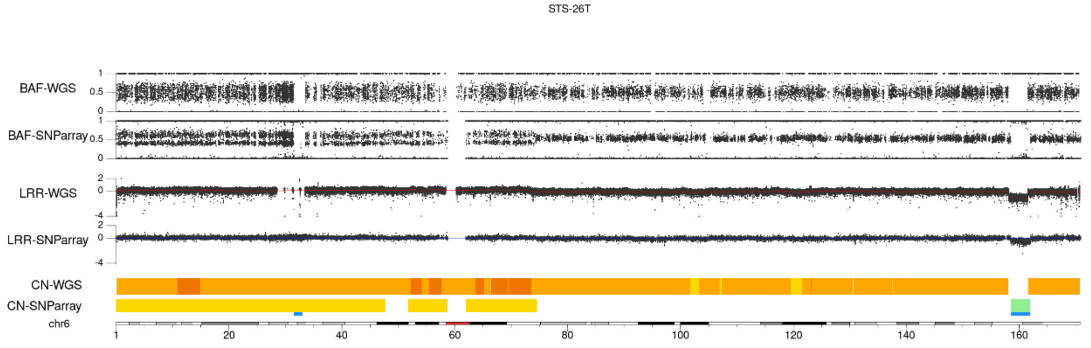
STS-26T
BAF-WGS
BAF-SNParray
LRR-WGS
LRR-SNParray
CN-WGS
CN-SNParray
chr6
1 20 40 60 80 100 120 140 160

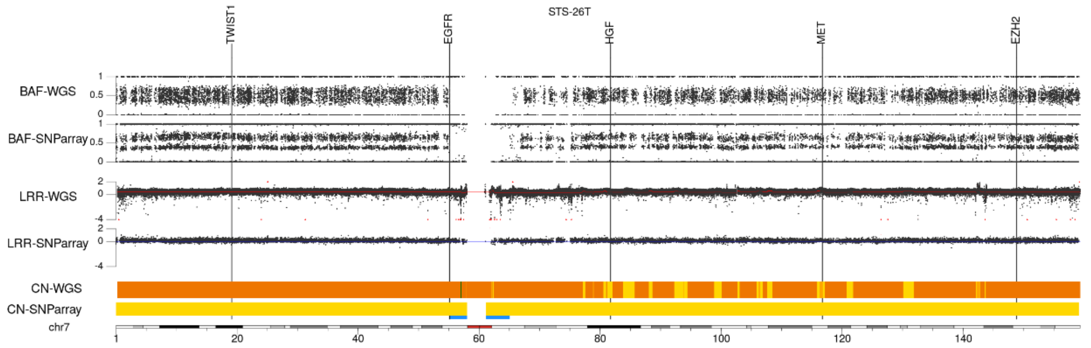
TWIST1
EGFR
STS-26T
HGF
MET
EZH2
BAF-WGS
BAF-SNParray
LRR-WGS
LRR-SNParray
CN-WGS
CN-SNParray
chr7
1 20 40 60 80 100 120 140

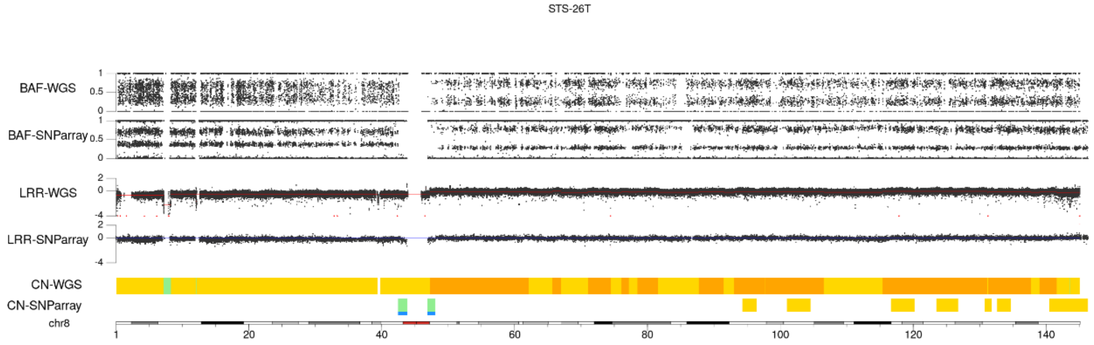
STS-26T
BAF-WGS
BAF-SNParray
LRR-WGS
LRR-SNParray
CN-WGS
CN-SNParray
chr8
1 20 40 60 80 100 120 140

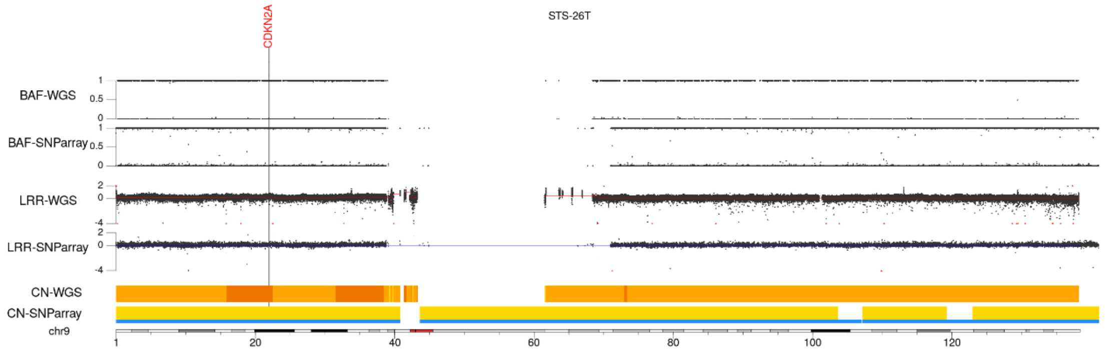
CDKN2A
STS-26T
BAF-WGS
BAF-SNParray
LRR-WGS
LRR-SNParray
CN-WGS
CN-SNParray
chr9
1 20 40 60 80 100 120

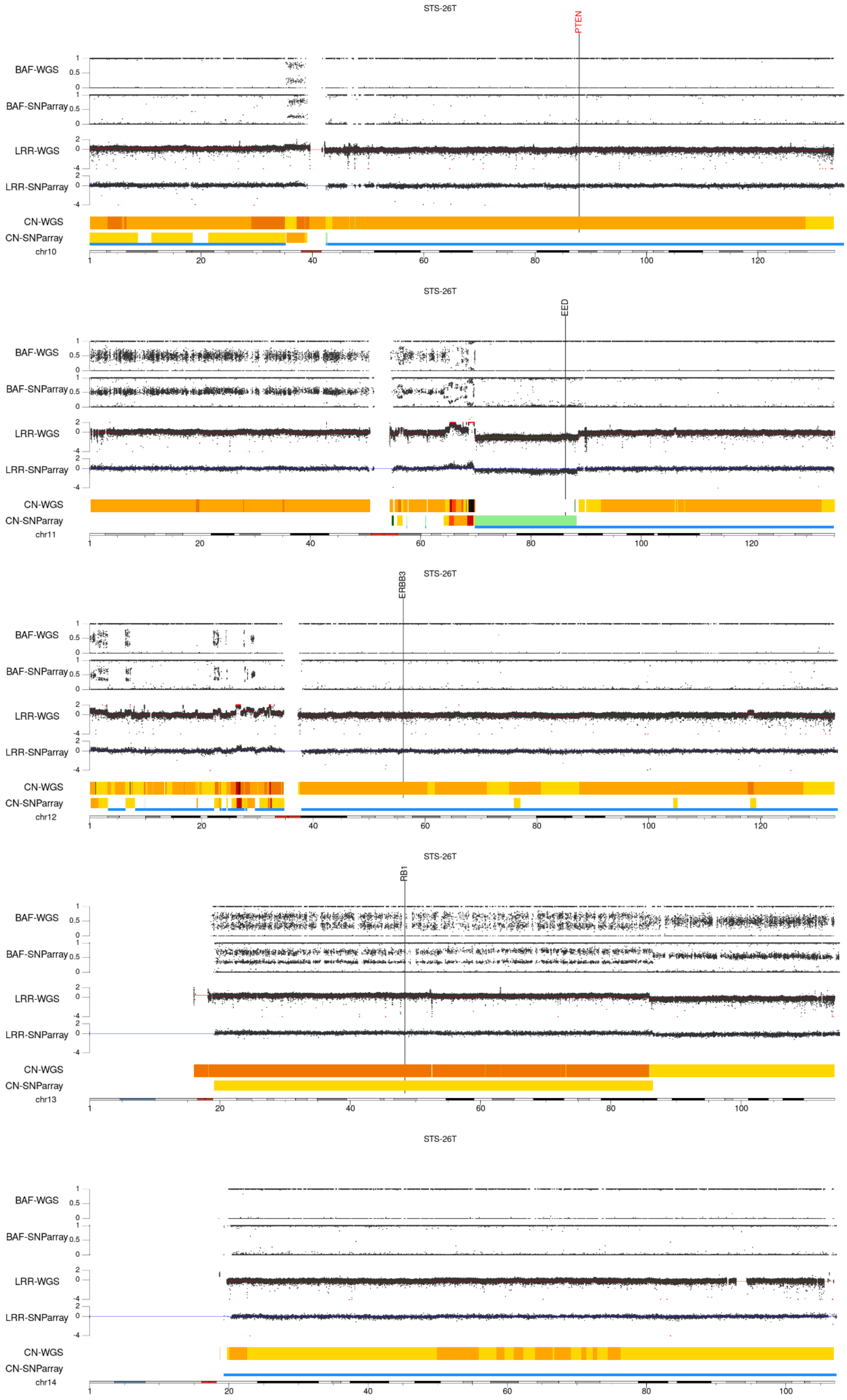
STS-26T
PTEN
BAF-WGS
BAF-SNParray
LRR-WGS
LRR-SNParray
CN-WGS
CN-SNParray
chr10
STS-26T
EED
BAF-WGS
BAF-SNParray
LRR-WGS
LRR-SNParray
CN-WGS
CN-SNParray
chr11
STS-26T
ERBB3
BAF-WGS
BAF-SNParray
LRR-WGS
LRR-SNParray
CN-WGS
CN-SNParray
chr12
STS-26T
RB1
BAF-WGS
BAF-SNParray
LRR-WGS
LRR-SNParray
CN-WGS
CN-SNParray
chr13
STS-26T
BAF-WGS
BAF-SNParray
LRR-WGS
LRR-SNParray
CN-WGS
CN-SNParray
chr14

STS-26T

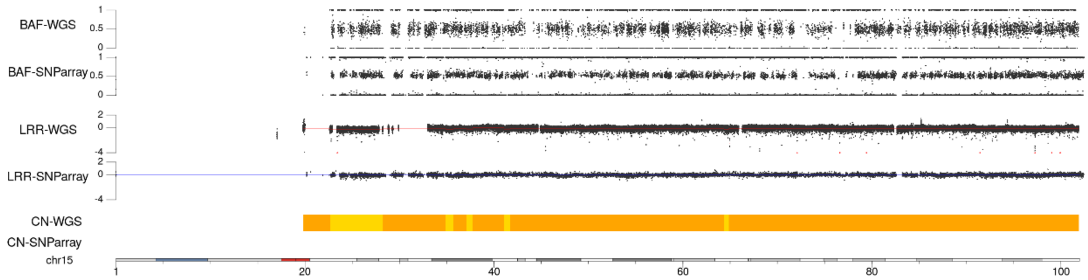


STS-26T

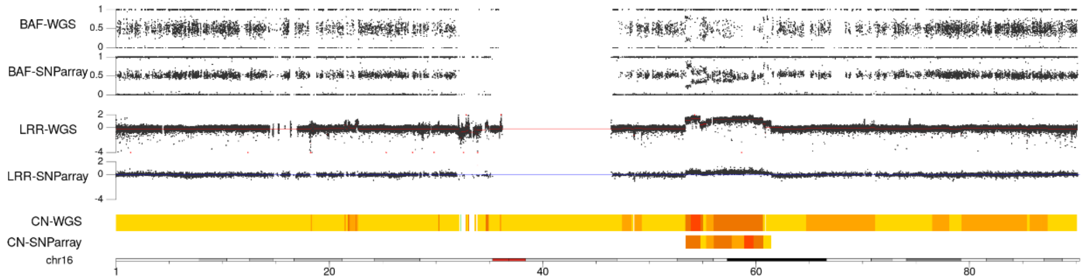


STS-26T

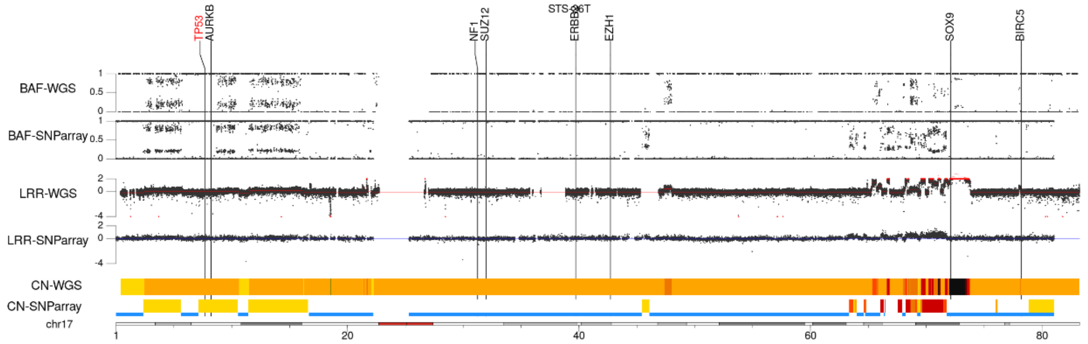


STS-26T

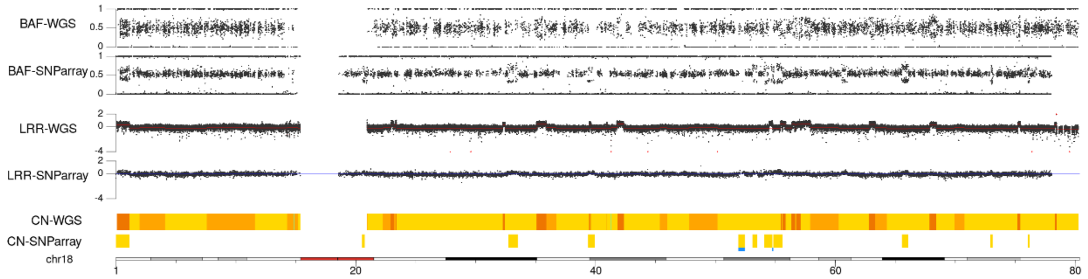


STS-26T

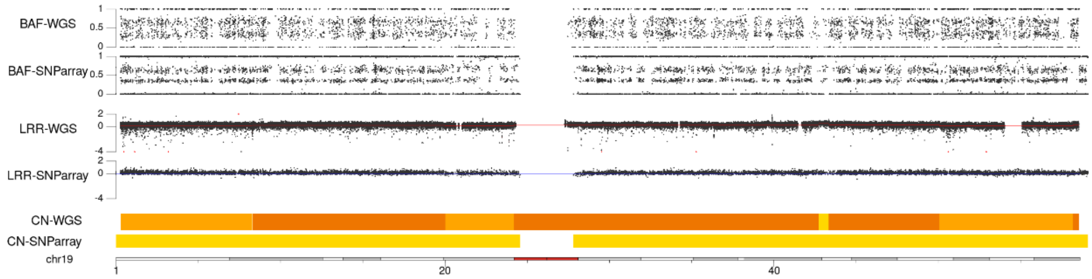

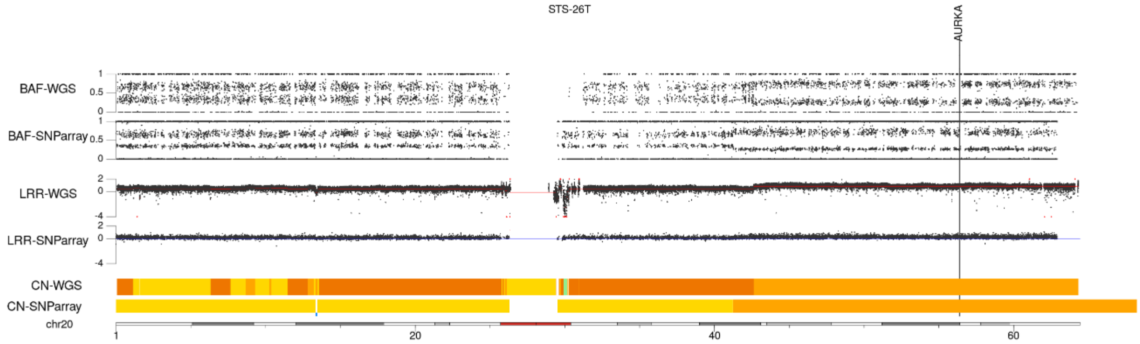
STS-26T
AURKA
BAF-WGS
BAF-SNParray
LRR-WGS
LRR-SNParray
CN-WGS
CN-SNParray
chr20
1
20
40
60

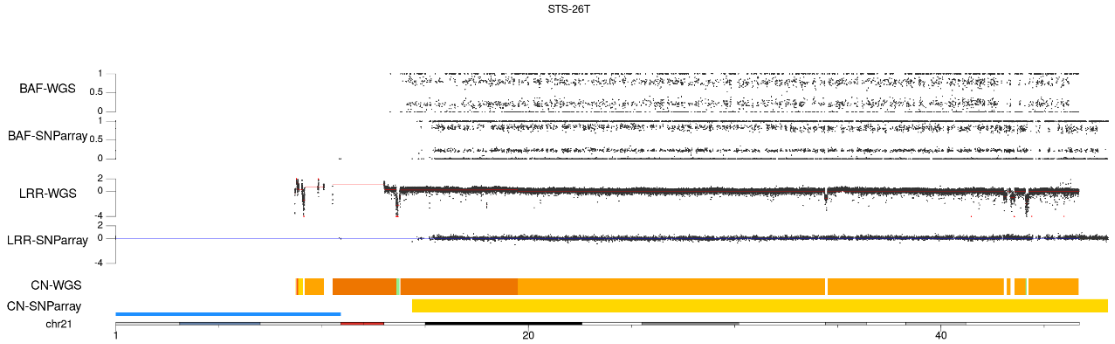
STS-26T
BAF-WGS
BAF-SNParray
LRR-WGS
LRR-SNParray
CN-WGS
CN-SNParray
chr21
1
20
40

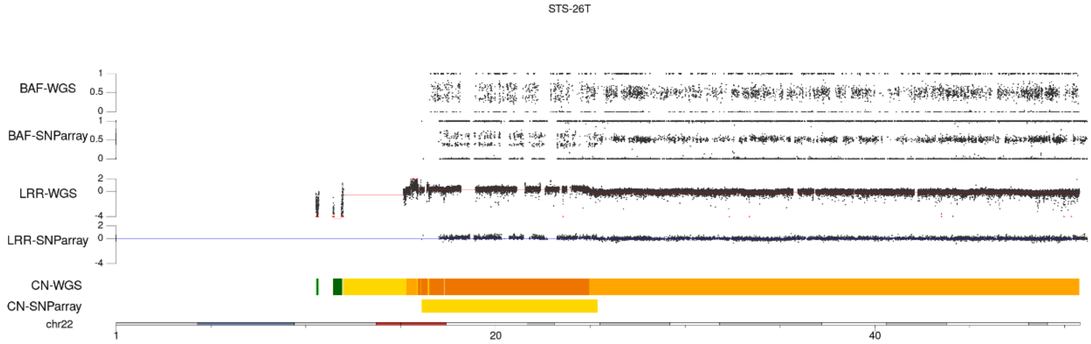
STS-26T
BAF-WGS
BAF-SNParray
LRR-WGS
LRR-SNParray
CN-WGS
CN-SNParray
chr22
1
20
40

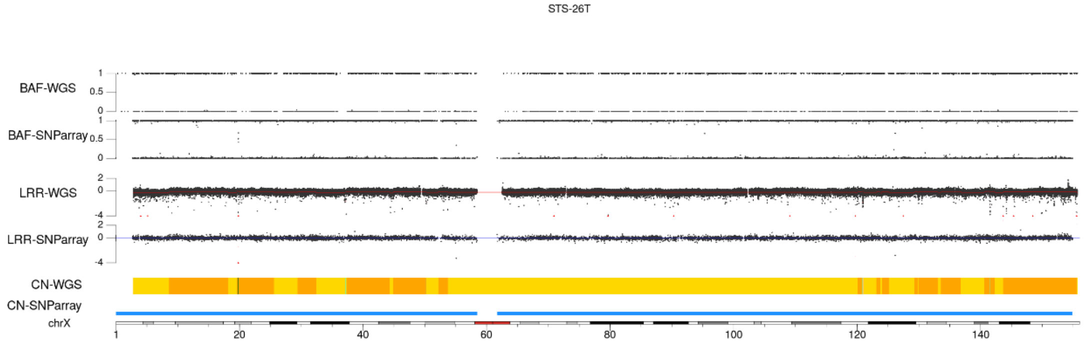
STS-26T
BAF-WGS
BAF-SNParray
LRR-WGS
LRR-SNParray
CN-WGS
CN-SNParray
chrX
1
20
40
60
80
100
120
140

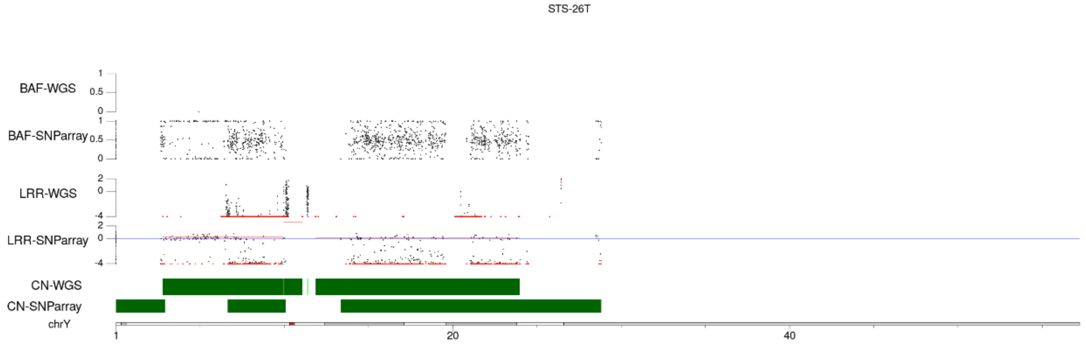
STS-26T
BAF-WGS
BAF-SNParray
LRR-WGS
LRR-SNParray
CN-WGS
CN-SNParray
chrY
1
20
40

#### HS-Sch-2

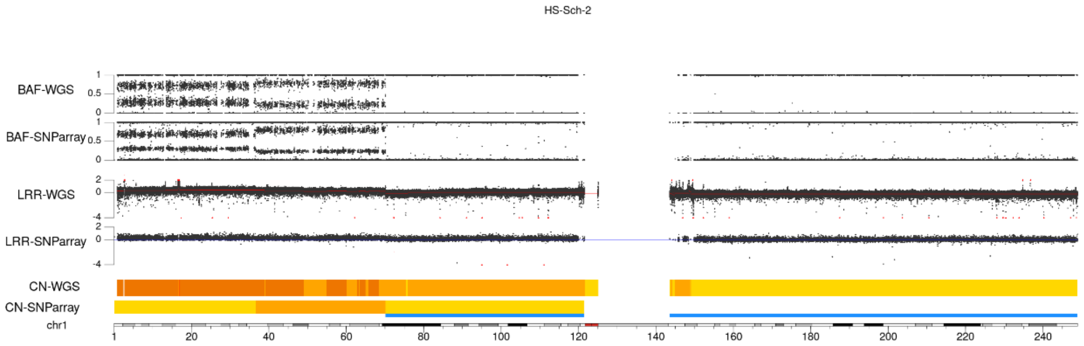
HS-Sch-2
BAF-WGS
BAF-SNParray
LRR-WGS
LRR-SNParray
CN-WGS
CN-SNParray
chr1
1 20 40 60 80 100 120 140 160 180 200 220 240

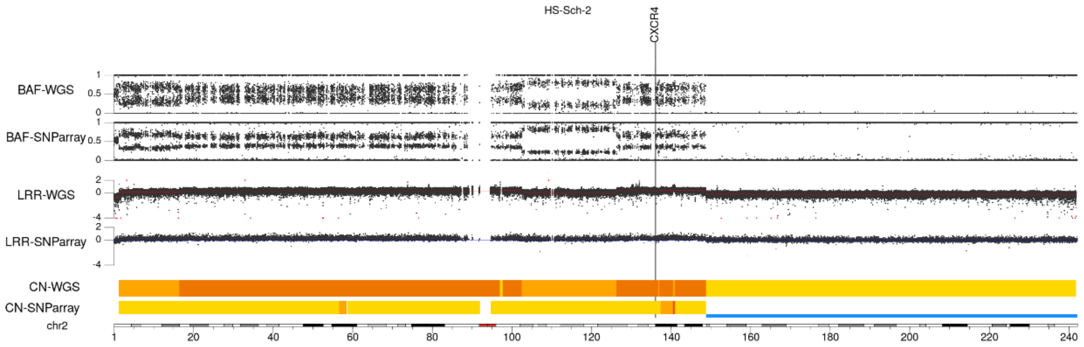
HS-Sch-2
CXCR4
BAF-WGS
BAF-SNParray
LRR-WGS
LRR-SNParray
CN-WGS
CN-SNParray
chr2
1 20 40 60 80 100 120 140 160 180 200 220 240

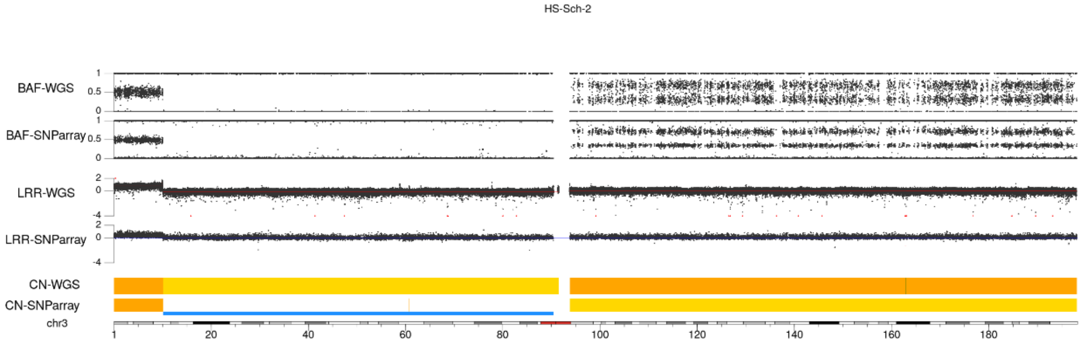
HS-Sch-2
BAF-WGS
BAF-SNParray
LRR-WGS
LRR-SNParray
CN-WGS
CN-SNParray
chr3
1 20 40 60 80 100 120 140 160 180

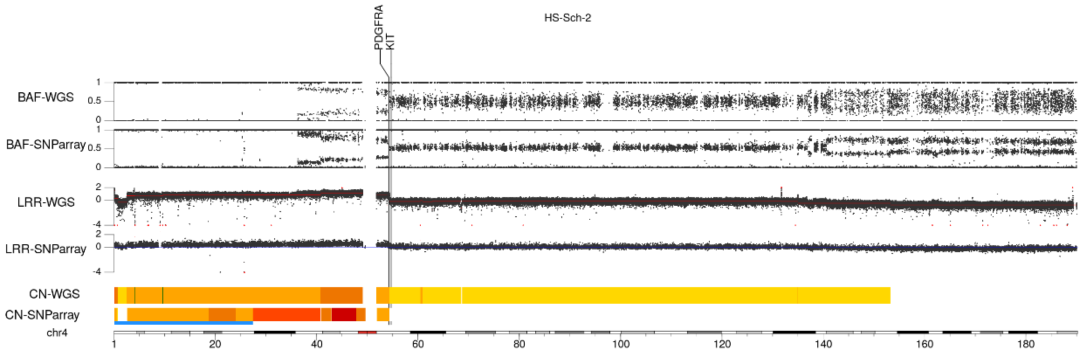
HS-Sch-2
PDGFRA
KIT
BAF-WGS
BAF-SNParray
LRR-WGS
LRR-SNParray
CN-WGS
CN-SNParray
chr4
1 20 40 60 80 100 120 140 160 180

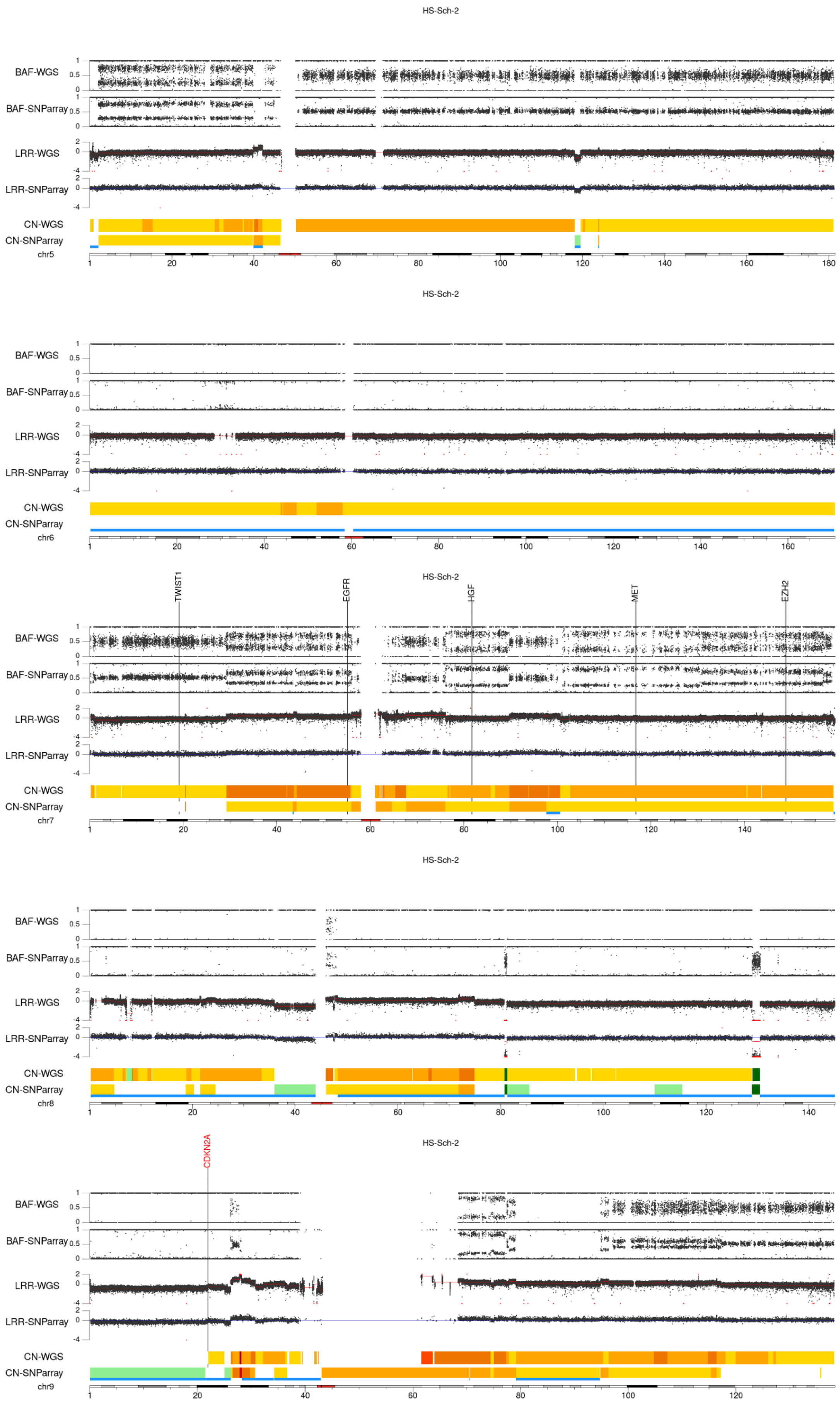
HS-Sch-2
BAF-WGS
BAF-SNParray
LRR-WGS
LRR-SNParray
CN-WGS
CN-SNParray
chr5
1 20 40 60 80 100 120 140 160 180
HS-Sch-2
BAF-WGS
BAF-SNParray
LRR-WGS
LRR-SNParray
CN-WGS
CN-SNParray
chr6
1 20 40 60 80 100 120 140 160
HS-Sch-2
TWIST1
EGFR
HGF
MET
EZH2
BAF-WGS
BAF-SNParray
LRR-WGS
LRR-SNParray
CN-WGS
CN-SNParray
chr7
1 20 40 60 80 100 120 140
HS-Sch-2
BAF-WGS
BAF-SNParray
LRR-WGS
LRR-SNParray
CN-WGS
CN-SNParray
chr8
1 20 40 60 80 100 120 140
HS-Sch-2
CDKN2A
BAF-WGS
BAF-SNParray
LRR-WGS
LRR-SNParray
CN-WGS
CN-SNParray
chr9
1 20 40 60 80 100 120

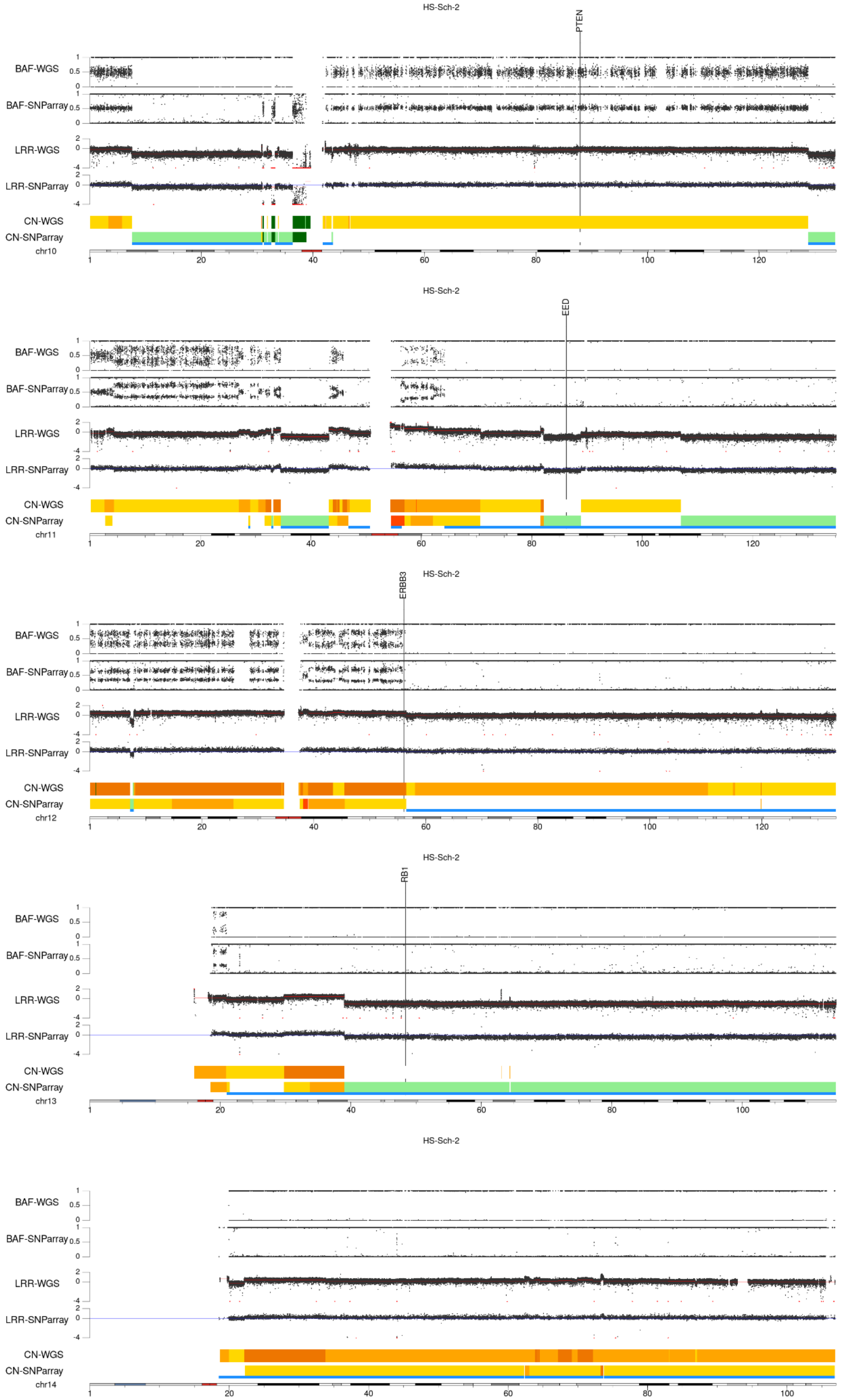
HS-Sch-2
PTEN
BAF-WGS
BAF-SNParray
LRR-WGS
LRR-SNParray
CN-WGS
CN-SNParray
chr10
HS-Sch-2
EED
chr11
HS-Sch-2
ERBB3
chr12
HS-Sch-2
RB1
chr13
HS-Sch-2
chr14

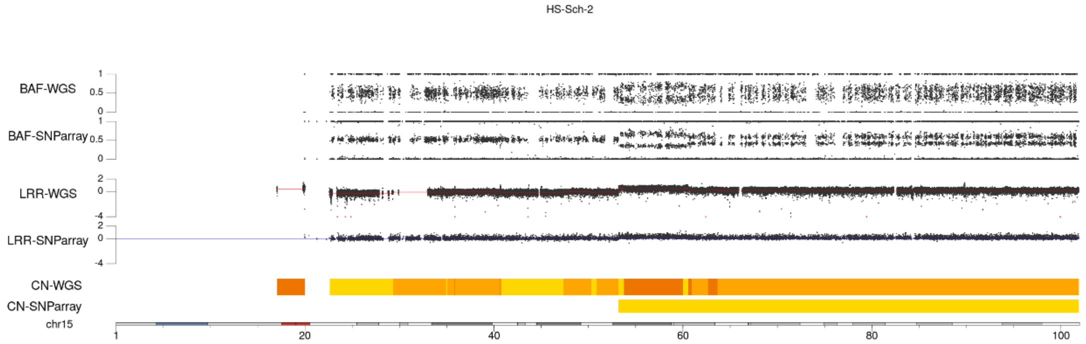
HS-Sch-2
BAF-WGS
BAF-SNParray
LRR-WGS
LRR-SNParray
CN-WGS
CN-SNParray
chr15
1
20
40
60
80
100

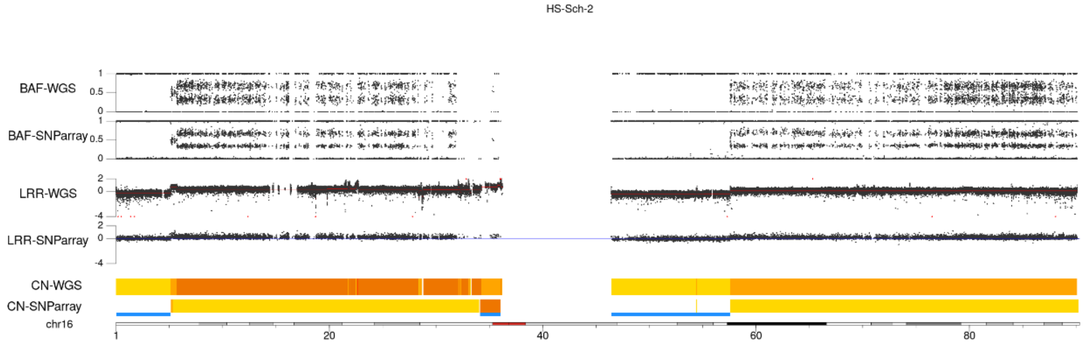
HS-Sch-2
BAF-WGS
BAF-SNParray
LRR-WGS
LRR-SNParray
CN-WGS
CN-SNParray
chr16
1
20
40
60
80

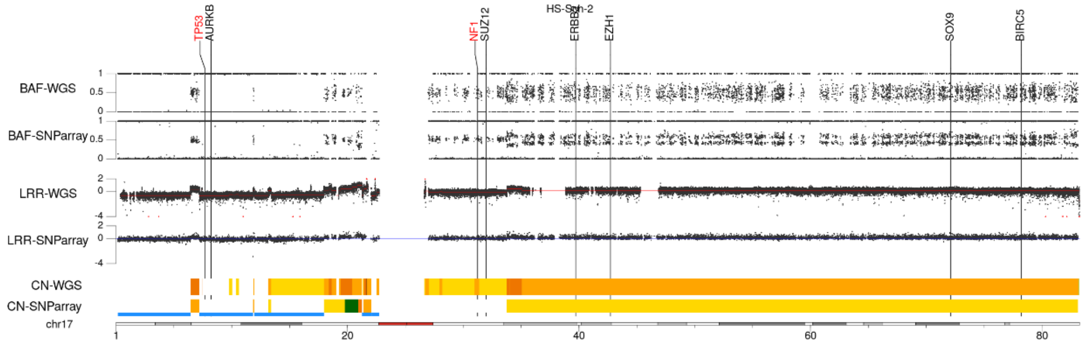
HS-Sch-2
TP53
AURKB
NF1
SUZ12
ERBB2
EZH1
SOX9
BIRC5
BAF-WGS
BAF-SNParray
LRR-WGS
LRR-SNParray
CN-WGS
CN-SNParray
chr17
1
20
40
60
80

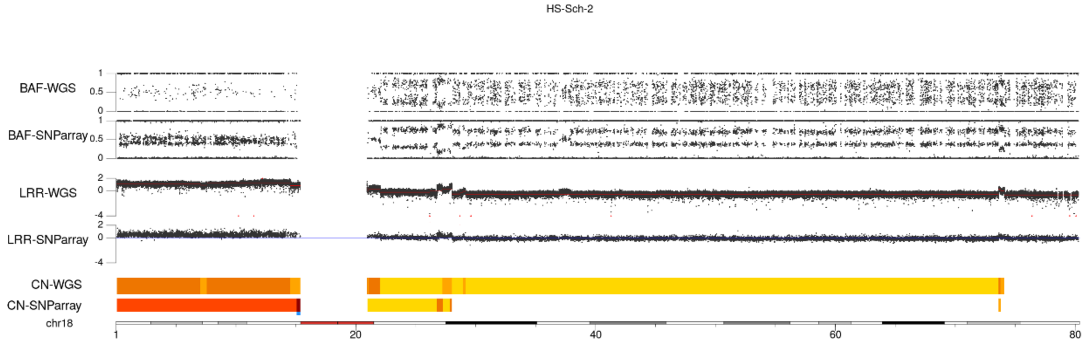
HS-Sch-2
BAF-WGS
BAF-SNParray
LRR-WGS
LRR-SNParray
CN-WGS
CN-SNParray
chr18
1
20
40
60
80

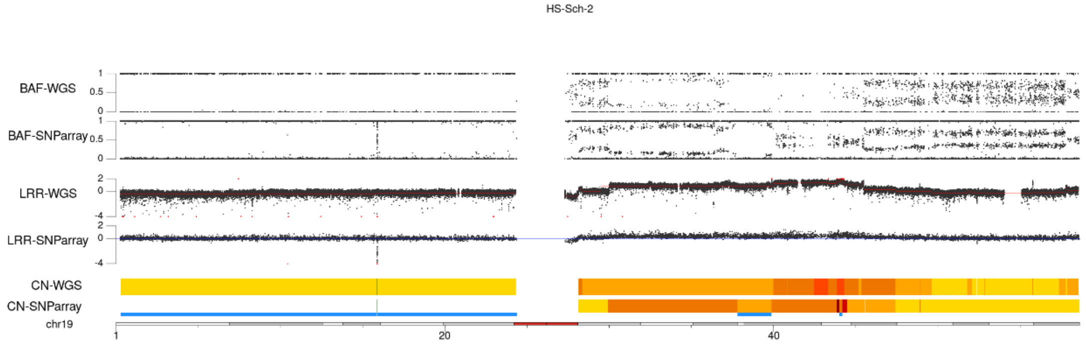
HS-Sch-2
BAF-WGS
BAF-SNParray
LRR-WGS
LRR-SNParray
CN-WGS
CN-SNParray
chr19
1
20
40

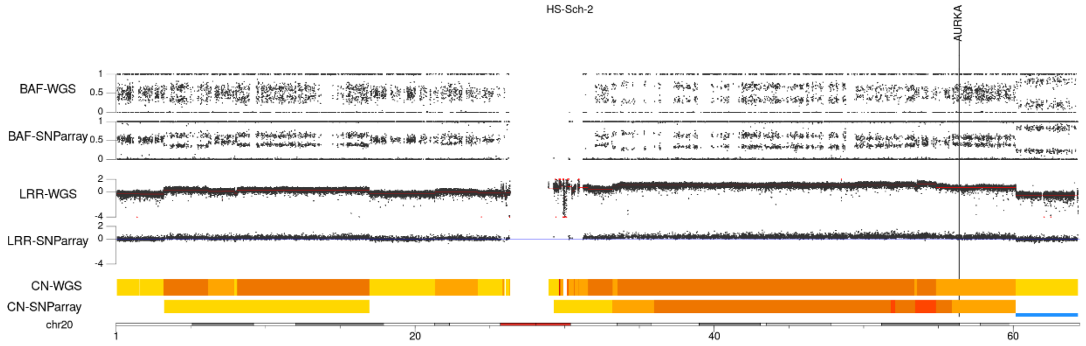
HS-Sch-2
AURKA
BAF-WGS
BAF-SNParray
LRR-WGS
LRR-SNParray
CN-WGS
CN-SNParray
chr20
1
20
40
60

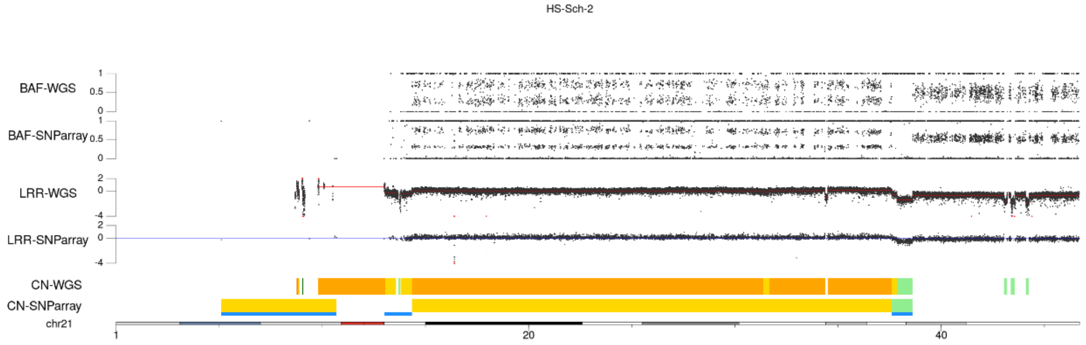
HS-Sch-2
BAF-WGS
BAF-SNParray
LRR-WGS
LRR-SNParray
CN-WGS
CN-SNParray
chr21
1
20
40

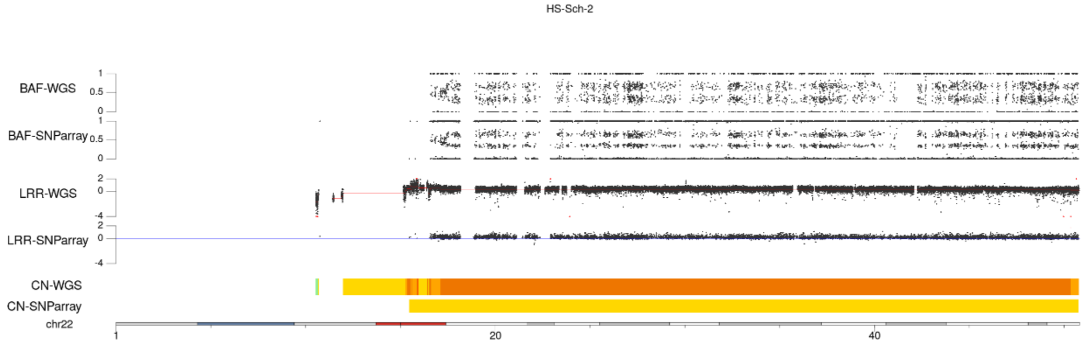
HS-Sch-2
BAF-WGS
BAF-SNParray
LRR-WGS
LRR-SNParray
CN-WGS
CN-SNParray
chr22
1
20
40

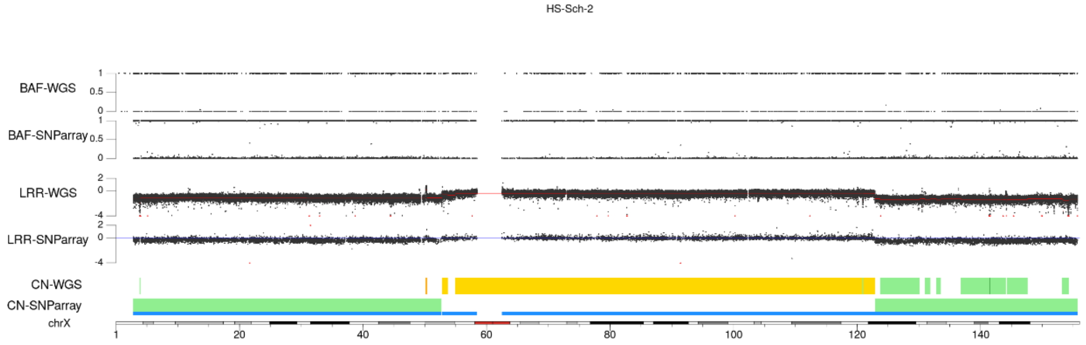
HS-Sch-2
BAF-WGS
BAF-SNParray
LRR-WGS
LRR-SNParray
CN-WGS
CN-SNParray
chrX
1
20
40
60
80
100
120
140

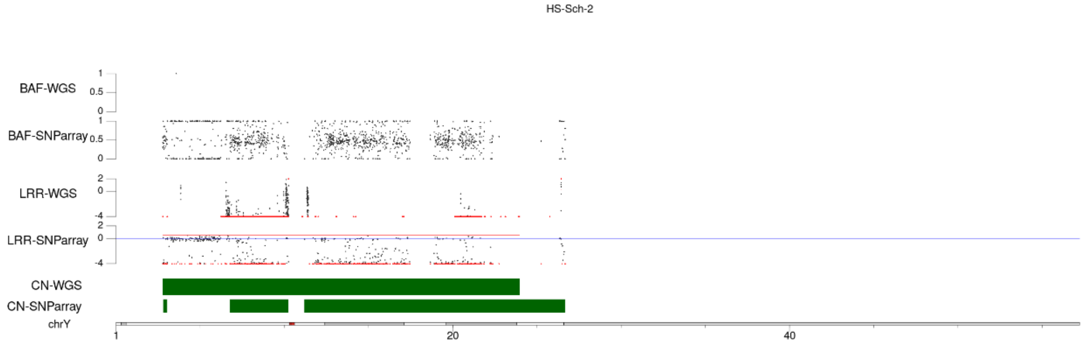
HS-Sch-2
BAF-WGS
BAF-SNParray
LRR-WGS
LRR-SNParray
CN-WGS
CN-SNParray
chrY
1
20
40

#### HS-PSS

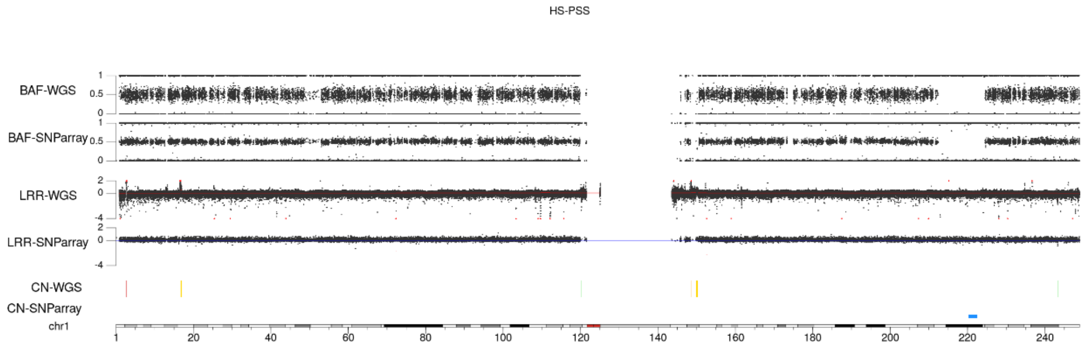
HS-PSS
BAF-WGS
BAF-SNParray
LRR-WGS
LRR-SNParray
CN-WGS
CN-SNParray
chr1
1 20 40 60 80 100 120 140 160 180 200 220 240

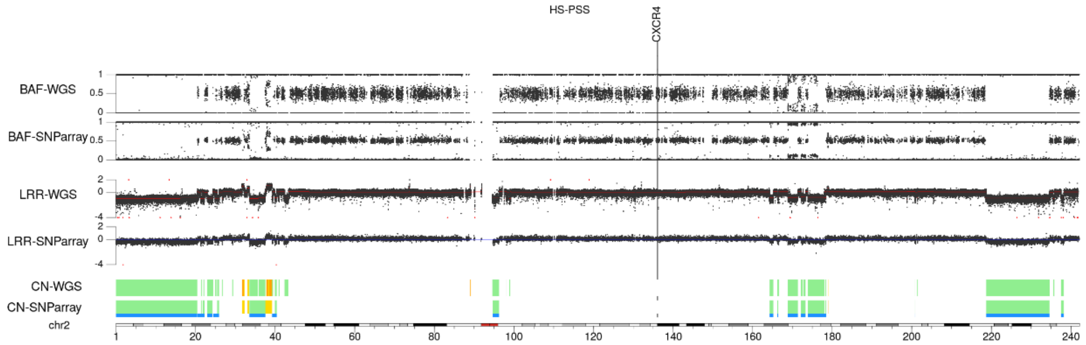
HS-PSS
CXCR4
BAF-WGS
BAF-SNParray
LRR-WGS
LRR-SNParray
CN-WGS
CN-SNParray
chr2
1 20 40 60 80 100 120 140 160 180 200 220 240

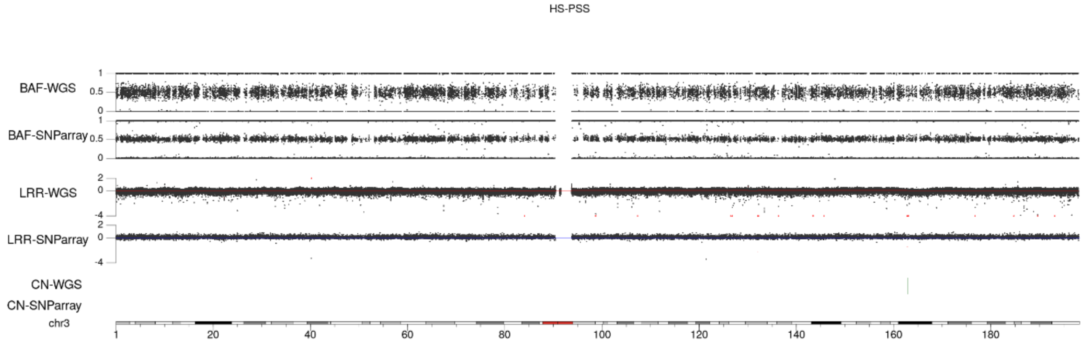
HS-PSS
BAF-WGS
BAF-SNParray
LRR-WGS
LRR-SNParray
CN-WGS
CN-SNParray
chr3
1 20 40 60 80 100 120 140 160 180

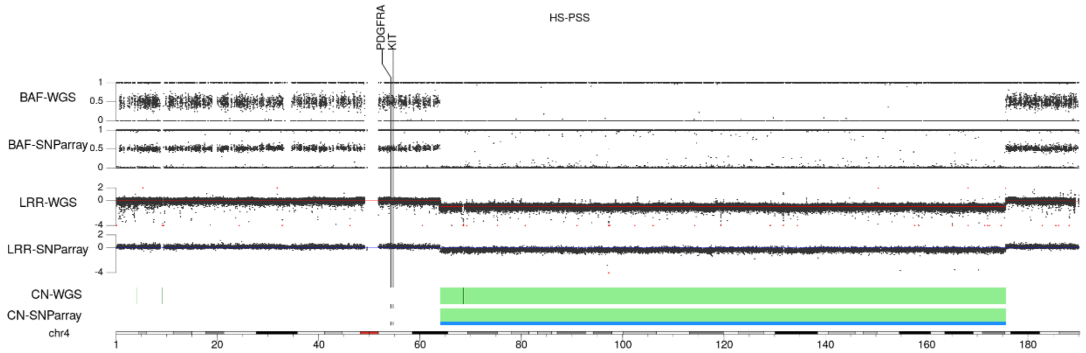
HS-PSS
PDGFRA
KIT
BAF-WGS
BAF-SNParray
LRR-WGS
LRR-SNParray
CN-WGS
CN-SNParray
chr4
1 20 40 60 80 100 120 140 160 180

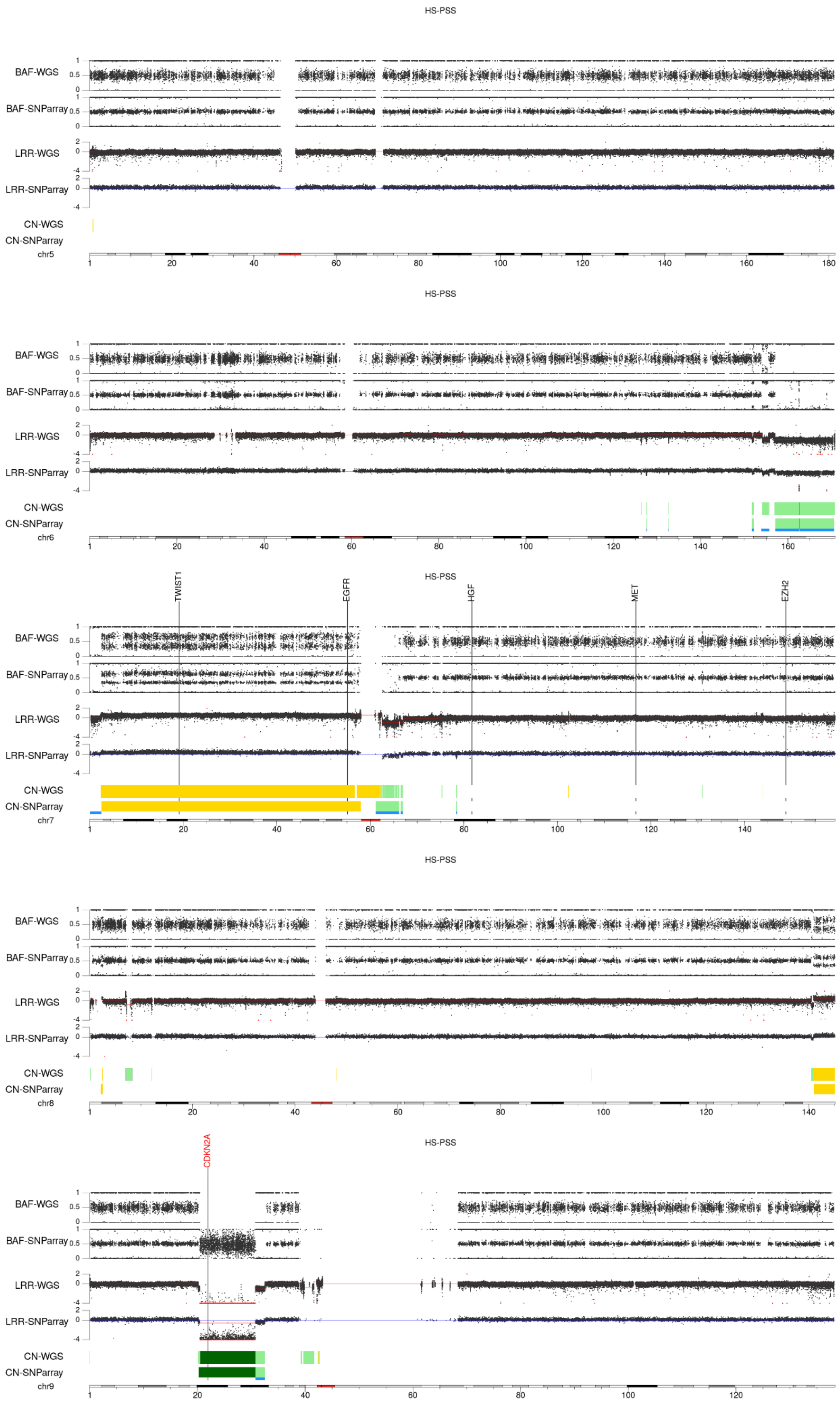
HS-PSS
BAF-WGS
BAF-SNParray
LRR-WGS
LRR-SNParray
CN-WGS
CN-SNParray
chr5
HS-PSS
BAF-WGS
BAF-SNParray
LRR-WGS
LRR-SNParray
CN-WGS
CN-SNParray
chr6
HS-PSS
TWIST1
EGFR
HGF
MET
EZH2
BAF-WGS
BAF-SNParray
LRR-WGS
LRR-SNParray
CN-WGS
CN-SNParray
chr7
HS-PSS
BAF-WGS
BAF-SNParray
LRR-WGS
LRR-SNParray
CN-WGS
CN-SNParray
chr8
HS-PSS
CDKN2A
BAF-WGS
BAF-SNParray
LRR-WGS
LRR-SNParray
CN-WGS
CN-SNParray
chr9

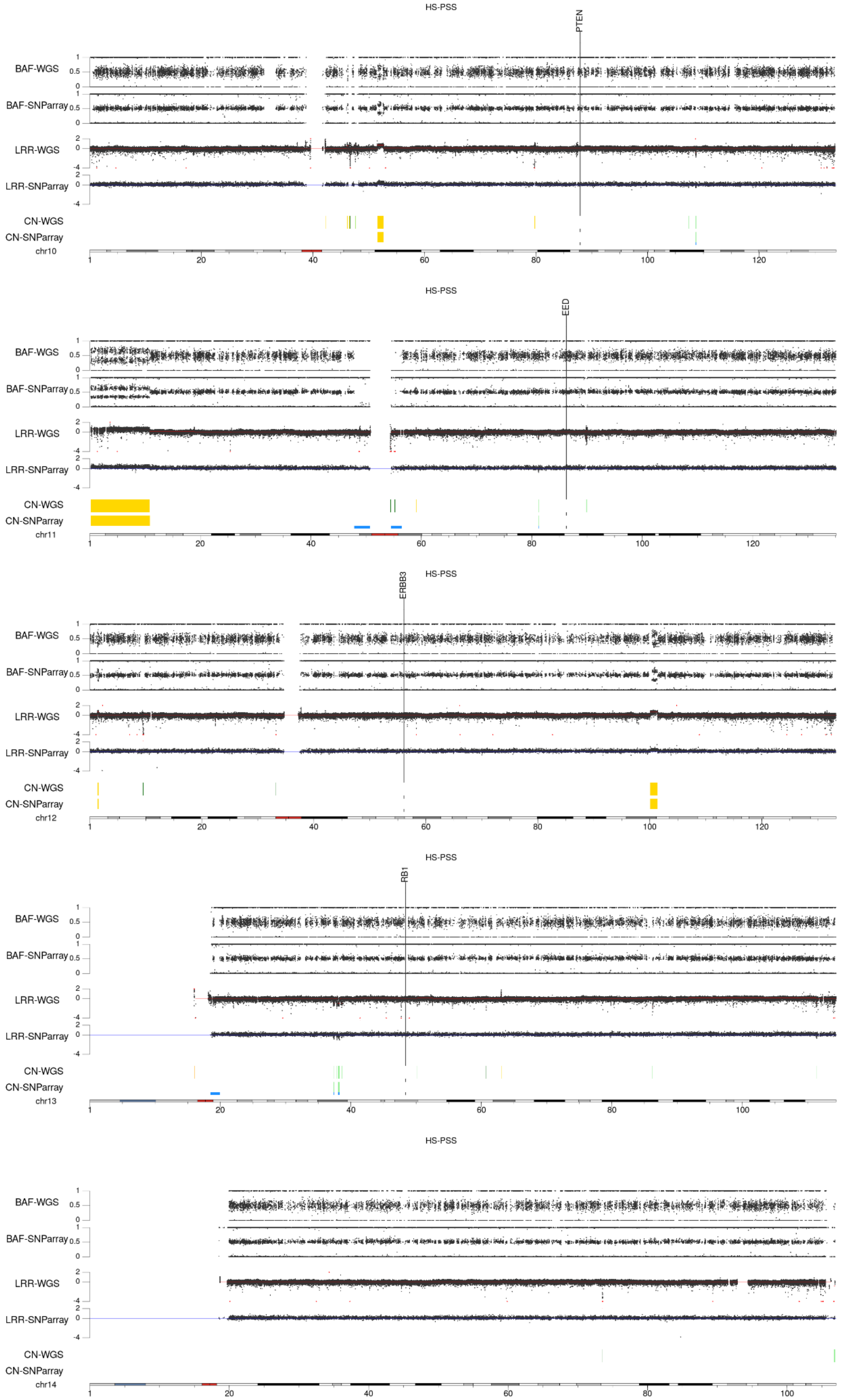
HS-PSS
PTEN
BAF-WGS
BAF-SNParray
LRR-WGS
LRR-SNParray
CN-WGS
CN-SNParray
chr10
HS-PSS
EED
chr11
HS-PSS
ERBB3
chr12
HS-PSS
RB1
chr13
HS-PSS
chr14

HS-PSS

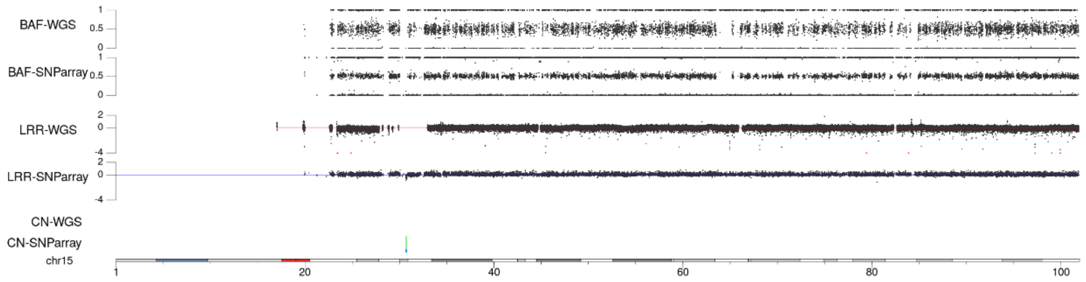


HS-PSS

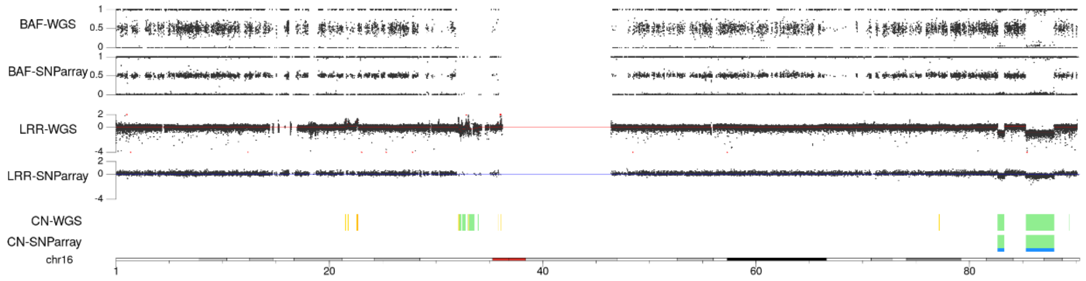


HS-PSS

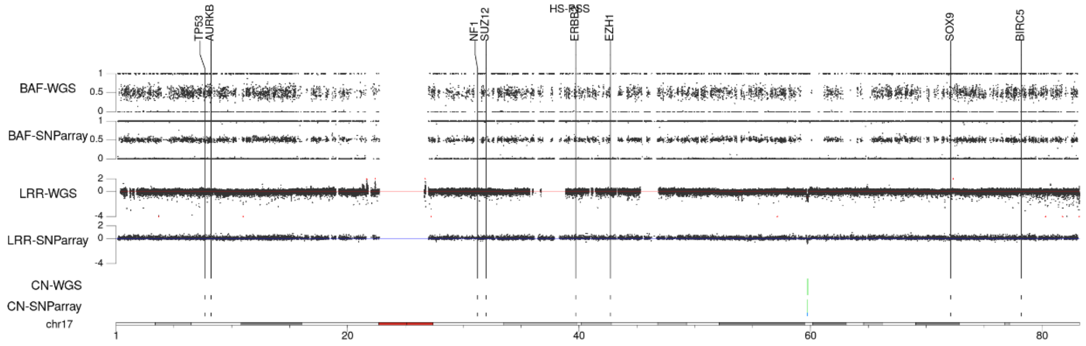


HS-PSS

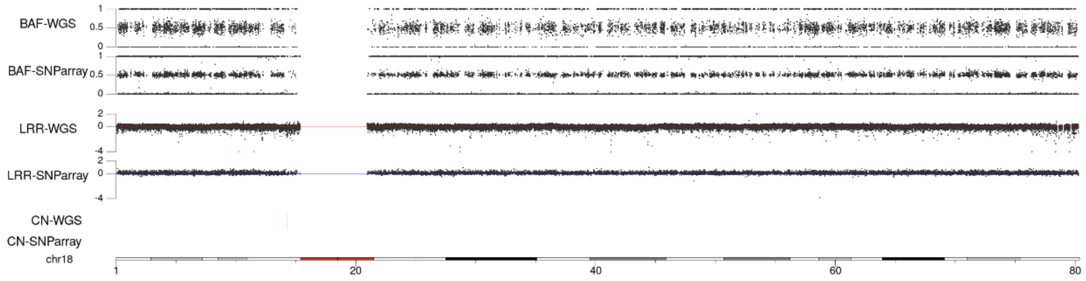


HS-PSS

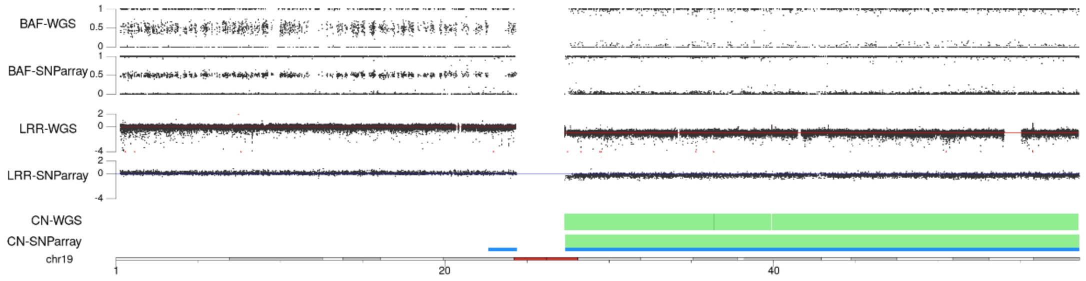

HS-PSS

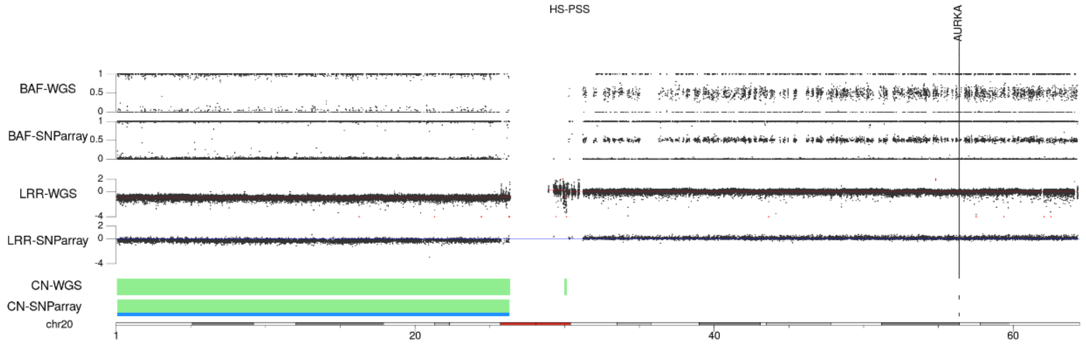


HS-PSS

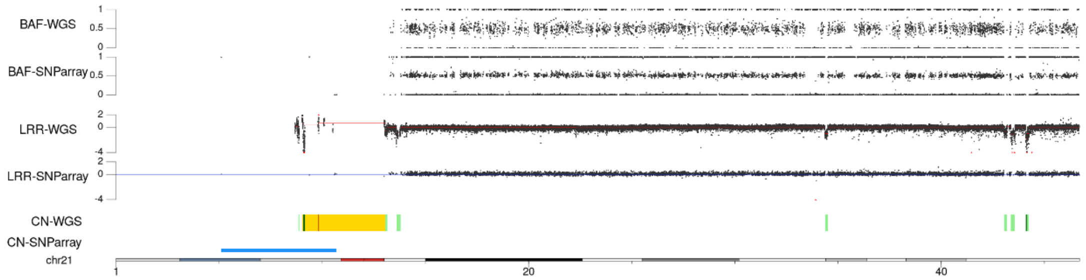


HS-PSS

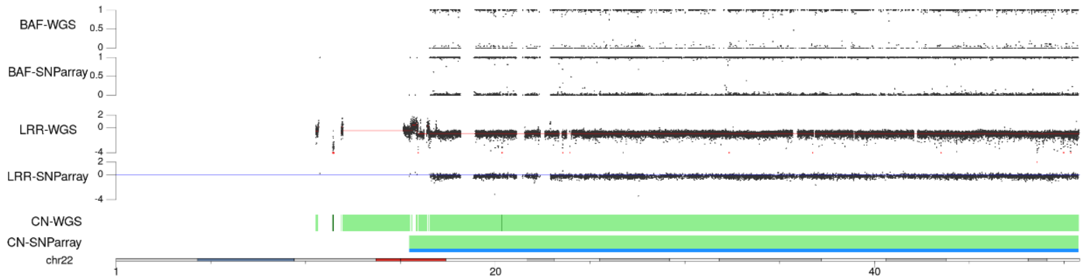


HS-PSS

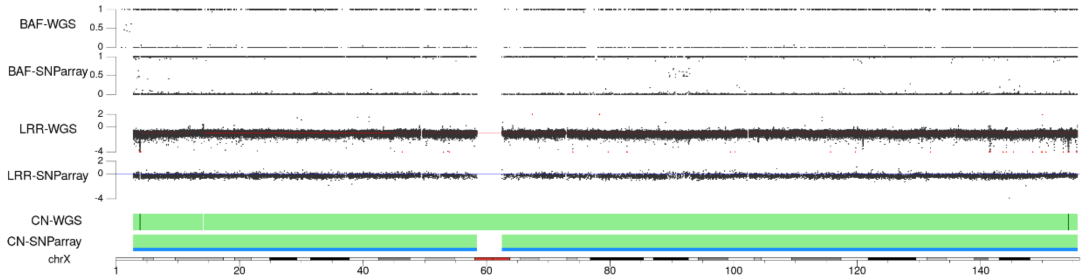


HS-PSS

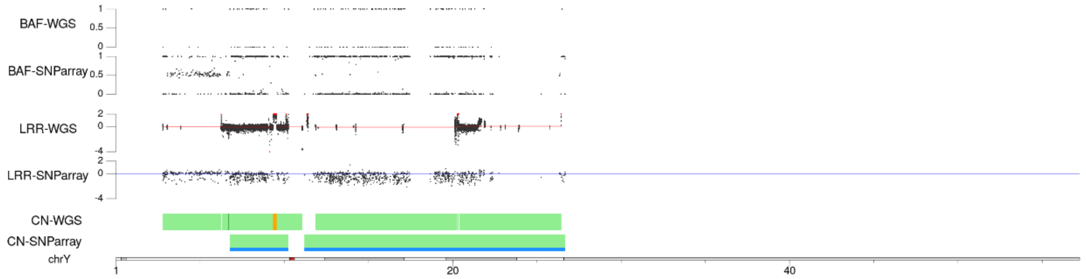

### Supplementary document 5:
Co-evolution examples by SNP-array profile analysis.

By comparing the genomic profiles of different batches of the same cell lines obtained from distinct labs we show that, despite being generally stable, the genomes of MPNST cell lines accumulate changes over time.

**Figure 1** shows the samples analyzed to illustrate two cases of co-evolution of the same cell lines that have been expanded and preserved in independent labs.

The genomic profiles of these two cell lines, ST88-14 and S462, can be seen in **Figure 2** and **Figure 3** respectively. In both figures, a purple shadow highlights the regions with a different genomic profile among the different samples cell batches analyzed.

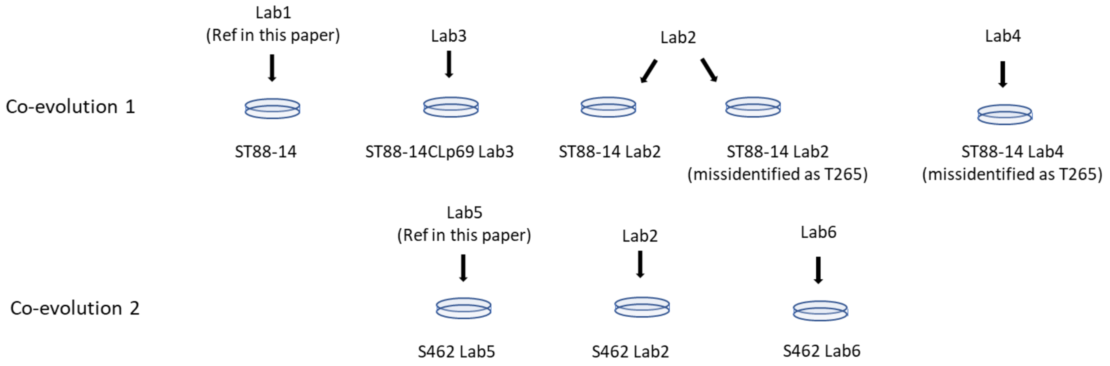


***Figure 1**. Scheme explaining the different samples analyzed by SNP-array to illustrate co-evolution of cell lines expanded and preserved in independent labs.*

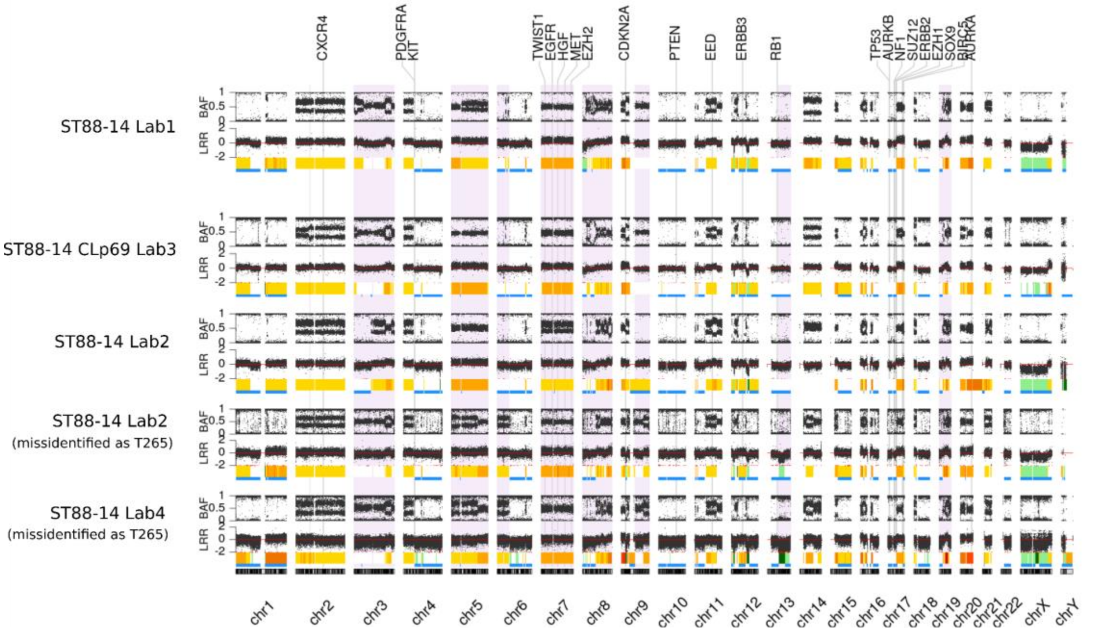


***Figure 2**. SNP-array profiles show the co-evolution of the ST88-14 cell line in samples from different laboratories. A purple shadow highlights the differences between them*

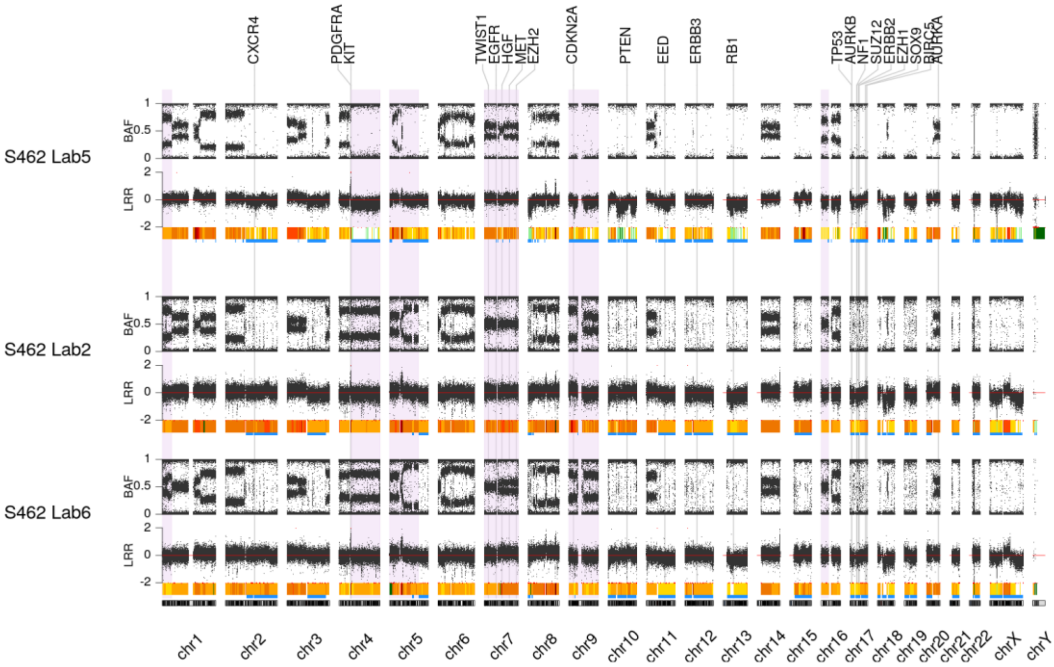


***Figure 3**. SNP-array profiles show the co-evolution of the S462 cell line in samples from different laboratories. A purple shadow highlights the differences between them*

### Supplementary Document 6: Examples of rearrangements affecting MPNST-related genes

#### Breakpoint detection using Integrative Genomic Viewer (IGV)

Integrative Genomic Viewer [1] (IGV) allow us to visualize NGS data mapped against a reference genome. In IGV, we can see the different alteration patterns created by different structural variants (SV). Genomic breakpoints cause a drastic drop in coverage and the presence of soft-clipped reads showing that part of the reads do not match the reference genome at that position. Rearrangements also produce a discordant mapping of reads in a read pair, where each read maps on different chromosomes or unexpected locations in the same chromosome. These discordant read pairs are represented by different colors, so the presence of colored reads close to the breakpoints reinforce the breakpoint identification and provide information about the translocation partner (**Figure 1**).

In the following sections, we show some breakpoints originated from inter-chromosomal rearrangements affecting different genes on different MPNSTs cell lines.

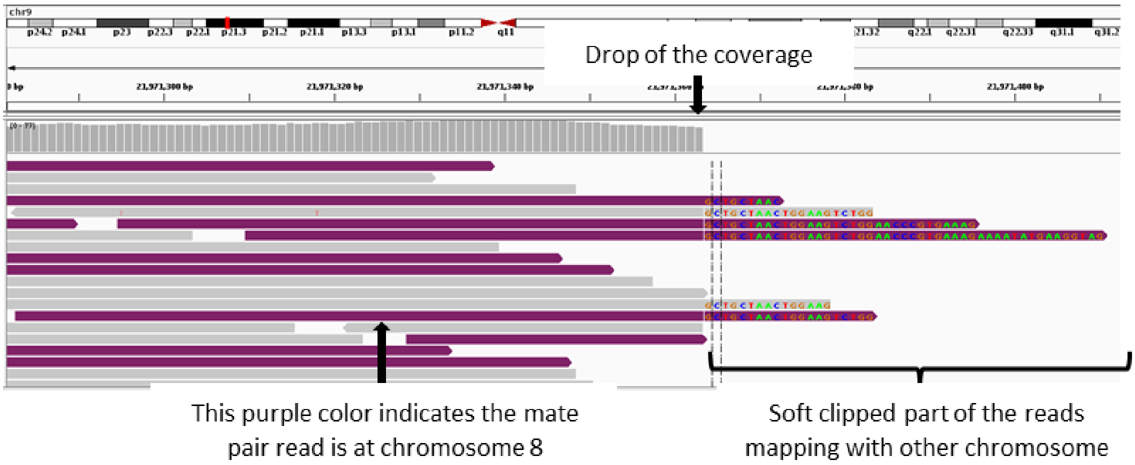


***Figure 1.*** *Screenshot from IGV showing a genomic breakpoint. Annotations point to the effects of breakpoints: coverage drop, soft-clipped reads and colored reads.*

### NMS-2

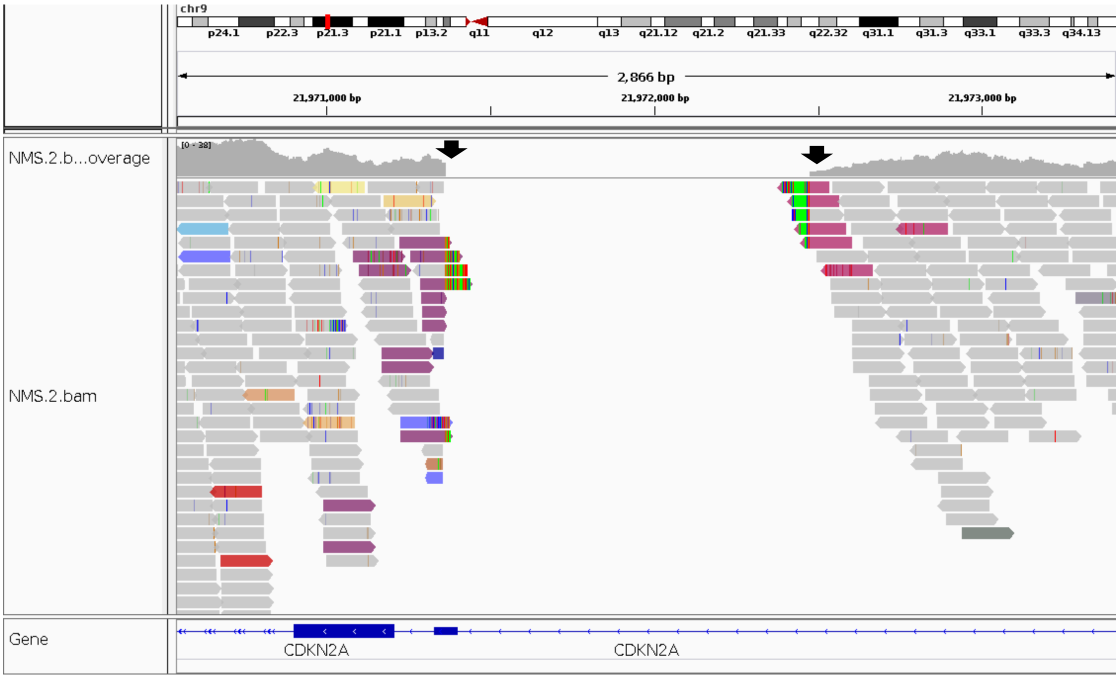


***Figure 2**. Screenshot from IGV showing the breakpoints affecting CDKN2A in NMS-2. Black arrows indicate the positions of two translocation breakpoints, one with chromosome 8 and one with chromosome 20.*

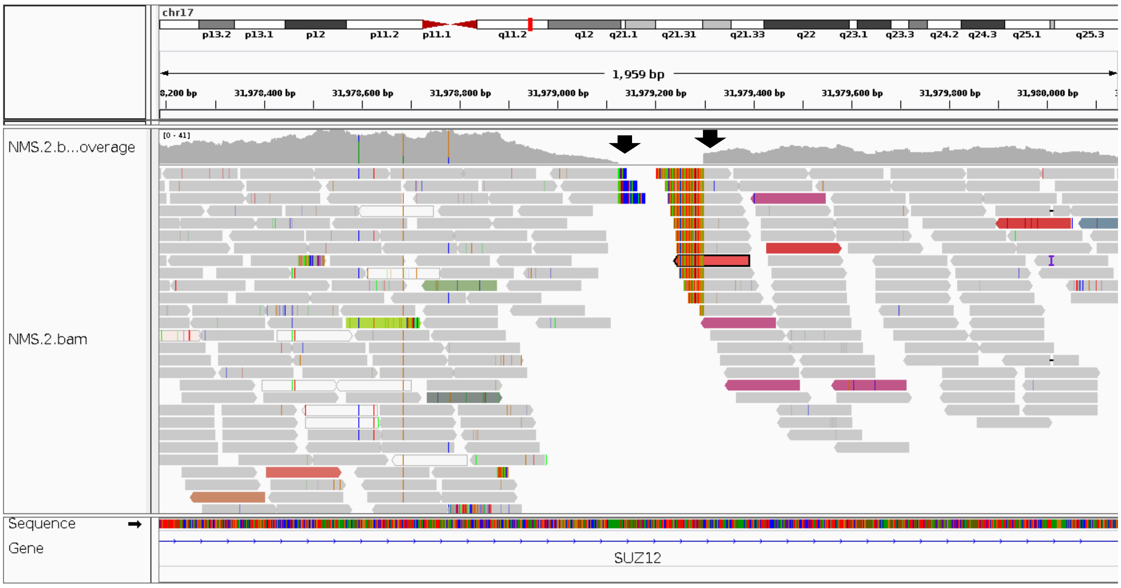


***Figure 3**. Screenshot from IGV showing a translocation of SUZ12 with a telomeric region in NMS-2. Black arrows indicate breakpoint positions*

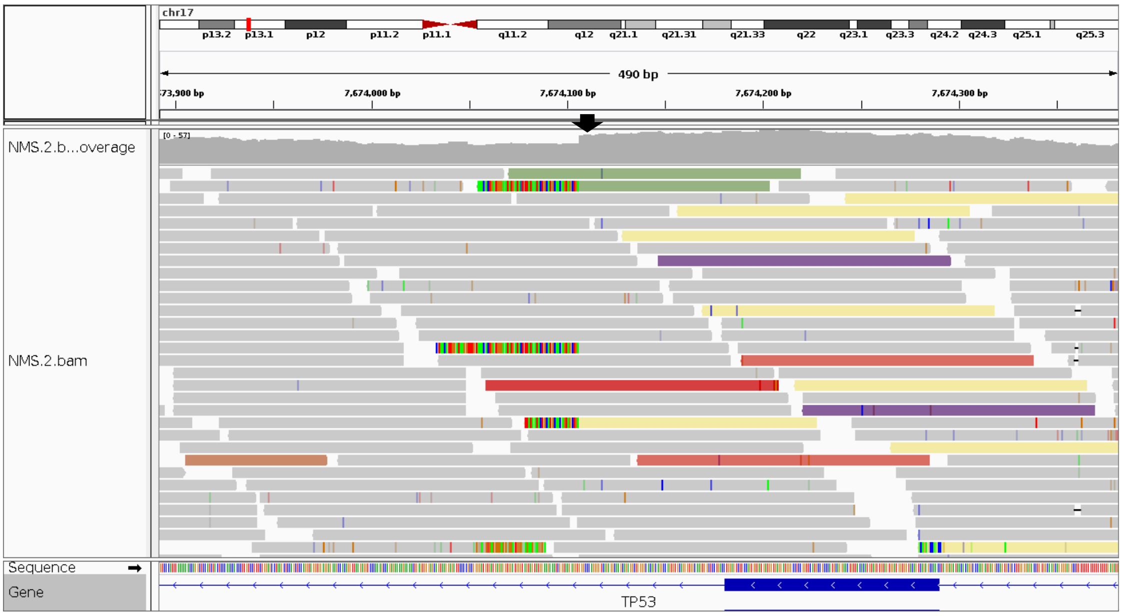


***Figure 4**. Screenshot from IGV showing the breakpoints of a translocation between* TP53 *and chromosome 4 in NMS-2. Black arrow indicates the breakpoint position.*

## ST88-14

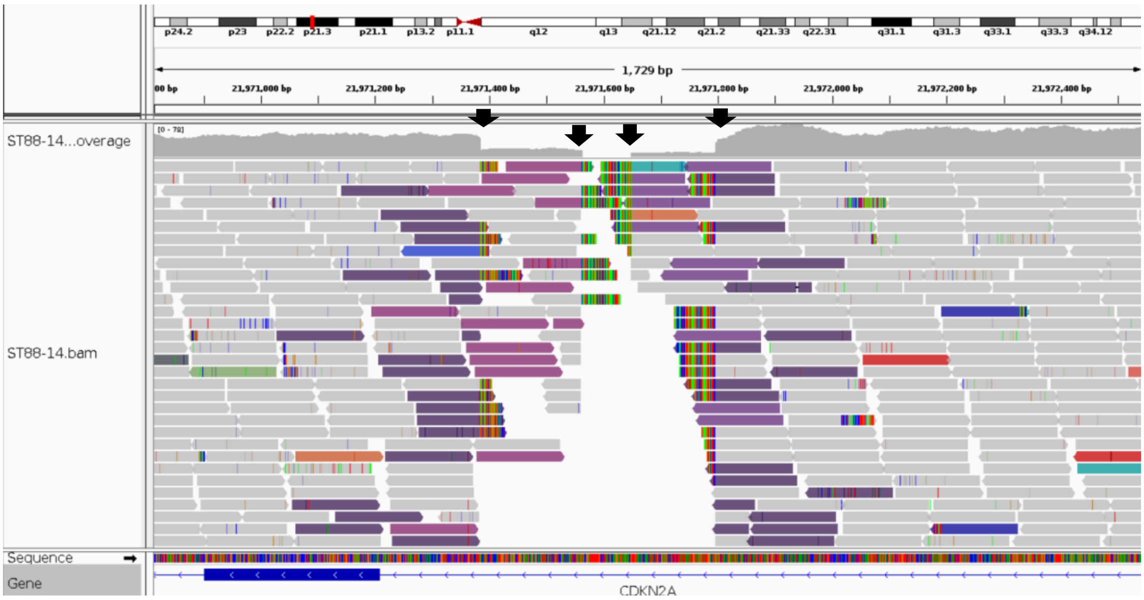


***Figure 5**. Screenshot from IGV showing multiple breakpoints close to exon 2 of CDKN2A in ST88-14. Black arrows indicate breakpoint positions. This is a complex rearrangement involving chromosomes 9, 17 and 19.*

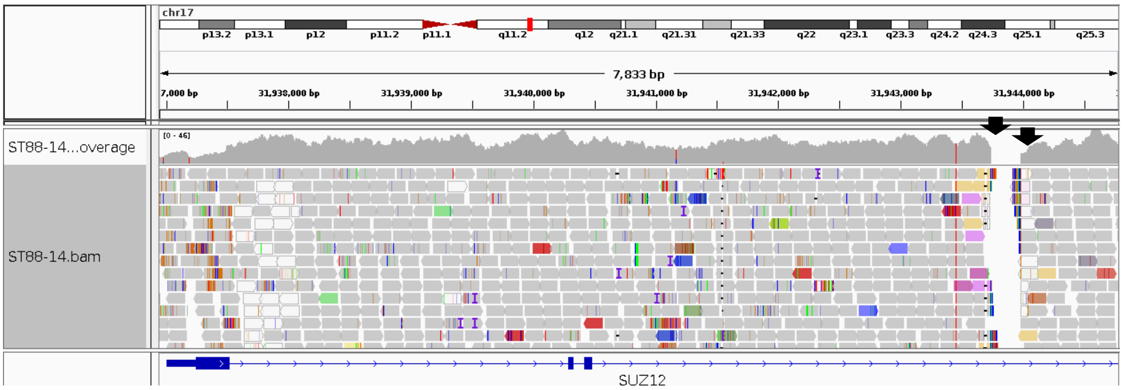


***Figure 6***. *Screenshot from IGV showing the translocation breakpoints affecting SUZ12 in ST88-14. Black arrows indicate breakpoint positions.*

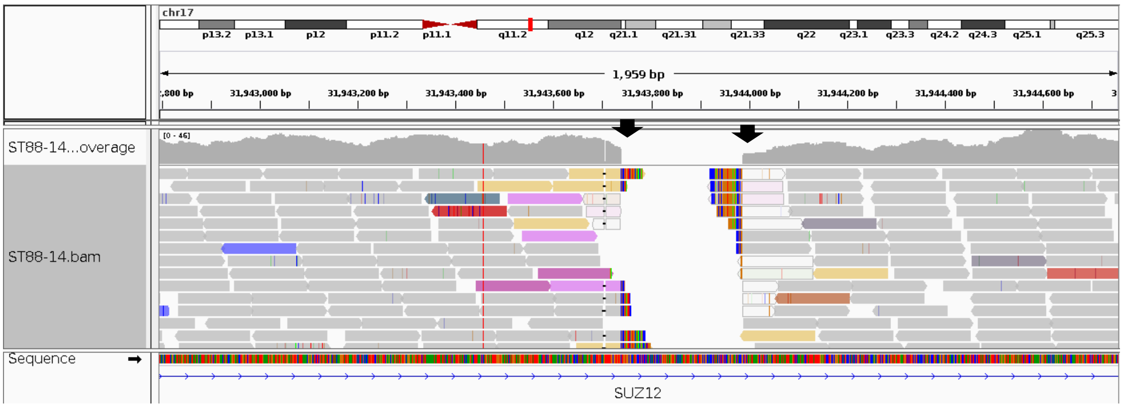


***Figure 7***. *Screenshot from IGV showing closer image of breakpoints affecting SUZ12 in ST88-14. Black arrows indicate breakpoint positions. The very light colors of reads next to the breakpoint show low-mapping quality since this breakpoint seems to involve repetitive regions in chromosome 21.*

#### Validation by PCR and Sanger sequencing of some breakpoints.

Inter-chromosomal rearrangements detected by LUMPY or CliffHunteR affecting some of the commonly altered genes in MPNST were validated by PCR and Sanger sequencing. The PCR primers, annealing temperatures and amplicon lengths are summarized in the following table:

**Table 1**. List of primers used for the breakpoint validation

| Name | Sequence (5'->3') Template | Length | Tm | Atm |
|---|---|---|---|---|
| ST8814_F_chr9-chr17_rigth_CDKN2A | CTGGATAATCTCCCTCAAGACCA | 23 | 59,03 | 58,79 |
| ST8814_R_chr9-chr17_rigth_CDKN2A | ACAGTGGACAATAAGCACTTGAA | 23 | 58,54 | |
| ST8814_F_chr9-chr8_CDKN2A | AAGCCCCAGGTGTCTAATTACC | 22 | 59,76 | 59,96 |
| ST8814_R_chr9-chr8_CDKN2A | TGCTCCATTGGTCTTTGTGACT | 22 | 60,16 | |
| NMS_2_F_chr4-chr17_TP53 | AGCATGTTAGTCAAGGTAGCCC | 22 | 60,09 | 60,03 |
| NMS_2_R_chr4-chr17_TP53 | TCCTAGGTTGGCTCTGACTGTA | 22 | 59,96 | |

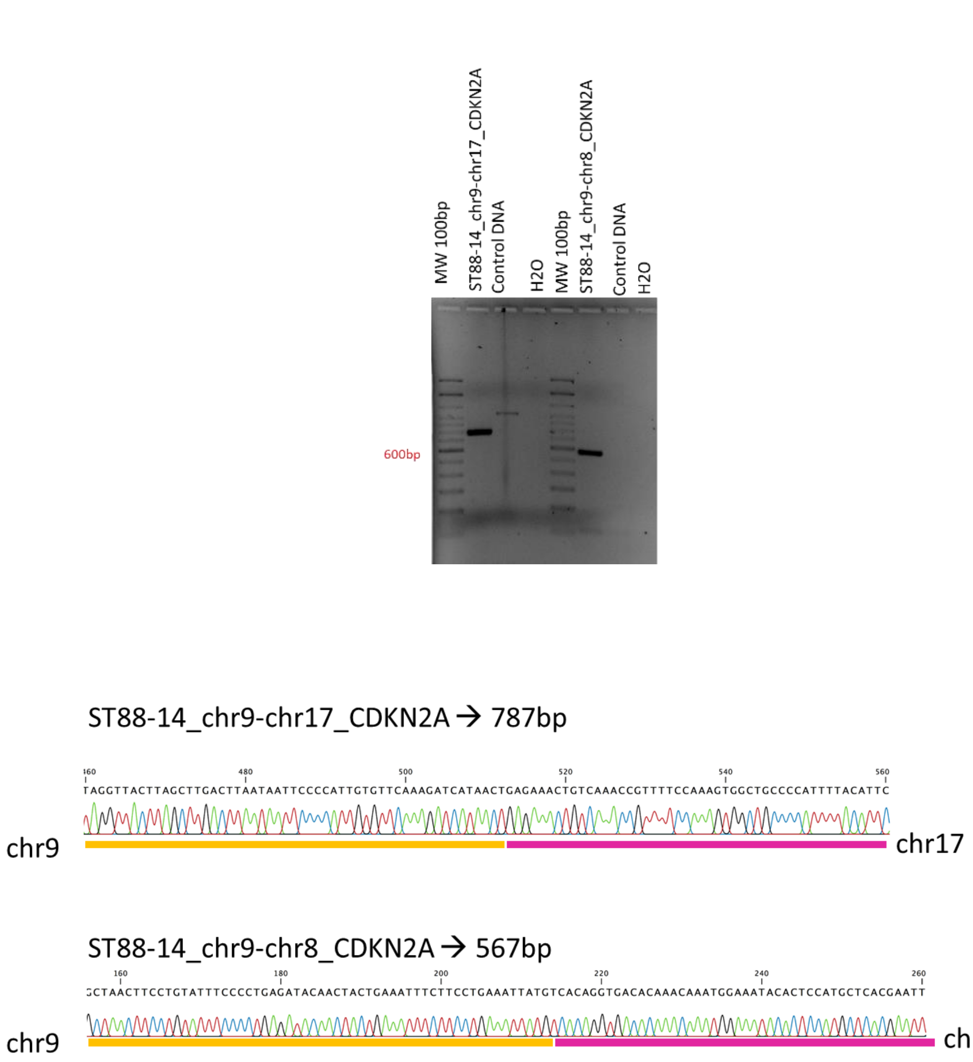


***Figure 8.*** *Validation of two of the breakpoints inactivating CDKN2A in ST88-14 cell line.**(a).** PCR validation. **(b).** Sanger sequencing breakpoint validation*

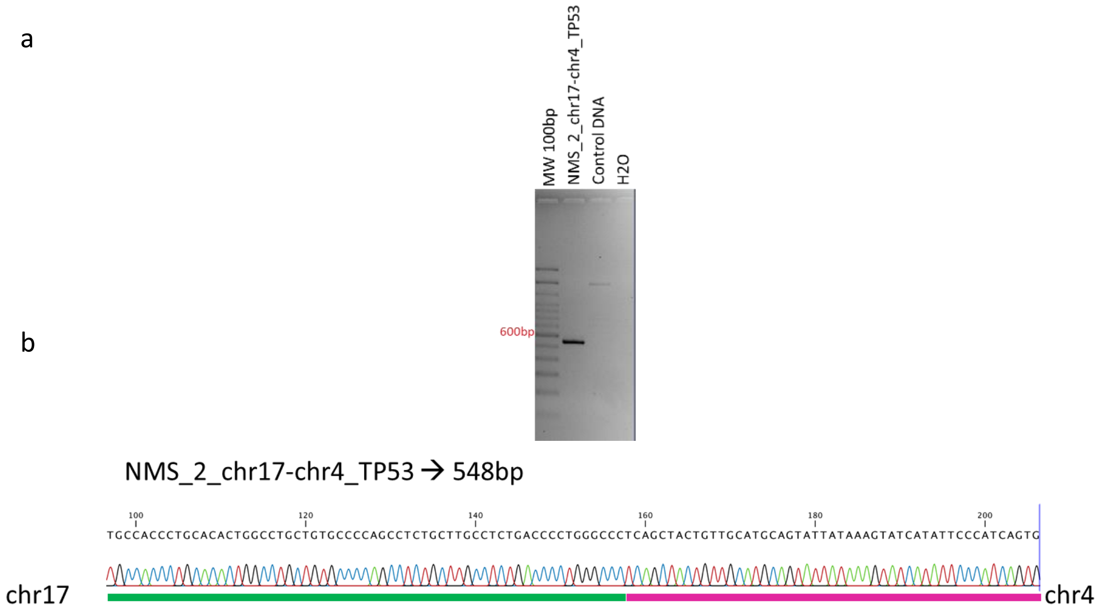


***Figure 9.*** *Validation of the breakpoint affecting TP53 in NMS-2 cell line..(a). PCR validation. (b). Sanger sequencing breakpoint validation*

### Validation by RT-PCR and Sanger sequencing of the EML4-ALK fusion gene breakpoints detected in the HS-PSS cell line.

Validation of the breakpoints detected in HS-PSS cell line by WGS data. We used the primers and conditions described previously in Takeuchi et al. (2008)[2].

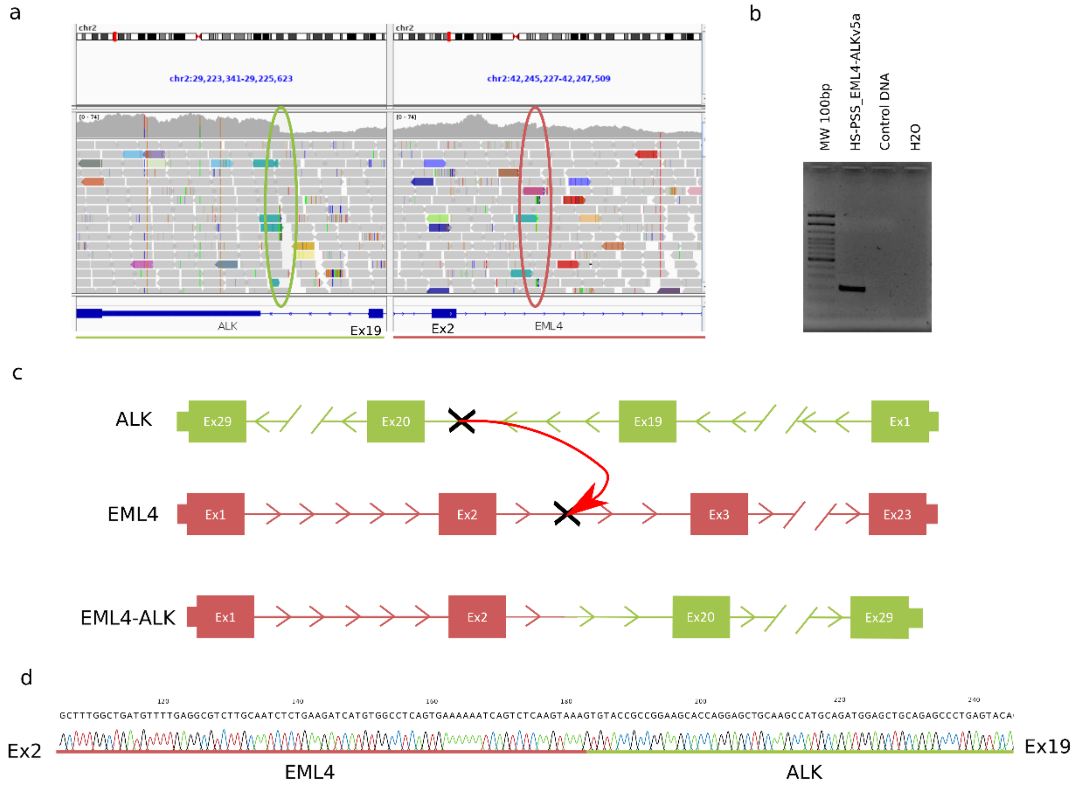


*Figure 10*. *EML4-ALK v5a validation.* ***(a)***. *IGV screenshot showing the inversion breakpoints.* ***(b)***. *RT-PCR gel confirming the amplification of the fusion gene.* ***(c)***. *Scheme of the fusion gene generated.* ***(d)***. *Sanger sequencing validating the EML4-ALK v5a fusion gene found in HS-PSS cell line.*

##### *References*

1. Robinson JT, Thorvaldsdóttir H, Winckler W, Guttman M, Lander ES, Getz G, et al. Integrative genomics viewer. Nature Biotechnology. 2011;29:24–6.

2. Takeuchi K, Choi YL, Soda M, Inamura K, Togashi Y, Hatano S, et al. Multiplex reverse transcription-PCR screening for EML4-ALK fusion transcripts. Clinical Cancer Research. 2008;14:6618–24.
