## Supplemental Table 1 for "A detailed landscape of genomic alterations in malignant peripheral nerve sheath tumor cell lines challenges the current MPNST diagnosis"

| Microsatellite | Chr. location | S462 | ST88-14 | NF90-8 | sNF96.2 | NMS-2 | STS26T | HS-Sch-2 |
|---|---|---|---|---|---|---|---|---|
| D8S1179 | 8 | 10, 12 | 14 | 10, 12 | 13 | 12, 13 | 13, 14 | 14 |
| D21S11 | 21q11.2-q21 | 29 | 29, 32.3 | 30 | 30 | 30 | 30, 31 | 29, 30 |
| D7S820 | 7q11.21-22 | 8, 10 | 8 | 11 | 10, 11 | 9, 12 | 8, 11 | 8, 11 |
| CSF1PO | 5q33.3-34 | 12 | 9, 12 | 12 | 12 | 12 | 10, 13 | 11, 13 |
| D3S1358 | 3p | 14, 17 | 15, 18 | 15 | 16 | 15, 16 | 14 | 18 |
| TH01 | 11p15.5 | 7, 8 | 9 | 6 | 6 | 6 | 6, 9.3 | 6, 7 |
| D13S317 | 13q22-31 | 12 | 12 | 8 | 10 | 12 | 9, 10 | 11 |
| D16S539 | 16q24-qter | 11, 13 | 13 | 9 | 11 | 9 | 12, 13 | 11, 12 |
| D2S1338 | 2q35-37.1 | 23 | 17, 23 | 16, 24 | 17 | 18, 19 | 20 | 20 |
| D19S433 | 19q12-13.1 | 14 | 13, 14 | 13, 16 | 13, 14 | 13.2, 14.2 | 14 | 14.2 |
| vWA | 12p12-pter | 19 | 16 | 17 | 17, 19 | 14, 19 | 17 | 15, 16 |
| TPOX | 2p23-2per | 8 | 11, 12 | 11 | 11 | 8, 9 | 8 | 8 |
| D18S51 | 18q21.3 | 16 | 12 | 14, 18 | 16 | 17 | 17, 18 | 13, 14 |
| AMEL | Xp22.1-22.3 Yp11.2 | X | X, Y | X | X | X, Y | X | X |
| D5S818 | 5q21-31 | 12 | 12, 13 | 11 | 11 | 12, 13 | 11, 12 | 13 |
| FGA | 4q28 | 20 | 21 | 25 | 22 | 21, 22 | 22, 23 | 23, 26 |

| HS-PSS |
| --- |
| 14, 15 |
| 28, 32 |
| 10, 12 |
| 10, 12 |
| 16 |
| 6, 9 |
| 8, 10 |
| 10 |
| 19 |
| 13 |
| 14, 15 |
| 8 |
| 14, 15 |
| X, Y |
| 9, 11 |
| 22 |
