## Supplemental Table 2 for "A detailed landscape of genomic alterations in malignant peripheral nerve sheath tumor cell lines challenges the current MPNST diagnosis"

### S462 cell line variants with a potential impact on protein function

| chr | start | end | Sample | Gene.refGen | REF | ALT | Func.refGene | ExonicFunc.r | VAF | avsnp150 |
|---|---|---|---|---|---|---|---|---|---|---|
| chr1 | 7736404 | 7736404 | S462 | CAMTA1 | G | T | exonic | stopgain | 0.565217391 | . |
| chr1 | 12725527 | 12725527 | S462 | AADACL3 | T | G | exonic | nonsynonym | 0.787234042 | rs7513079 |
| chr1 | 12760937 | 12760937 | S462 | C1orf158 | C | T | exonic | nonsynonym | 0.686274509 | rs1132185 |
| chr1 | 21833280 | 21833280 | S462 | HSPG2 | C | A | exonic | nonsynonym | 0.209302325 | . |
| chr1 | 26282344 | 26282362 | S462 | UBXN11 | ACCGGGACC | A | exonic | nonframeshi | 0.9 | . |
| chr1 | 26337258 | 26337258 | S462 | CRYBG2 | G | A | exonic | nonsynonym | 0.275862068 | rs146453215 |
| chr1 | 26345087 | 26345087 | S462 | CRYBG2 | A | G | exonic | nonsynonym | 0.535714285 | rs11579835 |
| chr1 | 34904984 | 34904984 | S462 | DLGAP3 | C | CTTG | exonic | nonframeshi | 0.272727272 | . |
| chr1 | 59638370 | 59638370 | S462 | FGGY | G | GT | exonic | frameshift_ir | 0.216216216 | . |
| chr1 | 146066541 | 146066541 | S462 | NBPF10 | T | G | exonic | nonsynonym | 0.692307692 | . |
| chr1 | 146069686 | 146069686 | S462 | NBPF10 | C | T | exonic | nonsynonym | 0.4375 | . |
| chr1 | 148972321 | 148972321 | S462 | PDE4DIP | C | T | exonic | nonsynonym | 0.44 | rs1628172 |
| chr1 | 149029862 | 149029862 | S462 | PDE4DIP | A | G | exonic | nonsynonym | 0.44 | . |
| chr1 | 213997621 | 213997621 | S462 | PROX1 | A | C | exonic | nonsynonym | 0.205882352 | . |
| chr1 | 235342705 | 235342705 | S462 | GGPS1 | G | A | exonic | nonsynonym | 0.224137931 | . |
| chr1 | 241671141 | 241671141 | S462 | WDR64 | T | C | exonic | nonsynonym | 0.195652173 | . |
| chr2 | 63882649 | 63882649 | S462 | UGP2 | G | A | exonic | nonsynonym | 0.2 | . |
| chr2 | 112573788 | 112573788 | S462 | POLR1B | C | T | exonic | nonsynonym | 0.977777777 | . |
| chr2 | 127220161 | 127220161 | S462 | CYP27C1 | A | G | exonic | nonsynonym | 0.333333333 | rs950006231 |
| chr2 | 205740533 | 205740533 | S462 | NRP2 | C | A | exonic | nonsynonym | 0.290322580 | . |
| chr2 | 219235754 | 219235754 | S462 | ANKZF1 | G | A | exonic | nonsynonym | 1 | rs201069890 |
| chr2 | 221482675 | 221482675 | S462 | EPHA4 | G | A | exonic | nonsynonym | 0.166666666 | . |
| chr3 | 13483535 | 13483535 | S462 | HDAC11 | C | T | exonic | nonsynonym | 0.172413793 | . |
| chr3 | 37007053 | 37007053 | S462 | MLH1 | C | T | exonic | nonsynonym | 0.410714285 | . |
| chr3 | 46709583 | 46709589 | S462 | TMIE | TAAGAAG | TAAG | exonic | nonframeshi | 1 | . |
| chr3 | 111112316 | 111112316 | S462 | NECTIN3 | A | C | exonic | nonsynonym | 0.290322580 | . |
| chr3 | 197511986 | 197511986 | S462 | BDH1 | C | T | exonic | nonsynonym | 0.2 | rs144840608 |
| chr4 | 575941 | 575941 | S462 | TMEM271 | T | C | exonic | nonsynonym | 0.6 | . |
| chr4 | 9173181 | 9173182 | S462 | FAM90A26 | AT | A | exonic | frameshift_d | 1 | . |
| chr4 | 87615729 | 87615765 | S462 | DSPP | AATAGCAGT | AACAGCAGT | exonic | nonframeshi | 1 | . |
| chr4 | 133152506 | 133152506 | S462 | PCDH10 | T | C | exonic | nonsynonym | 0.785714285 | . |
| chr5 | 16670821 | 16670821 | S462 | MYO10 | C | T | exonic | nonsynonym | 0.764705882 | rs563993750 |
| chr5 | 37195860 | 37195860 | S462 | CPLANE1 | A | G | exonic | nonsynonym | 0.796296296 | . |
| chr5 | 55954959 | 55954959 | S462 | IL6ST | G | C | exonic | nonsynonym | 0.652173913 | rs140433866 |

| | | | | | | | | | | |
|---|---|---|---|---|---|---|---|---|---|---|
| chr5 | 56914919 | 56914931 | S462 | SETD9 | ACTGAAGCA( | A | exonic | nonframeshi | 0.494252873 | . |
| chr5 | 82304657 | 82304657 | S462 | ATP6AP1L | G | T | exonic | stopgain | 0.311111111 | . |
| chr5 | 82304658 | 82304658 | S462 | ATP6AP1L | A | C | exonic | nonsynonym | 0.318181818 | . |
| chr5 | 135009435 | 135009435 | S462 | CATSPER3 | A | G | exonic | nonsynonym | 1 | rs142494932 |
| chr5 | 141673686 | 141673686 | S462 | ARAP3 | G | A | exonic | nonsynonym | 0.25 | . |
| chr5 | 149374580 | 149374580 | S462 | IL17B | C | T | exonic | nonsynonym | 0.962962962 | rs373011468 |
| chr6 | 32530201 | 32530201 | S462 | HLA-DRB5 | T | TCCAG | exonic | frameshift_ir | 0.478260869 | . |
| chr6 | 82366051 | 82366051 | S462 | TPBG | G | T | exonic | nonsynonym | 0.212121212 | . |
| chr6 | 116925507 | 116925507 | S462 | RFX6 | G | C | exonic | nonsynonym | 0.673913043 | rs146115506 |
| chr6 | 144531050 | 144531050 | S462 | UTRN | A | G | splicing | . | 0.322580645 | . |
| chr6 | 170319057 | 170319057 | S462 | FAM120B | A | T | exonic | nonsynonym | 0.269230769 | . |
| chr7 | 100893830 | 100893830 | S462 | ACHE | G | C | exonic | nonsynonym | 0.426229508 | rs17885778 |
| chr7 | 138621107 | 138621107 | S462 | SVOPL | C | A | exonic | nonsynonym | 0.634146341 | rs142327276 |
| chr7 | 140674264 | 140674264 | S462 | ADCK2 | G | C | splicing | . | 0.492753623 | rs140400082 |
| chr7 | 151195597 | 151195598 | S462 | IQCA1L | TC | T | exonic | frameshift_d | 1 | . |
| chr7 | 152144739 | 152144739 | S462 | KMT2C | C | T | exonic | nonsynonym | 0.236842105 | . |
| chr8 | 8377352 | 8377358 | S462 | PRAG1 | CGCCGCT | C | exonic | nonframeshi | 1 | . |
| chr8 | 25366878 | 25366878 | S462 | DOCK5 | C | A | exonic | nonsynonym | 0.288461538 | . |
| chr8 | 43315905 | 43315905 | S462 | POTEA | C | A | exonic | nonsynonym | 0.224489795 | . |
| chr8 | 53250839 | 53250839 | S462 | OPRK1 | A | G | exonic | nonsynonym | 0.314285714 | . |
| chr8 | 56068532 | 56068532 | S462 | RPS20 | T | TAA | splicing | . | 1 | . |
| chr8 | 90051875 | 90051875 | S462 | DECR1 | T | G | exonic | nonsynonym | 0.307692307 | . |
| chr8 | 112301930 | 112301930 | S462 | CSMD3 | C | A | exonic | nonsynonym | 0.241379310 | . |
| chr8 | 112352468 | 112352468 | S462 | CSMD3 | T | C | exonic | nonsynonym | 0.8 | rs139786530 |
| chr8 | 117171168 | 117171176 | S462 | SLC30A8 | GGTCAGTGA | G | splicing | . | 0.227272727 | . |
| chr8 | 125049045 | 125049045 | S462 | WASHC5 | G | C | exonic | stopgain | 0.204545454 | . |
| chr8 | 144379425 | 144379425 | S462 | ADCK5 | C | A | exonic | nonsynonym | 0.714285714 | rs6599528 |
| chr9 | 41964577 | 41964577 | S462 | CNTNAP3B | A | C | exonic | nonsynonym | 1 | rs76032838 |
| chr9 | 41991627 | 41991627 | S462 | CNTNAP3B | G | A | exonic | nonsynonym | 1 | rs3739620 |
| chr9 | 41991703 | 41991703 | S462 | CNTNAP3B | G | T | exonic | nonsynonym | 1 | rs62555055 |
| chr9 | 87668645 | 87668645 | S462 | DAPK1 | G | T | exonic | nonsynonym | 1 | rs55994363 |
| chr9 | 87921255 | 87921255 | S462 | SPATA31C1 | C | A | exonic | nonsynonym | 0.358974358 | . |
| chr9 | 91409678 | 91409679 | S462 | NFIL3 | AG | A | exonic | frameshift_d | 0.242424242 | . |
| chr9 | 94445142 | 94445142 | S462 | MFSD14B | G | C | exonic | nonsynonym | 1 | rs144279626 |
| chr9 | 96484604 | 96484604 | S462 | HABP4 | G | T | exonic | nonsynonym | 0.225806451 | . |
| chr9 | 97371638 | 97371644 | S462 | CCDC180 | CCACCTT | C | exonic | nonframeshi | 0.219512195 | . |

| | | | | | | | | | | |
|---|---|---|---|---|---|---|---|---|---|---|
| chr9 | 127813567 | 127813567 | S462 | FPGS | G | T | exonic | nonsynonym | 0.32 | . |
| chr9 | 137110891 | 137110892 | S462 | DPP7 | GT | G | exonic | frameshift_d | 1 | . |
| chr10 | 45000413 | 45000413 | S462 | C10orf25 | G | A | exonic | nonsynonym | 1 | . |
| chr10 | 47502343 | 47502343 | S462 | AGAP9 | T | C | exonic | nonsynonym | 1 | rs1047447 |
| chr10 | 88594678 | 88594678 | S462 | LIPJ | T | C | exonic | nonsynonym | 0.387096774 | . |
| chr10 | 93076668 | 93076668 | S462 | CYP26A1 | G | A | exonic | nonsynonym | 0.277777777 | rs763817649 |
| chr10 | 104034648 | 104034648 | S462 | COL17A1 | G | A | exonic | nonsynonym | 1 | rs144220426 |
| chr10 | 120890878 | 120890878 | S462 | WDR11 | G | A | exonic | nonsynonym | 0.133333333 | . |
| chr11 | 4587098 | 4587098 | S462 | OR52I2 | C | A | exonic | nonsynonym | 0.25 | . |
| chr11 | 5200445 | 5200445 | S462 | OR51V1 | T | C | exonic | nonsynonym | 0.324324324 | . |
| chr11 | 61776320 | 61776320 | S462 | MYRF | A | T | splicing | . | 0.296296296 | . |
| chr11 | 78658315 | 78658315 | S462 | TENM4 | C | T | exonic | nonsynonym | 0.130434782 | . |
| chr11 | 92799473 | 92799473 | S462 | FAT3 | T | C | exonic | nonsynonym | 0.333333333 | . |
| chr12 | 6940901 | 6940901 | S462 | ATN1 | T | C | exonic | nonsynonym | 1 | . |
| chr12 | 16277369 | 16277369 | S462 | SLC15A5 | C | A | exonic | nonsynonym | 0.210526315 | . |
| chr12 | 25108825 | 25108825 | S462 | CASC1 | T | TAAAAAAAA/ | splicing | . | 1 | . |
| chr12 | 29489672 | 29489672 | S462 | OVCH1 | C | A | exonic | nonsynonym | 0.234042553 | . |
| chr12 | 109067607 | 109067607 | S462 | USP30 | A | T | exonic | nonsynonym | 0.266666666 | . |
| chr12 | 121626874 | 121626879 | S462 | ORAI1 | CCGCCA | C | exonic | frameshift_d | 0.291666666 | . |
| chr13 | 28713430 | 28713431 | S462 | SLC46A3 | TA | T | exonic | stopgain | 0.410256410 | . |
| chr13 | 33111806 | 33111806 | S462 | STARD13 | A | C | exonic | nonsynonym | 1 | . |
| chr13 | 110456862 | 110456862 | S462 | COL4A2-AS2 | G | A | exonic | nonsynonym | 0.2 | rs561182370 |
| chr13 | 113404216 | 113404216 | S462 | LOC1019288 | C | T | exonic | nonsynonym | 0.3 | rs104709439 |
| chr14 | 30956239 | 30956239 | S462 | STRN3 | G | A | exonic | nonsynonym | 0.190476190 | rs766739305 |
| chr14 | 31305118 | 31305118 | S462 | HEATR5A | C | T | exonic | nonsynonym | 0.338235294 | rs61754158 |
| chr14 | 91313087 | 91313087 | S462 | CCDC88C | A | C | exonic | nonsynonym | 0.369565217 | . |
| chr14 | 92071009 | 92071009 | S462 | ATXN3 | C | CGCTGCTGCT | exonic | nonframeshi | 0.603773584 | . |
| chr15 | 23440409 | 23440451 | S462 | GOLGA6L2 | ACATCTTCTC | A | exonic | nonframeshi | 0.25 | . |
| chr15 | 28132778 | 28132778 | S462 | HERC2 | C | T | exonic | nonsynonym | 1 | rs576250950 |
| chr15 | 33821295 | 33821295 | S462 | RYR3 | C | A | exonic | nonsynonym | 0.163265306 | . |
| chr15 | 40987844 | 40987844 | S462 | INO80 | A | G | exonic | nonsynonym | 0.355555555 | . |
| chr15 | 44665972 | 44665972 | S462 | PATL2 | C | A | splicing | . | 0.153846153 | . |
| chr16 | 12003036 | 12003036 | S462 | SNX29 | G | A | exonic | nonsynonym | 0.448275862 | rs748795008 |
| chr16 | 29383437 | 29383437 | S462 | NPIPB11 | G | T | exonic | nonsynonym | 1 | rs796341971 |
| chr17 | 7676040 | 7676040 | S462 | TP53 | C | G | exonic | nonsynonym | 1 | rs11540654 |
| chr17 | 10400910 | 10400910 | S462 | MYH8 | C | G | exonic | nonsynonym | 0.476190476 | . |

| | | | | | | | | | | |
|---|---|---|---|---|---|---|---|---|---|---|
| chr17 | 31338739 | 31338739 | S462 | NF1 | C | A | exonic | stopgain | 1 | rs772295894 |
| chr17 | 34287635 | 34287635 | S462 | CCL11 | G | T | exonic | nonsynonym | 0.5 | . |
| chr17 | 37624364 | 37624364 | S462 | DDX52 | C | T | exonic | nonsynonym | 1 | rs7216445 |
| chr17 | 40780410 | 40780410 | S462 | KRT27 | T | A | exonic | nonsynonym | 0.4375 | . |
| chr17 | 43884083 | 43884083 | S462 | MPP2 | C | T | splicing | . | 1 | rs231518 |
| chr17 | 76088561 | 76088561 | S462 | EXOC7 | C | A | exonic | nonsynonym | 0.224137931 | . |
| chr17 | 79731335 | 79731335 | S462 | ENPP7 | T | C | exonic | nonsynonym | 0.211538461 | . |
| chr18 | 14852205 | 14852205 | S462 | ANKRD30B | A | T | exonic | stopgain | 0.243243243 | . |
| chr19 | 1065019 | 1065019 | S462 | ABCA7 | G | C | exonic | nonsynonym | 1 | . |
| chr19 | 10985393 | 10985393 | S462 | SMARCA4 | C | T | exonic | stopgain | 0.3 | . |
| chr19 | 11725227 | 11725227 | S462 | ZNF823 | T | C | exonic | nonsynonym | 0.272727272 | . |
| chr19 | 16773311 | 16773311 | S462 | NWD1 | G | A | exonic | nonsynonym | 1 | rs146121969 |
| chr19 | 39905924 | 39905924 | S462 | FCGBP | T | C | exonic | nonsynonym | 1 | . |
| chr19 | 42324318 | 42324318 | S462 | TMEM145 | T | G | exonic | nonsynonym | 0.4 | . |
| chr19 | 44908822 | 44908822 | S462 | APOE | C | G | exonic | nonsynonym | 1 | . |
| chr19 | 47545723 | 47545723 | S462 | ZNF541 | A | G | exonic | nonsynonym | 0.153846153 | . |
| chr19 | 52212745 | 52212745 | S462 | PPP2R1A | A | T | exonic | nonsynonym | 0.205128205 | . |
| chr19 | 56088901 | 56088901 | S462 | ZNF787 | C | T | exonic | nonsynonym | 0.192307692 | . |
| chr20 | 57365388 | 57365388 | S462 | RAE1 | T | G | exonic | nonsynonym | 0.177777777 | . |
| chr21 | 10569454 | 10569454 | S462 | TPTE | G | A | exonic | nonsynonym | 0.186666666 | rs1810856 |
| chr21 | 10569701 | 10569701 | S462 | TPTE | C | T | exonic | stopgain | 0.307692307 | rs1810540 |
| chr21 | 14592640 | 14592640 | S462 | LOC388813 | C | A | splicing | . | 0.238095238 | . |
| chr21 | 41889346 | 41889346 | S462 | C2CD2 | T | A | splicing | . | 0.166666666 | . |
| chr21 | 44289773 | 44289773 | S462 | AIRE | C | T | exonic | stopgain | 1 | rs121434254 |
| chr21 | 45530897 | 45530897 | S462 | SLC19A1 | C | A | exonic | nonsynonym | 1 | rs142899279 |
| chr22 | 20242802 | 20242802 | S462 | RTN4R | C | G | exonic | nonsynonym | 0.22 | . |
| chr22 | 25029315 | 25029315 | S462 | KIAA1671 | A | G | exonic | nonsynonym | 1 | rs17667531 |
| chr22 | 40301172 | 40301181 | S462 | TNRC6B | TGCAGCAGC/ | T | exonic | nonframeshi | 1 | . |
| chr22 | 42127941 | 42127941 | S462 | CYP2D6\x3b( | G | A | exonic | nonsynonym | 1 | rs16947 |
| chrX | 318607 | 318607 | S462 | GTPBP6 | C | G | exonic | nonsynonym | 1 | . |
| chrX | 12721595 | 12721595 | S462 | FRMPD4 | T | C | exonic | nonsynonym | 0.352941176 | . |
| chrX | 12721814 | 12721814 | S462 | FRMPD4 | G | A | exonic | nonsynonym | 0.454545454 | . |
| chrX | 23000451 | 23000452 | S462 | DDX53 | AC | A | exonic | frameshift_d | 0.32 | . |
| chrX | 121048045 | 121048045 | S462 | GLUD2 | C | T | exonic | nonsynonym | 0.117647058 | . |
| chrX | 124334929 | 124334929 | S462 | TEX13D | A | T | exonic | nonsynonym | 0.363636363 | . |
| chrX | 136536129 | 136536129 | S462 | VGLL1 | A | G | exonic | nonsynonym | 0.5 | . |

| | | | | | | | | | | |
|---|---|---|---|---|---|---|---|---|---|---|
| chrX | 152759523 | 152759523 | S462 | CSAG3 | A | T | exonic | nonsynonym | 0.272727272 | . |
| chrX | 153058185 | 153058185 | S462 | PNMA3 | C | T | exonic | nonsynonym | 1 | rs6526155 |

| Variant | AAChange.re | GeneDetail.r | AF | AF_popmax | prop_pathog | CLNALLELEID | CLNDN | CLNDISDB | CLNREVSTAT |
|---|---|---|---|---|---|---|---|---|---|
| chr1:7736404_G/T | | . | . | . | 1 | . | . | . | . |
| chr1:12725527_T/G | AADACL3:NM | . | . | . | NA | . | . | . | . |
| chr1:12760937_C/T | C1orf158:NM | . | . | . | NA | . | . | . | . |
| chr1:21833280_C/A | HSPG2:NM_( | . | . | . | 0.714285714 | . | . | . | . |
| chr1:26282344_ACCGGGACCGG | UBXN11:NM_ | . | 0.0004 | 0.0015 | NA | . | . | . | . |
| chr1:26337258_G/A | CRYBG2:NM_ | . | 0.0048 | 0.0057 | 0.714285714 | . | . | . | . |
| chr1:26345087_A/G | CRYBG2:NM_ | . | 0.0016 | 0.0039 | 1 | . | . | . | . |
| chr1:34904984_C/CTTG | DLGAP3:NM_ | . | . | . | NA | . | . | . | . |
| chr1:59638370_G/GT | | . | . | . | NA | . | . | . | . |
| chr1:146066541_T/G | NBPF10:NM_ | . | . | . | NA | . | . | . | . |
| chr1:146069686_C/T | NBPF10:NM_ | . | . | . | NA | . | . | . | . |
| chr1:148972321_C/T | | . | . | . | NA | . | . | . | . |
| chr1:149029862_A/G | | . | . | . | NA | . | . | . | . |
| chr1:213997621_A/C | PROX1:NM_( | . | . | . | 0.857142857 | . | . | . | . |
| chr1:235342705_G/A | GGPS1:NM_( | . | . | . | 0.857142857 | . | . | . | . |
| chr1:241671141_T/C | WDR64:NM_ | . | . | . | 0.8 | . | . | . | . |
| chr2:63882649_G/A | UGP2:NM_0( | . | . | . | 0.714285714 | . | . | . | . |
| chr2:112573788_C/T | | . | . | . | 0.714285714 | . | . | . | . |
| chr2:127220161_A/G | CYP27C1:NM | . | . | . | NA | . | . | . | . |
| chr2:205740533_C/A | NRP2:NM_0( | . | . | . | 1 | . | . | . | . |
| chr2:219235754_G/A | ANKZF1:NM_ | . | 0.0011 | 0.0017 | 1 | . | . | . | . |
| chr2:221482675_G/A | EPHA4:NM_( | . | . | . | 0.857142857 | . | . | . | . |
| chr3:13483535_C/T | HDAC11:NM_ | . | . | . | 1 | . | . | . | . |
| chr3:37007053_C/T | | . | . | . | 1 | . | . | . | . |
| chr3:46709583_TAAGAAG/TAA( | TMIE:NM_00 | . | . | . | NA | . | . | . | . |
| chr3:111112316_A/C | NECTIN3:NM | . | . | . | 0.857142857 | . | . | . | . |
| chr3:197511986_C/T | BDH1:NM_0( | . | 0.0015 | 3 | 1 | . | . | . | . |
| chr4:575941_T/C | TMEM271:NI | . | . | . | NA | . | . | . | . |
| chr4:9173181_AT/A | FAM90A26:N | . | 0 | . | NA | . | . | . | . |
| chr4:87615729_AATAGCAGTGA | DSPP:NM_01 | . | . | . | NA | . | . | . | . |
| chr4:133152506_T/C | PCDH10:NM_ | . | . | . | 0.714285714 | . | . | . | . |
| chr5:16670821_C/T | MYO10:NM_ | . | . | . | 0.833333333 | . | . | . | . |
| chr5:37195860_A/G | CPLANE1:NM | . | . | . | 0.714285714 | . | . | . | . |
| chr5:55954959_G/C | IL6ST:NM_00 | . | 0.0005 | 0.0012 | 0.857142857 | . | . | . | . |

| | | | | | | | | | |
|---|---|---|---|---|---|---|---|---|---|
| chr5:56914919_ACTGAAGCAGT/ | SETD9:NM_0 | . | . | . | NA | . | . | . | . |
| chr5:82304657_G/T | ATP6AP1L:NI | . | . | . | NA | . | . | . | . |
| chr5:82304658_A/C | ATP6AP1L:NI | . | . | . | NA | . | . | . | . |
| chr5:135009435_A/G | CATSPER3:NI | . | 0.0053 | 0.0062 | 0.714285714 | . | . | . | . |
| chr5:141673686_G/A | ARAP3:NM_( | . | . | . | 0.714285714 | . | . | . | . |
| chr5:149374580_C/T | IL17B:NM_0( | . | . | . | 0.857142857 | . | . | . | . |
| chr6:32530201_T/TCCAG | HLA-DRB5:NI | . | . | . | NA | . | . | . | . |
| chr6:82366051_G/T | TPBG:NM_0( | . | . | . | 0.857142857 | . | . | . | . |
| chr6:116925507_G/C | RFX6:NM_17 | . | 0.0014 | 0.0047 | 0.714285714 | 428498 | Monogenic_( | MedGen:C38 | criteria_prov |
| chr6:144531050_A/G | . | NM_007124: | . | . | 1 | . | . | . | . |
| chr6:170319057_A/T | FAM120B:NN | . | . | . | 0.714285714 | . | . | . | . |
| chr7:100893830_G/C | | . | 0.0006 | 0.0012 | 0.857142857 | . | . | . | . |
| chr7:138621107_C/A | SVOPL:NM_1 | . | 0.0011 | 0.0016 | 0.857142857 | . | . | . | . |
| chr7:140674264_G/C | . | NM_052853: | 0.0025 | 0.0083 | 1 | . | . | . | . |
| chr7:151195597_TC/T | IQCA1L:NM_( | . | . | . | NA | . | . | . | . |
| chr7:152144739_C/T | KMT2C:NM_: | . | . | . | 0.857142857 | . | . | . | . |
| chr8:8377352_CGCCGCT/C | | . | . | . | NA | . | . | . | . |
| chr8:25366878_C/A | DOCK5:NM_( | . | . | . | 0.714285714 | . | . | . | . |
| chr8:43315905_C/A | POTEA:NM_( | . | . | . | NA | . | . | . | . |
| chr8:53250839_A/G | OPRK1:NM_( | . | . | . | 0.714285714 | . | . | . | . |
| chr8:56068532_T/TAA | . | NM_0011462 | 0.0022 | 0.0049 | NA | . | . | . | . |
| chr8:90051875_T/G | DECR1:NM_( | . | . | . | 1 | . | . | . | . |
| chr8:112301930_C/A | CSMD3:NM_( | . | . | . | 0.857142857 | . | . | . | . |
| chr8:112352468_T/C | CSMD3:NM_( | . | 0.0009 | 0.0016 | 0.857142857 | . | . | . | . |
| chr8:117171168_GGTCAGTGA/( | . | . | . | . | NA | . | . | . | . |
| chr8:125049045_G/C | WASHC5:NM | . | . | . | 1 | . | . | . | . |
| chr8:144379425_C/A | ADCK5:NM_1 | . | . | . | NA | . | . | . | . |
| chr9:41964577_A/C | CNTNAP3B:N | . | . | . | NA | . | . | . | . |
| chr9:41991627_G/A | CNTNAP3B:N | . | . | . | NA | . | . | . | . |
| chr9:41991703_G/T | CNTNAP3B:N | . | . | . | NA | . | . | . | . |
| chr9:87668645_G/T | DAPK1:NM_( | . | 0.0025 | 4 | 0.714285714 | . | . | . | . |
| chr9:87921255_C/A | SPATA31C1:N | . | . | . | NA | . | . | . | . |
| chr9:91409678_AG/A | NFIL3:NM_0( | . | . | . | NA | . | . | . | . |
| chr9:94445142_G/C | MFSD14B:NN | . | 0.0009 | 0.0017 | 0.714285714 | . | . | . | . |
| chr9:96484604_G/T | HABP4:NM_( | . | . | . | 0.857142857 | . | . | . | . |
| chr9:97371638_CCACCTT/C | CCDC180:NN | . | . | . | NA | . | . | . | . |

| | | | | | | | | | |
|---|---|---|---|---|---|---|---|---|---|
| chr9:127813567_G/T | FPGS:NM_00 | . | . | . | 0.857142857 | . | . | . | . |
| chr9:137110891_GT/G | DPP7:NM_01 | . | . | . | NA | . | . | . | . |
| chr10:45000413_G/A | C10orf25:NM | . | . | . | NA | . | . | . | . |
| chr10:47502343_T/C | AGAP9:NM_( | . | . | . | NA | . | . | . | . |
| chr10:88594678_T/C | LIPJ:NM_001 | . | . | . | 0.857142857 | . | . | . | . |
| chr10:93076668_G/A | CYP26A1:NM | . | . | . | 0.857142857 | . | . | . | . |
| chr10:104034648_G/A | COL17A1:NM | . | 0.0004 | 0.0012 | 0.857142857 | . | . | . | . |
| chr10:120890878_G/A | WDR11:NM_ | . | . | . | 0.571428571 | . | . | . | . |
| chr11:4587098_C/A | OR52I2:NM_( | . | . | . | 0.714285714 | . | . | . | . |
| chr11:5200445_T/C | OR51V1:NM_ | . | . | . | 0.714285714 | . | . | . | . |
| chr11:61776320_A/T | . | NM_0011273 | . | . | 1 | . | . | . | . |
| chr11:78658315_C/T | TENM4:NM_( | . | . | . | 0.714285714 | . | . | . | . |
| chr11:92799473_T/C | FAT3:NM_00 | . | . | . | 0.833333333 | . | . | . | . |
| chr12:6940901_T/C | ATN1:NM_0C | . | . | . | 0.714285714 | . | . | . | . |
| chr12:16277369_C/A | SLC15A5:NM | . | . | . | 1 | . | . | . | . |
| chr12:25108825_T/TAAAAAAAA | . | | . | . | NA | . | . | . | . |
| chr12:29489672_C/A | OVCH1:NM_( | . | . | . | 0.666666666 | . | . | . | . |
| chr12:109067607_A/T | USP30:NM_0 | . | . | . | 0.714285714 | . | . | . | . |
| chr12:121626874_CCGCCA/C | ORAI1:NM_0 | . | . | . | NA | . | . | . | . |
| chr13:28713430_TA/T | SLC46A3:NM | . | . | . | NA | . | . | . | . |
| chr13:33111806_A/C | STARD13:NM | . | . | . | 0.857142857 | . | . | . | . |
| chr13:110456862_G/A | COL4A2-AS2: | . | 0.0006 | 0.0012 | NA | . | . | . | . |
| chr13:113404216_C/T | LOC1019288 | . | . | . | NA | . | . | . | . |
| chr14:30956239_G/A | STRN3:NM_0 | . | . | . | 1 | . | . | . | . |
| chr14:31305118_C/T | HEATR5A:NM | . | 0.0040 | 0.0065 | 0.857142857 | . | . | . | . |
| chr14:91313087_A/C | CCDC88C:NM | . | . | . | 0.714285714 | . | . | . | . |
| chr14:92071009_C/CGCTGCTGCTGCTGCTGCT | | . | . | . | NA | . | . | . | . |
| chr15:23440409_ACATCTTCTCC | GOLGA6L2:N | . | . | . | NA | . | . | . | . |
| chr15:28132778_C/T | HERC2:NM_C | . | . | . | 0.857142857 | . | . | . | . |
| chr15:33821295_C/A | RYR3:NM_00 | . | . | . | 0.714285714 | . | . | . | . |
| chr15:40987844_A/G | INO80:NM_0 | . | . | . | 0.857142857 | . | . | . | . |
| chr15:44665972_C/A | . | NM_0013302 | . | . | 1 | . | . | . | . |
| chr16:12003036_G/A | SNX29:NM_0 | . | . | . | 0.8 | . | . | . | . |
| chr16:29383437_G/T | NPIPB11:NM | . | . | . | 0.75 | . | . | . | . |
| chr17:7676040_C/G | | . | . | . | 0.714285714 | 236497 | Hereditary_c | MedGen:C00 | criteria_prov |
| chr17:10400910_C/G | MYH8:NM_0 | . | . | . | 0.857142857 | . | . | . | . |

| | | | | | | | | | |
|---|---|---|---|---|---|---|---|---|---|
| chr17:31338739_C/A | NF1:NM_000 | . | . | . | 1 | 184734 | Hereditary_c | MedGen:C00 | criteria_prov |
| chr17:34287635_G/T | CCL11:NM_0 | . | . | . | 0.833333333 | . | . | . | . |
| chr17:37624364_C/T | DDX52:NM_( | . | . | . | NA | . | . | . | . |
| chr17:40780410_T/A | KRT27:NM_1 | . | . | . | 0.857142857 | . | . | . | . |
| chr17:43884083_C/T | . | NM_0012783 | . | . | NA | . | . | . | . |
| chr17:76088561_C/A | EXOC7:NM_( | . | . | . | 1 | . | . | . | . |
| chr17:79731335_T/C | ENPP7:NM_1 | . | . | . | 0.857142857 | . | . | . | . |
| chr18:14852205_A/T | ANKRD30B:N | . | . | . | 1 | . | . | . | . |
| chr19:1065019_G/C | ABCA7:NM_( | . | . | . | 0.833333333 | . | . | . | . |
| chr19:10985393_C/T | | . | . | . | 1 | . | . | . | . |
| chr19:11725227_T/C | ZNF823:NM_ | . | . | . | 0.833333333 | . | . | . | . |
| chr19:16773311_G/A | NWD1:NM_( | . | 0.0050 | 0.0064 | 0.714285714 | . | . | . | . |
| chr19:39905924_T/C | FCGBP:NM_( | . | . | . | NA | . | . | . | . |
| chr19:42324318_T/G | TMEM145:NI | . | . | . | NA | . | . | . | . |
| chr19:44908822_C/G | APOE:NM_0( | . | . | . | 0.714285714 | . | . | . | . |
| chr19:47545723_A/G | ZNF541:NM_ | . | . | . | 0.666666666 | . | . | . | . |
| chr19:52212745_A/T | PPP2R1A:NM | . | . | . | 0.857142857 | . | . | . | . |
| chr19:56088901_C/T | ZNF787:NM_ | . | . | . | 0.75 | . | . | . | . |
| chr20:57365388_T/G | RAE1:NM_00 | . | . | . | 1 | . | . | . | . |
| chr21:10569454_G/A | TPTE:NM_00 | . | . | . | NA | . | . | . | . |
| chr21:10569701_C/T | TPTE:NM_00 | . | . | . | NA | . | . | . | . |
| chr21:14592640_C/A | . | NM_0012565 | . | . | NA | . | . | . | . |
| chr21:41889346_T/A | . | NM_015500: | . | . | 1 | . | . | . | . |
| chr21:44289773_C/T | AIRE:NM_00( | . | 0.0011 | 1 | 0.5 | 18346 | Polyglandula | MedGen:C00 | criteria_prov |
| chr21:45530897_C/A | | . | 0.0007 | 0.0012 | 0.714285714 | . | . | . | . |
| chr22:20242802_C/G | RTN4R:NM_( | . | . | . | 0.714285714 | . | . | . | . |
| chr22:25029315_A/G | KIAA1671:NN | . | . | . | 0.666666666 | . | . | . | . |
| chr22:40301172_TGCAGCAGCA, | TNRC6B:NM_ | . | . | . | NA | . | . | . | . |
| chr22:42127941_G/A | CYP2D6:NM_ | . | . | . | NA | . | . | . | . |
| chrX:318607_C/G | GTPBP6:NM_ | . | . | . | NA | . | . | . | . |
| chrX:12721595_T/C | FRMPD4:NM_ | . | . | . | NA | . | . | . | . |
| chrX:12721814_G/A | FRMPD4:NM_ | . | . | . | NA | . | . | . | . |
| chrX:23000451_AC/A | DDX53:NM_1 | . | . | . | NA | . | . | . | . |
| chrX:121048045_C/T | GLUD2:NM_( | . | . | . | 0.857142857 | . | . | . | . |
| chrX:124334929_A/T | TEX13D:NM_ | . | . | . | NA | . | . | . | . |
| chrX:136536129_A/G | VGLL1:NM_0 | . | . | . | 0.857142857 | . | . | . | . |

| | | | | | | | | | |
|---|---|---|---|---|---|---|---|---|---|
| chrX:152759523_A/T | CSAG3:NM_( | . | . | . | NA | . | . | . | . |
| chrX:153058185_C/T | PNMA3:NM_ | . | . | . | NA | . | . | . | . |

| CLNSIG | SIFT_pred | Polyphen2_H | Polyphen2_H | LRT_pred | FATHMM_pr | MutationTas | MutationAss | InterVar_automated |
|---|---|---|---|---|---|---|---|---|
| . | . | . | . | D | . | A | . | Pathogenic |
| . | . | . | . | . | . | . | . | Uncertain_significance |
| . | . | . | . | . | . | . | . | Uncertain_significance |
| . | D | D | D | N | T | D | M | Uncertain_significance |
| . | . | . | . | . | . | . | . | . |
| . | D | D | D | N | T | D | M | Uncertain_significance |
| . | . | . | . | . | . | P | . | . |
| . | . | . | . | . | . | . | . | . |
| . | . | . | . | . | . | . | . | . |
| . | . | . | . | . | . | . | . | . |
| . | . | . | . | . | . | . | . | Uncertain_significance |
| . | . | . | . | . | . | . | . | Uncertain_significance |
| . | . | . | . | . | . | . | . | Uncertain_significance |
| . | D | D | D | D | T | D | M | Uncertain_significance |
| . | D | D | D | D | T | D | M | Uncertain_significance |
| . | D | . | . | D | T | D | M | Uncertain_significance |
| . | T | P | P | D | T | D | M | Uncertain_significance |
| . | T | D | D | D | T | D | M | Uncertain_significance |
| . | . | . | . | . | . | . | . | . |
| . | D | D | D | D | D | D | H | Uncertain_significance |
| . | D | D | P | D | D | D | M | Uncertain_significance |
| . | D | D | D | D | T | D | H | . |
| . | . | D | D | D | D | D | M | Uncertain_significance |
| . | D | D | D | D | D | D | H | Uncertain_significance |
| . | . | . | . | . | . | . | . | . |
| . | D | D | D | D | T | D | M | Uncertain_significance |
| . | D | D | D | D | D | D | M | Uncertain_significance |
| . | . | . | . | . | . | . | . | . |
| . | . | . | . | . | . | . | . | . |
| . | . | . | . | . | . | . | . | . |
| . | D | D | D | N | T | D | M | Uncertain_significance |
| . | D | D | D | . | T | D | M | . |
| . | D | D | P | N | T | D | M | Uncertain_significance |
| . | D | D | D | D | T | D | M | Uncertain_significance |

| | | | | | | | | |
|---|---|---|---|---|---|---|---|---|
| . | . | . | . | . | . | . | . | . |
| . | . | . | . | . | . | . | . | Uncertain_significance |
| . | . | . | . | . | . | . | . | Uncertain_significance |
| . | D | P | B | D | D | N | M | Uncertain_significance |
| . | D | D | D | N | T | D | M | Uncertain_significance |
| . | D | D | D | D | T | D | M | Uncertain_significance |
| . | . | . | . | . | . | . | . | . |
| . | D | D | P | D | T | D | M | Uncertain_significance |
| Conflicting_ir | D | D | D | D | T | D | L | Uncertain_significance |
| . | . | . | . | . | . | D | . | . |
| . | D | D | P | N | T | D | M | Uncertain_significance |
| . | D | D | D | D | T | D | M | Uncertain_significance |
| . | D | D | P | D | T | D | H | Uncertain_significance |
| . | . | . | . | . | . | D | . | . |
| . | . | . | . | . | . | . | . | . |
| . | D | D | D | U | D | D | M | Uncertain_significance |
| . | . | . | . | . | . | . | . | . |
| . | D | P | B | D | T | D | M | Uncertain_significance |
| . | . | . | . | . | . | . | . | Uncertain_significance |
| . | T | D | D | D | T | D | M | Uncertain_significance |
| . | . | . | . | . | . | . | . | . |
| . | D | P | P | D | D | D | M | Uncertain_significance |
| . | D | D | D | D | T | D | H | Uncertain_significance |
| . | D | D | D | D | T | D | M | Uncertain_significance |
| . | . | . | . | . | . | . | . | . |
| . | . | . | . | D | . | A | . | Likely_pathogenic |
| . | . | . | . | . | . | . | . | Uncertain_significance |
| . | . | . | . | . | . | . | . | . |
| . | . | . | . | . | . | . | . | . |
| . | . | . | . | . | . | . | . | Uncertain_significance |
| . | T | P | P | D | T | D | M | Uncertain_significance |
| . | . | . | . | . | . | . | . | Uncertain_significance |
| . | . | . | . | . | . | . | . | . |
| . | T | P | P | D | T | D | M | Uncertain_significance |
| . | D | D | D | D | T | D | M | Uncertain_significance |
| . | . | . | . | . | . | . | . | . |

| | | | | | | | | |
|---|---|---|---|---|---|---|---|---|
| . | D | D | P | D | T | D | H | Uncertain_significance |
| . | . | . | . | . | . | . | . | . |
| . | . | . | . | . | . | . | . | Uncertain_significance |
| . | . | . | . | . | . | . | . | Uncertain_significance |
| . | D | D | D | D | T | D | H | Uncertain_significance |
| . | D | D | D | D | T | D | M | Uncertain_significance |
| . | D | D | P | N | D | D | M | Uncertain_significance |
| . | T | P | B | D | D | D | L | Likely_pathogenic |
| . | D | D | D | N | D | N | H | Uncertain_significance |
| . | D | D | D | D | T | N | M | Uncertain_significance |
| . | . | . | . | . | . | D | . | . |
| . | T | D | D | D | D | D | L | Likely_pathogenic |
| . | D | D | D | . | T | D | M | Uncertain_significance |
| . | D | D | D | U | T | D | M | Uncertain_significance |
| . | D | . | . | . | D | D | H | . |
| . | . | . | . | . | . | . | . | . |
| . | D | D | D | . | D | N | L | Uncertain_significance |
| . | D | P | P | D | T | D | L | Uncertain_significance |
| . | . | . | . | . | . | . | . | . |
| . | . | . | . | . | . | . | . | . |
| . | D | D | D | D | T | D | M | Uncertain_significance |
| . | . | . | . | . | . | . | . | Uncertain_significance |
| . | . | . | . | . | . | . | . | Uncertain_significance |
| . | D | D | D | D | D | D | M | . |
| . | D | D | D | D | T | D | M | Uncertain_significance |
| . | D | D | D | U | T | D | M | Uncertain_significance |
| . | . | . | . | . | . | . | . | . |
| . | . | . | . | . | . | . | . | . |
| . | D | D | P | D | D | D | N | Uncertain_significance |
| . | D | P | B | D | D | D | N | Uncertain_significance |
| . | D | D | D | D | T | D | H | Likely_pathogenic |
| . | . | . | . | . | . | D | . | . |
| . | D | . | . | D | T | D | M | Uncertain_significance |
| . | D | . | . | . | T | P | M | . |
| Pathogenic/L | D | D | P | N | D | N | M | Uncertain_significance |
| . | D | P | P | U | D | D | M | Uncertain_significance |

| | | | | | | | | |
|---|---|---|---|---|---|---|---|---|
| Pathogenic | . | . | . | D | . | A | . | Pathogenic |
| . | D | D | D | D | T | D | . | Uncertain_significance |
| . | . | . | . | . | . | . | . | . |
| . | D | D | D | N | D | D | M | . |
| . | . | . | . | . | . | . | . | . |
| . | D | P | P | D | . | D | M | . |
| . | D | D | D | D | T | D | M | Uncertain_significance |
| . | . | . | . | . | . | D | . | Uncertain_significance |
| . | D | D | D | . | D | N | M | Uncertain_significance |
| . | . | . | . | D | . | A | . | Pathogenic |
| . | D | D | D | . | T | D | H | Uncertain_significance |
| . | D | D | P | U | T | D | M | Uncertain_significance |
| . | . | . | . | . | . | . | . | Uncertain_significance |
| . | . | . | . | . | . | . | . | . |
| . | D | D | D | D | T | N | M | Uncertain_significance |
| . | D | D | D | . | T | D | L | Uncertain_significance |
| . | D | D | D | D | T | D | M | Uncertain_significance |
| . | . | D | P | . | . | D | L | Uncertain_significance |
| . | D | D | D | D | D | D | M | Uncertain_significance |
| . | . | . | . | . | . | . | . | Uncertain_significance |
| . | . | . | . | . | . | . | . | Uncertain_significance |
| . | . | . | . | . | . | . | . | . |
| . | . | . | . | . | . | D | . | . |
| Pathogenic | . | . | . | N | . | A | . | Pathogenic |
| . | T | D | P | D | D | D | L | Uncertain_significance |
| . | D | P | P | N | T | D | M | Uncertain_significance |
| . | D | D | P | N | . | P | L | Uncertain_significance |
| . | . | . | . | . | . | . | . | . |
| . | . | . | . | . | . | . | . | Uncertain_significance |
| . | . | . | . | . | . | . | . | Uncertain_significance |
| . | . | . | . | . | . | . | . | . |
| . | . | . | . | . | . | . | . | . |
| . | . | . | . | . | . | . | . | . |
| . | D | D | D | N | D | D | M | Uncertain_significance |
| . | . | . | . | . | . | . | . | . |
| . | D | D | D | D | T | D | M | Uncertain_significance |

| | | | | | | | | |
|---|---|---|---|---|---|---|---|---|
| . | . | . | . | . | . | . | . | Uncertain_significance |
| . | . | . | . | . | . | . | . | Uncertain_significance |

### ST88-14 cell line variants with a potential impact on protein function

| chr | start | end | Sample | Gene.refGen | REF | ALT | Func.refGene | ExonicFunc.r | VAF | avsnp150 |
|---|---|---|---|---|---|---|---|---|---|---|
| chr1 | 32363838 | 32363838 | ST88-14 | TSSK3 | T | C | exonic | nonsynonym | 1 | . |
| chr1 | 56908064 | 56908064 | ST88-14 | C8A | G | A | exonic | nonsynonym | 1 | rs143908758 |
| chr1 | 75300612 | 75300612 | ST88-14 | SLC44A5 | C | A | exonic | nonsynonym | 0.451612903 | . |
| chr1 | 146121510 | 146121510 | ST88-14 | NBPF10 | A | C | exonic | nonsynonym | 1 | . |
| chr1 | 146984940 | 146984940 | ST88-14 | NBPF12 | T | TACGTATTGC | exonic | nonframeshi | 0.95 | . |
| chr1 | 149003009 | 149003009 | ST88-14 | PDE4DIP | G | A | exonic | nonsynonym | 1 | . |
| chr2 | 95181295 | 95181295 | ST88-14 | ZNF2 | T | TGCG | exonic | nonframeshi | 0.274509803 | . |
| chr2 | 177264531 | 177264531 | ST88-14 | NFE2L2 | C | G | splicing | . | 0.6 | . |
| chr3 | 12004864 | 12004864 | ST88-14 | SYN2 | C | G | exonic | nonsynonym | 0.857142857 | rs2923857 |
| chr3 | 71755044 | 71755044 | ST88-14 | GPR27 | G | A | exonic | nonsynonym | 0.25 | rs867272686 |
| chr3 | 74285305 | 74285305 | ST88-14 | CNTN3 | G | A | exonic | nonsynonym | 0.258064516 | rs101610991 |
| chr3 | 75730493 | 75730493 | ST88-14 | ZNF717 | C | A | exonic | nonsynonym | 0.344827586 | rs112949302 |
| chr3 | 75730499 | 75730499 | ST88-14 | ZNF717 | G | C | exonic | stopgain | 0.333333333 | rs200767888 |
| chr3 | 98007903 | 98007903 | ST88-14 | GABRR3 | A | T | exonic | stopgain | 0.5 | rs832032 |
| chr3 | 127963356 | 127963356 | ST88-14 | KBTBD12 | C | T | exonic | nonsynonym | 0.321428571 | . |
| chr3 | 194137736 | 194137736 | ST88-14 | HES1 | G | A | exonic | nonsynonym | 0.166666666 | . |
| chr4 | 42401099 | 42401099 | ST88-14 | SHISA3 | C | G | exonic | nonsynonym | 0.395348837 | . |
| chr4 | 56929904 | 56929904 | ST88-14 | REST | G | A | exonic | nonsynonym | 0.466666666 | . |
| chr4 | 105940114 | 105940114 | ST88-14 | NPNT | G | T | exonic | nonsynonym | 0.583333333 | . |
| chr4 | 122274435 | 122274439 | ST88-14 | KIAA1109 | CAGTA | C | exonic | frameshift_d | 0.228571428 | . |
| chr4 | 182679690 | 182679690 | ST88-14 | TENM3 | G | A | exonic | nonsynonym | 0.423076923 | . |
| chr5 | 77433193 | 77433193 | ST88-14 | WDR41 | C | T | exonic | nonsynonym | 0.4 | rs41272254 |
| chr5 | 141137592 | 141137592 | ST88-14 | PCDHB5 | C | T | exonic | nonsynonym | 0.548387096 | rs400562 |
| chr5 | 149317938 | 149317938 | ST88-14 | AFAP1L1 | G | A | exonic | nonsynonym | 0.6 | . |
| chr6 | 16327633 | 16327633 | ST88-14 | ATXN1 | G | GTGCTGCTGC | exonic | nonframeshi | 1 | . |
| chr6 | 16327633 | 16327633 | ST88-14 | ATXN1 | G | GTGCTGCTGC | exonic | nonframeshi | 1 | . |
| chr6 | 28301034 | 28301034 | ST88-14 | PGBD1 | G | A | exonic | nonsynonym | 1 | rs776958236 |
| chr6 | 70956144 | 70956144 | ST88-14 | B3GAT2 | C | G | exonic | nonsynonym | 0.3 | . |
| chr6 | 109442223 | 109442223 | ST88-14 | SMPD2 | A | G | exonic | nonsynonym | 0.142857142 | . |
| chr6 | 151808004 | 151808004 | ST88-14 | ESR1 | T | C | exonic | nonsynonym | 0.166666666 | rs752794477 |
| chr6 | 167976591 | 167976591 | ST88-14 | HGC6.3 | G | A | exonic | nonsynonym | 0.230769230 | rs61740140 |
| chr7 | 6608019 | 6608019 | ST88-14 | C7orf26 | C | G | exonic | nonsynonym | 0.488372093 | . |
| chr7 | 32543261 | 32543261 | ST88-14 | AVL9 | G | A | exonic | nonsynonym | 0.218181818 | . |
| chr7 | 99707873 | 99707873 | ST88-14 | CYP3A7\x3bC | A | T | exonic | nonsynonym | 0.421052631 | . |

| | | | | | | | | | | |
|---|---|---|---|---|---|---|---|---|---|---|
| chr7 | 151195597 | 151195598 | ST88-14 | IQCA1L | TC | T | exonic | frameshift_d | 0.513157894 | . |
| chr8 | 450074 | 450074 | ST88-14 | FBXO25 | G | A | exonic | nonsynonym | 1 | rs562354345 |
| chr8 | 8318865 | 8318871 | ST88-14 | PRAG1 | CGGGGCG | C | exonic | nonframeshi | 1 | . |
| chr8 | 141494938 | 141494938 | ST88-14 | MROH5 | C | T | splicing | . | 0.5 | rs6578193 |
| chr9 | 66989995 | 66989995 | ST88-14 | SPATA31A3 | G | A | exonic | nonsynonym | 0.533333333 | rs273319 |
| chr9 | 136286366 | 136286366 | ST88-14 | CCDC187 | A | C | exonic | nonsynonym | 1 | . |
| chr9 | 137500775 | 137500775 | ST88-14 | PNPLA7 | A | G | exonic | nonsynonym | 0.166666666 | . |
| chr10 | 47368085 | 47368085 | ST88-14 | ZNF488 | G | A | exonic | nonsynonym | 1 | rs3814160 |
| chr10 | 47368615 | 47368615 | ST88-14 | ZNF488 | G | A | exonic | nonsynonym | 1 | rs35618062 |
| chr10 | 47502343 | 47502343 | ST88-14 | AGAP9 | T | C | exonic | nonsynonym | 1 | rs1047447 |
| chr10 | 47502604 | 47502604 | ST88-14 | AGAP9 | A | G | exonic | nonsynonym | 1 | rs4013543 |
| chr10 | 47502991 | 47502991 | ST88-14 | AGAP9 | C | T | exonic | nonsynonym | 1 | rs4013536 |
| chr10 | 103350986 | 103350986 | ST88-14 | PCGF6 | A | AGGCGGG | exonic | nonframeshi | 0.285714285 | . |
| chr10 | 124496962 | 124496962 | ST88-14 | LHPP | C | T | exonic | nonsynonym | 0.153846153 | rs199754897 |
| chr11 | 8717833 | 8717833 | ST88-14 | ST5 | G | A | exonic | stopgain | 1 | . |
| chr11 | 24738261 | 24738261 | ST88-14 | LUZP2 | C | G | exonic | nonsynonym | 0.4 | . |
| chr11 | 34161000 | 34161000 | ST88-14 | ABTB2 | A | T | exonic | nonsynonym | 0.142857142 | . |
| chr11 | 54603368 | 54603368 | ST88-14 | OR4C46 | G | A | exonic | nonsynonym | 0.725490196 | rs77689730 |
| chr11 | 54603640 | 54603640 | ST88-14 | OR4C46 | T | C | exonic | nonsynonym | 0.651162790 | rs74396937 |
| chr11 | 54603820 | 54603820 | ST88-14 | OR4C46 | G | A | exonic | nonsynonym | 1 | rs11246606 |
| chr11 | 54706901 | 54706901 | ST88-14 | OR4A5 | A | G | exonic | nonsynonym | 0.729166666 | rs10902343 |
| chr11 | 64607578 | 64607578 | ST88-14 | NRXN2 | G | C | exonic | nonsynonym | 0.655172413 | . |
| chr11 | 103064338 | 103064338 | ST88-14 | DCUN1D5 | T | C | exonic | nonsynonym | 0.341463414 | . |
| chr11 | 118901219 | 118901219 | ST88-14 | BCL9L | G | A | exonic | nonsynonym | 0.384615384 | rs766026718 |
| chr11 | 124250452 | 124250452 | ST88-14 | OR8G1 | C | G | exonic | stopgain | 1 | rs4268525 |
| chr11 | 125614101 | 125614101 | ST88-14 | STT3A | G | T | exonic | nonsynonym | 0.5 | . |
| chr11 | 125948031 | 125948031 | ST88-14 | VSIG10L2 | T | TGCTGGGTG/ | exonic | nonframeshi | 0.387096774 | . |
| chr12 | 6936728 | 6936731 | ST88-14 | ATN1 | ACAG | A | exonic | nonframeshi | 1 | . |
| chr12 | 49795696 | 49795696 | ST88-14 | NCKAP5L | T | G | exonic | nonsynonym | 1 | . |
| chr13 | 77081955 | 77081955 | ST88-14 | MYCBP2 | A | T | exonic | nonsynonym | 0.518518518 | . |
| chr14 | 19434064 | 19434064 | ST88-14 | POTEG | T | C | exonic | nonsynonym | 1 | rs57000026 |
| chr14 | 24057240 | 24057240 | ST88-14 | CARMIL3 | A | G | exonic | nonsynonym | 0.818181818 | rs45508393 |
| chr14 | 24177197 | 24177200 | ST88-14 | REC8 | GGAA | G | exonic | nonframeshi | 0.768115942 | . |
| chr14 | 68793049 | 68793049 | ST88-14 | ZFP36L1 | C | T | exonic | nonsynonym | 0.590909090 | . |
| chr15 | 21422162 | 21422162 | ST88-14 | POTEB3 | T | G | exonic | nonsynonym | 0.339622641 | . |
| chr15 | 22613584 | 22613584 | ST88-14 | GOLGA8J | A | G | exonic | nonsynonym | 1 | . |

| | | | | | | | | | | |
|---|---|---|---|---|---|---|---|---|---|---|
| chr15 | 22903836 | 22903836 | ST88-14 | CYFIP1 | C | T | exonic | nonsynonym | 1 | rs7170637 |
| chr16 | 1433059 | 1433059 | ST88-14 | PERCC1 | G | A | exonic | nonsynonym | 0.5 | . |
| chr16 | 71536914 | 71536914 | ST88-14 | CHST4 | G | C | exonic | nonsynonym | 0.289473684 | . |
| chr16 | 88951212 | 88951212 | ST88-14 | LOC1001296 | C | G | exonic | nonsynonym | 1 | rs71395352 |
| chr17 | 1643887 | 1643887 | ST88-14 | SCARF1 | C | T | exonic | nonsynonym | 0.181818181 | . |
| chr17 | 31221856 | 31221856 | ST88-14 | NF1 | C | CT | exonic | frameshift_ir | 1 | . |
| chr17 | 41097869 | 41097869 | ST88-14 | KRTAP4-8 | G | GGGGC | exonic | frameshift_ir | 0.3 | . |
| chr17 | 41097873 | 41097873 | ST88-14 | KRTAP4-8 | A | AG | exonic | frameshift_ir | 0.363636363 | . |
| chr17 | 41097875 | 41097875 | ST88-14 | KRTAP4-8 | A | AGCTGGAGA | exonic | frameshift_ir | 0.454545454 | . |
| chr17 | 41487398 | 41487398 | ST88-14 | KRT36 | G | A | exonic | nonsynonym | 1 | rs201168056 |
| chr17 | 50168409 | 50168409 | ST88-14 | SGCA | C | A | exonic | nonsynonym | 0.377049180 | rs35130237 |
| chr18 | 49912148 | 49912148 | ST88-14 | MYO5B | G | A | exonic | nonsynonym | 1 | rs771897253 |
| chr18 | 74447220 | 74447220 | ST88-14 | DIPK1C | C | A | exonic | stopgain | 1 | . |
| chr19 | 2076895 | 2076895 | ST88-14 | MOB3A | C | A | exonic | nonsynonym | 1 | . |
| chr19 | 8605046 | 8605046 | ST88-14 | ADAMTS10 | C | G | exonic | nonsynonym | 1 | rs7255721 |
| chr19 | 8698550 | 8698550 | ST88-14 | ACTL9 | G | C | exonic | nonsynonym | 1 | rs10410943 |
| chr19 | 41967684 | 41967684 | ST88-14 | ATP1A3 | C | A | exonic | nonsynonym | 0.271186440 | . |
| chr19 | 51768878 | 51768878 | ST88-14 | FPR2 | T | C | exonic | nonsynonym | 0.488372093 | rs74602258 |
| chr19 | 54735264 | 54735264 | ST88-14 | KIR3DL3 | A | T | exonic | nonsynonym | 0.422222222 | rs602444 |
| chr19 | 54832784 | 54832784 | ST88-14 | KIR2DS4 | T | G | exonic | nonsynonym | 0.391304347 | rs1130476 |
| chr19 | 54835110 | 54835110 | ST88-14 | KIR2DS4 | G | T | exonic | nonsynonym | 0.416666666 | rs1130478 |
| chr21 | 33794369 | 33794369 | ST88-14 | ITSN1 | A | G | exonic | nonsynonym | 0.305555555 | . |
| chr21 | 45504510 | 45504529 | ST88-14 | COL18A1 | CCGGCCCCC | CA | exonic | nonframeshi | 1 | . |
| chr22 | 21923433 | 21923433 | ST88-14 | PPM1F | C | T | exonic | nonsynonym | 1 | . |
| chr22 | 50283705 | 50283705 | ST88-14 | PLXNB2 | T | G | exonic | nonsynonym | 1 | . |
| chrX | 315246 | 315246 | ST88-14 | GTPBP6 | T | G | exonic | nonsynonym | 0.464285714 | . |
| chrX | 318607 | 318607 | ST88-14 | GTPBP6 | C | G | exonic | nonsynonym | 0.5 | . |
| chrX | 1193297 | 1193297 | ST88-14 | CRLF2 | T | C | exonic | nonsynonym | 0.370370370 | . |
| chrX | 15587851 | 15587851 | ST88-14 | ACE2 | G | A | exonic | nonsynonym | 0.157894736 | . |
| chrX | 27821947 | 27821947 | ST88-14 | MAGEB10 | A | AG | exonic | frameshift_ir | 0.333333333 | . |
| chrX | 84468584 | 84468584 | ST88-14 | HDX | T | C | exonic | nonsynonym | 1 | . |
| chrX | 152135086 | 152135086 | ST88-14 | MAGEA10 | C | A | exonic | nonsynonym | 0.285714285 | . |
| chrX | 152728157 | 152728157 | ST88-14 | CSAG1 | G | C | exonic | stopgain | 1 | rs1894360 |

| Variant | AAChange.re | GeneDetail.r | AF | AF_popmax | prop_pathog | CLNALLELEID | CLNDN | CLNDISDB | CLNREVSTAT | CLNSIG |
|---|---|---|---|---|---|---|---|---|---|---|
| chr1:323638 | TSSK3:NM_0 | . | . | . | 0.714285714 | . | . | . | . | . |
| chr1:569080 | C8A:NM_000 | . | 0.0031 | 0.0038 | 1 | . | . | . | . | . |
| chr1:753006 | SLC44A5:NM | . | . | . | 0.857142857 | . | . | . | . | . |
| chr1:146121 | NBPF10:NM_ | . | . | . | NA | . | . | . | . | . |
| chr1:146984 | NBPF12:NM_ | . | . | . | NA | . | . | . | . | . |
| chr1:149003009_G/A | | . | . | . | NA | . | . | . | . | . |
| chr2:95181295_T/TGCG | | . | . | . | NA | . | . | . | . | . |
| chr2:177264 | . | NM_001313 | . | . | 1 | . | . | . | . | . |
| chr3:120048 | SYN2:NM_00 | . | . | . | NA | . | . | . | . | . |
| chr3:717550 | GPR27:NM_ | . | . | . | 0.714285714 | . | . | . | . | . |
| chr3:742853 | CNTN3:NM_ | . | . | . | 0.857142857 | . | . | . | . | . |
| chr3:757304 | ZNF717:NM_ | . | 0.0016 | 0.0028 | NA | . | . | . | . | . |
| chr3:757304 | ZNF717:NM_ | . | 0.0038 | 0.0075 | NA | . | . | . | . | . |
| chr3:980079 | GABRR3:NM_ | . | . | . | NA | . | . | . | . | . |
| chr3:127963 | KBTBD12:NM | . | . | . | 0.857142857 | . | . | . | . | . |
| chr3:194137 | HES1:NM_00 | . | . | . | 0.857142857 | . | . | . | . | . |
| chr4:424010 | SHISA3:NM_ | . | . | . | 0.714285714 | . | . | . | . | . |
| chr4:569299 | REST:NM_00 | . | . | . | 0.857142857 | . | . | . | . | . |
| chr4:105940 | NPNT:NM_0 | . | . | . | 0.714285714 | . | . | . | . | . |
| chr4:122274 | KIAA1109:NM | . | . | . | NA | . | . | . | . | . |
| chr4:182679 | TENM3:NM_ | . | . | . | 0.833333333 | . | . | . | . | . |
| chr5:774331 | WDR41:NM_ | . | 0.0040 | 5 | 0.714285714 | . | . | . | . | . |
| chr5:141137 | PCDHB5:NM_ | . | . | . | NA | . | . | . | . | . |
| chr5:149317 | AFAP1L1:NM | . | . | . | 0.857142857 | . | . | . | . | . |
| chr6:163276 | ATXN1:NM_ | . | 0.0065 | 0.0091 | NA | . | . | . | . | . |
| chr6:163276 | ATXN1:NM_ | . | . | . | NA | . | . | . | . | . |
| chr6:283010 | PGBD1:NM_ | . | . | . | 0.857142857 | . | . | . | . | . |
| chr6:709561 | B3GAT2:NM_ | . | . | . | 0.857142857 | . | . | . | . | . |
| chr6:109442 | SMPD2:NM_ | . | . | . | 0.857142857 | . | . | . | . | . |
| chr6:151808 | ESR1:NM_00 | . | . | . | 0.571428571 | . | . | . | . | . |
| chr6:167976 | HGC6.3:NM_ | . | 0.0027 | 0.0064 | NA | . | . | . | . | . |
| chr7:660801 | C7orf26:NM_ | . | . | . | 0.833333333 | . | . | . | . | . |
| chr7:325432 | AVL9:NM_01 | . | . | . | 0.857142857 | . | . | . | . | . |
| chr7:997078 | CYP3A7:NM_ | . | 6.368e-05 | 0.0024 | 0.857142857 | . | . | . | . | . |

| | | | | | | | | | | |
|---|---|---|---|---|---|---|---|---|---|---|
| chr7:151195 | IQCA1L:NM_ | . | . | . | NA | . | . | . | . | . |
| chr8:450074_ | FBXO25:NM_ | . | . | . | 0.857142857 | . | . | . | . | . |
| chr8:8318865_CGGGGCG/ | | . | . | . | NA | . | . | . | . | . |
| chr8:141494 | . | NM_207414: | . | . | NA | . | . | . | . | . |
| chr9:669899 | SPATA31A3:N | . | . | . | NA | . | . | . | . | . |
| chr9:136286 | CCDC187:NM | . | . | . | NA | . | . | . | . | . |
| chr9:137500 | PNPLA7:NM_ | . | . | . | 0.857142857 | . | . | . | . | . |
| chr10:47368 | ZNF488:NM_ | . | . | . | NA | . | . | . | . | . |
| chr10:47368 | ZNF488:NM_ | . | . | . | NA | . | . | . | . | . |
| chr10:47502 | AGAP9:NM_ | . | . | . | NA | . | . | . | . | . |
| chr10:47502 | AGAP9:NM_ | . | . | . | NA | . | . | . | . | . |
| chr10:47502 | AGAP9:NM_ | . | . | . | NA | . | . | . | . | . |
| chr10:10335 | PCGF6:NM_0 | . | . | . | NA | . | . | . | . | . |
| chr10:12449 | LHPP:NM_00 | . | . | . | 0.857142857 | . | . | . | . | . |
| chr11:87178 | ST5:NM_139 | . | . | . | 0.5 | . | . | . | . | . |
| chr11:24738 | LUZP2:NM_0 | . | . | . | 0.714285714 | . | . | . | . | . |
| chr11:34161 | ABTB2:NM_1 | . | . | . | 0.857142857 | . | . | . | . | . |
| chr11:54603 | OR4C46:NM_ | . | . | . | NA | . | . | . | . | . |
| chr11:54603 | OR4C46:NM_ | . | . | . | NA | . | . | . | . | . |
| chr11:54603 | OR4C46:NM_ | . | . | . | NA | . | . | . | . | . |
| chr11:54706 | OR4A5:NM_ | . | . | . | NA | . | . | . | . | . |
| chr11:64607 | NRXN2:NM_ | . | . | . | 0.666666666 | . | . | . | . | . |
| chr11:10306 | DCUN1D5:NI | . | . | . | 0.857142857 | . | . | . | . | . |
| chr11:11890 | BCL9L:NM_1 | . | . | . | 0.714285714 | . | . | . | . | . |
| chr11:12425 | OR8G1:NM_ | . | . | . | NA | . | . | . | . | . |
| chr11:12561 | STT3A:NM_0 | . | . | . | 0.857142857 | . | . | . | . | . |
| chr11:12594 | VSIG10L2:NM | . | . | . | NA | . | . | . | . | . |
| chr12:69367 | ATN1:NM_00 | . | . | . | NA | . | . | . | . | . |
| chr12:49795 | NCKAP5L:NM | . | . | . | 0.714285714 | . | . | . | . | . |
| chr13:77081 | MYCBP2:NM_ | . | . | . | 0.714285714 | . | . | . | . | . |
| chr14:19434 | POTEG:NM_ | . | . | . | NA | . | . | . | . | . |
| chr14:24057 | CARMIL3:NM | . | 0.0048 | 0.0073 | 0.714285714 | . | . | . | . | . |
| chr14:24177 | REC8:NM_00 | . | . | . | NA | . | . | . | . | . |
| chr14:68793 | ZFP36L1:NM_ | . | . | . | 1 | . | . | . | . | . |
| chr15:21422 | POTEB3:NM_ | . | . | . | 1 | . | . | . | . | . |
| chr15:22613 | GOLGA8J:NM | . | . | . | NA | . | . | . | . | . |

| | | | | | | | | | | |
|---|---|---|---|---|---|---|---|---|---|---|
| chr15:22903836_C/T | | . | . | . | NA | . | . | . | . | . |
| chr16:14330! | PERCC1:NM_ | . | . | . | NA | . | . | . | . | . |
| chr16:71536! | CHST4:NM_0 | . | . | . | 1 | . | . | . | . | . |
| chr16:88951. | LOC1001296! | . | 0.0020 | 0.0052 | NA | . | . | . | . | . |
| chr17:16438! | SCARF1:NM_ | . | . | . | 0.857142857 | . | . | . | . | . |
| chr17:31221! | NF1:NM_000 | . | . | . | NA | . | . | . | . | . |
| chr17:41097! | KRTAP4-8:NN | . | . | . | NA | . | . | . | . | . |
| chr17:41097! | KRTAP4-8:NN | . | . | . | NA | . | . | . | . | . |
| chr17:41097! | KRTAP4-8:NN | . | . | . | NA | . | . | . | . | . |
| chr17:41487: | KRT36:NM_0 | . | 6.371e-05 | 0.0012 | 0.857142857 | . | . | . | . | . |
| chr17:50168 | SGCA:NM_00 | . | 0.0004 | 0.0035 | 0.285714285 | 194779 | Limb-girdle_r | MedGen:C29 | criteria_prov | Conflicting_ir |
| chr18:49912: | MYO5B:NM_ | . | . | . | 1 | . | . | . | . | . |
| chr18:74447. | DIPK1C:NM_( | . | . | . | 1 | . | . | . | . | . |
| chr19:20768! | MOB3A:NM_ | . | . | . | 1 | . | . | . | . | . |
| chr19:86050 | ADAMTS10:N | . | . | . | NA | . | . | . | . | . |
| chr19:86985! | ACTL9:NM_1 | . | . | . | NA | . | . | . | . | . |
| chr19:41967( | ATP1A3:NM_ | . | . | . | 0.571428571 | . | . | . | . | . |
| chr19:51768! | FPR2:NM_00 | . | 0.0046 | 0.0046 | 0.714285714 | . | . | . | . | . |
| chr19:54735. | KIR3DL3:NM_ | . | . | . | NA | . | . | . | . | . |
| chr19:54832 | KIR2DS4:NM_ | . | . | . | NA | . | . | . | . | . |
| chr19:54835: | KIR2DS4:NM_ | . | . | . | NA | . | . | . | . | . |
| chr21:33794369_A/G | | . | . | . | 0.857142857 | . | . | . | . | . |
| chr21:45504! | COL18A1:NM | . | . | . | NA | . | . | . | . | . |
| chr22:21923 | PPM1F:NM_( | . | . | . | 0.857142857 | . | . | . | . | . |
| chr22:50283 | PLXNB2:NM_ | . | . | . | 0.714285714 | . | . | . | . | . |
| chrX:315246_ | GTPBP6:NM_ | . | . | . | NA | . | . | . | . | . |
| chrX:318607_ | GTPBP6:NM_ | . | . | . | NA | . | . | . | . | . |
| chrX:119329 | CRLF2:NM_0( | . | . | . | NA | . | . | . | . | . |
| chrX:155878! | ACE2:NM_00 | . | . | . | 0.857142857 | . | . | . | . | . |
| chrX:278219 | MAGEB10:NI | . | . | . | NA | . | . | . | . | . |
| chrX:844685! | HDX:NM_001 | . | . | . | 0.714285714 | . | . | . | . | . |
| chrX:152135( | MAGEA10:NI | . | . | . | 0.714285714 | . | . | . | . | . |
| chrX:152728: | CSAG1:NM_C | . | . | . | NA | . | . | . | . | . |

| SIFT_pred | Polyphen2_H | Polyphen2_H | LRT_pred | FATHMM_pr | MutationTast | MutationAss | InterVar_automated |
|---|---|---|---|---|---|---|---|
| T | P | P | D | T | D | M | Uncertain_significance |
| D | D | D | D | D | D | M | Uncertain_significance |
| D | D | P | D | T | D | M | Uncertain_significance |
| . | . | . | . | . | . | . | Uncertain_significance |
| . | . | . | . | . | . | . | . |
| . | . | . | . | . | . | . | Uncertain_significance |
| . | . | . | . | . | . | . | . |
| . | . | . | . | . | D | . | . |
| . | . | . | . | . | . | . | Uncertain_significance |
| D | D | D | U | T | D | M | Uncertain_significance |
| D | D | P | D | T | D | M | Uncertain_significance |
| . | . | . | . | . | . | . | Uncertain_significance |
| . | . | . | . | . | . | . | Uncertain_significance |
| . | . | . | . | . | . | . | Uncertain_significance |
| D | D | D | D | T | D | H | Uncertain_significance |
| D | D | D | D | T | D | H | Uncertain_significance |
| D | D | D | D | T | D | L | Uncertain_significance |
| D | D | D | D | T | D | M | Uncertain_significance |
| T | D | D | D | D | D | L | Uncertain_significance |
| . | . | . | . | . | . | . | . |
| D | D | D | . | T | D | M | Uncertain_significance |
| D | D | D | D | T | D | L | Uncertain_significance |
| . | . | . | . | . | . | . | Uncertain_significance |
| D | P | P | D | T | D | M | Uncertain_significance |
| . | . | . | . | . | . | . | . |
| . | . | . | . | . | . | . | . |
| D | D | D | D | T | D | M | Uncertain_significance |
| D | D | D | D | T | D | M | Uncertain_significance |
| D | D | D | D | T | D | H | Uncertain_significance |
| D | D | P | N | T | D | L | Likely_pathogenic |
| . | . | . | . | . | . | . | . |
| D | D | P | D | T | D | . | Uncertain_significance |
| D | D | D | D | T | D | M | Uncertain_significance |
| D | D | D | D | T | D | H | Uncertain_significance |

| | | | | | | | |
|---|---|---|---|---|---|---|---|
| . | . | . | . | . | . | . | . |
| D | D | D | D | T | D | M | Uncertain_significance |
| . | . | . | . | . | . | . | . |
| . | . | . | . | . | . | . | . |
| . | . | . | . | . | . | . | Uncertain_significance |
| . | . | . | . | . | . | . | Uncertain_significance |
| T | D | D | D | D | D | M | Uncertain_significance |
| . | . | . | . | . | . | . | Uncertain_significance |
| . | . | . | . | . | . | . | . |
| . | . | . | . | . | . | . | Uncertain_significance |
| . | . | . | . | . | . | . | Uncertain_significance |
| . | . | . | . | . | . | . | Uncertain_significance |
| . | . | . | . | . | . | . | . |
| D | D | D | D | T | D | H | Uncertain_significance |
| . | . | . | N | . | A | . | Pathogenic |
| D | D | D | D | T | D | L | Uncertain_significance |
| D | D | D | D | T | D | M | Uncertain_significance |
| . | . | . | . | . | . | . | Uncertain_significance |
| . | . | . | . | . | . | . | Uncertain_significance |
| . | . | . | . | . | . | . | Uncertain_significance |
| . | . | . | . | . | . | . | Uncertain_significance |
| D | D | P | . | T | D | N | . |
| D | D | D | D | T | D | M | Uncertain_significance |
| T | D | D | D | T | D | M | Uncertain_significance |
| . | . | . | . | . | . | . | Uncertain_significance |
| D | P | B | D | D | D | M | Uncertain_significance |
| . | . | . | . | . | . | . | . |
| . | . | . | . | . | . | . | . |
| D | D | D | D | T | N | M | Uncertain_significance |
| D | P | P | D | T | D | L | Uncertain_significance |
| . | . | . | . | . | . | . | Uncertain_significance |
| D | D | D | D | T | D | L | Uncertain_significance |
| . | . | . | . | . | . | . | . |
| D | . | . | . | . | D | . | Uncertain_significance |
| . | D | D | . | . | . | . | Uncertain_significance |
| . | . | . | . | . | . | . | Uncertain_significance |

| | | | | | | | |
|---|---|---|---|---|---|---|---|
| . | . | . | . | . | . | . | Uncertain_significance |
| . | . | . | . | . | . | . | . |
| D | D | D | D | D | D | M | Uncertain_significance |
| . | . | . | . | . | . | . | Uncertain_significance |
| D | D | D | D | T | D | H | Uncertain_significance |
| . | . | . | . | . | . | . | . |
| . | . | . | . | . | . | . | . |
| . | . | . | . | . | . | . | . |
| . | . | . | . | . | . | . | . |
| T | P | P | D | D | D | M | Uncertain_significance |
| T | P | B | N | D | N | L | Uncertain_significance |
| D | D | D | D | D | D | H | Uncertain_significance |
| . | . | . | . | . | A | . | Uncertain_significance |
| D | D | D | D | . | D | M | . |
| . | . | . | . | . | . | . | Uncertain_significance |
| . | . | . | . | . | . | . | Uncertain_significance |
| D | B | B | N | D | D | M | Likely_pathogenic |
| D | P | P | U | T | D | M | Uncertain_significance |
| . | . | . | . | . | . | . | Uncertain_significance |
| . | . | . | . | . | . | . | Uncertain_significance |
| . | . | . | . | . | . | . | Uncertain_significance |
| D | D | D | D | T | D | M | Uncertain_significance |
| . | . | . | . | . | . | . | . |
| D | D | D | D | T | D | M | Uncertain_significance |
| D | D | D | N | T | D | M | . |
| . | . | . | . | . | . | . | Uncertain_significance |
| . | . | . | . | . | . | . | Uncertain_significance |
| . | . | . | . | . | . | . | Uncertain_significance |
| D | D | D | D | T | D | H | Uncertain_significance |
| . | . | . | . | . | . | . | . |
| D | D | D | D | T | D | L | Uncertain_significance |
| D | D | D | D | T | N | M | Uncertain_significance |
| . | . | . | . | . | . | . | Uncertain_significance |

### NF90-8 cell line variants with a potential impact on protein function

| chr | start | end | Sample | Gene.refGen | REF | ALT | Func.refGene | ExonicFunc.r | VAF | avsnp150 |
|---|---|---|---|---|---|---|---|---|---|---|
| chr1 | 26282320 | 26282344 | 90-8TL | UBXN11 | TCCAGGACA( | T | exonic | nonframeshi | 0.969696969 | . |
| chr1 | 26889877 | 26889877 | 90-8TL | GPN2 | G | A | exonic | nonsynonym | 1 | . |
| chr1 | 144435174 | 144435174 | 90-8TL | NBPF15 | C | T | exonic | nonsynonym | 0.396226415 | rs201553147 |
| chr1 | 145898994 | 145898994 | 90-8TL | ITGA10 | C | T | exonic | nonsynonym | 0.611940298 | rs2274616 |
| chr1 | 146126370 | 146126370 | 90-8TL | NBPF10 | A | T | exonic | nonsynonym | 0.549019607 | . |
| chr1 | 148972321 | 148972321 | 90-8TL | PDE4DIP | C | T | exonic | nonsynonym | 1 | rs1628172 |
| chr1 | 149071644 | 149071644 | 90-8TL | NBPF9 | T | C | exonic | nonsynonym | 1 | . |
| chr1 | 149390858 | 149390858 | 90-8TL | NOTCH2NLC | G | T | exonic | nonsynonym | 0.454545454 | . |
| chr1 | 149554508 | 149554508 | 90-8TL | NBPF19 | C | G | exonic | nonsynonym | 0.4375 | . |
| chr1 | 152709204 | 152709204 | 90-8TL | LCE4A | C | CAGCCCTGG( | exonic | nonframeshi | 1 | . |
| chr1 | 154959406 | 154959427 | 90-8TL | PYGO2 | CATGGTGGG | C | exonic | nonframeshi | 0.509433962 | . |
| chr1 | 154959776 | 154959776 | 90-8TL | PYGO2 | G | A | exonic | nonsynonym | 0.36 | . |
| chr2 | 46480669 | 46480669 | 90-8TL | TMEM247 | C | G | exonic | nonsynonym | 0.28 | rs74318890 |
| chr2 | 196433390 | 196433390 | 90-8TL | HECW2 | C | CA | exonic | frameshift_ir | 0.44 | . |
| chr2 | 200490234 | 200490234 | 90-8TL | KCTD18 | G | A | exonic | nonsynonym | 0.423076923 | . |
| chr2 | 218828546 | 218828546 | 90-8TL | PRKAG3 | A | G | exonic | nonsynonym | 0.512820512 | . |
| chr3 | 51988821 | 51988821 | 90-8TL | ABHD14A-AC | C | T | exonic | nonsynonym | 1 | rs121912698 |
| chr3 | 57375913 | 57375919 | 90-8TL | DNAH12 | CGCCAGT | C | exonic | nonframeshi | 1 | . |
| chr3 | 112078880 | 112078880 | 90-8TL | TMPRSS7 | T | A | splicing | . | 1 | . |
| chr3 | 115676067 | 115676067 | 90-8TL | GAP43 | G | A | exonic | nonsynonym | 0.142857142 | . |
| chr3 | 170084745 | 170084745 | 90-8TL | GPR160 | T | C | exonic | nonsynonym | 1 | . |
| chr3 | 183775955 | 183775961 | 90-8TL | YEATS2 | CGGAGGA | CGGA | exonic | nonframeshi | 1 | . |
| chr3 | 186572190 | 186572190 | 90-8TL | DNAJB11 | A | G | exonic | nonsynonym | 0.681818181 | . |
| chr4 | 1406310 | 1406310 | 90-8TL | NKX1-1 | G | T | exonic | nonsynonym | 0.391304347 | . |
| chr4 | 44650040 | 44650040 | 90-8TL | YIPF7 | G | T | exonic | nonsynonym | 0.414634146 | . |
| chr5 | 67142238 | 67142238 | 90-8TL | MAST4 | G | C | splicing | . | 0.673469387 | . |
| chr5 | 95682836 | 95682836 | 90-8TL | SPATA9 | C | T | exonic | nonsynonym | 0.142857142 | . |
| chr5 | 141184133 | 141184133 | 90-8TL | PCDHB16 | A | G | exonic | nonsynonym | 1 | rs17844651 |
| chr6 | 41070093 | 41070093 | 90-8TL | OARD1 | G | A | exonic | stopgain | 0.305084745 | rs144679838 |
| chr6 | 43046583 | 43046583 | 90-8TL | CUL7 | C | T | exonic | nonsynonym | 0.346938775 | rs200040003 |
| chr6 | 142753451 | 142753451 | 90-8TL | HIVEP2 | T | G | exonic | nonsynonym | 0.318181818 | . |
| chr7 | 33383786 | 33383786 | 90-8TL | BBS9 | C | T | exonic | nonsynonym | 0.603448275 | rs771777721 |
| chr7 | 73704186 | 73704186 | 90-8TL | STX1A | T | C | exonic | nonsynonym | 1 | . |
| chr8 | 120516097 | 120516097 | 90-8TL | MTBP | C | A | exonic | nonsynonym | 0.386363636 | . |

| | | | | | | | | | | |
|---|---|---|---|---|---|---|---|---|---|---|
| chr8 | 130783730 | 130783730 | 90-8TL | ADCY8 | T | A | exonic | nonsynonym | 0.275362318 | . |
| chr9 | 5920406 | 5920411 | 90-8TL | KIAA2026 | CGACTA | C | exonic | frameshift_d | 1 | . |
| chr9 | 34372875 | 34372875 | 90-8TL | MYORG | G | C | exonic | stopgain | 1 | rs4879782 |
| chr9 | 35906586 | 35906586 | 90-8TL | HRCT1 | T | TCCA | exonic | nonframeshif | 0.636363636 | . |
| chr9 | 41991703 | 41991703 | 90-8TL | CNTNAP3B | G | T | exonic | nonsynonym | 0.5 | rs62555055 |
| chr9 | 41991721 | 41991721 | 90-8TL | CNTNAP3B | G | A | exonic | nonsynonym | 0.514285714 | rs3739623 |
| chr9 | 75077186 | 75077186 | 90-8TL | NMRK1 | C | A | exonic | nonsynonym | 0.372093023 | . |
| chr10 | 45826273 | 45826273 | 90-8TL | AGAP4 | C | A | exonic | nonsynonym | 0.615384615 | . |
| chr10 | 47368085 | 47368085 | 90-8TL | ZNF488 | G | A | exonic | nonsynonym | 0.361702127 | rs3814160 |
| chr10 | 47368615 | 47368615 | 90-8TL | ZNF488 | G | A | exonic | nonsynonym | 0.318181818 | rs35618062 |
| chr10 | 122833767 | 122833767 | 90-8TL | CUZD1 | C | T | exonic | nonsynonym | 0.142857142 | . |
| chr10 | 124044601 | 124044601 | 90-8TL | CHST15 | A | T | exonic | nonsynonym | 0.318181818 | . |
| chr11 | 89968879 | 89968880 | 90-8TL | TRIM64 | AG | A | exonic | frameshift_d | 1 | . |
| chr11 | 96092210 | 96092222 | 90-8TL | MAML2 | TTGCTGCTGC | TTGC | exonic | nonframeshif | 1 | . |
| chr11 | 108541766 | 108541766 | 90-8TL | EXPH5 | T | C | exonic | nonsynonym | 0.166666666 | . |
| chr12 | 54576638 | 54576638 | 90-8TL | PDE1B | C | T | exonic | nonsynonym | 0.238095238 | rs764032346 |
| chr12 | 56322464 | 56322464 | 90-8TL | PAN2 | A | T | exonic | nonsynonym | 1 | . |
| chr12 | 103993455 | 103993455 | 90-8TL | GLT8D2 | T | C | exonic | nonsynonym | 1 | rs145520946 |
| chr13 | 78601731 | 78601731 | 90-8TL | POU4F1 | A | G | exonic | nonsynonym | 0.333333333 | . |
| chr14 | 64771032 | 64771032 | 90-8TL | SPTB | G | A | exonic | nonsynonym | 0.276595744 | rs148337824 |
| chr14 | 92071010 | 92071010 | 90-8TL | ATXN3 | C | CCTGCTGCTG | exonic | nonframeshif | 0.733333333 | . |
| chr15 | 51576183 | 51576183 | 90-8TL | DMXL2 | T | TAAAAAAAA | splicing | . | 0.629629629 | . |
| chr15 | 85244644 | 85244644 | 90-8TL | GOLGA6L3 | G | T | exonic | nonsynonym | 0.230769230 | . |
| chr16 | 53956733 | 53956733 | 90-8TL | FTO | G | C | exonic | nonsynonym | 125 | . |
| chr16 | 70468868 | 70468868 | 90-8TL | FCSK | T | A | exonic | nonsynonym | 1 | rs201690930 |
| chr17 | 7676381 | 7676381 | 90-8TL | TP53 | C | T | splicing | . | 1 | . |
| chr17 | 9771541 | 9771541 | 90-8TL | DHRS7C | C | T | exonic | nonsynonym | 0.509433962 | rs199841658 |
| chr17 | 10512902 | 10512902 | 90-8TL | MYH1 | T | TA | exonic | frameshift_ir | 1 | rs545765873 |
| chr17 | 21703340 | 21703340 | 90-8TL | KCNJ18 | C | T | exonic | nonsynonym | 0.24 | . |
| chr17 | 31235950 | 31235957 | 90-8TL | NF1 | GGATCCTT | G | exonic | frameshift_d | 1 | . |
| chr17 | 31993330 | 31993330 | 90-8TL | SUZ12 | T | A | exonic | stopgain | 1 | . |
| chr17 | 39727923 | 39727923 | 90-8TL | ERBB2 | C | A | exonic | nonsynonym | 0.762711864 | rs55943169 |
| chr17 | 43016519 | 43016519 | 90-8TL | VAT1 | G | A | exonic | nonsynonym | 0.354166666 | . |
| chr17 | 43884083 | 43884083 | 90-8TL | MPP2 | C | T | splicing | . | 0.707692307 | rs231518 |
| chr17 | 49973193 | 49973193 | 90-8TL | DLX4 | G | C | exonic | nonsynonym | 0.628571428 | rs151318731 |
| chr17 | 50999702 | 50999702 | 90-8TL | SPAG9 | A | C | exonic | nonsynonym | 0.553191489 | . |

| | | | | | | | | | | |
|---|---|---|---|---|---|---|---|---|---|---|
| chr17 | 81647906 | 81647906 | 90-8TL | TSPAN10 | T | TTAAC | exonic | stopgain | 1 | . |
| chr18 | 712349 | 712367 | 90-8TL | ENOSF1 | GCCGCGCGT/ | G | exonic | frameshift_d | 0.894736842 | . |
| chr18 | 14755006 | 14755006 | 90-8TL | ANKRD30B | G | T | splicing | . | 0.45 | . |
| chr18 | 46253735 | 46253738 | 90-8TL | C18orf25 | TCTG | T | exonic | nonframeshi | 0.735294117 | . |
| chr18 | 62575320 | 62575320 | 90-8TL | ZCCHC2 | A | G | exonic | nonsynonym | 0.344262295 | . |
| chr19 | 8086248 | 8086248 | 90-8TL | FBN3 | C | G | exonic | nonsynonym | 0.476190476 | rs144887983 |
| chr19 | 13207858 | 13207864 | 90-8TL | CACNA1A | CCTGCTG | CCTG | exonic | nonframeshi | 1 | . |
| chr19 | 41577497 | 41577497 | 90-8TL | CEACAM21 | A | C | exonic | nonsynonym | 0.515625 | rs714106 |
| chr19 | 41579520 | 41579520 | 90-8TL | CEACAM21 | G | A | exonic | nonsynonym | 0.492537313 | rs2302188 |
| chr19 | 41717688 | 41717688 | 90-8TL | CEACAM5 | G | A | exonic | nonsynonym | 0.524390243 | rs7249230 |
| chr19 | 41761981 | 41761981 | 90-8TL | CEACAM6 | T | G | exonic | nonsynonym | 1 | rs11548735 |
| chr19 | 54735264 | 54735264 | 90-8TL | KIR3DL3 | A | T | exonic | nonsynonym | 0.5 | rs602444 |
| chr19 | 54847200 | 54847200 | 90-8TL | KIR2DS1\x3b | A | G | exonic | nonsynonym | 0.339805825 | rs113664264 |
| chr20 | 4248011 | 4248011 | 90-8TL | ADRA1D | T | TCGGCGCC | exonic | frameshift_ir | 1 | . |
| chr20 | 16427109 | 16427109 | 90-8TL | KIF16B | T | C | exonic | nonsynonym | 1 | . |
| chr20 | 25292583 | 25292583 | 90-8TL | PYGB | T | G | exonic | nonsynonym | 0.571428571 | . |
| chr20 | 59023585 | 59023585 | 90-8TL | TUBB1 | G | A | exonic | nonsynonym | 0.447761194 | rs150551805 |
| chr20 | 63977367 | 63977367 | 90-8TL | SAMD10 | T | G | exonic | nonsynonym | 0.513513513 | rs145097158 |
| chr22 | 22515221 | 22515221 | 90-8TL | ZNF280A | G | T | exonic | nonsynonym | 0.289473684 | rs361580 |
| chr22 | 22515224 | 22515224 | 90-8TL | ZNF280A | C | T | exonic | nonsynonym | 0.289473684 | rs362011 |
| chr22 | 23998589 | 23998589 | 90-8TL | GSTT4 | A | G | exonic | nonsynonym | 0.371428571 | . |
| chr22 | 23998678 | 23998678 | 90-8TL | GSTT4 | T | C | exonic | nonsynonym | 0.358974358 | . |
| chrX | 46607148 | 46607148 | 90-8TL | SLC9A7 | A | G | exonic | nonsynonym | 0.181818181 | . |
| chrX | 51407918 | 51407951 | 90-8TL | EZHIP | CCCGAGGCC( | C | exonic | nonframeshi | 1 | . |
| chrX | 153073248 | 153073248 | 90-8TL | PNMA6A | C | CA | exonic | frameshift_ir | 0.5 | . |

| Variant | AAChange.re | GeneDetail.r | AF | AF_popmax | prop_pathog | CLNALLELEID | CLNDN | CLNDISDB | CLNREVSTAT | CLNSIG |
|---|---|---|---|---|---|---|---|---|---|---|
| chr1:262823 | UBXN11:NM_ | . | 0.0027 | 0.0035 | NA | . | . | . | . | . |
| chr1:268898 | GPN2:NM_0 | . | . | . | 0.714285714 | . | . | . | . | . |
| chr1:144435 | NBPF15:NM_ | . | . | . | NA | . | . | . | . | . |
| chr1:145898 | ITGA10:NM_ | . | . | . | NA | . | . | . | . | . |
| chr1:146126 | NBPF10:NM_ | . | . | . | NA | . | . | . | . | . |
| chr1:148972321_C/T | | . | . | . | NA | . | . | . | . | . |
| chr1:149071 | NBPF9:NM_0 | . | . | . | NA | . | . | . | . | . |
| chr1:149390 | NOTCH2NLC: | . | . | . | NA | . | . | . | . | . |
| chr1:149554 | NBPF19:NM_ | . | . | . | 0.75 | . | . | . | . | . |
| chr1:152709 | LCE4A:NM_1 | . | . | . | NA | . | . | . | . | . |
| chr1:154959 | PYGO2:NM_ | . | . | . | NA | . | . | . | . | . |
| chr1:154959 | PYGO2:NM_ | . | . | . | 0.714285714 | . | . | . | . | . |
| chr2:464806 | TMEM247:N | . | . | . | 0.8 | . | . | . | . | . |
| chr2:196433 | HECW2:NM_ | . | . | . | NA | . | . | . | . | . |
| chr2:200490 | KCTD18:NM_ | . | . | . | 0.714285714 | . | . | . | . | . |
| chr2:218828 | PRKAG3:NM_ | . | . | . | 1 | . | . | . | . | . |
| chr3:519888 | ACY1:NM_00 | . | 0.0030 | 0.0048 | 0.857142857 | 33149 | Aminoacylas | MedGen:C18 | criteria_prov | Conflicting_i |
| chr3:573759 | DNAH12:NM | . | . | . | NA | . | . | . | . | . |
| chr3:1120788 | . | NM_0010425 | . | . | 1 | . | . | . | . | . |
| chr3:115676 | GAP43:NM_ | . | . | . | 0.857142857 | . | . | . | . | . |
| chr3:170084 | GPR160:NM_ | . | . | . | 1 | . | . | . | . | . |
| chr3:183775 | YEATS2:NM_ | . | . | . | NA | . | . | . | . | . |
| chr3:186572 | DNAJB11:NM | . | . | . | 1 | . | . | . | . | . |
| chr4:140631 | NKX1-1:NM_ | . | 0.0007 | 0.0013 | 1 | . | . | . | . | . |
| chr4:446500 | YIPF7:NM_18 | . | . | . | 0.857142857 | . | . | . | . | . |
| chr5:671422 | . | NM_0011646 | . | . | 1 | . | . | . | . | . |
| chr5:956828 | SPATA9:NM_ | . | . | . | 0.714285714 | . | . | . | . | . |
| chr5:141184 | PCDHB16:NM | . | . | . | NA | . | . | . | . | . |
| chr6:41070093_G/A | | . | 0.0020 | 0.0025 | 1 | . | . | . | . | . |
| chr6:430465 | CUL7:NM_00 | . | 0.0003 | 0.0012 | 0.714285714 | 189014 | not_providec | MedGen:CN | criteria_prov | Likely_patho |
| chr6:142753 | HIVEP2:NM_ | . | . | . | 0.857142857 | . | . | . | . | . |
| chr7:33383786_C/T | | . | . | . | 0.714285714 | 306191 | Bardet-Biedl_ | MedGen:C07 | criteria_prov | Uncertain_si |
| chr7:737041 | STX1A:NM_0 | . | . | . | 0.714285714 | . | . | . | . | . |
| chr8:120516 | MTBP:NM_0 | . | . | . | 1 | . | . | . | . | . |

| | | | | | | | | | | |
|---|---|---|---|---|---|---|---|---|---|---|
| chr8:130783 | ADCY8:NM_C | . | . | . | 0.714285714 | . | . | . | . | . |
| chr9:592040 | KIAA2026:NM | . | . | . | NA | . | . | . | . | . |
| chr9:343728 | MYORG:NM_ | . | . | . | NA | . | . | . | . | . |
| chr9:359065 | HRCT1:NM_C | . | . | . | NA | . | . | . | . | . |
| chr9:419917 | CNTNAP3B:N | . | . | . | NA | . | . | . | . | . |
| chr9:419917 | CNTNAP3B:N | . | . | . | NA | . | . | . | . | . |
| chr9:750771 | NMRK1:NM_ | . | . | . | 0.857142857 | . | . | . | . | . |
| chr10:45826 | AGAP4:NM_ | . | 4.786e-05 | 1 | 0.857142857 | . | . | . | . | . |
| chr10:47368 | ZNF488:NM_ | . | . | . | NA | . | . | . | . | . |
| chr10:47368 | ZNF488:NM_ | . | . | . | NA | . | . | . | . | . |
| chr10:12283 | CUZD1:NM_C | . | . | . | 0.857142857 | . | . | . | . | . |
| chr10:12404 | CHST15:NM_ | . | . | . | 0.857142857 | . | . | . | . | . |
| chr11:89968 | TRIM64:NM_ | . | 0.0026 | 0.0071 | NA | . | . | . | . | . |
| chr11:96092 | MAML2:NM_ | . | . | . | NA | . | . | . | . | . |
| chr11:10854 | EXPH5:NM_0 | . | . | . | 0.714285714 | . | . | . | . | . |
| chr12:54576 | PDE1B:NM_C | . | . | . | 0.714285714 | . | . | . | . | . |
| chr12:56322 | PAN2:NM_0C | . | . | . | 0.857142857 | . | . | . | . | . |
| chr12:10399 | GLT8D2:NM_ | . | 0.0057 | 0.0092 | 0.714285714 | . | . | . | . | . |
| chr13:78601 | POU4F1:NM_ | . | . | . | 1 | . | . | . | . | . |
| chr14:64771 | SPTB:NM_00 | . | 0.0023 | 0.0039 | 0.857142857 | 264543 | not_specifiec | MedGen:CN1 | criteria_prov | Uncertain_sig |
| chr14:92071010_C/CCTGC | | | . | . | NA | . | . | . | . | . |
| chr15:51576 | . | NM_015263: | . | . | NA | . | . | . | . | . |
| chr15:85244 | GOLGA6L3:N | . | 6.869e-05 | 0.0014 | NA | . | . | . | . | . |
| chr16:53956 | FTO:NM_001 | . | . | . | NA | . | . | . | . | . |
| chr16:70468 | FCSK:NM_14 | . | 0.0035 | 0.0046 | 0.857142857 | . | . | . | . | . |
| chr17:76763 | . | | . | . | 1 | . | . | . | . | . |
| chr17:97715 | DHRS7C:NM_ | . | 0.0009 | 0.0014 | 0.714285714 | . | . | . | . | . |
| chr17:10512 | MYH1:NM_0 | . | 0.0009 | 0.0012 | NA | . | . | . | . | . |
| chr17:21703 | KCNJ18:NM_ | . | . | . | 0.857142857 | . | . | . | . | . |
| chr17:31235 | NF1:NM_000 | . | . | . | NA | . | . | . | . | . |
| chr17:31993 | SUZ12:NM_0 | . | . | . | 1 | . | . | . | . | . |
| chr17:39727 | ERBB2:NM_0 | . | 0.0063 | 7 | 0.666666666 | 137823 | not_specifiec | MedGen:CN1 | no_assertion | not_providec |
| chr17:43016 | VAT1:NM_00 | . | . | . | 0.857142857 | . | . | . | . | . |
| chr17:43884 | . | NM_001278 | . | . | NA | . | . | . | . | . |
| chr17:49973 | DLX4:NM_00 | . | 0.0028 | 0.0042 | 1 | . | . | . | . | . |
| chr17:50999 | SPAG9:NM_C | . | . | . | 0.714285714 | . | . | . | . | . |

| | | | | | | | | | | |
|---|---|---|---|---|---|---|---|---|---|---|
| chr17:81647 | TSPAN10:NM | . | . | . | NA | . | . | . | . | . |
| chr18:71234 | ENOSF1:NM_ | . | . | . | NA | . | . | . | . | . |
| chr18:14755 | . | NM_001367 | . | . | 0 | . | . | . | . | . |
| chr18:46253 | C18orf25:NM | . | . | . | NA | . | . | . | . | . |
| chr18:62575 | ZCCHC2:NM_ | . | . | . | 0.857142857 | . | . | . | . | . |
| chr19:80862 | FBN3:NM_00 | . | 0.0061 | 0.0088 | 0.714285714 | . | . | . | . | . |
| chr19:13207 | CACNA1A:NN | . | . | . | NA | . | . | . | . | . |
| chr19:41577 | CEACAM21:N | . | . | . | NA | . | . | . | . | . |
| chr19:41579 | CEACAM21:N | . | . | . | NA | . | . | . | . | . |
| chr19:41717 | CEACAM5:NN | . | . | . | NA | . | . | . | . | . |
| chr19:41761 | CEACAM6:NN | . | . | . | NA | . | . | . | . | . |
| chr19:54735 | KIR3DL3:NM_ | . | . | . | NA | . | . | . | . | . |
| chr19:54847 | KIR2DS4:NM_ | . | . | . | NA | . | . | . | . | . |
| chr20:42480 | ADRA1D:NM_ | . | . | . | NA | . | . | . | . | . |
| chr20:16427 | KIF16B:NM_ | . | . | . | 0.714285714 | . | . | . | . | . |
| chr20:25292 | PYGB:NM_00 | . | . | . | 0.714285714 | . | . | . | . | . |
| chr20:59023 | TUBB1:NM_ | . | . | . | 0.857142857 | . | . | . | . | . |
| chr20:63977 | SAMD10:NM_ | . | 0.0067 | 0.0082 | 0.833333333 | . | . | . | . | . |
| chr22:22515 | ZNF280A:NM | . | . | . | NA | . | . | . | . | . |
| chr22:22515 | ZNF280A:NM | . | . | . | NA | . | . | . | . | . |
| chr22:23998 | GSTT4:NM_0 | . | . | . | NA | . | . | . | . | . |
| chr22:23998 | GSTT4:NM_0 | . | 0 | . | NA | . | . | . | . | . |
| chrX:466071 | SLC9A7:NM_ | . | . | . | 0.857142857 | . | . | . | . | . |
| chrX:514079 | EZHIP:NM_2 | . | 0.0006 | 0.0016 | NA | . | . | . | . | . |
| chrX:153073 | PNMA6A:NM | . | . | . | NA | . | . | . | . | . |

| SIFT_pred | Polyphen2_H | Polyphen2_H | LRT_pred | FATHMM_pr | MutationTast | MutationAss | InterVar_automated |
|---|---|---|---|---|---|---|---|
| . | . | . | . | . | . | . | . |
| D | P | P | N | T | D | M | Uncertain_significance |
| . | . | . | . | . | . | . | . |
| . | . | . | . | . | . | . | . |
| . | . | . | . | . | . | . | . |
| . | . | . | . | . | . | . | Uncertain_significance |
| . | . | . | . | . | . | . | Uncertain_significance |
| . | . | . | . | . | . | . | . |
| . | D | P | . | . | N | M | Uncertain_significance |
| . | . | . | . | . | . | . | . |
| . | . | . | . | . | . | . | . |
| D | D | P | D | T | D | L | Uncertain_significance |
| D | D | D | N | . | . | M | Uncertain_significance |
| . | . | . | . | . | . | . | . |
| T | D | D | D | T | D | M | Uncertain_significance |
| D | D | D | D | D | D | H | . |
| D | D | D | D | T | A | H | Uncertain_significance |
| . | . | . | . | . | . | . | . |
| . | . | . | . | . | D | . | . |
| D | D | D | D | T | D | M | Uncertain_significance |
| D | D | D | D | . | D | M | Uncertain_significance |
| . | . | . | . | . | . | . | . |
| D | D | D | D | D | D | H | Uncertain_significance |
| D | . | . | . | D | . | . | . |
| D | D | D | D | T | D | M | Uncertain_significance |
| . | . | . | . | . | D | . | . |
| D | D | D | D | T | D | L | Uncertain_significance |
| . | . | . | . | . | . | . | Uncertain_significance |
| . | . | . | D | . | A | . | Uncertain_significance |
| D | D | D | D | T | D | L | Uncertain_significance |
| D | D | D | D | T | D | M | Uncertain_significance |
| D | P | B | D | T | D | M | Uncertain_significance |
| D | P | B | D | T | D | M | Uncertain_significance |
| D | D | P | D | . | D | M | Uncertain_significance |

| | | | | | | | |
|---|---|---|---|---|---|---|---|
| D | P | P | D | T | D | L | Uncertain_significance |
| . | . | . | . | . | . | . | . |
| . | . | . | . | . | . | . | Uncertain_significance |
| . | . | . | . | . | . | . | . |
| . | . | . | . | . | . | . | Uncertain_significance |
| . | . | . | . | . | . | . | . |
| D | D | D | D | T | D | H | . |
| D | P | P | D | T | D | M | Uncertain_significance |
| . | . | . | . | . | . | . | Uncertain_significance |
| . | . | . | . | . | . | . | . |
| D | D | D | D | T | D | M | Uncertain_significance |
| D | D | D | D | T | D | M | Uncertain_significance |
| . | . | . | . | . | . | . | . |
| . | . | . | . | . | . | . | . |
| D | D | D | D | T | D | L | Uncertain_significance |
| D | D | D | D | T | D | L | Uncertain_significance |
| D | D | D | D | T | D | M | . |
| T | D | D | D | T | D | M | Uncertain_significance |
| D | D | P | D | D | D | M | Uncertain_significance |
| D | D | P | D | T | D | M | Uncertain_significance |
| . | . | . | . | . | . | . | . |
| . | . | . | . | . | . | . | . |
| . | . | . | . | . | . | . | Uncertain_significance |
| . | . | . | . | . | . | . | . |
| D | D | D | D | T | D | M | Uncertain_significance |
| . | . | . | . | . | D | . | . |
| T | D | D | D | T | D | M | Uncertain_significance |
| . | . | . | . | . | . | . | . |
| D | D | B | D | D | D | M | Uncertain_significance |
| . | . | . | . | . | . | . | . |
| . | . | . | D | . | A | . | Likely_pathogenic |
| D | B | B | . | D | D | M | Uncertain_significance |
| D | D | D | D | T | D | M | Uncertain_significance |
| . | . | . | . | . | . | . | . |
| D | D | D | . | D | D | M | Uncertain_significance |
| T | D | D | D | T | D | M | . |

| | | | | | | | |
|---|---|---|---|---|---|---|---|
| . | . | . | . | . | . | . | . |
| . | . | . | . | . | . | . | . |
| . | . | . | . | . | N | . | . |
| . | . | . | . | . | . | . | . |
| D | D | D | D | T | D | M | Uncertain_significance |
| D | D | D | U | D | D | N | . |
| . | . | . | . | . | . | . | . |
| . | . | . | . | . | . | . | Uncertain_significance |
| . | . | . | . | . | . | . | Uncertain_significance |
| . | . | . | . | . | . | . | Uncertain_significance |
| . | . | . | . | . | . | . | Uncertain_significance |
| . | . | . | . | . | . | . | Uncertain_significance |
| . | . | . | . | . | . | . | Uncertain_significance |
| . | . | . | . | . | . | . | . |
| T | P | P | D | D | D | N | . |
| D | B | B | D | D | D | H | Uncertain_significance |
| D | D | P | D | T | D | M | Uncertain_significance |
| D | D | D | D | . | D | L | Uncertain_significance |
| . | . | . | . | . | . | . | Uncertain_significance |
| . | . | . | . | . | . | . | Uncertain_significance |
| . | . | . | . | . | . | . | . |
| . | . | . | . | . | . | . | . |
| D | D | D | D | T | D | M | Uncertain_significance |
| . | . | . | . | . | . | . | . |
| . | . | . | . | . | . | . | . |

### SNF96-2 cell line variants with a potential impact on protein function

| chr | start | end | Sample | Gene.refGen | REF | ALT | Func.refGene | ExonicFunc.r | VAF | avsnp150 |
|---|---|---|---|---|---|---|---|---|---|---|
| chr1 | 26282344 | 26282362 | SNF96.2 | UBXN11 | ACCGGGACC | A | exonic | nonframeshi | 1 | . |
| chr1 | 26345087 | 26345087 | SNF96.2 | CRYBG2 | A | G | exonic | nonsynonym | 1 | rs11579835 |
| chr1 | 40862096 | 40862096 | SNF96.2 | CITED4 | T | G | exonic | nonsynonym | 0.4 | . |
| chr1 | 145873487 | 145873487 | SNF96.2 | ANKRD35 | G | A | exonic | nonsynonym | 1 | rs6670984 |
| chr1 | 146067225 | 146067225 | SNF96.2 | NBPF10 | C | A | exonic | nonsynonym | 1 | . |
| chr1 | 146067283 | 146067283 | SNF96.2 | NBPF10 | C | T | exonic | nonsynonym | 1 | . |
| chr1 | 148595576 | 148595576 | SNF96.2 | NBPF14 | C | A | exonic | nonsynonym | 1 | . |
| chr1 | 150077136 | 150077136 | SNF96.2 | VPS45 | G | A | exonic | nonsynonym | 1 | rs200134576 |
| chr1 | 154010094 | 154010094 | SNF96.2 | NUP210L | G | A | exonic | nonsynonym | 0.136363636 | . |
| chr1 | 159890480 | 159890481 | SNF96.2 | CFAP45 | AT | A | exonic | frameshift_d | 0.414634146 | . |
| chr1 | 173914743 | 173914743 | SNF96.2 | SERPINC1 | G | A | exonic | nonsynonym | 1 | rs121909551 |
| chr2 | 99554341 | 99554341 | SNF96.2 | AFF3 | T | A | exonic | nonsynonym | 1 | . |
| chr2 | 112564485 | 112564485 | SNF96.2 | POLR1B | C | G | exonic | nonsynonym | 1 | . |
| chr2 | 117944023 | 117944023 | SNF96.2 | CCDC93 | C | A | splicing | . | 1 | . |
| chr2 | 166199787 | 166199787 | SNF96.2 | SCN9A | C | A | exonic | nonsynonym | 0.521739130 | . |
| chr2 | 227008117 | 227008117 | SNF96.2 | COL4A4 | C | A | exonic | nonsynonym | 0.441176470 | . |
| chr2 | 236580947 | 236580947 | SNF96.2 | ACKR3 | G | A | exonic | nonsynonym | 0.384615384 | . |
| chr3 | 8768070 | 8768070 | SNF96.2 | OXTR | G | A | exonic | nonsynonym | 1 | . |
| chr3 | 12004864 | 12004864 | SNF96.2 | SYN2 | C | G | exonic | nonsynonym | 1 | rs2923857 |
| chr3 | 49125327 | 49125327 | SNF96.2 | LAMB2 | A | T | exonic | nonsynonym | 1 | . |
| chr3 | 121576878 | 121576879 | SNF96.2 | ARGFX | CT | C | exonic | frameshift_d | 0.409090909 | . |
| chr3 | 127575494 | 127575502 | SNF96.2 | TPRA1 | AGCTCCTCC | A | exonic | frameshift_d | 0.386363636 | . |
| chr3 | 165030771 | 165030771 | SNF96.2 | SI | G | A | exonic | nonsynonym | 1 | . |
| chr4 | 139730430 | 139730436 | SNF96.2 | MAML3 | TCTGCTG | TCTG | exonic | nonframeshi | 1 | . |
| chr4 | 184011422 | 184011422 | SNF96.2 | STOX2 | C | T | exonic | nonsynonym | 1 | rs201760642 |
| chr5 | 141184153 | 141184153 | SNF96.2 | PCDHB16 | G | A | exonic | nonsynonym | 1 | rs2697532 |
| chr6 | 16327684 | 16327684 | SNF96.2 | ATXN1 | A | ATGCTGCTG | exonic | nonframeshi | 0.947368421 | . |
| chr6 | 32530201 | 32530201 | SNF96.2 | HLA-DRB5 | T | TCCAG | exonic | frameshift_ir | 0.357142857 | . |
| chr6 | 167976591 | 167976591 | SNF96.2 | HGC6.3 | G | A | exonic | nonsynonym | 0.333333333 | rs61740140 |
| chr7 | 984491 | 984491 | SNF96.2 | CYP2W1 | T | A | exonic | nonsynonym | 0.771428571 | . |
| chr7 | 45920875 | 45920875 | SNF96.2 | IGFBP3 | C | T | exonic | nonsynonym | 0.266666666 | . |
| chr7 | 74779322 | 74779322 | SNF96.2 | NCF1 | A | G | exonic | nonsynonym | 0.255813953 | rs10614 |
| chr7 | 104328901 | 104328901 | SNF96.2 | LHFPL3 | C | T | exonic | nonsynonym | 0.322580645 | rs373201390 |
| chr7 | 123452787 | 123452788 | SNF96.2 | IQUB | CT | C | exonic | frameshift_d | 0.254901960 | . |

| | | | | | | | | | | |
|---|---|---|---|---|---|---|---|---|---|---|
| chr7 | 129727248 | 129727248 | SNF96.2 | NRF1 | G | C | exonic | nonsynonym | 0.347826086 | . |
| chr8 | 8318865 | 8318871 | SNF96.2 | PRAG1 | CGGGGCG | C | exonic | nonframeshi | 0.521739130 | . |
| chr8 | 8377352 | 8377358 | SNF96.2 | PRAG1 | CGCCGCT | C | exonic | nonframeshi | 1 | . |
| chr8 | 11808709 | 11808715 | SNF96.2 | FDFT1 | GTCCCAC | GTCCCACTCC | exonic | nonframeshi | 1 | . |
| chr8 | 94519116 | 94519117 | SNF96.2 | VIRMA | TC | T | exonic | frameshift_d | 0.24 | . |
| chr8 | 131944802 | 131944802 | SNF96.2 | EFR3A | G | A | exonic | nonsynonym | 0.319444444 | . |
| chr8 | 141494938 | 141494938 | SNF96.2 | MROH5 | C | T | splicing | . | 0.640625 | rs6578193 |
| chr8 | 144137996 | 144137996 | SNF96.2 | HGH1 | G | T | exonic | nonsynonym | 0.428571428 | . |
| chr8 | 144379425 | 144379425 | SNF96.2 | ADCK5 | C | A | exonic | nonsynonym | 1 | rs6599528 |
| chr9 | 35295783 | 35295783 | SNF96.2 | UNC13B | C | T | exonic | nonsynonym | 1 | rs138440338 |
| chr9 | 66141076 | 66141080 | SNF96.2 | ANKRD20A1\ | TTAAG | T | exonic | frameshift_d | 0.354838709 | . |
| chr9 | 93263923 | 93263923 | SNF96.2 | WNK2 | A | G | exonic | nonsynonym | 0.142857142 | . |
| chr9 | 97650564 | 97650565 | SNF96.2 | NCBP1 | GC | G | exonic | stopgain | 0.458333333 | . |
| chr9 | 129868144 | 129868144 | SNF96.2 | USP20 | G | A | exonic | nonsynonym | 1 | rs148425010 |
| chr9 | 129900016 | 129900016 | SNF96.2 | FNBP1 | C | A | exonic | stopgain | 0.578947368 | . |
| chr9 | 130617799 | 130617799 | SNF96.2 | FUBP3 | G | C | exonic | nonsynonym | 0.405405405 | . |
| chr10 | 46584599 | 46584599 | SNF96.2 | SYT15 | G | C | exonic | nonsynonym | 0.387755102 | rs3127785 |
| chr10 | 95968782 | 95968782 | SNF96.2 | CC2D2B | T | C | exonic | nonsynonym | 0.451612903 | . |
| chr10 | 119172416 | 119172416 | SNF96.2 | PRDX3 | C | A | exonic | nonsynonym | 1 | rs11554923 |
| chr11 | 85725357 | 85725357 | SNF96.2 | SYTL2 | G | A | exonic | nonsynonym | 1 | rs147682106 |
| chr11 | 86264258 | 86264258 | SNF96.2 | EED | A | C | exonic | nonsynonym | 1 | . |
| chr12 | 11186007 | 11186007 | SNF96.2 | TAS2R42 | G | C | exonic | nonsynonym | 0.682539682 | rs1650017 |
| chr12 | 11186175 | 11186175 | SNF96.2 | TAS2R42 | A | C | exonic | nonsynonym | 0.657894736 | rs1669413 |
| chr12 | 25108825 | 25108825 | SNF96.2 | CASC1 | T | TAAAAAAAA/ | splicing | . | 0.909090909 | . |
| chr12 | 26055402 | 26055404 | SNF96.2 | RASSF8 | TCA | T | exonic | frameshift_d | 0.337349397 | . |
| chr12 | 65326930 | 65326930 | SNF96.2 | MSRB3 | G | A | exonic | nonsynonym | 0.58 | rs776973896 |
| chr13 | 110456904 | 110456904 | SNF96.2 | COL4A2-AS2 | G | A | exonic | nonsynonym | 0.428571428 | rs370055942 |
| chr14 | 20295444 | 20295444 | SNF96.2 | TTC5 | T | C | exonic | nonsynonym | 1 | rs61741728 |
| chr14 | 21665434 | 21665434 | SNF96.2 | OR4E2 | A | G | exonic | nonsynonym | 1 | rs2874103 |
| chr14 | 21665783 | 21665783 | SNF96.2 | OR4E2 | G | A | exonic | nonsynonym | 1 | rs970382 |
| chr14 | 31302389 | 31302390 | SNF96.2 | HEATR5A | TA | T | exonic | frameshift_d | 0.392156862 | . |
| chr14 | 81324057 | 81324057 | SNF96.2 | STON2 | C | A | exonic | nonsynonym | 0.153846153 | . |
| chr14 | 92071009 | 92071009 | SNF96.2 | ATXN3 | C | CGCTGCTGCT | exonic | nonframeshi | 1 | . |
| chr14 | 100539024 | 100539024 | SNF96.2 | BEGAIN | G | A | exonic | nonsynonym | 1 | . |
| chr15 | 75639625 | 75639625 | SNF96.2 | IMP3 | G | C | exonic | nonsynonym | 1 | rs148358312 |
| chr15 | 83897858 | 83897858 | SNF96.2 | ADAMTSL3 | T | G | exonic | nonsynonym | 375 | . |

| | | | | | | | | | | |
|---|---|---|---|---|---|---|---|---|---|---|
| chr16 | 87765862 | 87765862 | SNF96.2 | KLHDC4 | T | C | exonic | nonsynonym | 0.357142857 | . |
| chr17 | 31233187 | 31233188 | SNF96.2 | NF1 | GC | G | exonic | frameshift_d | 1 | . |
| chr17 | 36451746 | 36451746 | SNF96.2 | TBC1D3D\x3l | A | G | exonic | nonsynonym | 1 | . |
| chr17 | 41190343 | 41190343 | SNF96.2 | KRTAP9-1 | T | TGCTGTGGG | exonic | nonframeshi | 1 | . |
| chr17 | 50356597 | 50356597 | SNF96.2 | XYLT2 | T | A | exonic | stopgain | 1 | . |
| chr17 | 76018931 | 76018931 | SNF96.2 | EVPL | C | T | exonic | nonsynonym | 0.133333333 | . |
| chr17 | 81647906 | 81647906 | SNF96.2 | TSPAN10 | T | TTAAC | exonic | stopgain | 1 | . |
| chr19 | 8605046 | 8605046 | SNF96.2 | ADAMTS10 | C | G | exonic | nonsynonym | 0.6 | rs7255721 |
| chr19 | 9848097 | 9848097 | SNF96.2 | PIN1 | C | A | exonic | stopgain | 0.466666666 | . |
| chr19 | 15682425 | 15682425 | SNF96.2 | CYP4F12 | C | T | exonic | nonsynonym | 0.541666666 | rs2285888 |
| chr19 | 21116968 | 21116971 | SNF96.2 | ZNF714 | TATA | T | exonic | nonframeshi | 0.5 | . |
| chr19 | 33205444 | 33205444 | SNF96.2 | LRP3 | C | T | exonic | nonsynonym | 0.571428571 | rs201590022 |
| chr19 | 35753198 | 35753198 | SNF96.2 | LIN37 | G | A | exonic | nonsynonym | 0.3125 | rs201290153 |
| chr19 | 39902274 | 39902274 | SNF96.2 | FCGBP | G | C | exonic | nonsynonym | 0.615384615 | rs139175656 |
| chr19 | 41981759 | 41981763 | SNF96.2 | ATP1A3 | ACAGC | A | exonic | frameshift_d | 0.245283018 | . |
| chr19 | 54726150 | 54726150 | SNF96.2 | KIR3DL3 | T | A | exonic | nonsynonym | 1 | rs2075731 |
| chr19 | 57477298 | 57477298 | SNF96.2 | ZNF772 | A | ACCC | exonic | nonframeshi | 1 | . |
| chr20 | 64098440 | 64098440 | SNF96.2 | OPRL1 | C | T | exonic | stopgain | 1 | rs534873633 |
| chr21 | 41168492 | 41168501 | SNF96.2 | BACE2 | TTCTTGGCCA | T | exonic | nonframeshi | 0.533333333 | . |
| chr22 | 36763949 | 36763949 | SNF96.2 | IFT27 | G | A | exonic | nonsynonym | 1 | rs143842386 |
| chr22 | 43168044 | 43168044 | SNF96.2 | TTLL12 | G | C | exonic | nonsynonym | 1 | rs147704786 |
| chrX | 318747 | 318747 | SNF96.2 | GTPBP6 | A | G | exonic | nonsynonym | 375 | . |
| chrX | 24364299 | 24364326 | SNF96.2 | SUPT20HL1 | TGCTGCTGCT | T | exonic | nonframeshi | 1 | . |
| chrX | 63678573 | 63678573 | SNF96.2 | ARHGEF9 | C | T | splicing | . | 0.142857142 | . |
| chrX | 102127322 | 102127322 | SNF96.2 | TCEAL2 | T | A | exonic | nonsynonym | 0.222222222 | . |
| chrX | 119011856 | 119011856 | SNF96.2 | LONRF3 | A | G | exonic | nonsynonym | 1 | . |
| chrX | 153182483 | 153182483 | SNF96.2 | MAGEA1 | A | G | exonic | nonsynonym | 1 | rs2008160 |

| Variant | AAChange.re | GeneDetail.r | AF | AF_popmax | prop_pathog | CLNALLELEID | CLNDN | CLNDISDB | CLNREVSTAT | CLNSIG |
|---|---|---|---|---|---|---|---|---|---|---|
| chr1:262823 | UBXN11:NM_ | . | 0.0004 | 0.0015 | NA | . | . | . | . | . |
| chr1:263450 | CRYBG2:NM_ | . | 0.0016 | 0.0039 | 1 | . | . | . | . | . |
| chr1:408620 | CITED4:NM_ | . | . | . | 0.714285714 | . | . | . | . | . |
| chr1:145873 | ANKRD35:NM | . | . | . | NA | . | . | . | . | . |
| chr1:146067 | NBPF10:NM_ | . | . | . | NA | . | . | . | . | . |
| chr1:146067 | NBPF10:NM_ | . | . | . | NA | . | . | . | . | . |
| chr1:148595 | NBPF14:NM_ | . | . | . | NA | . | . | . | . | . |
| chr1:150077 | VPS45:NM_0 | . | . | . | 0.857142857 | . | . | . | . | . |
| chr1:154010 | NUP210L:NM | . | . | . | 0.714285714 | . | . | . | . | . |
| chr1:159890 | CFAP45:NM_ | . | . | . | NA | . | . | . | . | . |
| chr1:173914 | SERPINC1:NM | . | 0.0011 | 1 | 0.714285714 | 33050 | Reduced_ant | Human_Pher | criteria_prov | Pathogenic |
| chr2:995543 | AFF3:NM_00 | . | . | . | 0.714285714 | . | . | . | . | . |
| chr2:112564485_C/G | | . | . | . | 0.857142857 | . | . | . | . | . |
| chr2:117944 | . | NM_019044: | . | . | 1 | . | . | . | . | . |
| chr2:166199 | SCN9A:NM_ | . | . | . | 1 | . | . | . | . | . |
| chr2:227008 | COL4A4:NM_ | . | . | . | 1 | . | . | . | . | . |
| chr2:236580 | ACKR3:NM_ | . | . | . | 0.857142857 | . | . | . | . | . |
| chr3:876807 | OXTR:NM_00 | . | . | . | 0.714285714 | . | . | . | . | . |
| chr3:120048 | SYN2:NM_00 | . | . | . | NA | . | . | . | . | . |
| chr3:491253 | LAMB2:NM_ | . | . | . | 0.714285714 | . | . | . | . | . |
| chr3:121576 | ARGFX:NM_ | . | . | . | NA | . | . | . | . | . |
| chr3:127575494_AGCTCCT | | . | . | . | NA | . | . | . | . | . |
| chr3:165030 | SI:NM_0010 | . | . | . | 0.857142857 | . | . | . | . | . |
| chr4:139730 | MAML3:NM_ | . | . | . | NA | . | . | . | . | . |
| chr4:184011 | STOX2:NM_0 | . | 0.0008 | 0.0014 | 0.857142857 | . | . | . | . | . |
| chr5:141184 | PCDHB16:NM | . | . | . | NA | . | . | . | . | . |
| chr6:163276 | ATXN1:NM_ | . | . | . | NA | . | . | . | . | . |
| chr6:325302 | HLA-DRB5:N | . | . | . | NA | . | . | . | . | . |
| chr6:167976 | HGC6.3:NM_ | . | 0.0027 | 0.0064 | NA | . | . | . | . | . |
| chr7:984491_ | CYP2W1:NM_ | . | . | . | 0.857142857 | . | . | . | . | . |
| chr7:459208 | IGFBP3:NM_ | . | . | . | 0.714285714 | . | . | . | . | . |
| chr7:747793 | NCF1:NM_00 | . | . | . | NA | . | . | . | . | . |
| chr7:104328 | LHFPL3:NM_ | . | . | . | 0.857142857 | . | . | . | . | . |
| chr7:123452 | IQUB:NM_00 | . | . | . | NA | . | . | . | . | . |

| | | | | | | | | | | |
|---|---|---|---|---|---|---|---|---|---|---|
| chr7:129727 | NRF1:NM_00 | . | . | . | 0.833333333 | . | . | . | . | . |
| chr8:8318865_CGGGGCG/ | | . | . | . | NA | . | . | . | . | . |
| chr8:8377352_CGCCGCT/C | | . | . | . | NA | . | . | . | . | . |
| chr8:11808709_GTCCCAC/ | | . | . | . | NA | . | . | . | . | . |
| chr8:945191 | VIRMA:NM_( | . | . | . | NA | . | . | . | . | . |
| chr8:131944802_G/A | | . | . | . | 0.857142857 | . | . | . | . | . |
| chr8:141494 | . | NM_207414: | . | . | NA | . | . | . | . | . |
| chr8:144137 | HGH1:NM_0 | . | . | . | 0.833333333 | . | . | . | . | . |
| chr8:144379 | ADCK5:NM_1 | . | . | . | NA | . | . | . | . | . |
| chr9:352957 | UNC13B:NM_ | . | 0.0046 | 0.0049 | 1 | . | . | . | . | . |
| chr9:661410 | ANKRD20A3: | . | . | . | NA | . | . | . | . | . |
| chr9:932639 | WNK2:NM_0 | . | . | . | 0.714285714 | . | . | . | . | . |
| chr9:976505 | NCBP1:NM_( | . | . | . | NA | . | . | . | . | . |
| chr9:129868 | USP20:NM_0 | . | 0.0033 | 0.0054 | 0.714285714 | . | . | . | . | . |
| chr9:129900 | FNBP1:NM_C | . | . | . | 1 | . | . | . | . | . |
| chr9:130617 | FUBP3:NM_C | . | . | . | 0.857142857 | . | . | . | . | . |
| chr10:46584 | SYT15:NM_0 | . | . | . | NA | . | . | . | . | . |
| chr10:95968 | CC2D2B:NM_ | . | . | . | 1 | . | . | . | . | . |
| chr10:11917 | PRDX3:NM_C | . | 0.0039 | 0.0059 | 0.857142857 | . | . | . | . | . |
| chr11:85725 | SYTL2:NM_2( | . | 0.0006 | 1 | NA | . | . | . | . | . |
| chr11:86264 | EED:NM_001 | . | . | . | 0.857142857 | . | . | . | . | . |
| chr12:11186 | TAS2R42:NM | . | . | . | NA | . | . | . | . | . |
| chr12:11186 | TAS2R42:NM | . | . | . | NA | . | . | . | . | . |
| chr12:25108 | . | | . | . | NA | . | . | . | . | . |
| chr12:26055 | RASSF8:NM_ | . | . | . | NA | . | . | . | . | . |
| chr12:65326 | MSRB3:NM_ | . | . | . | 1 | . | . | . | . | . |
| chr13:11045 | COL4A2-AS2: | . | 0.0019 | 3 | NA | . | . | . | . | . |
| chr14:20295 | TTC5:NM_13 | . | 0.0052 | 0.0087 | 0.714285714 | . | . | . | . | . |
| chr14:21665 | OR4E2:NM_C | . | . | . | NA | . | . | . | . | . |
| chr14:21665 | OR4E2:NM_C | . | . | . | NA | . | . | . | . | . |
| chr14:31302 | HEATR5A:NM | . | . | . | NA | . | . | . | . | . |
| chr14:81324 | STON2:NM_C | . | . | . | NA | . | . | . | . | . |
| chr14:92071009_C/CGCTG | | . | . | . | NA | . | . | . | . | . |
| chr14:10053 | BEGAIN:NM_ | . | . | . | 0.857142857 | . | . | . | . | . |
| chr15:75639 | IMP3:NM_01 | . | 0.0026 | 0.0042 | 0.833333333 | . | . | . | . | . |
| chr15:83897 | ADAMTSL3:N | . | . | . | 1 | . | . | . | . | . |

| | | | | | | | | | | |
|---|---|---|---|---|---|---|---|---|---|---|
| chr16:87765 | KLHDC4:NM_ | . | . | . | 0.857142857 | . | . | . | . | . |
| chr17:31233 | NF1:NM_000 | . | . | . | NA | . | . | . | . | . |
| chr17:36451 | TBC1D3J:NM | . | . | . | NA | . | . | . | . | . |
| chr17:41190 | KRTAP9-1:NM | . | . | . | NA | . | . | . | . | . |
| chr17:50356 | XYLT2:NM_0 | . | . | . | 1 | . | . | . | . | . |
| chr17:76018 | EVPL:NM_00 | . | . | . | 1 | . | . | . | . | . |
| chr17:81647 | TSPAN10:NM | . | . | . | NA | . | . | . | . | . |
| chr19:86050 | ADAMTS10:N | . | . | . | NA | . | . | . | . | . |
| chr19:98480 | PIN1:NM_00 | . | . | . | 1 | . | . | . | . | . |
| chr19:15682 | CYP4F12:NM | . | . | . | NA | . | . | . | . | . |
| chr19:21116 | ZNF714:NM_ | . | . | . | NA | . | . | . | . | . |
| chr19:33205 | LRP3:NM_00 | . | 0.0099 | 9 | 0.857142857 | . | . | . | . | . |
| chr19:35753 | LIN37:NM_0 | . | 0.0021 | 0.0031 | 0.857142857 | . | . | . | . | . |
| chr19:39902 | FCGBP:NM_0 | . | . | . | NA | . | . | . | . | . |
| chr19:41981 | ATP1A3:NM_ | . | . | . | NA | . | . | . | . | . |
| chr19:54726 | KIR3DL3:NM | . | 0 | . | NA | . | . | . | . | . |
| chr19:57477 | ZNF772:NM_ | . | . | . | NA | . | . | . | . | . |
| chr20:64098 | OPRL1:NM_0 | . | . | . | 1 | . | . | . | . | . |
| chr21:41168 | BACE2:NM_0 | . | . | . | NA | . | . | . | . | . |
| chr22:36763 | IFT27:NM_0 | . | 0.0010 | 0.0019 | 0.857142857 | . | . | . | . | . |
| chr22:43168 | TTLL12:NM_ | . | 0.0031 | 0.0049 | 0.714285714 | . | . | . | . | . |
| chrX:318747_ | GTPBP6:NM_ | . | . | . | NA | . | . | . | . | . |
| chrX:243642 | SUPT20HL1:N | . | . | . | NA | . | . | . | . | . |
| chrX:636785 | . | . | . | . | 1 | . | . | . | . | . |
| chrX:102127 | TCEAL2:NM_ | . | . | . | 0.714285714 | . | . | . | . | . |
| chrX:119011 | LONRF3:NM_ | . | . | . | 0.857142857 | . | . | . | . | . |
| chrX:153182 | MAGEA1:NM | . | . | . | NA | . | . | . | . | . |

| SIFT_pred | Polyphen2_H | Polyphen2_H | LRT_pred | FATHMM_pr | MutationTast | MutationAss | InterVar_automated |
|---|---|---|---|---|---|---|---|
| . | . | . | . | . | . | . | . |
| . | . | . | . | . | P | . | . |
| D | D | P | U | T | D | M | Uncertain_significance |
| . | . | . | . | . | . | . | Uncertain_significance |
| . | . | . | . | . | . | . | Uncertain_significance |
| . | . | . | . | . | . | . | Uncertain_significance |
| . | . | . | . | . | . | . | . |
| D | D | D | D | T | D | H | Likely_pathogenic |
| T | D | D | D | T | D | M | Uncertain_significance |
| . | . | . | . | . | . | . | . |
| D | D | D | N | T | A | M | Uncertain_significance |
| D | B | P | D | T | D | M | Uncertain_significance |
| D | D | D | D | T | D | M | Uncertain_significance |
| . | . | . | . | . | D | . | . |
| D | D | D | D | D | D | M | Uncertain_significance |
| D | D | D | . | D | D | H | . |
| D | D | P | D | T | D | M | Uncertain_significance |
| D | D | P | D | T | D | L | Uncertain_significance |
| . | . | . | . | . | . | . | Uncertain_significance |
| D | P | B | D | T | D | H | Uncertain_significance |
| . | . | . | . | . | . | . | . |
| . | . | . | . | . | . | . | . |
| D | D | D | D | T | D | H | Uncertain_significance |
| . | . | . | . | . | . | . | . |
| D | D | D | D | D | D | L | Uncertain_significance |
| . | . | . | . | . | . | . | Uncertain_significance |
| . | . | . | . | . | . | . | . |
| . | . | . | . | . | . | . | . |
| . | . | . | . | . | . | . | . |
| D | D | D | D | T | D | H | Uncertain_significance |
| D | D | D | N | T | D | M | Uncertain_significance |
| . | . | . | . | . | . | . | Uncertain_significance |
| D | P | P | D | T | D | M | Uncertain_significance |
| . | . | . | . | . | . | . | . |

| | | | | | | | |
|---|---|---|---|---|---|---|---|
| D | D | D | D | . | D | L | Uncertain_significance |
| . | . | . | . | . | . | . | . |
| . | . | . | . | . | . | . | . |
| . | . | . | . | . | . | . | . |
| . | . | . | . | . | . | . | . |
| D | D | D | D | T | D | M | Uncertain_significance |
| . | . | . | . | . | . | . | . |
| D | D | D | D | . | N | M | Uncertain_significance |
| . | . | . | . | . | . | . | Uncertain_significance |
| D | D | P | D | D | D | M | Uncertain_significance |
| . | . | . | . | . | . | . | . |
| T | D | D | D | T | D | M | Uncertain_significance |
| . | . | . | . | . | . | . | . |
| D | P | P | N | T | D | M | Uncertain_significance |
| . | . | . | D | . | A | . | Uncertain_significance |
| D | D | D | D | T | D | M | Uncertain_significance |
| . | . | . | . | . | . | . | Uncertain_significance |
| . | . | . | . | . | D | . | Uncertain_significance |
| D | D | D | D | T | D | H | Uncertain_significance |
| . | . | . | . | . | . | . | . |
| D | D | D | D | T | D | M | Uncertain_significance |
| . | . | . | . | . | . | . | Uncertain_significance |
| . | . | . | . | . | . | . | Uncertain_significance |
| . | . | . | . | . | . | . | . |
| . | . | . | . | . | . | . | . |
| D | D | D | D | D | D | H | Uncertain_significance |
| . | . | . | . | . | . | . | Uncertain_significance |
| D | D | P | D | T | D | L | Uncertain_significance |
| . | . | . | . | . | . | . | Uncertain_significance |
| . | . | . | . | . | . | . | Uncertain_significance |
| . | . | . | . | . | . | . | . |
| . | . | . | . | . | . | . | . |
| . | . | . | . | . | . | . | . |
| D | D | D | D | T | D | M | . |
| D | P | B | D | . | D | M | . |
| D | D | D | D | D | D | H | Uncertain_significance |

| | | | | | | | |
|---|---|---|---|---|---|---|---|
| D | D | D | D | T | D | M | Uncertain_significance |
| . | . | . | . | . | . | . | . |
| . | . | . | . | . | . | . | Uncertain_significance |
| . | . | . | . | . | . | . | . |
| . | . | . | D | . | A | . | Likely_pathogenic |
| D | D | D | D | D | D | M | Uncertain_significance |
| . | . | . | . | . | . | . | . |
| . | . | . | . | . | . | . | Uncertain_significance |
| . | . | . | D | . | A | . | Uncertain_significance |
| . | . | . | . | . | . | . | Uncertain_significance |
| . | . | . | . | . | . | . | . |
| T | D | D | D | D | D | M | Uncertain_significance |
| D | D | D | D | T | D | M | Uncertain_significance |
| . | . | . | . | . | . | . | Uncertain_significance |
| . | . | . | . | . | . | . | . |
| . | . | . | . | . | . | . | Uncertain_significance |
| . | . | . | . | . | . | . | . |
| . | . | . | D | . | D | . | Uncertain_significance |
| . | . | . | . | . | . | . | . |
| D | D | D | D | T | D | M | Uncertain_significance |
| T | D | P | D | T | D | M | . |
| . | . | . | . | . | . | . | . |
| . | . | . | . | . | . | . | . |
| . | . | . | . | . | D | . | . |
| D | P | P | D | T | N | M | Uncertain_significance |
| D | D | D | D | T | D | M | Uncertain_significance |
| . | . | . | . | . | . | . | Uncertain_significance |

#NMS-2 cell line variants with a potential impact on protein function

| chr | start | end | Sample | Gene.refGen | REF | ALT | Func.refGene | ExonicFunc.r | VAF | avsnp150 |
|---|---|---|---|---|---|---|---|---|---|---|
| chr1 | 1437382 | 1437382 | NMS.2 | VWA1 | G | A | exonic | nonsynonym | 0.392857142 | . |
| chr1 | 43606433 | 43606433 | NMS.2 | PTPRF | C | G | exonic | nonsynonym | 1 | . |
| chr1 | 46533277 | 46533277 | NMS.2 | TMEM275 | C | G | exonic | nonsynonym | 0.3 | . |
| chr1 | 121365328 | 121365328 | NMS.2 | SRGAP2C | G | C | exonic | nonsynonym | 1 | . |
| chr1 | 149390802 | 149390802 | NMS.2 | NOTCH2NLC | A | AGGCGGCGG | exonic | nonframeshi | 1 | . |
| chr1 | 155208991 | 155208991 | NMS.2 | MTX1 | T | A | exonic | nonsynonym | 0.533333333 | rs760077 |
| chr1 | 180154945 | 180154945 | NMS.2 | QSOX1 | C | CGCT | exonic | nonframeshi | 0.294117647 | . |
| chr1 | 241958819 | 241958819 | NMS.2 | BECN2 | G | A | exonic | nonsynonym | 0.166666666 | . |
| chr2 | 1137652 | 1137652 | NMS.2 | SNTG2 | A | T | exonic | nonsynonym | 0.529411764 | rs748927979 |
| chr2 | 43889784 | 43889784 | NMS.2 | LRPPRC | C | T | exonic | nonsynonym | 0.441176470 | rs147302249 |
| chr2 | 113192531 | 113192531 | NMS.2 | PSD4 | C | T | exonic | nonsynonym | 0.25 | rs117870995 |
| chr2 | 157321335 | 157321335 | NMS.2 | ERMN | C | T | exonic | nonsynonym | 0.340909090 | rs143517901 |
| chr2 | 200892074 | 200892074 | NMS.2 | NIF3L1 | C | T | exonic | nonsynonym | 0.707317073 | rs144466763 |
| chr2 | 240021253 | 240021253 | NMS.2 | NDUFA10 | A | G | exonic | nonsynonym | 0.3 | rs140776586 |
| chr3 | 6861728 | 6861728 | NMS.2 | GRM7 | T | TA | exonic | stopgain | 0.2 | . |
| chr3 | 9827207 | 9827207 | NMS.2 | ARPC4-TTLL3 | C | T | exonic | nonsynonym | 0.756097560 | . |
| chr3 | 51980458 | 51980458 | NMS.2 | ABHD14A | C | T | exonic | nonsynonym | 0.289473684 | rs145685600 |
| chr3 | 57368154 | 57368154 | NMS.2 | DNAH12 | T | G | exonic | nonsynonym | 0.7 | . |
| chr3 | 111133860 | 111133860 | NMS.2 | NECTIN3 | G | C | exonic | nonsynonym | 0.309523809 | rs15611 |
| chr3 | 113657263 | 113657272 | NMS.2 | USF3 | TTGCTGCTGC | TTGCTGC | exonic | nonframeshi | 1 | . |
| chr3 | 167315801 | 167315804 | NMS.2 | ZBBX | TCTG | T | exonic | nonframeshi | 0.692307692 | . |
| chr4 | 130852 | 130852 | NMS.2 | ZNF718 | C | T | exonic | nonsynonym | 1 | . |
| chr4 | 131416 | 131416 | NMS.2 | ZNF718 | G | C | exonic | nonsynonym | 1 | . |
| chr4 | 8606806 | 8606806 | NMS.2 | CPZ | C | T | exonic | nonsynonym | 0.25 | . |
| chr4 | 37845708 | 37845708 | NMS.2 | PGM2 | G | A | exonic | nonsynonym | 1 | . |
| chr4 | 52003267 | 52003271 | NMS.2 | LRRC66 | ATATC | A | exonic | frameshift_d | 0.707317073 | . |
| chr4 | 145096107 | 145096107 | NMS.2 | ANAPC10 | A | G | exonic | nonsynonym | 0.615384615 | . |
| chr5 | 139805 | 139805 | NMS.2 | PLEKHG4B | G | A | exonic | nonsynonym | 0.777777777 | rs964226641 |
| chr5 | 139394120 | 139394120 | NMS.2 | PROB1 | C | T | exonic | nonsynonym | 0.340909090 | . |
| chr5 | 168423571 | 168423571 | NMS.2 | WWC1 | G | A | exonic | nonsynonym | 475 | . |
| chr6 | 7229741 | 7229741 | NMS.2 | RREB1 | A | C | exonic | nonsynonym | 0.55 | rs116672033 |
| chr6 | 16327660 | 16327660 | NMS.2 | ATXN1 | C | CTGCTGA | exonic | nonframeshi | 0.357142857 | . |
| chr6 | 129600780 | 129600780 | NMS.2 | ARHGAP18 | T | TA | exonic | frameshift_ir | 0.428571428 | . |
| chr7 | 2513247 | 2513251 | NMS.2 | LFNG | AGATG | AGATGGATG | exonic | frameshift_ir | 1 | . |

| | | | | | | | | | | |
|---|---|---|---|---|---|---|---|---|---|---|
| chr7 | 26184853 | 26184853 | NMS.2 | NFE2L3 | C | CATAA | exonic | frameshift_ir | 0.243902439 | . |
| chr7 | 34051944 | 34051944 | NMS.2 | BMPER | G | A | exonic | nonsynonym | 0.470588235 | rs267601491 |
| chr7 | 56106409 | 56106409 | NMS.2 | CHCHD2 | G | A | exonic | nonsynonym | 0.461538461 | rs142444896 |
| chr7 | 121331831 | 121331831 | NMS.2 | WNT16 | G | A | exonic | nonsynonym | 0.604651162 | . |
| chr7 | 127106631 | 127106631 | NMS.2 | GRM8 | G | A | exonic | nonsynonym | 0.153846153 | . |
| chr7 | 128673398 | 128673398 | NMS.2 | FAM71F2 | G | T | splicing | . | 0.928571428 | . |
| chr8 | 41723615 | 41723615 | NMS.2 | ANK1 | G | A | exonic | nonsynonym | 0.304347826 | . |
| chr8 | 109087637 | 109087637 | NMS.2 | TRHR | G | C | exonic | nonsynonym | 0.324675324 | . |
| chr8 | 143865354 | 143865360 | NMS.2 | EPPK1 | GGGCGGC | GGGC | exonic | nonframeshi | 1 | . |
| chr8 | 143865354 | 143865360 | NMS.2 | EPPK1 | GGGCGGC | G | exonic | nonframeshi | 1 | . |
| chr8 | 144138429 | 144138429 | NMS.2 | HGH1 | G | T | splicing | . | 0.422222222 | . |
| chr9 | 41991721 | 41991721 | NMS.2 | CNTNAP3B | G | A | exonic | nonsynonym | 1 | rs3739623 |
| chr9 | 110375465 | 110375465 | NMS.2 | SVEP1 | T | TA | splicing | . | 0.560975609 | . |
| chr10 | 46584599 | 46584599 | NMS.2 | SYT15 | G | C | exonic | nonsynonym | 0.352941176 | rs3127785 |
| chr10 | 99880233 | 99880233 | NMS.2 | DNMBP | G | A | exonic | nonsynonym | 0.349206349 | rs375209058 |
| chr10 | 99884127 | 99884127 | NMS.2 | DNMBP | C | T | exonic | nonsynonym | 0.388888888 | . |
| chr11 | 4091404 | 4091404 | NMS.2 | STIM1 | G | T | exonic | nonsynonym | 1 | rs200525396 |
| chr11 | 14259653 | 14259653 | NMS.2 | SPON1 | T | G | exonic | nonsynonym | 1 | . |
| chr11 | 46542834 | 46542836 | NMS.2 | AMBRA1 | AAG | A | exonic | frameshift_d | 1 | . |
| chr11 | 70326625 | 70326626 | NMS.2 | PPFIA1 | AG | A | exonic | frameshift_d | 1 | . |
| chr11 | 85664560 | 85664560 | NMS.2 | CREBZF | C | T | exonic | nonsynonym | 0.238095238 | . |
| chr11 | 133925866 | 133925866 | NMS.2 | IGSF9B | G | A | exonic | nonsynonym | 1 | . |
| chr12 | 6621133 | 6621134 | NMS.2 | LPAR5 | AG | A | exonic | frameshift_d | 0.615384615 | . |
| chr12 | 6667903 | 6667903 | NMS.2 | ZNF384 | T | TTGCTGC | exonic | nonframeshi | 0.4 | . |
| chr12 | 55837233 | 55837233 | NMS.2 | MMP19 | T | C | exonic | nonsynonym | 0.552631578 | . |
| chr12 | 76031021 | 76031021 | NMS.2 | PHLDA1 | C | T | exonic | nonsynonym | 0.173913043 | . |
| chr12 | 121626874 | 121626879 | NMS.2 | ORAI1 | CCGCCA | C | exonic | frameshift_d | 1 | . |
| chr12 | 124402512 | 124402521 | NMS.2 | NCOR2 | GGCTGCTGCT | GGCTGCTGCT | exonic | nonframeshi | 0.923076923 | . |
| chr12 | 132808144 | 132808144 | NMS.2 | GOLGA3 | C | T | exonic | nonsynonym | 0.592592592 | . |
| chr13 | 25097025 | 25097025 | NMS.2 | PABPC3 | T | A | exonic | nonsynonym | 0.595744680 | rs104430191 |
| chr14 | 19434064 | 19434064 | NMS.2 | POTEG | T | C | exonic | nonsynonym | 0.25 | rs57000026 |
| chr14 | 77026607 | 77026607 | NMS.2 | IRF2BPL | G | A | exonic | nonsynonym | 0.153846153 | . |
| chr14 | 88426650 | 88426650 | NMS.2 | SPATA7 | G | GA | exonic | frameshift_ir | 0.696969696 | . |
| chr14 | 92071009 | 92071009 | NMS.2 | ATXN3 | C | CGCTGCTGCT | exonic | nonframeshi | 1 | . |
| chr14 | 104943593 | 104943593 | NMS.2 | AHNAK2 | T | C | exonic | nonsynonym | 0.333333333 | . |
| chr15 | 51576183 | 51576183 | NMS.2 | DMXL2 | T | TAAAAAAAA | splicing | . | 0.529411764 | . |

| | | | | | | | | | | |
|---|---|---|---|---|---|---|---|---|---|---|
| chr15 | 82345046 | 82345046 | NMS.2 | GOLGA6L10 | G | A | exonic | nonsynonym | 375 | rs866101815 |
| chr15 | 85244770 | 85244770 | NMS.2 | GOLGA6L3 | G | T | exonic | nonsynonym | 0.5 | . |
| chr16 | 30654688 | 30654688 | NMS.2 | PRR14 | C | T | exonic | nonsynonym | 0.315789473 | rs746088400 |
| chr16 | 56884111 | 56884111 | NMS.2 | SLC12A3 | G | A | exonic | nonsynonym | 1 | rs139329616 |
| chr16 | 84657228 | 84657228 | NMS.2 | KLHL36 | C | T | exonic | stopgain | 0.166666666 | . |
| chr17 | 4553812 | 4553812 | NMS.2 | MYBBP1A | C | A | exonic | stopgain | 0.346153846 | . |
| chr17 | 7916002 | 7916002 | NMS.2 | RNF227 | G | C | exonic | nonsynonym | 0.4 | . |
| chr17 | 31340646 | 31340646 | NMS.2 | NF1 | G | T | splicing | . | 1 | . |
| chr17 | 38724731 | 38724731 | NMS.2 | MLLT6 | G | T | exonic | stopgain | 0.40625 | . |
| chr17 | 38951650 | 38951650 | NMS.2 | FBXO47 | C | T | exonic | nonsynonym | 725 | rs78686509 |
| chr17 | 41528000 | 41528000 | NMS.2 | KRT19 | G | C | exonic | nonsynonym | 0.857142857 | . |
| chr17 | 63943014 | 63943014 | NMS.2 | SCN4A | A | G | exonic | nonsynonym | 0.661290322 | . |
| chr17 | 69193139 | 69193139 | NMS.2 | ABCA10 | T | C | exonic | nonsynonym | 0.285714285 | . |
| chr17 | 82084839 | 82084839 | NMS.2 | FASN | G | C | exonic | nonsynonym | 1 | . |
| chr18 | 12452388 | 12452388 | NMS.2 | SPIRE1 | G | A | exonic | nonsynonym | 1 | rs568852514 |
| chr18 | 46253735 | 46253738 | NMS.2 | C18orf25 | TCTG | T | exonic | nonframeshi | 1 | . |
| chr18 | 63720882 | 63720882 | NMS.2 | SERPINB11 | G | C | exonic | nonsynonym | 0.285714285 | . |
| chr19 | 1221320 | 1221320 | NMS.2 | STK11 | C | T | exonic | nonsynonym | 1 | rs121913322 |
| chr19 | 8605046 | 8605046 | NMS.2 | ADAMTS10 | C | G | exonic | nonsynonym | 0.36 | rs7255721 |
| chr19 | 41717688 | 41717688 | NMS.2 | CEACAM5 | G | A | exonic | nonsynonym | 1 | rs7249230 |
| chr19 | 41761981 | 41761981 | NMS.2 | CEACAM6 | T | G | exonic | nonsynonym | 1 | rs11548735 |
| chr19 | 43185001 | 43185001 | NMS.2 | PSG5 | G | A | exonic | stopgain | 1 | rs760079282 |
| chr19 | 44430314 | 44430314 | NMS.2 | ZNF229 | A | G | exonic | nonsynonym | 1 | rs2571174 |
| chr19 | 54096945 | 54096945 | NMS.2 | OSCAR | C | A | exonic | nonsynonym | 1 | rs1657535 |
| chr19 | 54726150 | 54726150 | NMS.2 | KIR3DL3 | T | A | exonic | nonsynonym | 0.934782608 | rs2075731 |
| chr19 | 54832784 | 54832784 | NMS.2 | KIR2DS4 | T | G | exonic | nonsynonym | 1 | rs1130476 |
| chr19 | 54835079 | 54835079 | NMS.2 | KIR2DS4 | G | C | exonic | nonsynonym | 1 | rs112522228 |
| chr19 | 54835110 | 54835110 | NMS.2 | KIR2DS4 | G | T | exonic | nonsynonym | 1 | rs1130478 |
| chr19 | 54837865 | 54837865 | NMS.2 | KIR2DS4 | G | C | exonic | nonsynonym | 1 | rs1130494 |
| chr19 | 54837873 | 54837873 | NMS.2 | KIR2DS4 | T | C | exonic | nonsynonym | 1 | rs201518955 |
| chr19 | 55490895 | 55490895 | NMS.2 | SSC5D | G | A | exonic | nonsynonym | 0.40625 | . |
| chr20 | 2617169 | 2617169 | NMS.2 | TMC2 | G | A | exonic | nonsynonym | 1 | rs145587686 |
| chr20 | 14327280 | 14327280 | NMS.2 | FLRT3 | G | T | exonic | nonsynonym | 1 | . |
| chr20 | 63945900 | 63945900 | NMS.2 | UCKL1 | C | T | exonic | nonsynonym | 0.7 | rs201285389 |
| chr21 | 44600952 | 44600952 | NMS.2 | KRTAP10-7 | G | A | exonic | nonsynonym | 0.476190476 | rs944419 |
| chr21 | 44601264 | 44601264 | NMS.2 | KRTAP10-7 | A | C | exonic | nonsynonym | 0.558823529 | rs363877 |

| | | | | | | | | | | |
|---|---|---|---|---|---|---|---|---|---|---|
| chr21 | 44601475 | 44601475 | NMS.2 | KRTAP10-7 | C | G | exonic | nonsynonym | 0.68 | rs446817 |
| chr21 | 44601579 | 44601579 | NMS.2 | KRTAP10-7 | G | A | exonic | nonsynonym | 0.5 | rs369720 |
| chr22 | 22488632 | 22488632 | NMS.2 | ZNF280B | G | T | exonic | nonsynonym | 1 | rs2236729 |
| chr22 | 22515221 | 22515221 | NMS.2 | ZNF280A | G | T | exonic | nonsynonym | 1 | rs361580 |
| chr22 | 22515224 | 22515224 | NMS.2 | ZNF280A | C | T | exonic | nonsynonym | 1 | rs362011 |
| chr22 | 22550942 | 22550942 | NMS.2 | PRAME | A | G | exonic | nonsynonym | 0.387096774 | rs769323726 |
| chr22 | 31806127 | 31806127 | NMS.2 | DEPDC5 | A | G | exonic | nonsynonym | 0.363636363 | . |
| chrX | 2955503 | 2955503 | NMS.2 | ARSE | C | T | exonic | nonsynonym | 1 | rs150756612 |
| chrX | 36159479 | 36159479 | NMS.2 | CFAP47 | A | T | exonic | nonsynonym | 0.217391304 | . |
| chrX | 141002745 | 141002754 | NMS.2 | SPANXB1 | TCCAACGAG( | T | exonic | nonframeshi1 | 0.783783783 | . |
| chrX | 154653251 | 154653251 | NMS.2 | CTAG2 | C | G | exonic | nonsynonym | 1 | rs17328091 |

| Variant | AAChange.re | GeneDetail.r | AF | AF_popmax | prop_pathog | CLNALLELEID | CLNDN | CLNDISDB | CLNREVSTAT | CLNSIG |
|---|---|---|---|---|---|---|---|---|---|---|
| chr1:143738 | VWA1:NM_0 | . | . | . | 0.833333333 | . | . | . | . | . |
| chr1:436064 | PTPRF:NM_0 | . | . | . | 0.714285714 | . | . | . | . | . |
| chr1:465332 | TMEM275:N | . | . | . | NA | . | . | . | . | . |
| chr1:121365 | SRGAP2C:NN | . | . | . | NA | . | . | . | . | . |
| chr1:149390 | NOTCH2NLC: | . | . | . | NA | . | . | . | . | . |
| chr1:155208 | MTX1:NM_0 | . | . | . | NA | . | . | . | . | . |
| chr1:180154 | QSOX1:NM_ | . | 0.0014 | 0.0024 | NA | . | . | . | . | . |
| chr1:241958 | BECN2:NM_ | . | . | . | NA | . | . | . | . | . |
| chr2:113765 | SNTG2:NM_ | . | . | . | 0.857142857 | . | . | . | . | . |
| chr2:438897 | LRPPRC:NM_ | . | 0.0004 | 0.0083 | 0.857142857 | . | . | . | . | . |
| chr2:113192 | PSD4:NM_01 | . | 0.0001 | 0.0019 | 0.857142857 | . | . | . | . | . |
| chr2:157321 | ERMN:NM_0 | . | 0.0008 | 0.0013 | 1 | . | . | . | . | . |
| chr2:200892074_C/T | | . | 0.0006 | 0.0051 | 0.857142857 | . | . | . | . | . |
| chr2:240021 | NDUFA10:NN | . | 0.0003 | 0.0051 | 1 | 210803 | not_specifiec | MedGen:CN1 | criteria_prov | Uncertain_si |
| chr3:686172 | GRM7:NM_0 | . | . | . | NA | . | . | . | . | . |
| chr3:982720 | TTLL3:NM_0 | . | . | . | 0.714285714 | . | . | . | . | . |
| chr3:519804 | ABHD14A:NN | . | . | . | 0.666666666 | . | . | . | . | . |
| chr3:573681 | DNAH12:NM | . | . | . | NA | . | . | . | . | . |
| chr3:111133 | NECTIN3:NM | . | . | . | 0.857142857 | . | . | . | . | . |
| chr3:113657 | USF3:NM_00 | . | . | . | NA | . | . | . | . | . |
| chr3:167315 | ZBBX:NM_00 | . | . | . | NA | . | . | . | . | . |
| chr4:130852_ | ZNF718:NM_ | . | . | . | NA | . | . | . | . | . |
| chr4:131416_ | ZNF718:NM_ | . | . | . | NA | . | . | . | . | . |
| chr4:860680 | CPZ:NM_003 | . | . | . | 0.714285714 | . | . | . | . | . |
| chr4:378457 | PGM2:NM_0 | . | . | . | 0.857142857 | . | . | . | . | . |
| chr4:520032 | LRRC66:NM_ | . | . | . | NA | . | . | . | . | . |
| chr4:145096 | ANAPC10:NN | . | . | . | NA | . | . | . | . | . |
| chr5:139805_ | PLEKHG4B:N | . | 6.406e-05 | 0.0013 | NA | . | . | . | . | . |
| chr5:139394 | PROB1:NM_ | . | . | . | 0.8 | . | . | . | . | . |
| chr5:168423 | WWC1:NM_ | . | . | . | 0.857142857 | . | . | . | . | . |
| chr6:722974 | RREB1:NM_0 | . | 0.0005 | 9 | 0.666666666 | . | . | . | . | . |
| chr6:163276 | ATXN1:NM_ | . | 0.0001 | 0.0028 | NA | . | . | . | . | . |
| chr6:129600 | ARHGAP18:N | . | . | . | NA | . | . | . | . | . |
| chr7:251324 | LFNG:NM_00 | . | . | . | NA | . | . | . | . | . |

| | | | | | | | | | | |
|---|---|---|---|---|---|---|---|---|---|---|
| chr7:261848! | NFE2L3:NM_ | . | . | . | NA | . | . | . | . | . |
| chr7:340519· | BMPER:NM_( | . | . | . | 0.714285714 | . | . | . | . | . |
| chr7:561064( | CHCHD2:NM_ | . | 0.0132 | 0.0064 | 0.857142857 | . | . | . | . | . |
| chr7:121331! | WNT16:NM_ | . | . | . | 1 | . | . | . | . | . |
| chr7:127106631_G/A | | . | . | . | 0.714285714 | . | . | . | . | . |
| chr7:128673: | . | NM_0012902 | . | . | NA | . | . | . | . | . |
| chr8:417236: | ANK1:NM_0( | . | . | . | 0.857142857 | . | . | . | . | . |
| chr8:109087( | TRHR:NM_0( | . | . | . | 0.833333333 | . | . | . | . | . |
| chr8:143865: | EPPK1:NM_0 | . | . | . | NA | . | . | . | . | . |
| chr8:143865: | EPPK1:NM_0 | . | . | . | NA | . | . | . | . | . |
| chr8:144138· | . | NM_016458: | . | . | 1 | . | . | . | . | . |
| chr9:419917: | CNTNAP3B:N | . | . | . | NA | . | . | . | . | . |
| chr9:110375· | . | NM_153366: | . | . | NA | . | . | . | . | . |
| chr10:46584! | SYT15:NM_0: | . | . | . | NA | . | . | . | . | . |
| chr10:99880: | DNMBP:NM_ | . | . | . | 0.857142857 | . | . | . | . | . |
| chr10:99884: | DNMBP:NM_ | . | . | . | 0.857142857 | . | . | . | . | . |
| chr11:40914( | STIM1:NM_0 | . | . | . | 0.714285714 | . | . | . | . | . |
| chr11:14259( | SPON1:NM_( | . | . | . | 0.75 | . | . | . | . | . |
| chr11:46542834_AAG/A | | . | . | . | NA | . | . | . | . | . |
| chr11:70326( | PPFIA1:NM_( | . | . | . | NA | . | . | . | . | . |
| chr11:85664! | CREBZF:NM_ | . | . | . | 0.666666666 | . | . | . | . | . |
| chr11:13392! | IGSF9B:NM_( | . | . | . | 0.714285714 | . | . | . | . | . |
| chr12:66211: | LPAR5:NM_0 | . | . | . | NA | . | . | . | . | . |
| chr12:66679( | ZNF384:NM_ | . | 0.0016 | 0.0018 | NA | . | . | . | . | . |
| chr12:55837: | MMP19:NM_ | . | . | . | 0.857142857 | . | . | . | . | . |
| chr12:76031( | PHLDA1:NM_ | . | . | . | 0.857142857 | . | . | . | . | . |
| chr12:12162( | ORAI1:NM_0 | . | . | . | NA | . | . | . | . | . |
| chr12:12440: | NCOR2:NM_( | . | . | . | NA | . | . | . | . | . |
| chr12:13280! | GOLGA3:NM_ | . | . | . | 0.714285714 | . | . | . | . | . |
| chr13:25097( | PABPC3:NM_ | . | . | . | 0.714285714 | . | . | . | . | . |
| chr14:19434( | POTEG:NM_( | . | . | . | NA | . | . | . | . | . |
| chr14:77026( | IRF2BPL:NM_ | . | . | . | 0.714285714 | . | . | . | . | . |
| chr14:88426( | SPATA7:NM_ | . | . | . | NA | . | . | . | . | . |
| chr14:92071009_C/CGCTG | | . | . | . | NA | . | . | . | . | . |
| chr14:10494: | AHNAK2:NM_ | . | . | . | 0.666666666 | . | . | . | . | . |
| chr15:51576: | . | NM_015263: | . | . | NA | . | . | . | . | . |

| | | | | | | | | | | |
|---|---|---|---|---|---|---|---|---|---|---|
| chr15:82345 | GOLGA6L10:I | . | 0.0008 | 0.0031 | NA | . | . | . | . | . |
| chr15:85244 | GOLGA6L3:N | . | . | . | NA | . | . | . | . | . |
| chr16:30654 | PRR14:NM_0 | . | . | . | 0.714285714 | . | . | . | . | . |
| chr16:56884 | SLC12A3:NM | . | 0.0003 | 0.0051 | 1 | 227377 | Familial_hyp | MedGen:C02 | criteria_prov | Uncertain_sig |
| chr16:84657 | KLHL36:NM_ | . | . | . | 0.5 | . | . | . | . | . |
| chr17:45538 | MYBBP1A:NM | . | . | . | 1 | . | . | . | . | . |
| chr17:79160 | RNF227:NM_ | . | . | . | NA | . | . | . | . | . |
| chr17:31340 | . | NM_000267: | . | . | 1 | . | . | . | . | . |
| chr17:38724 | MLLT6:NM_C | . | . | . | 0.5 | . | . | . | . | . |
| chr17:38951 | FBXO47:NM_ | . | . | . | 0.714285714 | . | . | . | . | . |
| chr17:41528 | KRT19:NM_0 | . | . | . | 1 | . | . | . | . | . |
| chr17:63943 | SCN4A:NM_C | . | . | . | 1 | . | . | . | . | . |
| chr17:69193 | ABCA10:NM_ | . | . | . | 0.666666666 | . | . | . | . | . |
| chr17:82084 | FASN:NM_00 | . | . | . | 0.714285714 | . | . | . | . | . |
| chr18:12452 | SPIRE1:NM_C | . | 0.0001 | 0.0026 | 0.857142857 | . | . | . | . | . |
| chr18:46253 | C18orf25:NM | . | . | . | NA | . | . | . | . | . |
| chr18:63720 | SERPINB11:N | . | . | . | 1 | . | . | . | . | . |
| chr19:12213 | STK11:NM_0 | . | 6.376e-05 | 0.0012 | 0.285714285 | 151829 | Neoplasm\|P | Human_Pher | criteria_prov | Conflicting_ir |
| chr19:86050 | ADAMTS10:N | . | . | . | NA | . | . | . | . | . |
| chr19:41717 | CEACAM5:NM | . | . | . | NA | . | . | . | . | . |
| chr19:41761 | CEACAM6:NM | . | . | . | NA | . | . | . | . | . |
| chr19:43185 | PSG5:NM_00 | . | . | . | 1 | . | . | . | . | . |
| chr19:44430 | ZNF229:NM_ | . | . | . | NA | . | . | . | . | . |
| chr19:54096 | OSCAR:NM_C | . | . | . | NA | . | . | . | . | . |
| chr19:54726 | KIR3DL3:NM_ | . | 0 | . | NA | . | . | . | . | . |
| chr19:54832 | KIR2DS4:NM_ | . | . | . | NA | . | . | . | . | . |
| chr19:54835 | KIR2DS4:NM_ | . | . | . | NA | . | . | . | . | . |
| chr19:54835 | KIR2DS4:NM_ | . | . | . | NA | . | . | . | . | . |
| chr19:54837 | KIR2DS4:NM_ | . | . | . | NA | . | . | . | . | . |
| chr19:54837 | KIR2DS4:NM_ | . | . | . | NA | . | . | . | . | . |
| chr19:55490 | SSC5D:NM_0 | . | . | . | 0.714285714 | . | . | . | . | . |
| chr20:26171 | TMC2:NM_0 | . | 0.0001 | 0.0026 | 0.714285714 | . | . | . | . | . |
| chr20:14327 | FLRT3:NM_0 | . | . | . | 0.857142857 | . | . | . | . | . |
| chr20:63945900_C/T | | . | 9.565e-05 | 0.0013 | 1 | . | . | . | . | . |
| chr21:44600 | KRTAP10-7:N | . | . | . | NA | . | . | . | . | . |
| chr21:44601 | KRTAP10-7:N | . | . | . | NA | . | . | . | . | . |

| | | | | | | | | | | |
|---|---|---|---|---|---|---|---|---|---|---|
| chr21:44601 | KRTAP10-7:N | . | . | . | NA | . | . | . | . | . |
| chr21:44601 | KRTAP10-7:N | . | . | . | NA | . | . | . | . | . |
| chr22:22488 | ZNF280B:NM | . | . | . | NA | . | . | . | . | . |
| chr22:22515 | ZNF280A:NM | . | . | . | NA | . | . | . | . | . |
| chr22:22515 | ZNF280A:NM | . | . | . | NA | . | . | . | . | . |
| chr22:22550942_A/G | | . | . | . | 0.714285714 | . | . | . | . | . |
| chr22:31806127_A/G | | . | . | . | 0.857142857 | . | . | . | . | . |
| chrX:295550 | ARSE:NM_00 | . | 0.0001 | 2 | 0.857142857 | . | . | . | . | . |
| chrX:361594 | CFAP47:NM_ | . | . | . | NA | . | . | . | . | . |
| chrX:141002 | SPANXB1:NM | . | . | . | NA | . | . | . | . | . |
| chrX:154653 | CTAG2:NM_ | . | . | . | NA | . | . | . | . | . |

| SIFT_pred | Polyphen2_H | Polyphen2_H | LRT_pred | FATHMM_pr | MutationTast | MutationAss | InterVar_automated |
|---|---|---|---|---|---|---|---|
| D | D | P | U | D | D | . | Uncertain_significance |
| D | D | D | N | T | D | M | Uncertain_significance |
| . | . | . | . | . | . | . | . |
| . | . | . | . | . | . | . | Uncertain_significance |
| . | . | . | . | . | . | . | . |
| . | . | . | . | . | . | . | Uncertain_significance |
| . | . | . | . | . | . | . | . |
| . | . | . | . | . | . | . | Uncertain_significance |
| D | D | D | D | T | D | H | Uncertain_significance |
| D | D | D | D | T | D | M | Uncertain_significance |
| D | D | D | D | T | D | M | Uncertain_significance |
| D | D | D | D | D | D | M | . |
| D | D | P | D | T | D | H | Uncertain_significance |
| D | D | D | D | D | D | M | . |
| . | . | . | . | . | . | . | . |
| D | D | D | U | T | D | M | Uncertain_significance |
| D | D | P | . | T | N | M | Uncertain_significance |
| . | . | . | . | . | . | . | Uncertain_significance |
| D | D | D | D | T | D | M | Uncertain_significance |
| . | . | . | . | . | . | . | . |
| . | . | . | . | . | . | . | . |
| . | . | . | . | . | . | . | Uncertain_significance |
| . | . | . | . | . | . | . | Uncertain_significance |
| D | D | D | U | T | D | H | Uncertain_significance |
| D | D | D | D | T | D | M | Uncertain_significance |
| . | . | . | . | . | . | . | . |
| . | . | . | . | . | . | . | Uncertain_significance |
| . | . | . | . | . | . | . | Uncertain_significance |
| D | D | D | . | . | D | L | . |
| D | D | D | D | T | D | M | Uncertain_significance |
| D | P | P | . | T | D | L | Uncertain_significance |
| . | . | . | . | . | . | . | . |
| . | . | . | . | . | . | . | . |
| . | . | . | . | . | . | . | . |

| | | | | | | | |
|---|---|---|---|---|---|---|---|
| . | . | . | . | . | . | . | . |
| D | D | D | N | T | D | M | Uncertain_significance |
| D | D | D | D | T | D | M | Uncertain_significance |
| D | D | D | D | D | D | H | Uncertain_significance |
| T | D | D | D | D | D | N | . |
| . | . | . | . | . | . | . | . |
| D | D | D | D | T | D | M | Uncertain_significance |
| D | D | D | D | T | D | . | Uncertain_significance |
| . | . | . | . | . | . | . | . |
| . | . | . | . | . | . | . | . |
| . | . | . | . | . | D | . | . |
| . | . | . | . | . | . | . | . |
| . | . | . | . | . | . | . | . |
| . | . | . | . | . | . | . | Uncertain_significance |
| D | D | D | D | T | D | M | . |
| D | D | D | D | T | D | M | Uncertain_significance |
| D | P | B | D | T | D | M | Uncertain_significance |
| . | D | P | D | T | . | . | Uncertain_significance |
| . | . | . | . | . | . | . | . |
| . | . | . | . | . | . | . | . |
| D | D | D | N | . | D | N | Uncertain_significance |
| D | D | P | D | T | D | L | Uncertain_significance |
| . | . | . | . | . | . | . | . |
| . | . | . | . | . | . | . | . |
| D | D | D | D | T | D | M | Uncertain_significance |
| D | D | D | D | T | D | M | Uncertain_significance |
| . | . | . | . | . | . | . | . |
| . | . | . | . | . | . | . | . |
| D | D | P | D | T | D | L | Uncertain_significance |
| D | D | D | U | T | D | H | Uncertain_significance |
| . | . | . | . | . | . | . | Uncertain_significance |
| D | D | D | U | T | D | M | Uncertain_significance |
| . | . | . | . | . | . | . | . |
| . | . | . | . | . | . | . | . |
| D | D | P | . | T | N | M | Uncertain_significance |
| . | . | . | . | . | . | . | . |

| | | | | | | | |
|---|---|---|---|---|---|---|---|
| . | . | . | . | . | . | . | Uncertain_significance |
| . | . | . | . | . | . | . | Uncertain_significance |
| D | D | D | D | T | D | L | Uncertain_significance |
| D | D | D | D | D | D | M | Uncertain_significance |
| . | . | . | N | . | A | . | Pathogenic |
| . | . | . | D | . | A | . | Uncertain_significance |
| . | . | . | . | . | . | . | . |
| . | . | . | . | . | D | . | . |
| . | . | . | N | . | A | . | Pathogenic |
| T | D | D | D | T | D | M | Uncertain_significance |
| D | D | D | D | D | D | H | Uncertain_significance |
| D | D | D | D | D | D | M | Uncertain_significance |
| D | P | P | . | T | D | L | Uncertain_significance |
| D | D | D | N | T | D | M | . |
| D | D | D | D | T | D | M | Uncertain_significance |
| . | . | . | . | . | . | . | . |
| . | D | D | . | D | D | M | Uncertain_significance |
| T | B | B | D | T | D | L | Uncertain_significance |
| . | . | . | . | . | . | . | Uncertain_significance |
| . | . | . | . | . | . | . | Uncertain_significance |
| . | . | . | . | . | . | . | Uncertain_significance |
| . | . | . | . | . | A | . | . |
| . | . | . | . | . | . | . | Uncertain_significance |
| . | . | . | . | . | . | . | . |
| . | . | . | . | . | . | . | Uncertain_significance |
| . | . | . | . | . | . | . | Uncertain_significance |
| . | . | . | . | . | . | . | Uncertain_significance |
| . | . | . | . | . | . | . | Uncertain_significance |
| . | . | . | . | . | . | . | Uncertain_significance |
| . | . | . | . | . | . | . | Uncertain_significance |
| D | D | D | U | T | D | M | Uncertain_significance |
| D | D | D | D | T | D | L | Uncertain_significance |
| D | D | D | D | T | D | M | Uncertain_significance |
| D | D | P | D | . | D | M | Uncertain_significance |
| . | . | . | . | . | . | . | Uncertain_significance |
| . | . | . | . | . | . | . | Uncertain_significance |

| | | | | | | | |
|---|---|---|---|---|---|---|---|
| . | . | . | . | . | . | . | Uncertain_significance |
| . | . | . | . | . | . | . | Uncertain_significance |
| . | . | . | . | . | . | . | Uncertain_significance |
| . | . | . | . | . | . | . | Uncertain_significance |
| . | . | . | . | . | . | . | Uncertain_significance |
| D | D | D | D | T | N | M | . |
| D | D | D | D | T | D | M | Uncertain_significance |
| D | D | D | D | D | D | L | Uncertain_significance |
| . | . | . | . | . | . | . | Uncertain_significance |
| . | . | . | . | . | . | . | . |
| . | . | . | . | . | . | . | Uncertain_significance |

#STS-26T cell line variants with a potential impact on protein function

| chr | start | end | Sample | Gene.refGene | REF |
|---|---|---|---|---|---|
| chr1 | 24074186 | 24074186 | STS-26T | MYOM3 | G |
| chr1 | 26282344 | 26282362 | STS-26T | UBXN11 | ACCGGGACCGGGACTG |
| chr1 | 26345087 | 26345087 | STS-26T | CRYBG2 | A |
| chr1 | 29258679 | 29258679 | STS-26T | PTPRU | C |
| chr1 | 32780239 | 32780239 | STS-26T | YARS | C |
| chr1 | 42789795 | 42789795 | STS-26T | TMEM269 | G |
| chr1 | 54800992 | 54800992 | STS-26T | TTC22 | G |
| chr1 | 56742201 | 56742201 | STS-26T | FYB2 | C |
| chr1 | 74336077 | 74336077 | STS-26T | FPGT-TNNI3K\x3b | C |
| chr1 | 84965613 | 84965613 | STS-26T | MCOLN2 | G |
| chr1 | 94043362 | 94043362 | STS-26T | ABCA4 | C |
| chr1 | 145856613 | 145856613 | STS-26T | PIAS3 | C |
| chr1 | 145960573 | 145960573 | STS-26T | ANKRD34A | C |
| chr1 | 146066541 | 146066541 | STS-26T | NBPF10 | T |
| chr1 | 146067225 | 146067225 | STS-26T | NBPF10 | C |
| chr1 | 146069569 | 146069569 | STS-26T | NBPF10 | C |
| chr1 | 149390802 | 149390802 | STS-26T | NOTCH2NLC | A |
| chr1 | 150557962 | 150557962 | STS-26T | ADAMTSL4 | G |
| chr1 | 152911463 | 152911463 | STS-26T | IVL | C |
| chr1 | 155208991 | 155208991 | STS-26T | MTX1 | T |
| chr1 | 158626214 | 158626214 | STS-26T | SPTA1 | C |
| chr1 | 167920631 | 167920631 | STS-26T | MPC2 | C |
| chr1 | 173557431 | 173557431 | STS-26T | SLC9C2 | A |
| chr1 | 176765790 | 176765790 | STS-26T | PAPPA2 | G |
| chr1 | 205059090 | 205059090 | STS-26T | CNTN2 | C |
| chr1 | 241957774 | 241957774 | STS-26T | BECN2 | C |
| chr1 | 246900339 | 246900339 | STS-26T | AHCTF1 | T |
| chr2 | 50922637 | 50922637 | STS-26T | NRXN1 | G |
| chr2 | 112332465 | 112332465 | STS-26T | ZC3H6 | C |
| chr2 | 132320213 | 132320213 | STS-26T | ZNF806 | A |
| chr2 | 137056548 | 137056548 | STS-26T | THSD7B | G |
| chr2 | 137115148 | 137115148 | STS-26T | THSD7B | G |
| chr2 | 168907568 | 168907568 | STS-26T | G6PC2 | G |
| chr2 | 168927319 | 168927319 | STS-26T | ABCB11 | A |

| | | | | | |
|---|---|---|---|---|---|
| chr2 | 171725924 | 171725924 | STS-26T | DYNC1I2 | C |
| chr2 | 188989432 | 188989432 | STS-26T | COL3A1 | G |
| chr2 | 189455433 | 189455433 | STS-26T | WDR75 | T |
| chr2 | 218428306 | 218428306 | STS-26T | VIL1 | G |
| chr2 | 240590833 | 240590833 | STS-26T | CAPN10 | C |
| chr3 | 16885062 | 16885062 | STS-26T | PLCL2 | G |
| chr3 | 38096696 | 38096696 | STS-26T | DLEC1 | G |
| chr3 | 38096697 | 38096697 | STS-26T | DLEC1 | G |
| chr3 | 98007903 | 98007903 | STS-26T | GABRR3 | A |
| chr3 | 125177889 | 125177889 | STS-26T | SLC12A8 | C |
| chr3 | 126472043 | 126472043 | STS-26T | ZXDC | G |
| chr3 | 130391203 | 130391203 | STS-26T | COL6A5 | A |
| chr3 | 130662309 | 130662309 | STS-26T | COL6A6 | G |
| chr4 | 1172686 | 1172686 | STS-26T | SPON2 | T |
| chr4 | 5989064 | 5989064 | STS-26T | C4orf50 | A |
| chr4 | 11399567 | 11399567 | STS-26T | HS3ST1 | G |
| chr4 | 11399822 | 11399822 | STS-26T | HS3ST1 | C |
| chr4 | 37443856 | 37443856 | STS-26T | NWD2 | G |
| chr4 | 55896772 | 55896772 | STS-26T | EXOC1 | G |
| chr4 | 56354103 | 56354103 | STS-26T | AASDH | A |
| chr4 | 56446740 | 56446740 | STS-26T | PAICS | T |
| chr4 | 68668125 | 68668125 | STS-26T | UGT2B15 | C |
| chr4 | 69485338 | 69485338 | STS-26T | UGT2B4 | G |
| chr4 | 78911125 | 78911125 | STS-26T | BMP2K | G |
| chr4 | 119031942 | 119031942 | STS-26T | SYNPO2 | C |
| chr4 | 143214167 | 143214167 | STS-26T | USP38 | C |
| chr4 | 150683625 | 150683633 | STS-26T | LRBA | TGTCACGTG |
| chr4 | 154570575 | 154570575 | STS-26T | FGB | G |
| chr5 | 35659193 | 35659193 | STS-26T | SPEF2 | C |
| chr5 | 62353284 | 62353284 | STS-26T | KIF2A | A |
| chr5 | 112838721 | 112838721 | STS-26T | APC | C |
| chr5 | 132559359 | 132559359 | STS-26T | RAD50 | G |
| chr5 | 136356930 | 136356930 | STS-26T | TRPC7 | G |
| chr5 | 140552043 | 140552043 | STS-26T | SRA1 | A |
| chr5 | 141137592 | 141137592 | STS-26T | PCDHB5 | C |
| chr5 | 141184133 | 141184133 | STS-26T | PCDHB16 | A |

| | | | | | |
|---|---|---|---|---|---|
| chr5 | 141184153 | 141184153 | STS-26T | PCDHB16 | G |
| chr5 | 141375100 | 141375100 | STS-26T | PCDHGA6 | T |
| chr5 | 146878727 | 146878730 | STS-26T | PPP2R2B | AGCT |
| chr5 | 149980428 | 149980428 | STS-26T | SLC26A2 | C |
| chr6 | 10751156 | 10751156 | STS-26T | TMEM14B | G |
| chr6 | 16327684 | 16327684 | STS-26T | ATXN1 | A |
| chr6 | 31815978 | 31815978 | STS-26T | HSPA1A | T |
| chr6 | 36899601 | 36899601 | STS-26T | C6orf89 | C |
| chr6 | 41061555 | 41061555 | STS-26T | APOBEC2 | C |
| chr6 | 43448894 | 43448894 | STS-26T | ABCC10 | A |
| chr6 | 62047954 | 62047954 | STS-26T | KHDRBS2 | G |
| chr6 | 87589938 | 87589939 | STS-26T | RARS2 | CG |
| chr6 | 89387512 | 89387512 | STS-26T | RRAGD | G |
| chr6 | 127829497 | 127829497 | STS-26T | THEMIS | C |
| chr6 | 130878120 | 130878120 | STS-26T | EPB41L2 | G |
| chr6 | 136346027 | 136346027 | STS-26T | MAP7 | C |
| chr6 | 146398946 | 146398946 | STS-26T | GRM1 | C |
| chr7 | 1547017 | 1547017 | STS-26T | TMEM184A | A |
| chr7 | 18829459 | 18829459 | STS-26T | HDAC9 | A |
| chr7 | 19116876 | 19116876 | STS-26T | TWIST1 | A |
| chr7 | 20681607 | 20681607 | STS-26T | ABCB5 | G |
| chr7 | 21744447 | 21744447 | STS-26T | DNAH11 | T |
| chr7 | 23260029 | 23260029 | STS-26T | GPNMB | C |
| chr7 | 25228344 | 25228344 | STS-26T | NPVF | C |
| chr7 | 26193346 | 26193346 | STS-26T | HNRNPA2B1 | T |
| chr7 | 34684653 | 34684653 | STS-26T | NPSR1 | G |
| chr7 | 48389204 | 48389204 | STS-26T | ABCA13 | G |
| chr7 | 70771559 | 70771559 | STS-26T | AUTS2 | C |
| chr7 | 71335709 | 71335709 | STS-26T | GALNT17 | C |
| chr7 | 74779322 | 74779322 | STS-26T | NCF1 | A |
| chr7 | 87566858 | 87566858 | STS-26T | ABCB1 | G |
| chr7 | 88282672 | 88282672 | STS-26T | STEAP4 | C |
| chr7 | 96080367 | 96080367 | STS-26T | DYNC1I1 | C |
| chr7 | 99520306 | 99520306 | STS-26T | ZKSCAN5 | T |
| chr7 | 107915609 | 107915609 | STS-26T | DLD | G |
| chr7 | 140753336 | 140753336 | STS-26T | BRAF | A |

| | | | | | |
|---|---|---|---|---|---|
| chr7 | 142036239 | 142036239 | STS-26T | MGAM | G |
| chr7 | 142040156 | 142040156 | STS-26T | MGAM | G |
| chr7 | 142131021 | 142131021 | STS-26T | MGAM2 | C |
| chr7 | 142864293 | 142864296 | STS-26T | EPHB6 | CCCT |
| chr7 | 142961033 | 142961033 | STS-26T | KEL | G |
| chr7 | 143756314 | 143756314 | STS-26T | CTAGE6 | G |
| chr7 | 151195597 | 151195598 | STS-26T | IQCA1L | TC |
| chr8 | 1676560 | 1676560 | STS-26T | DLGAP2 | G |
| chr8 | 2962543 | 2962543 | STS-26T | CSMD1 | G |
| chr8 | 3214634 | 3214634 | STS-26T | CSMD1 | C |
| chr8 | 8377352 | 8377358 | STS-26T | PRAG1 | CGCCGCT |
| chr8 | 19822512 | 19822512 | STS-26T | INTS10 | A |
| chr8 | 54621018 | 54621018 | STS-26T | RP1 | G |
| chr8 | 94838575 | 94838575 | STS-26T | INTS8 | C |
| chr8 | 113019172 | 113019172 | STS-26T | CSMD3 | C |
| chr8 | 141494938 | 141494938 | STS-26T | MROH5 | C |
| chr8 | 143921023 | 143921023 | STS-26T | PLEC | A |
| chr8 | 144379425 | 144379425 | STS-26T | ADCK5 | C |
| chr9 | 21971018 | 21971018 | STS-26T | CDKN2A | G |
| chr9 | 39102694 | 39102694 | STS-26T | CNTNAP3 | G |
| chr9 | 41991703 | 41991703 | STS-26T | CNTNAP3B | G |
| chr9 | 104569865 | 104569865 | STS-26T | OR13C8 | G |
| chr9 | 110549959 | 110549959 | STS-26T | SVEP1 | C |
| chr9 | 116235622 | 116235622 | STS-26T | PAPPA | G |
| chr9 | 123136105 | 123136105 | STS-26T | STRBP | G |
| chr9 | 127516899 | 127516899 | STS-26T | NIBAN2 | G |
| chr9 | 134701287 | 134701287 | STS-26T | COL5A1 | G |
| chr10 | 19088097 | 19088097 | STS-26T | MALRD1 | C |
| chr10 | 46549754 | 46549754 | STS-26T | GPRIN2 | G |
| chr10 | 49625530 | 49625530 | STS-26T | CHAT | G |
| chr10 | 68946400 | 68946400 | STS-26T | DDX50 | C |
| chr10 | 75022147 | 75022153 | STS-26T | KAT6B | GGAAGAA |
| chr10 | 75398985 | 75398985 | STS-26T | ZNF503 | T |
| chr10 | 87961113 | 87961113 | STS-26T | PTEN | T |
| chr10 | 87961114 | 87961114 | STS-26T | PTEN | T |
| chr10 | 101010757 | 101010757 | STS-26T | PDZD7 | T |

| | | | | | | |
|---|---|---|---|---|---|---|
| chr11 | 488638 | 488638 | STS-26T | | PTDSS2 | C |
| chr11 | 4882141 | 4882141 | STS-26T | | OR51T1 | C |
| chr11 | 5248480 | 5248480 | STS-26T | | HBG1 | C |
| chr11 | 55554873 | 55554873 | STS-26T | | OR4C15 | G |
| chr11 | 86951776 | 86951776 | STS-26T | | FZD4 | C |
| chr11 | 92353616 | 92353616 | STS-26T | | FAT3 | G |
| chr11 | 94973303 | 94973303 | STS-26T | | CWC15 | G |
| chr11 | 101042064 | 101042064 | STS-26T | | PGR | C |
| chr11 | 121053623 | 121053623 | STS-26T | | TBCEL | G |
| chr11 | 124023373 | 124023373 | STS-26T | | OR10G9 | C |
| chr11 | 134360257 | 134360257 | STS-26T | | GLB1L2 | G |
| chr12 | 6936728 | 6936728 | STS-26T | | ATN1 | A |
| chr12 | 11186007 | 11186007 | STS-26T | | TAS2R42 | G |
| chr12 | 11186175 | 11186175 | STS-26T | | TAS2R42 | A |
| chr12 | 13608823 | 13608823 | STS-26T | | GRIN2B | C |
| chr12 | 22644023 | 22644023 | STS-26T | | ETNK1 | G |
| chr12 | 25108825 | 25108825 | STS-26T | | CASC1 | T |
| chr12 | 33407055 | 33407055 | STS-26T | | SYT10 | C |
| chr12 | 49326136 | 49326136 | STS-26T | | TROAP | A |
| chr12 | 52058871 | 52058871 | STS-26T | | NR4A1 | A |
| chr12 | 66463071 | 66463071 | STS-26T | | GRIP1 | C |
| chr12 | 70353913 | 70353914 | STS-26T | | CNOT2 | TA |
| chr12 | 85057186 | 85057186 | STS-26T | | LRRIQ1 | T |
| chr13 | 102748299 | 102748299 | STS-26T | | CCDC168 | C |
| chr13 | 106493168 | 106493168 | STS-26T | | EFNB2 | G |
| chr14 | 18985482 | 18985482 | STS-26T | | POTEM | G |
| chr14 | 21525364 | 21525364 | STS-26T | | SALL2 | A |
| chr14 | 21665434 | 21665434 | STS-26T | | OR4E2 | A |
| chr14 | 21665783 | 21665783 | STS-26T | | OR4E2 | G |
| chr14 | 22877070 | 22877070 | STS-26T | | LRP10 | G |
| chr14 | 36711448 | 36711448 | STS-26T | | SLC25A21 | A |
| chr14 | 89931608 | 89931608 | STS-26T | | EFCAB11 | C |
| chr14 | 92071010 | 92071010 | STS-26T | | ATXN3 | C |
| chr14 | 93586807 | 93586822 | STS-26T | | UNC79 | TTGCCTGAAGATTCTC |
| chr15 | 20534729 | 20534729 | STS-26T | | GOLGA6L6 | C |
| chr15 | 22608089 | 22608089 | STS-26T | | GOLGA8J | C |

| | | | | | |
|---|---|---|---|---|---|
| chr15 | 22612072 | 22612072 | STS-26T | GOLGA8J | T |
| chr15 | 22903836 | 22903836 | STS-26T | CYFIP1 | C |
| chr15 | 23647279 | 23647279 | STS-26T | MAGEL2 | G |
| chr15 | 31229302 | 31229304 | STS-26T | LOC283710 | CGG |
| chr15 | 33773593 | 33773593 | STS-26T | RYR3 | G |
| chr15 | 40407805 | 40407807 | STS-26T | IVD | GTT |
| chr15 | 41184530 | 41184530 | STS-26T | EXD1 | C |
| chr15 | 41888042 | 41888042 | STS-26T | SPTBN5 | G |
| chr15 | 52253328 | 52253328 | STS-26T | MYO5C | C |
| chr15 | 58599600 | 58599600 | STS-26T | ADAM10 | G |
| chr15 | 64418656 | 64418656 | STS-26T | TRIP4 | A |
| chr15 | 85244616 | 85244616 | STS-26T | GOLGA6L3 | G |
| chr15 | 88881558 | 88881558 | STS-26T | HAPLN3 | C |
| chr16 | 260008 | 260015 | STS-26T | FAM234A | GGGCCAAC |
| chr16 | 5028162 | 5028162 | STS-26T | NAGPA | G |
| chr16 | 9763519 | 9763519 | STS-26T | GRIN2A | G |
| chr16 | 67987919 | 67987919 | STS-26T | DPEP2 | C |
| chr16 | 71069298 | 71069298 | STS-26T | HYDIN | G |
| chr16 | 71284878 | 71284878 | STS-26T | CMTR2 | G |
| chr16 | 87851781 | 87851781 | STS-26T | SLC7A5 | C |
| chr17 | 7676127 | 7676137 | STS-26T | TP53 | GTAGGAGCTGC |
| chr17 | 11923879 | 11923879 | STS-26T | DNAH9 | G |
| chr17 | 41528150 | 41528150 | STS-26T | KRT19 | G |
| chr17 | 41854575 | 41854575 | STS-26T | KLHL11 | T |
| chr17 | 43884083 | 43884083 | STS-26T | MPP2 | C |
| chr17 | 45274184 | 45274185 | STS-26T | MAP3K14 | GC |
| chr17 | 75813215 | 75813215 | STS-26T | UNK | T |
| chr18 | 712349 | 712367 | STS-26T | ENOSF1 | GCCGCGCGTACCATGG( |
| chr18 | 11886988 | 11886988 | STS-26T | MPPE1 | C |
| chr18 | 23810503 | 23810503 | STS-26T | LAMA3 | G |
| chr18 | 31326941 | 31326941 | STS-26T | DSG1 | G |
| chr19 | 1065019 | 1065019 | STS-26T | ABCA7 | G |
| chr19 | 8917625 | 8917625 | STS-26T | MUC16 | C |
| chr19 | 10453304 | 10453304 | STS-26T | PDE4A | C |
| chr19 | 11447525 | 11447531 | STS-26T | PRKCSH | AGAGGAG |
| chr19 | 15682425 | 15682425 | STS-26T | CYP4F12 | C |

| | | | | | |
|---|---|---|---|---|---|
| chr19 | 17821347 | 17821347 | STS-26T | INSL3 | C |
| chr19 | 17834887 | 17834887 | STS-26T | JAK3 | C |
| chr19 | 21383406 | 21383406 | STS-26T | ZNF738 | A |
| chr19 | 35351919 | 35351919 | STS-26T | FFAR1 | C |
| chr19 | 39894126 | 39894126 | STS-26T | FCGBP | C |
| chr19 | 39902274 | 39902274 | STS-26T | FCGBP | G |
| chr19 | 39918471 | 39918471 | STS-26T | FCGBP | C |
| chr19 | 41577497 | 41577497 | STS-26T | CEACAM21 | A |
| chr19 | 41579520 | 41579520 | STS-26T | CEACAM21 | G |
| chr19 | 41586462 | 41586462 | STS-26T | CEACAM21 | A |
| chr19 | 41717688 | 41717688 | STS-26T | CEACAM5 | G |
| chr19 | 41761981 | 41761981 | STS-26T | CEACAM6 | T |
| chr19 | 41999272 | 41999272 | STS-26T | GRIK5 | C |
| chr19 | 42910648 | 42910648 | STS-26T | PSG6 | G |
| chr19 | 49670259 | 49670259 | STS-26T | BCL2L12 | C |
| chr19 | 53634561 | 53634561 | STS-26T | DPRX | G |
| chr19 | 56192035 | 56192035 | STS-26T | ZSCAN5B | C |
| chr19 | 56824615 | 56824615 | STS-26T | PEG3 | T |
| chr19 | 57941112 | 57941112 | STS-26T | ZNF256 | G |
| chr20 | 14326155 | 14326155 | STS-26T | FLRT3 | G |
| chr20 | 17481834 | 17481834 | STS-26T | PCSK2 | G |
| chr20 | 23365094 | 23365094 | STS-26T | GZF1 | A |
| chr20 | 44501182 | 44501182 | STS-26T | SERINC3 | G |
| chr20 | 46725215 | 46725215 | STS-26T | SLC2A10 | T |
| chr20 | 47310147 | 47310147 | STS-26T | ZMYND8 | G |
| chr20 | 53254377 | 53254377 | STS-26T | TSHZ2 | G |
| chr20 | 58715174 | 58715174 | STS-26T | NPEPL1 | C |
| chr21 | 10569534 | 10569534 | STS-26T | TPTE | C |
| chr22 | 15690174 | 15690174 | STS-26T | POTEH | G |
| chr22 | 18121695 | 18121695 | STS-26T | TUBA8 | G |
| chr22 | 25029315 | 25029315 | STS-26T | KIAA1671 | A |
| chr22 | 32398013 | 32398013 | STS-26T | RTCB | T |
| chr22 | 35330371 | 35330371 | STS-26T | TOM1 | A |
| chr22 | 37070604 | 37070604 | STS-26T | TMPRSS6 | G |
| chr22 | 39678064 | 39678064 | STS-26T | CACNA1I | C |
| chr22 | 42127941 | 42127941 | STS-26T | CYP2D6\x3bCYP2 | G |

| | | | | | |
|---|---|---|---|---|---|
| chr22 | 50548686 | 50548686 | STS-26T | KLHDC7B | G |
| chrX | 318607 | 318607 | STS-26T | GTPBP6 | C |
| chrX | 1193297 | 1193297 | STS-26T | CRLF2 | T |
| chrX | 3103825 | 3103825 | STS-26T | ARSF | G |
| chrX | 50420491 | 50420491 | STS-26T | DGKK | C |
| chrX | 75784274 | 75784274 | STS-26T | MAGEE2 | C |
| chrX | 91877385 | 91877385 | STS-26T | PCDH11X | T |
| chrX | 115170393 | 115170393 | STS-26T | LRCH2 | T |
| chrX | 124420330 | 124420330 | STS-26T | TENM1 | C |
| chrX | 141003560 | 141003560 | STS-26T | SPANXB1 | C |
| chrX | 141905702 | 141905702 | STS-26T | MAGEC1 | C |
| chrX | 148662440 | 148662440 | STS-26T | AFF2 | C |

| ALT | Func.refGene | ExonicFunc.refGene | VAF |
|---|---|---|---|
| A | exonic | nonsynonymous_SNV | 0.8 |
| A | exonic | nonframeshift_deletion | 0.88888888888 |
| G | exonic | nonsynonymous_SNV | 0.70588235294 |
| A | exonic | nonsynonymous_SNV | 0.77272727272 |
| T | exonic | nonsynonymous_SNV | 0.29090909090 |
| A | splicing | . | 0.76923076923 |
| A | exonic | stopgain | 0.70833333333 |
| T | exonic | nonsynonymous_SNV | 0.75 |
| G | exonic | nonsynonymous_SNV | 0.66666666666 |
| A | exonic | nonsynonymous_SNV | 775 |
| T | exonic | nonsynonymous_SNV | 0.29545454545 |
| T | exonic | nonsynonymous_SNV | 0.55 |
| T | exonic | nonsynonymous_SNV | 0.43333333333 |
| G | exonic | nonsynonymous_SNV | 0.55555555555 |
| A | exonic | nonsynonymous_SNV | 0.53333333333 |
| T | exonic | nonsynonymous_SNV | 1 |
| AGGCGGCGGCGGCGGCGGCGGCGGCGGC | exonic | nonframeshift_insertion | 1 |
| A | exonic | stopgain | 0.37931034482 |
| T | exonic | stopgain | 0.5 |
| A | exonic | nonsynonymous_SNV | 1 |
| T | exonic | nonsynonymous_SNV | 0.51020408163 |
| T | exonic | nonsynonymous_SNV | 0.50980392156 |
| C | exonic | nonsynonymous_SNV | 0.54545454545 |
| A | exonic | nonsynonymous_SNV | 0.4 |
| T | exonic | nonsynonymous_SNV | 575 |
| T | exonic | nonsynonymous_SNV | 0.42105263157 |
| A | exonic | nonsynonymous_SNV | 0.43902439024 |
| A | exonic | nonsynonymous_SNV | 1 |
| T | exonic | nonsynonymous_SNV | 1 |
| G | exonic | nonsynonymous_SNV | 1 |
| A | exonic | nonsynonymous_SNV | 1 |
| A | exonic | stopgain | 1 |
| C | exonic | nonsynonymous_SNV | 1 |
| C | exonic | nonsynonymous_SNV | 1 |

| | | | |
|---|---|---|---|
| T | exonic | nonsynonymous_SNV | 1 |
| A | exonic | nonsynonymous_SNV | 1 |
| G | exonic | nonsynonymous_SNV | 1 |
| A | exonic | nonsynonymous_SNV | 1 |
| T | exonic | nonsynonymous_SNV | 1 |
| A | exonic | nonsynonymous_SNV | 1 |
| A | exonic | nonsynonymous_SNV | 1 |
| A | exonic | nonsynonymous_SNV | 1 |
| T | exonic | stopgain | 0.24657534246 |
| T | exonic | nonsynonymous_SNV | 0.54761904761 |
| A | exonic | nonsynonymous_SNV | 0.40540540540 |
| C | exonic | nonsynonymous_SNV | 0.45161290322 |
| A | splicing | . | 0.6 |
| A | splicing | . | 0.4 |
| T | exonic | nonsynonymous_SNV | 1 |
| A | exonic | stopgain | 0.14285714285 |
| T | exonic | nonsynonymous_SNV | 0.28571428571 |
| A | exonic | stopgain | 0.13333333333 |
| A | exonic | nonsynonymous_SNV | 0.43589743589 |
| C | exonic | nonsynonymous_SNV | 0.22580645161 |
| C | exonic | nonsynonymous_SNV | 0.32 |
| A | exonic | nonsynonymous_SNV | 0.3 |
| A | exonic | nonsynonymous_SNV | 0.33333333333 |
| A | exonic | nonsynonymous_SNV | 0.47368421052 |
| T | exonic | nonsynonymous_SNV | 0.48648648648 |
| T | exonic | nonsynonymous_SNV | 0.55555555555 |
| T | exonic | frameshift_deletion | 0.26315789473 |
| A | exonic | stopgain | 0.57692307692 |
| T | exonic | nonsynonymous_SNV | 0.77142857142 |
| C | exonic | nonsynonymous_SNV | 0.14285714285 |
| T | exonic | nonsynonymous_SNV | 0.65517241379 |
| A | exonic | nonsynonymous_SNV | 0.52941176470 |
| A | exonic | nonsynonymous_SNV | 0.3 |
| AGTC | exonic | nonframeshift_insertion | 1 |
| T | exonic | nonsynonymous_SNV | 0.54285714285 |
| G | exonic | nonsynonymous_SNV | 1 |

| | | | |
|---|---|---|---|
| A | exonic | nonsynonymous_SNV | 1 |
| G | exonic | nonsynonymous_SNV | 0.75 |
| AGCTGCTGCTGCTGCT | exonic | nonframeshift_insertior | 1 |
| T | exonic | nonsynonymous_SNV | 0.62 |
| T | exonic | nonsynonymous_SNV | 0.68181818181 |
| ATGCTGCTGCTGCTGC | exonic | nonframeshift_insertior | 0.38461538461 |
| G | exonic | nonsynonymous_SNV | 1 |
| T | exonic | nonsynonymous_SNV | 0.28 |
| A | exonic | nonsynonymous_SNV | 0.72 |
| T | exonic | nonsynonymous_SNV | 0.5 |
| A | exonic | nonsynonymous_SNV | 0.6 |
| C | exonic | frameshift_deletion | 0.64 |
| A | exonic | nonsynonymous_SNV | 0.59615384615 |
| T | exonic | nonsynonymous_SNV | 0.44897959183 |
| A | exonic | nonsynonymous_SNV | 0.35294117647 |
| T | exonic | nonsynonymous_SNV | 0.38461538461 |
| G | exonic | nonsynonymous_SNV | 0.5 |
| ACCC | exonic | nonframeshift_insertior | 0.38461538461 |
| C | splicing | . | 0.35849056603 |
| G | exonic | nonsynonymous_SNV | 0.15384615384 |
| A | exonic | nonsynonymous_SNV | 0.41379310344 |
| G | exonic | nonsynonymous_SNV | 0.29577464788 |
| G | exonic | nonsynonymous_SNV | 0.453125 |
| G | exonic | nonsynonymous_SNV | 0.68085106382 |
| C | exonic | nonsynonymous_SNV | 0.4 |
| A | exonic | nonsynonymous_SNV | 0.46666666666 |
| A | exonic | nonsynonymous_SNV | 0.63793103448 |
| T | exonic | nonsynonymous_SNV | 0.35849056603 |
| T | exonic | nonsynonymous_SNV | 0.36363636363 |
| G | exonic | nonsynonymous_SNV | 0.63333333333 |
| A | exonic | nonsynonymous_SNV | 0.65573770491 |
| A | exonic | nonsynonymous_SNV | 0.59649122807 |
| T | exonic | nonsynonymous_SNV | 0.31914893617 |
| C | splicing | . | 0.59574468085 |
| A | exonic | nonsynonymous_SNV | 0.53846153846 |
| T | exonic | nonsynonymous_SNV | 0.53191489361 |

| | | | |
|---|---|---|---|
| A | exonic | nonsynonymous_SNV | 0.48387096774 |
| A | exonic | stopgain | 0.67391304347 |
| T | exonic | nonsynonymous_SNV | 0.3125 |
| CCCTCCT | exonic | nonframeshift_insertior | 1 |
| A | exonic | nonsynonymous_SNV | 0.50980392156 |
| A | exonic | nonsynonymous_SNV | 0.37037037037 |
| T | exonic | frameshift_deletion | 0.55 |
| A | exonic | nonsynonymous_SNV | 0.60714285714 |
| T | exonic | nonsynonymous_SNV | 0.71428571428 |
| T | exonic | nonsynonymous_SNV | 0.4 |
| C | exonic | nonframeshift_deletion | 1 |
| C | exonic | nonsynonymous_SNV | 0.57692307692 |
| A | exonic | nonsynonymous_SNV | 0.8 |
| T | exonic | nonsynonymous_SNV | 0.34615384615 |
| T | exonic | nonsynonymous_SNV | 0.70370370370 |
| T | splicing | . | 0.86111111111 |
| G | exonic | nonsynonymous_SNV | 0.78571428571 |
| A | exonic | nonsynonymous_SNV | 1 |
| A | exonic | nonsynonymous_SNV | 1 |
| A | exonic | nonsynonymous_SNV | 0.94285714285 |
| T | exonic | nonsynonymous_SNV | 0.65306122448 |
| A | exonic | nonsynonymous_SNV | 1 |
| T | exonic | nonsynonymous_SNV | 1 |
| A | exonic | nonsynonymous_SNV | 1 |
| T | exonic | nonsynonymous_SNV | 1 |
| A | exonic | stopgain | 1 |
| A | exonic | nonsynonymous_SNV | 1 |
| T | exonic | nonsynonymous_SNV | 1 |
| C | exonic | nonsynonymous_SNV | 0.60714285714 |
| A | exonic | nonsynonymous_SNV | 1 |
| T | exonic | nonsynonymous_SNV | 1 |
| GGAA | exonic | nonframeshift_deletion | 1 |
| C | exonic | nonsynonymous_SNV | 0.45 |
| A | exonic | nonsynonymous_SNV | 1 |
| A | exonic | nonsynonymous_SNV | 1 |
| G | exonic | nonsynonymous_SNV | 1 |

| | | | |
|---|---|---|---|
| T | exonic | nonsynonymous_SNV | 0.51515151515 |
| T | exonic | nonsynonymous_SNV | 0.5 |
| T | exonic | nonsynonymous_SNV | 0.48648648648 |
| A | exonic | nonsynonymous_SNV | 0.30555555555 |
| A | exonic | nonsynonymous_SNV | 0.13333333333 |
| A | exonic | nonsynonymous_SNV | 1 |
| A | splicing | . | 1 |
| T | exonic | nonsynonymous_SNV | 1 |
| GT | exonic | frameshift_insertion | 1 |
| T | exonic | nonsynonymous_SNV | 1 |
| C | exonic | nonsynonymous_SNV | 0.5 |
| ACAGCAGCAGCAGCAGCAG | exonic | nonframeshift_insertior | 875 |
| C | exonic | nonsynonymous_SNV | 1 |
| C | exonic | nonsynonymous_SNV | 1 |
| T | exonic | nonsynonymous_SNV | 0.4 |
| GT | splicing | . | 0.28260869565 |
| TAAAAAAAAAAAAAAAAAAAA | splicing | . | 0.92857142857 |
| T | exonic | nonsynonymous_SNV | 1 |
| G | exonic | nonsynonymous_SNV | 1 |
| T | exonic | nonsynonymous_SNV | 0.68571428571 |
| T | exonic | nonsynonymous_SNV | 1 |
| TAA | exonic | nonframeshift_insertior | 0.85714285714 |
| C | splicing | . | 1 |
| T | exonic | nonsynonymous_SNV | 0.5 |
| A | exonic | nonsynonymous_SNV | 0.56097560975 |
| A | splicing | . | 0.58333333333 |
| G | exonic | nonsynonymous_SNV | 1 |
| G | exonic | nonsynonymous_SNV | 1 |
| A | exonic | nonsynonymous_SNV | 1 |
| A | exonic | nonsynonymous_SNV | 1 |
| T | exonic | nonsynonymous_SNV | 1 |
| T | exonic | nonsynonymous_SNV | 1 |
| CCTGCTGCTGCTGCTGCTGCTGCTGCTGCTGCTGCTGCTG | exonic | nonframeshift_insertior | 1 |
| T | exonic | nonframeshift_deletion | 0.57142857142 |
| T | exonic | nonsynonymous_SNV | 0.27272727272 |
| T | exonic | nonsynonymous_SNV | 0.5 |

| | | | |
|---|---|---|---|
| A | exonic | nonsynonymous_SNV | 0.37254901960 |
| T | exonic | nonsynonymous_SNV | 0.51282051282 |
| A | exonic | nonsynonymous_SNV | 0.36363636363 |
| CG | exonic | frameshift_deletion | 0.85714285714 |
| A | exonic | nonsynonymous_SNV | 0.46428571428 |
| G | exonic | frameshift_deletion | 0.44444444444 |
| T | exonic | nonsynonymous_SNV | 0.5625 |
| A | exonic | nonsynonymous_SNV | 0.52 |
| T | exonic | nonsynonymous_SNV | 0.53333333333 |
| A | exonic | nonsynonymous_SNV | 0.55555555555 |
| G | exonic | nonsynonymous_SNV | 0.25 |
| A | exonic | nonsynonymous_SNV | 0.19047619047 |
| T | exonic | nonsynonymous_SNV | 0.5 |
| G | exonic | frameshift_deletion | 0.65517241379 |
| A | exonic | nonsynonymous_SNV | 0.44444444444 |
| A | exonic | nonsynonymous_SNV | 0.47222222222 |
| T | exonic | stopgain | 0.43243243243 |
| A | exonic | nonsynonymous_SNV | 0.61764705882 |
| A | exonic | nonsynonymous_SNV | 0.54285714285 |
| T | exonic | nonsynonymous_SNV | 0.48275862068 |
| G | exonic | frameshift_deletion | 1 |
| A | exonic | nonsynonymous_SNV | 0.20833333333 |
| A | exonic | nonsynonymous_SNV | 1 |
| A | exonic | nonsynonymous_SNV | 1 |
| T | splicing | . | 1 |
| G | exonic | frameshift_deletion | 1 |
| C | splicing | . | 0.6 |
| G | exonic | frameshift_deletion | 0.84210526315 |
| CGG | exonic | frameshift_insertion | 0.39473684210 |
| A | exonic | nonsynonymous_SNV | 0.48275862068 |
| A | exonic | stopgain | 0.63333333333 |
| A | exonic | nonsynonymous_SNV | 1 |
| T | exonic | stopgain | 0.76595744680 |
| T | exonic | nonsynonymous_SNV | 0.35294117647 |
| AGAG | exonic | nonframeshift_deletion | 0.9375 |
| T | exonic | nonsynonymous_SNV | 0.29629629629 |

| | | | |
|---|---|---|---|
| T | exonic | nonsynonymous_SNV | 0.59090909090 |
| T | exonic | nonsynonymous_SNV | 0.35294117647 |
| C | exonic | nonsynonymous_SNV | 375 |
| T | exonic | nonsynonymous_SNV | 0.55882352941 |
| T | exonic | nonsynonymous_SNV | 0.18947368421 |
| C | exonic | nonsynonymous_SNV | 0.44444444444 |
| G | exonic | nonsynonymous_SNV | 0.47916666666 |
| C | exonic | nonsynonymous_SNV | 0.765625 |
| A | exonic | nonsynonymous_SNV | 0.51666666666 |
| T | splicing | . | 0.40909090909 |
| A | exonic | nonsynonymous_SNV | 0.66666666666 |
| G | exonic | nonsynonymous_SNV | 1 |
| T | exonic | nonsynonymous_SNV | 0.54166666666 |
| A | exonic | nonsynonymous_SNV | 0.38888888888 |
| T | exonic | nonsynonymous_SNV | 0.34693877551 |
| A | exonic | nonsynonymous_SNV | 0.39130434782 |
| G | exonic | nonsynonymous_SNV | 0.66 |
| A | exonic | nonsynonymous_SNV | 0.24561403508 |
| A | exonic | stopgain | 0.54385964912 |
| A | exonic | nonsynonymous_SNV | 0.36170212765 |
| A | exonic | nonsynonymous_SNV | 0.74509803921 |
| C | exonic | nonsynonymous_SNV | 0.41860465116 |
| A | exonic | nonsynonymous_SNV | 0.26595744680 |
| A | exonic | nonsynonymous_SNV | 0.3125 |
| A | exonic | nonsynonymous_SNV | 0.28205128205 |
| A | exonic | nonsynonymous_SNV | 0.27941176470 |
| G | exonic | nonsynonymous_SNV | 0.71929824561 |
| T | exonic | nonsynonymous_SNV | 0.34666666666 |
| C | exonic | nonsynonymous_SNV | 0.42857142857 |
| C | exonic | nonsynonymous_SNV | 0.25490196078 |
| G | exonic | nonsynonymous_SNV | 0.41666666666 |
| A | exonic | nonsynonymous_SNV | 0.5 |
| G | exonic | nonsynonymous_SNV | 0.38888888888 |
| A | exonic | nonsynonymous_SNV | 0.19047619047 |
| T | exonic | nonsynonymous_SNV | 0.67857142857 |
| A | exonic | nonsynonymous_SNV | 1 |

| | | | |
|---|---|---|---|
| A | exonic | nonsynonymous_SNV | 0.47368421052 |
| G | exonic | nonsynonymous_SNV | 1 |
| C | exonic | nonsynonymous_SNV | 1 |
| A | exonic | nonsynonymous_SNV | 1 |
| T | exonic | nonsynonymous_SNV | 1 |
| T | exonic | nonsynonymous_SNV | 1 |
| C | exonic | nonsynonymous_SNV | 0.52941176470 |
| G | exonic | nonsynonymous_SNV | 1 |
| T | exonic | nonsynonymous_SNV | 1 |
| G | exonic | nonsynonymous_SNV | 1 |
| T | exonic | stopgain | 1 |
| A | exonic | stopgain | 1 |

avsnp150
.
.
rs11579835
.
rs863224717
.
.
rs138668500
.
.
rs187071406
.
.
.
.
.
.
.
.
rs760077
.
.
.
.
.
.
.
.
.
rs12468523
rs752809070
.
.
.

.
.
.
.
.
.
.
.
rs832032
.
.
.
rs201329510
.
.
rs771189478
.
.
rs35001804
rs148777026
.
.
.
rs143310663
.
.
.
.
.
.
.
rs370769989
.
.
rs400562
rs17844651

rs2697532
.
.
rs104893915
rs17849403
.
.
.
rs41273362
.
.
.
.
.
.
.
.
.
.
.
.
.
rs17147995
rs886354
.
.
rs775942969
.
.
rs10614
.
rs73205916
.
.
rs145670503
rs113488022

.
.
.
.
.
.
.
.
.
.
.
.
.
.
.
rs6578193
.
rs6599528
rs121913386
.
rs62555055
rs150858980
.
.
.
.
.
.
rs4445576
.
.
.
.
.
.
.

rs753104582
.
.
rs771730511
.
.
rs2276350
.
.
rs747156389
.
.
rs1650017
rs1669413
.
.
.
rs370609371
.
.
.
.
.
.
.
rs201661142
rs1263811
rs2874103
rs970382
rs142153001
.
.
.
.
.
.

rs146352451
rs7170637
rs1002045692
.
.
.
.
.
.
.
.
rs200601580
.
.
.
.
.
.
.
rs151257488
.
.
.
.
rs231518
.
.
.
rs561966131
.
.
.
.
.
.
rs2285888

rs754081979
rs3213409
rs62110372
rs578010155
rs201693960
rs139175656
.
rs714106
rs2302188
rs3745936
rs7249230
rs11548735
.
.
.
rs761892759
.
rs559813104
rs776056252
.
.
.
rs758858636
.
.
.
.
rs76723236
rs201322170
.
rs17667531
.
rs34371697
.
.
rs16947

.
.
.
.
rs376014418
rs267606512
.
.
.
rs878856926
rs145052423
.

| Variant | AAChange.refGene | GeneDetail.refGene |
| --- | --- | --- |
| chr1:24074186_G/A | MYOM3:NM_152372:e | . |
| chr1:26282344_ACCGGGACCGGGACTGGGG/A | UBXN11:NM_0010772 | . |
| chr1:26345087_A/G | CRYBG2:NM_0010397 | . |
| chr1:29258679_C/A | PTPRU:NM_00119500 | . |
| chr1:32780239_C/T | YARS:NM_003680:exo | . |
| chr1:42789795_G/A | . | NM_001354602:exon2: |
| chr1:54800992_G/A | TTC22:NM_00111410 | . |
| chr1:56742201_C/T | FYB2:NM_001004303: | . |
| chr1:74336077_C/G | TNNI3K:NM_015978:e | . |
| chr1:84965613_G/A | MCOLN2:NM_153259: | . |
| chr1:94043362_C/T | ABCA4:NM_000350:ex | . |
| chr1:145856613_C/T | PIAS3:NM_006099:exc | . |
| chr1:145960573_C/T | ANKRD34A:NM_00103 | . |
| chr1:146066541_T/G | NBPF10:NM_0010397 | . |
| chr1:146067225_C/A | NBPF10:NM_0010397 | . |
| chr1:146069569_C/T | NBPF10:NM_0010397 | . |
| chr1:149390802_A/AGGCGGCGGCGGCGGCGGCGGCGGC | NOTCH2NLC:NM_0013 | . |
| chr1:150557962_G/A | ADAMTSL4:NM_01903 | . |
| chr1:152911463_C/T | IVL:NM_005547:exon2 | . |
| chr1:155208991_T/A | MTX1:NM_002455:exc | . |
| chr1:158626214_C/T | SPTA1:NM_003126:ex | . |
| chr1:167920631_C/T | MPC2:NM_001143674 | . |
| chr1:173557431_A/C | SLC9C2:NM_178527:e | . |
| chr1:176765790_G/A | PAPPA2:NM_020318:e | . |
| chr1:205059090_C/T | CNTN2:NM_00134608 | . |
| chr1:241957774_C/T | BECN2:NM_00129069 | . |
| chr1:246900339_T/A | AHCTF1:NM_0013233 | . |
| chr2:50922637_G/A | NRXN1:NM_00133008 | . |
| chr2:112332465_C/T | ZC3H6:NM_198581:ex | . |
| chr2:132320213_A/G | ZNF806:NM_0013554 | . |
| chr2:137056548_G/A | THSD7B:NM_0013163 | . |
| chr2:137115148_G/A | THSD7B:NM_0013163 | . |
| chr2:168907568_G/C | G6PC2:NM_00108168 | . |
| chr2:168927319_A/C | ABCB11:NM_003742:e | . |

| | | |
|---|---|---|
| chr2:171725924_C/T | | . |
| chr2:188989432_G/A | COL3A1:NM_000090:e | . |
| chr2:189455433_T/G | WDR75:NM_032168:e | . |
| chr2:218428306_G/A | VIL1:NM_007127:exon | . |
| chr2:240590833_C/T | CAPN10:NM_023083:e | . |
| chr3:16885062_G/A | PLCL2:NM_001144382 | . |
| chr3:38096696_G/A | DLEC1:NM_001321153 | . |
| chr3:38096697_G/A | DLEC1:NM_001321153 | . |
| chr3:98007903_A/T | GABRR3:NM_0011055 | . |
| chr3:125177889_C/T | SLC12A8:NM_0011954 | . |
| chr3:126472043_G/A | ZXDC:NM_001040653: | . |
| chr3:130391203_A/C | COL6A5:NM_00127829 | . |
| chr3:130662309_G/A | . | NM_001102608:exon34 |
| chr4:1172686_T/A | . | NM_001199021:exon3: |
| chr4:5989064_A/T | C4orf50:NM_00136469 | . |
| chr4:11399567_G/A | HS3ST1:NM_005114:e | . |
| chr4:11399822_C/T | HS3ST1:NM_005114:e | . |
| chr4:37443856_G/A | NWD2:NM_00114499( | . |
| chr4:55896772_G/A | EXOC1:NM_178237:ex | . |
| chr4:56354103_A/C | | . |
| chr4:56446740_T/C | PAICS:NM_001079524 | . |
| chr4:68668125_C/A | UGT2B15:NM_001076 | . |
| chr4:69485338_G/A | UGT2B4:NM_021139:e | . |
| chr4:78911125_G/A | BMP2K:NM_198892:e | . |
| chr4:119031942_C/T | SYNPO2:NM_0011289 | . |
| chr4:143214167_C/T | USP38:NM_001290325 | . |
| chr4:150683625_TGTCACGTG/T | LRBA:NM_001199282: | . |
| chr4:154570575_G/A | FGB:NM_001184741:e | . |
| chr5:35659193_C/T | SPEF2:NM_024867:ex | . |
| chr5:62353284_A/C | KIF2A:NM_001098511 | . |
| chr5:112838721_C/T | | . |
| chr5:132559359_G/A | RAD50:NM_005732:ex | . |
| chr5:136356930_G/A | TRPC7:NM_001167576 | . |
| chr5:140552043_A/AGTC | SRA1:NM_001035235: | . |
| chr5:141137592_C/T | PCDHB5:NM_015669:e | . |
| chr5:141184133_A/G | PCDHB16:NM_020957 | . |

| | | |
|---|---|---|
| chr5:141184153_G/A | PCDHB16:NM_020957 | . |
| chr5:141375100_T/G | PCDHGA6:NM_01891 | . |
| chr5:146878727_AGCT/AGCTGCTGCTGCTGCT | PPP2R2B:NM_181675: | . |
| chr5:149980428_C/T | SLC26A2:NM_000112: | . |
| chr6:10751156_G/T | TMEM14B:NM_00128 | . |
| chr6:16327684_A/ATGCTGCTGCTGCTGC | ATXN1:NM_00112816 | . |
| chr6:31815978_T/G | HSPA1A:NM_005345:e | . |
| chr6:36899601_C/T | C6orf89:NM_152734:e | . |
| chr6:41061555_C/A | APOBEC2:NM_006789 | . |
| chr6:43448894_A/T | ABCC10:NM_033450:e | . |
| chr6:62047954_G/A | KHDRBS2:NM_00135 | . |
| chr6:87589938_CG/C | RARS2:NM_00135050 | . |
| chr6:89387512_G/A | RRAGD:NM_021244:e | . |
| chr6:127829497_C/T | THEMIS:NM_0010109 | . |
| chr6:130878120_G/A | EPB41L2:NM_0013502 | . |
| chr6:136346027_C/T | | . |
| chr6:146398946_C/G | GRM1:NM_001278064 | . |
| chr7:1547017_A/ACCC | TMEM184A:NM_0010 | . |
| chr7:18829459_A/C | . | NM_001321868:exon1 |
| chr7:19116876_A/G | TWIST1:NM_000474:e | . |
| chr7:20681607_G/A | ABCB5:NM_178559:ex | . |
| chr7:21744447_T/G | DNAH11:NM_0012771 | . |
| chr7:23260029_C/G | GPNMB:NM_0010053 | . |
| chr7:25228344_C/G | NPVF:NM_022150:exo | . |
| chr7:26193346_T/C | HNRNPA2B1:NM_0021 | . |
| chr7:34684653_G/A | NPSR1:NM_00130093 | . |
| chr7:48389204_G/A | ABCA13:NM_152701:e | . |
| chr7:70771559_C/T | AUTS2:NM_00112723 | . |
| chr7:71335709_C/T | GALNT17:NM_022479 | . |
| chr7:74779322_A/G | NCF1:NM_000265:exo | . |
| chr7:87566858_G/A | ABCB1:NM_00134894 | . |
| chr7:88282672_C/A | STEAP4:NM_024636:e | . |
| chr7:96080367_C/T | DYNC1I1:NM_0011355 | . |
| chr7:99520306_T/C | . | NM_001318082:exon5: |
| chr7:107915609_G/A | DLD:NM_001289750:e | . |
| chr7:140753336_A/T | BRAF:NM_001354609: | . |

| | | |
|---|---|---|
| chr7:142036239_G/A | MGAM:NM_00136569 | . |
| chr7:142040156_G/A | MGAM:NM_00136569 | . |
| chr7:142131021_C/T | MGAM2:NM_0012936 | . |
| chr7:142864293_CCCT/CCCTCCT | EPHB6:NM_004445:ex | . |
| chr7:142961033_G/A | KEL:NM_000420:exon | . |
| chr7:143756314_G/A | CTAGE6:NM_178561:e | . |
| chr7:151195597_TC/T | IQCA1L:NM_00130441 | . |
| chr8:1676560_G/A | DLGAP2:NM_001346 | . |
| chr8:2962543_G/T | CSMD1:NM_033225:e | . |
| chr8:3214634_C/T | CSMD1:NM_033225:e | . |
| chr8:8377352_CGCCGCT/C | | . |
| chr8:19822512_A/C | | . |
| chr8:54621018_G/A | RP1:NM_006269:exon | . |
| chr8:94838575_C/T | INTS8:NM_017864:exo | . |
| chr8:113019172_C/T | CSMD3:NM_00136318 | . |
| chr8:141494938_C/T | . | NM_207414:exon4:c.4 |
| chr8:143921023_A/G | | . |
| chr8:144379425_C/A | ADCK5:NM_174922:ex | . |
| chr9:21971018_G/A | CDKN2A:NM_000077: | . |
| chr9:39102694_G/A | CNTNAP3:NM_033655 | . |
| chr9:41991703_G/T | CNTNAP3B:NM_00120 | . |
| chr9:104569865_G/A | OR13C8:NM_0010044 | . |
| chr9:110549959_C/T | SVEP1:NM_153366:ex | . |
| chr9:116235622_G/A | PAPPA:NM_002581:ex | . |
| chr9:123136105_G/T | STRBP:NM_00117113 | . |
| chr9:127516899_G/A | NIBAN2:NM_0010355 | . |
| chr9:134701287_G/A | COL5A1:NM_000093:e | . |
| chr10:19088097_C/T | MALRD1:NM_0011423 | . |
| chr10:46549754_G/C | GPRIN2:NM_014696:e | . |
| chr10:49625530_G/A | | . |
| chr10:68946400_C/T | DDX50:NM_024045:ex | . |
| chr10:75022147_GGAAGAA/GGAA | | . |
| chr10:75398985_T/C | ZNF503:NM_032772:e | . |
| chr10:87961113_T/A | PTEN:NM_000314:exo | . |
| chr10:87961114_T/A | PTEN:NM_000314:exo | . |
| chr10:101010757_T/G | PDZD7:NM_0011952 | . |

| | | |
|---|---|---|
| chr11:488638_C/T | PTDSS2:NM_00132954 | . |
| chr11:4882141_C/T | OR51T1:NM_0010047 | . |
| chr11:5248480_C/T | HBG1:NM_000559:exc | . |
| chr11:55554873_G/A | OR4C15:NM_001001 | . |
| chr11:86951776_C/A | FZD4:NM_012193:exo | . |
| chr11:92353616_G/A | FAT3:NM_001008781: | . |
| chr11:94973303_G/A | . | NM_001363372:exon1: |
| chr11:101042064_C/T | PGR:NM_001271161:e | . |
| chr11:121053623_G/GT | TBCEL:NM_001130047 | . |
| chr11:124023373_C/T | OR10G9:NM_0010019 | . |
| chr11:134360257_G/C | GLB1L2:NM_0013704 | . |
| chr12:6936728_A/ACAGCAGCAGCAGCAGCAG | ATN1:NM_001007026 | . |
| chr12:11186007_G/C | TAS2R42:NM_181429: | . |
| chr12:11186175_A/C | TAS2R42:NM_181429: | . |
| chr12:13608823_C/T | GRIN2B:NM_000834:e | . |
| chr12:22644023_G/GT | . | NM_018638:exon2:c.41 |
| chr12:25108825_T/TAAAAAAAAAAAAAAAAAAAA | . | |
| chr12:33407055_C/T | SYT10:NM_198992:ex | . |
| chr12:49326136_A/G | TROAP:NM_005480:ex | . |
| chr12:52058871_A/T | NR4A1:NM_173157:ex | . |
| chr12:66463071_C/T | GRIP1:NM_001178074 | . |
| chr12:70353913_TA/TAA | CNOT2:NM_00119930 | . |
| chr12:85057186_T/C | . | NM_001079910:exon8: |
| chr13:102748299_C/T | CCDC168:NM_001146 | . |
| chr13:106493168_G/A | EFNB2:NM_004093:ex | . |
| chr14:18985482_G/A | . | NM_001145442:exon7: |
| chr14:21525364_A/G | SALL2:NM_001291446 | . |
| chr14:21665434_A/G | OR4E2:NM_00100191 | . |
| chr14:21665783_G/A | OR4E2:NM_00100191 | . |
| chr14:22877070_G/A | LRP10:NM_014045:ex | . |
| chr14:36711448_A/T | SLC25A21:NM_001171 | . |
| chr14:89931608_C/T | EFCAB11:NM_001284 | . |
| chr14:92071010_C/CCTGCTGCTGCTGCTGCTGCTGCTGCTGCTGCTGCTGCTG | | . |
| chr14:93586807_TTGCCTGAAGATTCTC/T | UNC79:NM_00134621 | . |
| chr15:20534729_C/T | GOLGA6L6:NM_00114 | . |
| chr15:22608089_C/T | GOLGA8J:NM_001282 | . |

| | | |
|---|---|---|
| chr15:22612072_T/A | GOLGA8J:NM_001282 | . |
| chr15:22903836_C/T | | . |
| chr15:23647279_G/A | MAGEL2:NM_019066: | . |
| chr15:31229302_CGG/CG | LOC283710:NM_00124 | . |
| chr15:33773593_G/A | RYR3:NM_001036:exo | . |
| chr15:40407805_GTT/G | IVD:NM_001354599:e | . |
| chr15:41184530_C/T | EXD1:NM_152596:exo | . |
| chr15:41888042_G/A | SPTBN5:NM_016642:e | . |
| chr15:52253328_C/T | MYO5C:NM_018728:e | . |
| chr15:58599600_G/A | ADAM10:NM_001320 | . |
| chr15:64418656_A/G | TRIP4:NM_001321924 | . |
| chr15:85244616_G/A | GOLGA6L3:NM_00131 | . |
| chr15:88881558_C/T | HAPLN3:NM_178232: | . |
| chr16:260008_GGGCCAAC/G | FAM234A:NM_001284 | . |
| chr16:5028162_G/A | NAGPA:NM_016256:e | . |
| chr16:9763519_G/A | GRIN2A:NM_0011344 | . |
| chr16:67987919_C/T | DPEP2:NM_00132415 | . |
| chr16:71069298_G/A | HYDIN:NM_00119854 | . |
| chr16:71284878_G/A | CMTR2:NM_00132437 | . |
| chr16:87851781_C/T | SLC7A5:NM_003486:e | . |
| chr17:7676127_GTAGGAGCTGC/G | | . |
| chr17:11923879_G/A | DNAH9:NM_004662:e | . |
| chr17:41528150_G/A | KRT19:NM_002276:ex | . |
| chr17:41854575_T/A | KLHL11:NM_018143:e | . |
| chr17:43884083_C/T | . | NM_001278372:exon4: |
| chr17:45274184_GC/G | MAP3K14:NM_003954 | . |
| chr17:75813215_T/C | . | NM_001080419:exon5: |
| chr18:712349_GCCGCGCGTACCATGGCGT/G | ENOSF1:NM_0013540 | . |
| chr18:11886988_C/CGG | MPPE1:NM_00124290 | . |
| chr18:23810503_G/A | LAMA3:NM_00112771 | . |
| chr18:31326941_G/A | DSG1:NM_001942:exo | . |
| chr19:1065019_G/A | ABCA7:NM_019112:ex | . |
| chr19:8917625_C/T | MUC16:NM_024690:e | . |
| chr19:10453304_C/T | PDE4A:NM_006202:ex | . |
| chr19:11447525_AGAGGAG/AGAG | PRKCSH:NM_0012891 | . |
| chr19:15682425_C/T | CYP4F12:NM_023944: | . |

| | | |
|---|---|---|
| chr19:17821347_C/T | INSL3:NM_001265587 | . |
| chr19:17834887_C/T | JAK3:NM_000215:exon | . |
| chr19:21383406_A/C | ZNF738:NM_00135523 | . |
| chr19:35351919_C/T | FFAR1:NM_005303:exo | . |
| chr19:39894126_C/T | FCGBP:NM_003890:ex | . |
| chr19:39902274_G/C | FCGBP:NM_003890:ex | . |
| chr19:39918471_C/G | FCGBP:NM_003890:ex | . |
| chr19:41577497_A/C | CEACAM21:NM_00109 | . |
| chr19:41579520_G/A | CEACAM21:NM_00109 | . |
| chr19:41586462_A/T | . | NM_001288773:exon8: |
| chr19:41717688_G/A | CEACAM5:NM_001291 | . |
| chr19:41761981_T/G | CEACAM6:NM_002483 | . |
| chr19:41999272_C/T | GRIK5:NM_002088:exo | . |
| chr19:42910648_G/A | PSG6:NM_001031850: | . |
| chr19:49670259_C/T | BCL2L12:NM_0010406 | . |
| chr19:53634561_G/A | DPRX:NM_001012728: | . |
| chr19:56192035_C/G | ZSCAN5B:NM_001080 | . |
| chr19:56824615_T/A | PEG3:NM_001146184: | . |
| chr19:57941112_G/A | ZNF256:NM_005773:e | . |
| chr20:14326155_G/A | FLRT3:NM_013281:exo | . |
| chr20:17481834_G/A | PCSK2:NM_00120152 | . |
| chr20:23365094_A/C | GZF1:NM_001317019: | . |
| chr20:44501182_G/A | SERINC3:NM_006811:e | . |
| chr20:46725215_T/A | SLC2A10:NM_030777: | . |
| chr20:47310147_G/A | | . |
| chr20:53254377_G/A | TSHZ2:NM_001193421 | . |
| chr20:58715174_C/G | NPEPL1:NM_024663:e | . |
| chr21:10569534_C/T | TPTE:NM_001290224: | . |
| chr22:15690174_G/C | POTEH:NM_00113621 | . |
| chr22:18121695_G/C | TUBA8:NM_00119341 | . |
| chr22:25029315_A/G | KIAA1671:NM_001145 | . |
| chr22:32398013_T/A | RTCB:NM_014306:exo | . |
| chr22:35330371_A/G | TOM1:NM_00113573C | . |
| chr22:37070604_G/A | TMPRSS6:NM_00128 | . |
| chr22:39678064_C/T | CACNA1I:NM_001003 | . |
| chr22:42127941_G/A | CYP2D6:NM_001025 | . |

| | | |
|---|---|---|
| chr22:50548686_G/A | KLHDC7B:NM_138433 | . |
| chrX:318607_C/G | GTPBP6:NM_012227:e | . |
| chrX:1193297_T/C | CRLF2:NM_001012288 | . |
| chrX:3103825_G/A | ARSF:NM_001201538: | . |
| chrX:50420491_C/T | DGKK:NM_001013742 | . |
| chrX:75784274_C/T | MAGEE2:NM_138703: | . |
| chrX:91877385_T/C | | . |
| chrX:115170393_T/G | LRCH2:NM_00124396 | . |
| chrX:124420330_C/T | TENM1:NM_014253:e | . |
| chrX:141003560_C/G | SPANXB1:NM_032461 | . |
| chrX:141905702_C/T | MAGEC1:NM_005462: | . |
| chrX:148662440_C/A | AFF2:NM_001169122: | . |

| AF | AF_popmax | prop_pathogenic_pred | CLNALLELEID |
|---|---|---|---|
| . | . | 0.857142857142857 | . |
| 0.0004 | 0.0015 | NA | . |
| 0.0016 | 0.0039 | 1 | . |
| . | . | 0.857142857142857 | . |
| . | . | 0.857142857142857 | 212086 |
| . | . | NA | . |
| . | . | 1 | . |
| 0.0039 | 0.0065 | 0.857142857142857 | . |
| . | . | 0.857142857142857 | . |
| . | . | 0.857142857142857 | . |
| . | . | 0.714285714285714 | . |
| . | . | NA | . |
| . | . | NA | . |
| . | . | NA | . |
| . | . | NA | . |
| . | . | NA | . |
| . | . | NA | . |
| . | . | 1 | . |
| . | . | 1 | . |
| . | . | NA | . |
| . | . | 0.8 | . |
| . | . | 0.857142857142857 | . |
| . | . | 0.714285714285714 | . |
| . | . | 1 | . |
| . | . | 0.714285714285714 | . |
| . | . | NA | . |
| . | . | 0.857142857142857 | . |
| . | . | NA | . |
| . | . | 0.714285714285714 | . |
| . | . | NA | . |
| . | . | 0.714285714285714 | . |
| . | . | 1 | . |
| . | . | 1 | . |
| . | . | 0.857142857142857 | . |

| | | | |
|---|---|---|---|
| . | . | 0.714285714285714 | . |
| . | . | 1 | . |
| . | . | 0.714285714285714 | . |
| . | . | 1 | . |
| . | . | 0.857142857142857 | . |
| . | . | 1 | . |
| . | . | 0.857142857142857 | . |
| . | . | 0.857142857142857 | . |
| . | . | NA | . |
| . | . | 1 | . |
| . | . | 0.857142857142857 | . |
| . | . | 1 | . |
| 0.0006 | 1 | 1 | . |
| . | . | NA | . |
| . | . | 0.666666666666667 | . |
| . | . | 1 | . |
| . | . | 1 | . |
| . | . | 1 | . |
| 0.0053 | 0.0086 | 1 | . |
| 0.0003 | 0.0095 | 0.857142857142857 | . |
| . | . | 0.714285714285714 | . |
| . | . | 0.714285714285714 | . |
| . | . | 0.857142857142857 | . |
| 0.0039 | 6 | 0.714285714285714 | . |
| . | . | 0.714285714285714 | . |
| . | . | 0.857142857142857 | . |
| . | . | NA | . |
| . | . | 1 | . |
| . | . | 0.857142857142857 | . |
| . | . | 0.714285714285714 | . |
| . | . | 0.857142857142857 | 559909 |
| . | . | 0.857142857142857 | 182499 |
| . | . | 0.857142857142857 | . |
| . | . | NA | . |
| . | . | NA | . |
| . | . | NA | . |

| | | | |
|---|---|---|---|
| . | . | NA | . |
| . | . | 0.714285714285714 | . |
| . | . | NA | . |
| 0.0008 | 0.0024 | 1 | 19128 |
| 0.0051 | 0.0054 | 0.857142857142857 | . |
| . | . | NA | . |
| . | . | 0.666666666666667 | . |
| . | . | 0.714285714285714 | . |
| 0.0048 | 8 | 0.857142857142857 | . |
| . | . | 0.857142857142857 | . |
| . | . | 0.857142857142857 | . |
| . | . | NA | . |
| . | . | 1 | . |
| . | . | 0.857142857142857 | . |
| . | . | 0.714285714285714 | . |
| . | . | 0.714285714285714 | . |
| . | . | 0.714285714285714 | . |
| . | . | NA | . |
| . | . | 1 | . |
| . | . | 1 | . |
| . | . | 0.857142857142857 | . |
| . | . | 0.857142857142857 | . |
| 0.0021 | 0.0037 | 0.857142857142857 | . |
| . | . | NA | . |
| . | . | 0.714285714285714 | . |
| . | . | 0.714285714285714 | . |
| . | . | 1 | . |
| . | . | 0.714285714285714 | . |
| . | . | 0.714285714285714 | . |
| . | . | NA | . |
| . | . | 0.857142857142857 | . |
| 0.0057 | 8 | 0.714285714285714 | . |
| . | . | 0.857142857142857 | . |
| . | . | 1 | . |
| 0.0006 | 0.0024 | 0.857142857142857 | 200140 |
| . | . | 0.857142857142857 | 29000 |

| | | | | |
|---|---|---|---|---|
| . | . | 1 | . | |
| . | . | 1 | . | |
| . | . | 0.75 | . | |
| . | . | NA | . | |
| . | . | 0.714285714285714 | . | |
| . | . | 0.75 | . | |
| . | . | NA | . | |
| . | . | 0.857142857142857 | . | |
| . | . | 0.714285714285714 | . | |
| . | . | 0.857142857142857 | . | |
| . | . | NA | . | |
| . | . | 0.833333333333333 | . | |
| . | . | 0.857142857142857 | . | |
| . | . | 0.833333333333333 | . | |
| . | . | 0.714285714285714 | . | |
| . | . | NA | . | |
| . | . | 0.714285714285714 | . | |
| . | . | NA | . | |
| . | . | 0.8 | | 88528 |
| . | . | 0.833333333333333 | . | |
| . | . | NA | . | |
| . | . | 0.714285714285714 | . | |
| . | . | 0.857142857142857 | . | |
| . | . | 0.714285714285714 | . | |
| . | . | 1 | . | |
| . | . | 1 | . | |
| . | . | 0.714285714285714 | . | |
| . | . | NA | . | |
| . | . | NA | . | |
| . | . | 0.714285714285714 | . | |
| . | . | 0.714285714285714 | . | |
| . | . | NA | . | |
| . | . | 0.714285714285714 | . | |
| . | . | 1 | . | |
| . | . | 1 | . | |
| . | . | NA | . | |

| | | | |
|---|---|---|---|
| . | . | 1 | . |
| . | . | 0.857142857142857 | . |
| . | . | 0.666666666666667 | . |
| . | . | 0.666666666666667 | . |
| . | . | 1 | . |
| . | . | 0.833333333333333 | . |
| . | . | NA | . |
| . | . | 0.857142857142857 | . |
| . | . | NA | . |
| . | . | 0.714285714285714 | . |
| . | . | NA | . |
| . | . | NA | . |
| . | . | NA | . |
| . | . | NA | . |
| . | . | 0.8 | . |
| . | . | NA | . |
| . | . | NA | . |
| . | . | 0.714285714285714 | . |
| . | . | 1 | . |
| . | . | 0.714285714285714 | . |
| . | . | 0.857142857142857 | . |
| . | . | NA | . |
| . | . | 1 | . |
| . | . | NA | . |
| . | . | 1 | . |
| . | . | 0 | . |
| . | . | NA | . |
| . | . | NA | . |
| . | . | NA | . |
| 0.0078 | 0.0095 | 0.857142857142857 | . |
| . | . | 0.857142857142857 | . |
| . | . | 0.714285714285714 | . |
| . | . | NA | . |
| . | . | NA | . |
| . | . | 0.75 | . |
| . | . | NA | . |

| | | | | |
|---|---|---|---|---|
| . | . | NA | | . |
| . | . | NA | | . |
| . | . | NA | | . |
| . | . | NA | | . |
| . | . | 0.714285714285714 | | . |
| . | . | NA | | . |
| . | . | 0.714285714285714 | | . |
| . | . | 0.857142857142857 | | . |
| . | . | | 1 | . |
| . | . | 0.857142857142857 | | . |
| . | . | | 1 | . |
| 0.0019 | 0.0025 | NA | | . |
| . | . | 0.857142857142857 | | . |
| . | . | NA | | . |
| . | . | 0.714285714285714 | | . |
| . | . | 0.857142857142857 | | . |
| . | . | | 1 | . |
| . | . | 0.714285714285714 | | . |
| . | . | 0.857142857142857 | | . |
| 0.0006 | 0.0024 | 0.714285714285714 | | . |
| . | . | NA | | . |
| . | . | 0.857142857142857 | | . |
| . | . | 0.857142857142857 | | . |
| . | . | | 1 | . |
| . | . | NA | | . |
| . | . | NA | | . |
| . | . | | 1 | . |
| . | . | NA | | . |
| 0.0008 | 0.0024 | NA | | . |
| . | . | 0.833333333333333 | | . |
| . | . | | 1 | . |
| . | . | 0.833333333333333 | | . |
| . | . | | 1 | . |
| . | . | 0.8 | | . |
| . | . | NA | | . |
| . | . | NA | | . |

| | | | | |
|---|---|---|---|---|
| . | . | 0.833333333333333 | . | |
| 0.0064 | 0.0099 | 0.142857142857143 | | 138312 |
| . | . | NA | . | |
| . | . | 0.857142857142857 | . | |
| 0.0048 | 0.0077 | 1 | . | |
| . | . | NA | . | |
| . | . | 1 | . | |
| . | . | NA | . | |
| . | . | NA | . | |
| . | . | NA | . | |
| . | . | NA | . | |
| . | . | NA | . | |
| . | . | 0.666666666666667 | . | |
| . | . | 0.666666666666667 | . | |
| . | . | 0.714285714285714 | . | |
| . | . | 0.833333333333333 | . | |
| . | . | 0.666666666666667 | . | |
| . | . | 0.666666666666667 | . | |
| . | . | 1 | . | |
| . | . | 0.714285714285714 | . | |
| . | . | 0.857142857142857 | . | |
| . | . | 0.714285714285714 | . | |
| . | . | 0.714285714285714 | . | |
| . | . | 0.714285714285714 | . | |
| . | . | 0.857142857142857 | . | |
| . | . | 0.857142857142857 | . | |
| . | . | 0.714285714285714 | . | |
| 3.573e-05 | . | NA | . | |
| . | . | NA | . | |
| . | . | 0.857142857142857 | . | |
| . | . | 0.666666666666667 | . | |
| . | . | 0.857142857142857 | . | |
| 0.0007 | 0.0013 | 0.857142857142857 | . | |
| . | . | 1 | . | |
| . | . | 1 | . | |
| . | . | NA | . | |

| | | | |
|---|---|---|---|
| . | . | 0.666666666666667 | . |
| . | . | NA | . |
| . | . | NA | . |
| . | . | 0.857142857142857 | . |
| 0.0003 | 1 | 1 | . |
| . | . | 0.666666666666667 | . |
| . | . | 0.857142857142857 | . |
| . | . | 0.857142857142857 | . |
| . | . | 0.857142857142857 | . |
| . | . | NA | . |
| . | . | 1 | . |
| . | . | 0.5 | . |

CLNDN
.
.
.
.
Charcot-Marie-Tooth_disease,_dominant_intermediate_C
.
.
.
.
.
.
.
.
.
.
.
.
.
.
.
.
.
.
.
.
.
.
.
.
.
.
.
.
.

.
.
.
.
.
.
.
.
.
.
.
.
.
.
.
.
.
.
.
.
.
.
.
.
.
.
.
.
.
.
Familial_adenomatous_polyposis_1
Hereditary_cancer-predisposing_syndrome
.
.
.
.

.
.
.
Inborn_genetic_diseases|Osteochondrodysplasia|Diastrophic_dysplasia|Achondrogenesis,_type_IB|[illegible]
.
.
.
.
.
.
.
.
.
.
.
.
.
.
.
.
.
.
.
.
.
.
.
.
.
.
.
.
.
.
Maple_syrup_urine_disease,_type_3|not_specified|not_provided

.
.
.
.
.
.
.
.
.
.
.
.
.
.
.
.
.
.
Squamous_cell_carcinoma_of_the_skin|Cutaneous_melanoma|Squamous_cell_lung_carcinoma|Squ
.
.
.
.
.
.
.
.
.
.
.
.
.
.
.
.
.

.
Acute_megakaryoblastic_leukemia|Severe_combined_immunodeficiency,_autosomal_recessive,_T_c
.
.
.
.
.
.
.
.
.
.
.
.
.
.
.
.
.
.
.
.
.
.
.
.
.
.
.
.
.
.
.
.
.
.

CLNDISDB
.
.
.
.
MedGen:C1842237,OMIM:608323,Orphanet:ORPHA100045
.
.
.
.
.
.
.
.
.
.
.
.
.
.
.
.
.
.
.
.
.
.
.
.
.
.
.
.
.

.
.
.
.
.
.
.
.
.
.
.
.
.
.
.
.
.
.
.
.
.
.
.
.
.
.
.
.
.
.
MedGen:C2713442,OMIM:175100
MedGen:C0027672,SNOMED_CT:699346009
.
.
.
.

.
.
.
Multiple_epiphyseal_dysplasia_4|Atelosteogenesis_type_2|SLC26A2-Related_Disorders|not_provided
.
.
.
.
.
.
.
.
.
.
.
.
.
.
.
.
.
.
.
.
.
.
.
.
.
.
.
.
.
.
.
MedGen:CN043137,OMIM:246900|MedGen:CN169374|MedGen:CN517202

.
.
.
.
.
.
.
.
.
.
.
.
.
.
.
.
.
.
amous_cell_carcinoma_of_the_head_and_neck|Malignant_melanoma_of_skin|Hereditary_cancer-pr
.
.
.
.
.
.
.
.
.
.
.
.
.
.
.
.
.

.
MeSH:D007947,MedGen:C0023462,Orphanet:ORPHA518|MedGen:C1833275,OMIM:600802,Orphan[illegible]
.
.
.
.
.
.
.
.
.
.
.
.
.
.
.
.
.
.
.
.
.
.
.
.
.
.
.
.
.
.
.
.
.
.
.
.
.

| CLNREVSTAT | CLNSIG | SIFT_pred | Polyphen2_HDIV_pred |
|---|---|---|---|
| . | . | D | D |
| . | . | . | . |
| . | . | . | . |
| . | . | D | D |
| criteria_provided,_single_submitter | Uncertain_significance | D | D |
| . | . | . | . |
| . | . | . | . |
| . | . | D | D |
| . | . | D | D |
| . | . | D | D |
| . | . | T | D |
| . | . | . | . |
| . | . | . | . |
| . | . | . | . |
| . | . | . | . |
| . | . | . | . |
| . | . | . | . |
| . | . | . | . |
| . | . | . | . |
| . | . | . | . |
| . | . | . | D |
| . | . | D | D |
| . | . | D | D |
| . | . | D | D |
| . | . | D | D |
| . | . | . | . |
| . | . | D | D |
| . | . | . | . |
| . | . | D | D |
| . | . | . | . |
| . | . | D | P |
| . | . | . | . |
| . | . | D | D |
| . | . | D | D |

| | | | |
|---|---|---|---|
| . | . | D | P |
| . | . | D | D |
| . | . | D | P |
| . | . | D | D |
| . | . | T | D |
| . | . | . | . |
| . | . | D | D |
| . | . | D | D |
| . | . | . | . |
| . | . | D | D |
| . | . | D | D |
| . | . | D | D |
| . | . | . | . |
| . | . | . | . |
| . | . | D | . |
| . | . | . | . |
| . | . | D | D |
| . | . | . | . |
| . | . | D | D |
| . | . | D | D |
| . | . | D | P |
| . | . | D | D |
| . | . | D | D |
| . | . | D | D |
| . | . | D | P |
| . | . | D | D |
| . | . | . | . |
| . | . | . | . |
| . | . | D | D |
| . | . | D | P |
| criteria_provided,_single_submitter | Uncertain_significance | D | D |
| criteria_provided,_conflicting_interpretations | Conflicting_interpretations_of_pathogenicity | D | D |
| . | . | D | D |
| . | . | . | . |
| . | . | . | . |
| . | . | . | . |

| | | | |
|---|---|---|---|
| . | . | . | . |
| . | . | D | D |
| . | . | . | . |
| criteria_provided,_multiple_submitters,_no_conflicts | Pathogenic | D | D |
| . | . | D | D |
| . | . | . | . |
| . | . | D | P |
| . | . | D | D |
| . | . | D | D |
| . | . | D | P |
| . | . | D | D |
| . | . | . | . |
| . | . | D | D |
| . | . | D | D |
| . | . | D | D |
| . | . | D | D |
| . | . | T | P |
| . | . | . | . |
| . | . | . | . |
| . | . | D | D |
| . | . | D | D |
| . | . | D | D |
| . | . | D | D |
| . | . | . | . |
| . | . | T | D |
| . | . | D | D |
| . | . | D | D |
| . | . | D | D |
| . | . | D | D |
| . | . | . | . |
| . | . | D | D |
| . | . | D | D |
| . | . | D | D |
| . | . | . | . |
| criteria_provided,_multiple_submitters,_no_conflicts | Uncertain_significance | D | D |
| criteria_provided,_single_submitter | Pathogenic | D | D |

| | | | |
|---|---|---|---|
| . | . | D | D |
| . | . | . | . |
| . | . | D | P |
| . | . | . | . |
| . | . | D | D |
| . | . | . | P |
| . | . | . | . |
| . | . | D | D |
| . | . | D | D |
| . | . | D | D |
| . | . | . | . |
| . | . | D | D |
| . | . | D | D |
| . | . | T | D |
| . | . | D | P |
| . | . | . | . |
| . | . | D | D |
| . | . | . | . |
| criteria_provided,_single_submitter | Likely_pathogenic | D | D |
| . | . | D | D |
| . | . | . | . |
| . | . | D | D |
| . | . | D | D |
| . | . | T | D |
| . | . | D | D |
| . | . | . | . |
| . | . | D | D |
| . | . | . | . |
| . | . | . | . |
| . | . | D | B |
| . | . | D | D |
| . | . | . | . |
| . | . | T | P |
| . | . | D | D |
| . | . | D | D |
| . | . | . | . |

| | | | |
|---|---|---|---|
| . | . | D | D |
| . | . | D | D |
| . | . | D | D |
| . | . | D | P |
| . | . | D | D |
| . | . | D | D |
| . | . | . | . |
| . | . | D | D |
| . | . | . | . |
| . | . | D | D |
| . | . | . | . |
| . | . | . | . |
| . | . | . | . |
| . | . | . | . |
| . | . | . | D |
| . | . | . | . |
| . | . | . | . |
| . | . | D | D |
| . | . | D | D |
| . | . | D | P |
| . | . | D | D |
| . | . | . | . |
| . | . | . | . |
| . | . | . | . |
| . | . | D | D |
| . | . | . | . |
| . | . | . | . |
| . | . | . | . |
| . | . | . | . |
| . | . | T | D |
| . | . | D | D |
| . | . | T | D |
| . | . | . | . |
| . | . | . | . |
| . | . | . | P |
| . | . | . | . |

| | | | |
|---|---|---|---|
| . | . | . | . |
| . | . | . | . |
| . | . | . | . |
| . | . | . | . |
| . | . | T | D |
| . | . | . | . |
| . | . | D | D |
| . | . | D | D |
| . | . | D | D |
| . | . | D | D |
| . | . | D | D |
| . | . | . | . |
| . | . | D | D |
| . | . | . | . |
| . | . | D | D |
| . | . | D | D |
| . | . | . | . |
| . | . | D | P |
| . | . | D | P |
| . | . | T | D |
| . | . | . | . |
| . | . | D | D |
| . | . | T | D |
| . | . | D | D |
| . | . | . | . |
| . | . | . | . |
| . | . | . | . |
| . | . | . | . |
| . | . | . | . |
| . | . | D | D |
| . | . | . | . |
| . | . | D | P |
| . | . | . | . |
| . | . | D | D |
| . | . | . | . |
| . | . | . | . |

| | | | |
|---|---|---|---|
| . | . | D | D |
| criteria_provided,_conflicting_interpretations | Conflicting_interpretations_of_pathogenicity | D | B |
| . | . | . | . |
| . | . | D | D |
| . | . | . | D |
| . | . | . | . |
| . | . | . | D |
| . | . | . | . |
| . | . | . | . |
| . | . | . | . |
| . | . | . | . |
| . | . | . | . |
| . | . | D | D |
| . | . | D | D |
| . | . | D | D |
| . | . | D | D |
| . | . | D | D |
| . | . | D | D |
| . | . | . | . |
| . | . | D | D |
| . | . | D | D |
| . | . | D | D |
| . | . | D | D |
| . | . | D | D |
| . | . | D | D |
| . | . | D | D |
| . | . | D | D |
| . | . | . | . |
| . | . | . | . |
| . | . | D | D |
| . | . | D | D |
| . | . | D | P |
| . | . | D | D |
| . | . | D | D |
| . | . | D | D |
| . | . | . | . |

| | | | |
|---|---|---|---|
| . | . | D | P |
| . | . | . | . |
| . | . | . | . |
| . | . | D | P |
| . | . | . | D |
| . | . | D | D |
| . | . | D | D |
| . | . | D | P |
| . | . | D | D |
| . | . | . | . |
| . | . | . | . |
| . | . | . | . |

| Polyphen2_HVAR_pred | LRT_pred | FATHMM_pred | MutationTaster_pred | MutationAssessor_pred | InterVar_automated |
|---|---|---|---|---|---|
| D | D | T | D | M | Uncertain_significance |
| . | . | . | . | . | . |
| . | . | . | P | . | . |
| D | D | T | D | M | Uncertain_significance |
| P | D | T | D | M | Uncertain_significance |
| . | . | . | . | . | . |
| . | D | . | A | . | Uncertain_significance |
| P | D | T | D | M | Uncertain_significance |
| D | D | T | D | M | Likely_pathogenic |
| P | D | T | D | M | Uncertain_significance |
| P | D | D | D | L | . |
| . | . | . | . | . | . |
| . | . | . | . | . | Uncertain_significance |
| . | . | . | . | . | . |
| . | . | . | . | . | Uncertain_significance |
| . | . | . | . | . | Uncertain_significance |
| . | . | . | . | . | . |
| . | . | . | A | . | Pathogenic |
| . | . | . | D | . | Uncertain_significance |
| . | . | . | . | . | Uncertain_significance |
| D | N | . | D | M | Uncertain_significance |
| D | D | T | D | H | Uncertain_significance |
| P | D | T | N | M | Uncertain_significance |
| D | D | D | D | M | Uncertain_significance |
| D | N | T | D | M | Uncertain_significance |
| . | . | . | . | . | Uncertain_significance |
| D | D | T | D | M | Uncertain_significance |
| . | . | . | . | . | . |
| D | N | T | D | M | Uncertain_significance |
| . | . | . | . | . | . |
| P | D | T | D | L | Uncertain_significance |
| . | D | . | A | . | Uncertain_significance |
| P | . | D | D | M | Uncertain_significance |
| D | D | D | D | L | . |

| | | | | | |
|---|---|---|---|---|---|
| B | D | T | D | M | Uncertain_significance |
| D | D | D | D | H | Uncertain_significance |
| B | D | T | D | M | Uncertain_significance |
| D | D | D | D | H | Uncertain_significance |
| D | D | D | D | M | Uncertain_significance |
| . | . | . | D | . | Uncertain_significance |
| D | D | T | D | M | Uncertain_significance |
| D | D | T | D | M | Uncertain_significance |
| . | . | . | . | . | Uncertain_significance |
| D | . | D | D | M | Uncertain_significance |
| D | D | T | D | M | Uncertain_significance |
| D | . | D | D | H | Uncertain_significance |
| . | . | . | D | . | . |
| . | . | . | . | . | . |
| . | . | T | P | . | . |
| . | D | . | D | . | Uncertain_significance |
| D | D | D | D | M | Uncertain_significance |
| . | D | . | D | . | Uncertain_significance |
| D | D | . | D | M | Uncertain_significance |
| D | D | T | D | H | Uncertain_significance |
| P | N | T | D | M | Uncertain_significance |
| P | U | T | D | M | . |
| D | U | D | D | H | Uncertain_significance |
| P | N | T | D | M | Uncertain_significance |
| B | D | T | D | M | Uncertain_significance |
| D | D | T | D | M | Uncertain_significance |
| . | . | . | . | . | . |
| . | D | . | D | . | Uncertain_significance |
| D | D | T | D | M | Uncertain_significance |
| B | D | T | D | M | Uncertain_significance |
| D | D | D | D | L | Uncertain_significance |
| D | D | T | D | M | Uncertain_significance |
| D | D | T | D | M | Uncertain_significance |
| . | . | . | . | . | . |
| . | . | . | . | . | Uncertain_significance |
| . | . | . | . | . | Uncertain_significance |

| | | | | | |
|---|---|---|---|---|---|
| . | . | . | . | . | Uncertain_significance |
| D | U | T | D | M | Uncertain_significance |
| . | . | . | . | . | . |
| D | D | D | A | H | Likely_pathogenic |
| D | D | T | D | H | Uncertain_significance |
| . | . | . | . | . | . |
| D | N | T | D | . | Uncertain_significance |
| P | N | T | D | M | Uncertain_significance |
| D | D | T | D | H | Uncertain_significance |
| P | D | D | D | L | Uncertain_significance |
| D | D | T | D | M | Uncertain_significance |
| . | . | . | . | . | . |
| D | D | D | D | M | Uncertain_significance |
| D | D | T | D | M | Uncertain_significance |
| D | N | D | D | L | . |
| P | N | T | D | M | Uncertain_significance |
| P | D | D | D | L | Uncertain_significance |
| . | . | . | . | . | . |
| . | . | . | D | . | . |
| D | D | D | D | H | . |
| D | N | D | D | H | Uncertain_significance |
| P | D | T | D | M | Uncertain_significance |
| P | D | T | D | M | Uncertain_significance |
| . | . | . | . | . | Uncertain_significance |
| P | D | D | D | L | Uncertain_significance |
| D | D | T | D | L | Uncertain_significance |
| D | D | D | D | H | Uncertain_significance |
| P | D | T | D | L | Uncertain_significance |
| P | N | T | D | M | Uncertain_significance |
| . | . | . | . | . | Uncertain_significance |
| D | D | T | D | M | Uncertain_significance |
| D | N | D | N | M | . |
| D | D | T | D | M | Uncertain_significance |
| . | . | . | D | . | . |
| D | D | T | D | M | Uncertain_significance |
| D | D | D | D | N | Uncertain_significance |

| | | | | | |
|---|---|---|---|---|---|
| D | D | D | D | H | Uncertain_significance |
| . | D | . | A | . | Uncertain_significance |
| P | . | T | . | . | Uncertain_significance |
| . | . | . | . | . | . |
| P | N | D | D | L | Uncertain_significance |
| P | . | T | . | M | Uncertain_significance |
| . | . | . | . | . | . |
| P | D | T | D | M | Uncertain_significance |
| D | D | T | D | L | . |
| D | D | T | D | M | . |
| . | . | . | . | . | . |
| D | D | . | D | L | Uncertain_significance |
| D | D | T | D | M | Uncertain_significance |
| P | D | . | D | M | Uncertain_significance |
| B | D | T | D | H | Uncertain_significance |
| . | . | . | . | . | . |
| D | U | T | D | M | Uncertain_significance |
| . | . | . | . | . | Uncertain_significance |
| D | . | T | D | . | Likely_pathogenic |
| P | . | T | D | M | Uncertain_significance |
| . | . | . | . | . | Uncertain_significance |
| D | N | T | D | M | Uncertain_significance |
| P | D | T | D | M | Uncertain_significance |
| P | D | T | D | M | Uncertain_significance |
| D | D | D | D | H | Uncertain_significance |
| . | D | . | A | . | Uncertain_significance |
| D | U | T | D | H | Uncertain_significance |
| . | . | . | . | . | Uncertain_significance |
| . | . | . | . | . | . |
| B | D | D | D | M | Uncertain_significance |
| D | D | T | D | L | Uncertain_significance |
| . | . | . | . | . | . |
| P | D | T | D | M | Uncertain_significance |
| D | D | D | D | M | Uncertain_significance |
| D | D | D | D | M | Uncertain_significance |
| . | . | . | . | . | . |

| | | | | | |
|---|---|---|---|---|---|
| D | D | . | D | M | Uncertain_significance |
| D | D | T | D | M | Uncertain_significance |
| D | U | D | N | . | Uncertain_significance |
| P | . | T | N | M | Uncertain_significance |
| D | D | D | D | M | Uncertain_significance |
| D | . | T | D | M | Uncertain_significance |
| . | . | . | . | . | . |
| D | D | D | D | L | . |
| . | . | . | . | . | . |
| D | N | T | D | M | Uncertain_significance |
| . | . | . | . | . | . |
| . | . | . | . | . | . |
| . | . | . | . | . | Uncertain_significance |
| . | . | . | . | . | Uncertain_significance |
| D | D | . | D | L | Uncertain_significance |
| . | . | . | . | . | . |
| . | . | . | . | . | . |
| D | U | T | D | H | Uncertain_significance |
| D | D | . | D | M | Uncertain_significance |
| P | D | T | D | L | Uncertain_significance |
| D | D | T | D | H | Uncertain_significance |
| . | . | . | . | . | . |
| . | . | . | D | . | . |
| . | . | . | . | . | Uncertain_significance |
| D | D | D | D | M | Uncertain_significance |
| . | . | . | N | . | . |
| . | . | . | . | . | Uncertain_significance |
| . | . | . | . | . | Uncertain_significance |
| . | . | . | . | . | Uncertain_significance |
| D | D | D | D | M | Uncertain_significance |
| D | D | T | D | M | Uncertain_significance |
| D | D | T | D | M | Uncertain_significance |
| . | . | . | . | . | . |
| . | . | . | . | . | . |
| P | . | . | N | M | Uncertain_significance |
| . | . | . | . | . | Uncertain_significance |

| | | | | | |
|---|---|---|---|---|---|
| . | . | . | . | . | . |
| . | . | . | . | . | Uncertain_significance |
| . | . | . | . | . | . |
| . | . | . | . | . | . |
| D | D | D | D | L | Uncertain_significance |
| . | . | . | . | . | . |
| D | D | T | D | L | Uncertain_significance |
| D | N | D | D | M | Uncertain_significance |
| D | D | D | D | H | Uncertain_significance |
| D | D | T | D | M | Uncertain_significance |
| D | D | . | D | M | Uncertain_significance |
| . | . | . | . | . | Uncertain_significance |
| D | D | T | D | H | Uncertain_significance |
| . | . | . | . | . | . |
| P | D | T | D | L | Uncertain_significance |
| P | D | T | D | M | Uncertain_significance |
| . | D | . | A | . | Uncertain_significance |
| P | U | T | D | M | Uncertain_significance |
| P | D | T | D | M | Uncertain_significance |
| D | D | D | D | L | Uncertain_significance |
| . | . | . | . | . | . |
| D | D | T | D | H | Uncertain_significance |
| D | D | D | D | M | Uncertain_significance |
| D | D | D | D | H | Uncertain_significance |
| . | . | . | . | . | . |
| . | . | . | . | . | . |
| . | . | . | D | . | . |
| . | . | . | . | . | . |
| . | . | . | . | . | . |
| D | . | T | D | M | Uncertain_significance |
| . | D | . | A | . | Pathogenic |
| P | . | D | N | M | Uncertain_significance |
| . | . | . | A | . | Uncertain_significance |
| D | . | T | D | . | Uncertain_significance |
| . | . | . | . | . | . |
| . | . | . | . | . | Uncertain_significance |

| | | | | | |
|---|---|---|---|---|---|
| D | . | D | N | M | Uncertain_significance |
| B | N | T | N | N | . |
| . | . | . | . | . | . |
| D | D | T | D | M | Uncertain_significance |
| D | . | . | D | M | Uncertain_significance |
| . | . | . | . | . | Uncertain_significance |
| D | D | . | D | H | . |
| . | . | . | . | . | Uncertain_significance |
| . | . | . | . | . | Uncertain_significance |
| . | . | . | . | . | . |
| . | . | . | . | . | Uncertain_significance |
| . | . | . | . | . | Uncertain_significance |
| D | . | T | D | L | Uncertain_significance |
| D | . | T | N | M | Uncertain_significance |
| D | D | T | D | L | Uncertain_significance |
| D | . | D | N | H | Uncertain_significance |
| P | . | T | N | M | Uncertain_significance |
| P | D | T | . | N | Uncertain_significance |
| . | . | . | D | . | Uncertain_significance |
| P | D | T | D | L | . |
| D | D | T | D | H | Uncertain_significance |
| D | D | T | D | L | Uncertain_significance |
| D | N | T | D | H | Uncertain_significance |
| D | N | T | D | H | Uncertain_significance |
| D | D | D | D | L | . |
| D | D | T | D | M | Uncertain_significance |
| P | D | T | D | N | Uncertain_significance |
| . | . | . | . | . | Uncertain_significance |
| . | . | . | . | . | Uncertain_significance |
| P | D | T | D | H | Uncertain_significance |
| P | N | . | P | L | Uncertain_significance |
| P | D | T | D | M | Uncertain_significance |
| D | D | T | D | M | Uncertain_significance |
| D | D | D | D | M | Uncertain_significance |
| D | D | D | D | H | Uncertain_significance |
| . | . | . | . | . | Uncertain_significance |

| | | | | | |
|---|---|---|---|---|---|
| B | . | D | N | M | Uncertain_significance |
| . | . | . | . | . | Uncertain_significance |
| . | . | . | . | . | Uncertain_significance |
| P | U | D | D | M | Uncertain_significance |
| D | . | . | . | . | Uncertain_significance |
| D | . | T | N | M | Uncertain_significance |
| D | D | T | D | H | Uncertain_significance |
| P | D | T | D | M | Uncertain_significance |
| P | D | D | D | L | Uncertain_significance |
| . | . | . | . | . | Uncertain_significance |
| . | . | . | D | . | Uncertain_significance |
| . | N | . | A | . | Pathogenic |

#HS-Sch-2 cell line variants with a potential impact on protein function

| chr | start | end | Sample | Gene.refGene | REF | ALT | Func.refGene | ExonicFunc.refGene | VAF |
|---|---|---|---|---|---|---|---|---|---|
| chr1 | 942460 | 942460 | HS-Sch-2 | SAMD11 | G | A | exonic | nonsynonymous_SNV | 0.772727272 |
| chr1 | 2609364 | 2609364 | HS-Sch-2 | MMEL1 | C | CG | exonic | frameshift_insertion | 0.813953488 |
| chr1 | 24076186 | 24076186 | HS-Sch-2 | MYOM3 | C | A | exonic | nonsynonymous_SNV | 0.736111111 |
| chr1 | 26281107 | 26281108 | HS-Sch-2 | SH3BGRL3 | TC | T | exonic | stopgain | 0.746835443 |
| chr1 | 32964588 | 32964588 | HS-Sch-2 | RNF19B | A | G | exonic | nonsynonymous_SNV | 0.4 |
| chr1 | 45997719 | 45997726 | HS-Sch-2 | MAST2 | CACAGGTA | C | exonic | nonframeshift_deletion | 0.814814814 |
| chr1 | 53210792 | 53210792 | HS-Sch-2 | CPT2 | C | G | exonic | nonsynonymous_SNV | 0.807017543 |
| chr1 | 64178683 | 64178683 | HS-Sch-2 | ROR1 | A | C | exonic | nonsynonymous_SNV | 0.753846153 |
| chr1 | 120460621 | 120460621 | HS-Sch-2 | NBPF8 | C | A | exonic | nonsynonymous_SNV | 1 |
| chr1 | 145873487 | 145873487 | HS-Sch-2 | ANKRD35 | G | A | exonic | nonsynonymous_SNV | 1 |
| chr1 | 146067225 | 146067225 | HS-Sch-2 | NBPF10 | C | A | exonic | nonsynonymous_SNV | 1 |
| chr1 | 146067283 | 146067283 | HS-Sch-2 | NBPF10 | C | T | exonic | nonsynonymous_SNV | 1 |
| chr1 | 148557528 | 148557528 | HS-Sch-2 | NBPF14 | C | G | exonic | nonsynonymous_SNV | 1 |
| chr1 | 148559961 | 148559961 | HS-Sch-2 | NBPF14 | T | C | exonic | nonsynonymous_SNV | 1 |
| chr1 | 149062099 | 149062099 | HS-Sch-2 | NBPF9 | A | T | exonic | nonsynonymous_SNV | 1 |
| chr1 | 149071644 | 149071644 | HS-Sch-2 | NBPF9 | T | C | exonic | nonsynonymous_SNV | 1 |
| chr1 | 149390858 | 149390858 | HS-Sch-2 | NOTCH2NLC | G | T | exonic | nonsynonymous_SNV | 1 |
| chr1 | 149554508 | 149554508 | HS-Sch-2 | NBPF19 | C | G | exonic | nonsynonymous_SNV | 1 |
| chr1 | 149554661 | 149554661 | HS-Sch-2 | NBPF19 | T | G | exonic | nonsynonymous_SNV | 1 |
| chr1 | 151838529 | 151838529 | HS-Sch-2 | C2CD4D | G | T | exonic | nonsynonymous_SNV | 1 |
| chr1 | 156378043 | 156378043 | HS-Sch-2 | RHBG | T | G | exonic | nonsynonymous_SNV | 1 |
| chr1 | 228214420 | 228214420 | HS-Sch-2 | OBSCN | A | C | exonic | nonsynonymous_SNV | 1 |
| chr2 | 1648366 | 1648366 | HS-Sch-2 | PXDN | C | A | exonic | nonsynonymous_SNV | 0.393939393 |
| chr2 | 40429730 | 40429730 | HS-Sch-2 | SLC8A1 | G | T | exonic | nonsynonymous_SNV | 0.369230769 |
| chr2 | 54899725 | 54899725 | HS-Sch-2 | EML6 | C | T | exonic | nonsynonymous_SNV | 0.357142857 |
| chr2 | 63056255 | 63056255 | HS-Sch-2 | OTX1 | T | G | exonic | nonsynonymous_SNV | 0.3125 |
| chr2 | 73385903 | 73385909 | HS-Sch-2 | ALMS1 | TGGAGGA | TGGAGGAGG | exonic | nonframeshift_insertion | 1 |
| chr2 | 86766014 | 86766014 | HS-Sch-2 | RMND5A | C | G | exonic | nonsynonymous_SNV | 0.294117647 |
| chr2 | 95159238 | 95159238 | HS-Sch-2 | ZNF514 | A | AC | splicing | . | 675 |
| chr2 | 132320213 | 132320213 | HS-Sch-2 | ZNF806 | A | G | exonic | nonsynonymous_SNV | 0.370370370 |
| chr2 | 173198040 | 173198040 | HS-Sch-2 | MAP3K20 | G | C | exonic | nonsynonymous_SNV | 0.5 |
| chr2 | 178614556 | 178614556 | HS-Sch-2 | TTN | C | G | exonic | nonsynonymous_SNV | 0.567567567 |
| chr3 | 17238358 | 17238358 | HS-Sch-2 | TBC1D5 | C | T | exonic | nonsynonymous_SNV | 0.121212121 |
| chr3 | 38039608 | 38039608 | HS-Sch-2 | DLEC1 | A | G | exonic | nonsynonymous_SNV | 1 |

| | | | | | | | | | |
|---|---|---|---|---|---|---|---|---|---|
| chr3 | 109103857 | 109103857 | HS-Sch-2 | MORC1 | A | G | exonic | nonsynonymous_SNV | 0.309523809 |
| chr3 | 113604951 | 113604951 | HS-Sch-2 | SIDT1 | T | A | exonic | nonsynonymous_SNV | 0.733333333 |
| chr3 | 130992992 | 130992992 | HS-Sch-2 | ATP2C1 | A | C | exonic | nonsynonymous_SNV | 0.648648648 |
| chr3 | 130992996 | 130992996 | HS-Sch-2 | ATP2C1 | A | T | exonic | nonsynonymous_SNV | 0.315789473 |
| chr4 | 5990757 | 5990757 | HS-Sch-2 | C4orf50 | T | G | exonic | nonsynonymous_SNV | 1 |
| chr4 | 67681573 | 67681573 | HS-Sch-2 | UBA6 | G | T | exonic | nonsynonymous_SNV | 0.461538461 |
| chr4 | 73091539 | 73091539 | HS-Sch-2 | ANKRD17 | G | T | exonic | nonsynonymous_SNV | 0.558139534 |
| chr4 | 77949867 | 77949867 | HS-Sch-2 | MRPL1 | C | G | exonic | nonsynonymous_SNV | 0.5 |
| chr4 | 78870982 | 78870997 | HS-Sch-2 | BMP2K | ACAGCAGCA( | A | exonic | nonframeshift_deletion | 475 |
| chr4 | 104491793 | 104491793 | HS-Sch-2 | CXXC4 | T | A | exonic | nonsynonymous_SNV | 0.6 |
| chr4 | 112584170 | 112584170 | HS-Sch-2 | ZGRF1 | C | T | exonic | nonsynonymous_SNV | 0.540540540 |
| chr4 | 119319075 | 119319075 | HS-Sch-2 | FABP2 | C | T | exonic | nonsynonymous_SNV | 0.617647058 |
| chr4 | 161759504 | 161759504 | HS-Sch-2 | FSTL5 | C | A | exonic | nonsynonymous_SNV | 0.181818181 |
| chr4 | 183271242 | 183271242 | HS-Sch-2 | WWC2 | G | C | splicing | . | 0.333333333 |
| chr5 | 38412589 | 38412589 | HS-Sch-2 | EGFLAM | G | T | exonic | nonsynonymous_SNV | 0.253968253 |
| chr5 | 119150213 | 119150216 | HS-Sch-2 | DMXL1 | TGAA | T | exonic | nonframeshift_deletion | 1 |
| chr5 | 140552043 | 140552043 | HS-Sch-2 | SRA1 | A | AGTC | exonic | nonframeshift_insertion | 1 |
| chr5 | 141137592 | 141137592 | HS-Sch-2 | PCDHB5 | C | T | exonic | nonsynonymous_SNV | 0.454545454 |
| chr5 | 149217001 | 149217001 | HS-Sch-2 | ABLIM3 | C | T | exonic | nonsynonymous_SNV | 0.5 |
| chr6 | 30346533 | 30346533 | HS-Sch-2 | RPP21\x3bTRIM3 | G | A | exonic | nonsynonymous_SNV | 0.6 |
| chr6 | 132538470 | 132538470 | HS-Sch-2 | TAAR9 | A | T | exonic | stopgain | 1 |
| chr7 | 2513247 | 2513251 | HS-Sch-2 | LFNG | AGATG | AGATGGATG | exonic | frameshift_insertion | 1 |
| chr7 | 31847977 | 31847977 | HS-Sch-2 | PDE1C | T | C | exonic | nonsynonymous_SNV | 0.578947368 |
| chr7 | 99715886 | 99715886 | HS-Sch-2 | CYP3A7\x3bCYP3. | A | C | exonic | nonsynonymous_SNV | 0.416666666 |
| chr7 | 107580767 | 107580767 | HS-Sch-2 | DUS4L-BCAP29 | A | G | exonic | nonsynonymous_SNV | 0.717391304 |
| chr7 | 128866780 | 128866780 | HS-Sch-2 | ATP6V1FNB | C | T | exonic | nonsynonymous_SNV | 0.720930232 |
| chr7 | 130547646 | 130547646 | HS-Sch-2 | COPG2 | G | A | exonic | nonsynonymous_SNV | 0.245614035 |
| chr7 | 131556270 | 131556276 | HS-Sch-2 | PODXL | GGGCGAC | GGGCGACGG | exonic | nonframeshift_insertion | 1 |
| chr7 | 152158868 | 152158868 | HS-Sch-2 | KMT2C | T | G | exonic | nonsynonymous_SNV | 0.666666666 |
| chr8 | 6825251 | 6825251 | HS-Sch-2 | XKR5 | A | C | exonic | nonsynonymous_SNV | 1 |
| chr8 | 10610079 | 10610079 | HS-Sch-2 | RP1L1 | T | TCCTCTAACT( | exonic | nonframeshift_insertion | 1 |
| chr8 | 25503476 | 25503476 | HS-Sch-2 | CDCA2 | C | T | exonic | nonsynonymous_SNV | 1 |
| chr8 | 51475061 | 51475061 | HS-Sch-2 | PXDNL | C | A | exonic | nonsynonymous_SNV | 0.404761904 |
| chr8 | 70734336 | 70734337 | HS-Sch-2 | XKR9 | CA | C | exonic | frameshift_deletion | 0.416666666 |
| chr8 | 125008998 | 125008998 | HS-Sch-2 | SQLE | G | A | exonic | nonsynonymous_SNV | 125 |
| chr8 | 138812959 | 138812959 | HS-Sch-2 | COL22A1 | A | C | exonic | nonsynonymous_SNV | 0.166666666 |

| | | | | | | | | | |
|---|---|---|---|---|---|---|---|---|---|
| chr8 | 143859527 | 143859527 | HS-Sch-2 | EPPK1 | A | G | exonic | nonsynonymous_SNV | 1 |
| chr9 | 35906589 | 35906589 | HS-Sch-2 | HRCT1 | A | ACCACCACCA | exonic | nonframeshift_insertion | 1 |
| chr9 | 37429750 | 37429750 | HS-Sch-2 | GRHPR | G | A | exonic | nonsynonymous_SNV | 1 |
| chr9 | 41964577 | 41964577 | HS-Sch-2 | CNTNAP3B | A | C | exonic | nonsynonymous_SNV | 0.620689655 |
| chr9 | 41991721 | 41991721 | HS-Sch-2 | CNTNAP3B | G | A | exonic | nonsynonymous_SNV | 1 |
| chr10 | 25572707 | 25572707 | HS-Sch-2 | GPR158 | C | T | exonic | nonsynonymous_SNV | 1 |
| chr10 | 27735376 | 27735376 | HS-Sch-2 | MKX | T | G | splicing | . | 0.230769230 |
| chr10 | 47502343 | 47502343 | HS-Sch-2 | AGAP9 | T | C | exonic | nonsynonymous_SNV | 1 |
| chr10 | 47502604 | 47502604 | HS-Sch-2 | AGAP9 | A | G | exonic | nonsynonymous_SNV | 1 |
| chr10 | 47502991 | 47502991 | HS-Sch-2 | AGAP9 | C | T | exonic | nonsynonymous_SNV | 1 |
| chr10 | 66927883 | 66927883 | HS-Sch-2 | LRRTM3 | C | A | exonic | nonsynonymous_SNV | 0.452380952 |
| chr10 | 68459198 | 68459203 | HS-Sch-2 | DNA2 | CTTGTT | C | exonic | frameshift_deletion | 0.421052631 |
| chr10 | 72063097 | 72063097 | HS-Sch-2 | SPOCK2 | C | T | exonic | nonsynonymous_SNV | 0.4 |
| chr10 | 88817855 | 88817855 | HS-Sch-2 | LIPM | G | T | exonic | nonsynonymous_SNV | 0.296296296 |
| chr10 | 102599534 | 102599534 | HS-Sch-2 | SUFU | G | C | exonic | nonsynonymous_SNV | 0.509433962 |
| chr11 | 3828091 | 3828091 | HS-Sch-2 | RHOG | C | G | exonic | nonsynonymous_SNV | 0.220588235 |
| chr11 | 4803675 | 4803675 | HS-Sch-2 | OR52R1 | C | G | exonic | nonsynonymous_SNV | 0.461538461 |
| chr11 | 20160272 | 20160272 | HS-Sch-2 | DBX1 | C | A | exonic | nonsynonymous_SNV | 0.529411764 |
| chr11 | 20508340 | 20508340 | HS-Sch-2 | PRMT3 | A | G | exonic | nonsynonymous_SNV | 0.75 |
| chr11 | 47642396 | 47642396 | HS-Sch-2 | MTCH2 | C | T | exonic | nonsynonymous_SNV | 0.227272727 |
| chr11 | 47642399 | 47642399 | HS-Sch-2 | MTCH2 | A | G | exonic | nonsynonymous_SNV | 0.227272727 |
| chr11 | 47642407 | 47642407 | HS-Sch-2 | MTCH2 | G | C | exonic | nonsynonymous_SNV | 0.227272727 |
| chr11 | 47642411 | 47642411 | HS-Sch-2 | MTCH2 | G | A | exonic | stopgain | 0.227272727 |
| chr11 | 62529786 | 62529786 | HS-Sch-2 | AHNAK | G | C | exonic | nonsynonymous_SNV | 0.414634146 |
| chr11 | 111354478 | 111354478 | HS-Sch-2 | POU2AF1 | G | A | exonic | nonsynonymous_SNV | 0.166666666 |
| chr12 | 6936728 | 6936740 | HS-Sch-2 | ATN1 | ACAGCAGCA | ACAGCAGCA | exonic | nonframeshift_insertion | 1 |
| chr12 | 6936728 | 6936740 | HS-Sch-2 | ATN1 | ACAGCAGCA | A | exonic | nonframeshift_deletion | 1 |
| chr12 | 12822055 | 12822055 | HS-Sch-2 | DDX47 | G | A | exonic | nonsynonymous_SNV | 0.45 |
| chr12 | 25108825 | 25108825 | HS-Sch-2 | CASC1 | T | TAAAAAAAA | splicing | . | 1 |
| chr12 | 31104017 | 31104017 | HS-Sch-2 | DDX11 | G | A | exonic | stopgain | 0.314285714 |
| chr12 | 43376527 | 43376527 | HS-Sch-2 | ADAMTS20 | C | A | exonic | nonsynonymous_SNV | 375 |
| chr12 | 50727721 | 50727721 | HS-Sch-2 | DIP2B | G | T | exonic | nonsynonymous_SNV | 0.702380952 |
| chr12 | 65171059 | 65171059 | HS-Sch-2 | LEMD3 | T | C | exonic | nonsynonymous_SNV | 0.545454545 |
| chr12 | 115991155 | 115991155 | HS-Sch-2 | MED13L | C | G | exonic | nonsynonymous_SNV | 0.608695652 |
| chr13 | 51797843 | 51797843 | HS-Sch-2 | DHRS12 | G | A | exonic | nonsynonymous_SNV | 1 |
| chr13 | 102748379 | 102748379 | HS-Sch-2 | CCDC168 | G | T | exonic | nonsynonymous_SNV | 1 |

| | | | | | | | | | |
|---|---|---|---|---|---|---|---|---|---|
| chr15 | 22613584 | 22613584 | HS-Sch-2 | GOLGA8J | A | G | exonic | nonsynonymous_SNV | 1 |
| chr15 | 31435525 | 31435525 | HS-Sch-2 | KLF13 | A | G | exonic | nonsynonymous_SNV | 0.487179487 |
| chr15 | 73122735 | 73122735 | HS-Sch-2 | NEO1 | G | A | exonic | nonsynonymous_SNV | 1 |
| chr15 | 79025423 | 79025423 | HS-Sch-2 | RASGRF1 | C | T | exonic | nonsynonymous_SNV | 0.354166666 |
| chr15 | 85244770 | 85244770 | HS-Sch-2 | GOLGA6L3 | G | T | exonic | nonsynonymous_SNV | 625 |
| chr15 | 99712504 | 99712525 | HS-Sch-2 | MEF2A | CCAGCAGCA( | C | exonic | nonframeshift_deletion | 0.531914893 |
| chr15 | 101068804 | 101068804 | HS-Sch-2 | LRRK1 | G | A | exonic | nonsynonymous_SNV | 0.35 |
| chr16 | 985108 | 985108 | HS-Sch-2 | SOX8 | T | C | exonic | nonsynonymous_SNV | 0.133333333 |
| chr16 | 20322878 | 20322878 | HS-Sch-2 | GP2 | T | C | exonic | nonsynonymous_SNV | 0.633333333 |
| chr16 | 31464750 | 31464750 | HS-Sch-2 | ARMC5 | A | G | exonic | nonsynonymous_SNV | 0.378378378 |
| chr16 | 82071040 | 82071040 | HS-Sch-2 | HSD17B2 | T | G | exonic | nonsynonymous_SNV | 0.456521739 |
| chr17 | 6090444 | 6090444 | HS-Sch-2 | WSCD1 | C | CG | exonic | frameshift_insertion | 1 |
| chr17 | 7428940 | 7428940 | HS-Sch-2 | SPEM3 | C | G | exonic | nonsynonymous_SNV | 0.485714285 |
| chr17 | 7673802 | 7673802 | HS-Sch-2 | TP53 | C | T | exonic | nonsynonymous_SNV | 1 |
| chr17 | 8143358 | 8143358 | HS-Sch-2 | PER1 | G | A | exonic | nonsynonymous_SNV | 1 |
| chr17 | 12944131 | 12944131 | HS-Sch-2 | ARHGAP44 | C | G | exonic | nonsynonymous_SNV | 0.153846153 |
| chr17 | 17793779 | 17793785 | HS-Sch-2 | RAI1 | CCAGCAG | CCAG | exonic | nonframeshift_deletion | 1 |
| chr17 | 27765627 | 27765627 | HS-Sch-2 | NOS2 | T | A | exonic | nonsynonymous_SNV | 0.4 |
| chr17 | 31159074 | 31159102 | HS-Sch-2 | NF1 | TGGAAAAAT( | T | exonic | frameshift_deletion | 0.5 |
| chr17 | 31230383 | 31230383 | HS-Sch-2 | NF1 | G | A | splicing | . | 0.444444444 |
| chr17 | 36001422 | 36001422 | HS-Sch-2 | CCL15 | G | A | exonic | nonsynonymous_SNV | 0.655172413 |
| chr17 | 36013244 | 36013244 | HS-Sch-2 | CCL23 | T | C | exonic | nonsynonymous_SNV | 0.66 |
| chr17 | 36451697 | 36451697 | HS-Sch-2 | TBC1D3D\x3bTBC | C | A | exonic | nonsynonymous_SNV | 1 |
| chr17 | 36455955 | 36455955 | HS-Sch-2 | TBC1D3D\x3bTBC | T | C | exonic | nonsynonymous_SNV | 1 |
| chr17 | 37411381 | 37411381 | HS-Sch-2 | TADA2A | T | C | exonic | nonsynonymous_SNV | 0.487804878 |
| chr17 | 37624364 | 37624364 | HS-Sch-2 | DDX52 | C | T | exonic | nonsynonymous_SNV | 0.719298245 |
| chr17 | 39402308 | 39402308 | HS-Sch-2 | FBXL20 | T | TCCCCCGCCC | exonic | nonframeshift_insertior | 0.407407407 |
| chr17 | 48726906 | 48726906 | HS-Sch-2 | HOXB13 | G | T | exonic | nonsynonymous_SNV | 0.269230769 |
| chr17 | 48728199 | 48728199 | HS-Sch-2 | HOXB13 | C | T | exonic | nonsynonymous_SNV | 0.604166666 |
| chr17 | 62711335 | 62711335 | HS-Sch-2 | MARCHF10 | C | T | exonic | nonsynonymous_SNV | 0.637931034 |
| chr17 | 63941345 | 63941345 | HS-Sch-2 | SCN4A | G | T | exonic | nonsynonymous_SNV | 0.454545454 |
| chr17 | 74253250 | 74253250 | HS-Sch-2 | TTYH2 | C | A | exonic | nonsynonymous_SNV | 0.666666666 |
| chr18 | 13642733 | 13642734 | HS-Sch-2 | LDLRAD4 | TC | T | exonic | frameshift_deletion | 0.447058823 |
| chr19 | 47258628 | 47258628 | HS-Sch-2 | CCDC9 | C | T | exonic | nonsynonymous_SNV | 0.419354838 |
| chr19 | 54096945 | 54096945 | HS-Sch-2 | OSCAR | C | A | exonic | nonsynonymous_SNV | 0.230769230 |
| chr19 | 54832784 | 54832784 | HS-Sch-2 | KIR2DS4 | T | G | exonic | nonsynonymous_SNV | 0.571428571 |

| | | | | | | | | | |
|---|---|---|---|---|---|---|---|---|---|
| chr19 | 54835079 | 54835079 | HS-Sch-2 | KIR2DS4 | G | C | exonic | nonsynonymous_SNV | 0.461538461 |
| chr19 | 54835110 | 54835110 | HS-Sch-2 | KIR2DS4 | G | T | exonic | nonsynonymous_SNV | 0.580645161 |
| chr19 | 54837865 | 54837865 | HS-Sch-2 | KIR2DS4 | G | C | exonic | nonsynonymous_SNV | 0.269230769 |
| chr19 | 54837873 | 54837873 | HS-Sch-2 | KIR2DS4 | T | C | exonic | nonsynonymous_SNV | 0.269230769 |
| chr21 | 34716472 | 34716472 | HS-Sch-2 | CLIC6 | G | C | exonic | nonsynonymous_SNV | 0.319148936 |
| chr21 | 44600952 | 44600952 | HS-Sch-2 | KRTAP10-7 | G | A | exonic | nonsynonymous_SNV | 0.347826086 |
| chr21 | 44601264 | 44601264 | HS-Sch-2 | KRTAP10-7 | A | C | exonic | nonsynonymous_SNV | 0.647058823 |
| chr21 | 44601475 | 44601475 | HS-Sch-2 | KRTAP10-7 | C | G | exonic | nonsynonymous_SNV | 0.333333333 |
| chr21 | 44601579 | 44601579 | HS-Sch-2 | KRTAP10-7 | G | A | exonic | nonsynonymous_SNV | 0.523809523 |
| chr22 | 22515221 | 22515221 | HS-Sch-2 | ZNF280A | G | T | exonic | nonsynonymous_SNV | 0.621212121 |
| chr22 | 22515224 | 22515224 | HS-Sch-2 | ZNF280A | C | T | exonic | nonsynonymous_SNV | 0.621212121 |
| chr22 | 23998589 | 23998589 | HS-Sch-2 | GSTT4 | A | G | exonic | nonsynonymous_SNV | 0.305555555 |
| chr22 | 25029315 | 25029315 | HS-Sch-2 | KIAA1671 | A | G | exonic | nonsynonymous_SNV | 0.976744186 |
| chr22 | 29043333 | 29043333 | HS-Sch-2 | ZNRF3 | C | T | exonic | nonsynonymous_SNV | 0.440677966 |
| chr22 | 29328884 | 29328884 | HS-Sch-2 | AP1B1 | C | G | exonic | nonsynonymous_SNV | 0.220588235 |
| chr22 | 41972169 | 41972169 | HS-Sch-2 | SEPTIN3 | C | A | exonic | nonsynonymous_SNV | 0.350649350 |
| chrX | 14690744 | 14690744 | HS-Sch-2 | GLRA2 | C | T | exonic | nonsynonymous_SNV | 0.142857142 |
| chrX | 153182483 | 153182483 | HS-Sch-2 | MAGEA1 | A | G | exonic | nonsynonymous_SNV | 1 |
| chrX | 153182604 | 153182604 | HS-Sch-2 | MAGEA1 | G | A | exonic | nonsynonymous_SNV | 1 |
| chrX | 154653251 | 154653251 | HS-Sch-2 | CTAG2 | C | G | exonic | nonsynonymous_SNV | 1 |

| avsnp150 | Variant | AAChange.refGene | GeneDetail.refGene | AF | AF_popmax | prop_pathogenic_pred | CLNALLELEID |
|---|---|---|---|---|---|---|---|
| . | chr1:942460_ | SAMD11:NM_152486:e | . | . | . | 1 | . |
| . | chr1:2609364 | MMEL1:NM_033467:e | . | . | . | NA | . |
| . | chr1:2407618 | MYOM3:NM_152372:e | . | . | . | 0.857142857142857 | . |
| . | chr1:2628110 | SH3BGRL3:NM_031286 | . | . | . | NA | . |
| . | chr1:3296458 | RNF19B:NM_0011273€ | . | . | . | 0.666666666666667 | . |
| . | chr1:4599771 | MAST2:NM_00132432 | . | . | . | NA | . |
| . | chr1:5321079 | CPT2:NM_000098:exon | . | . | . | 1 | . |
| . | chr1:6417868 | ROR1:NM_005012:exo | . | . | . | 0.714285714285714 | . |
| rs587696045 | chr1:1204600 | NBPF8:NM_00103750 | . | . | . | NA | . |
| rs6670984 | chr1:1458734 | ANKRD35:NM_001280 | . | . | . | NA | . |
| . | chr1:1460672 | NBPF10:NM_0010397( | . | . | . | NA | . |
| . | chr1:1460672 | NBPF10:NM_0010397( | . | . | . | NA | . |
| . | chr1:1485575 | NBPF14:NM_015383:e | . | . | . | NA | . |
| . | chr1:1485599 | NBPF14:NM_015383:e | . | . | . | NA | . |
| . | chr1:1490620 | NBPF9:NM_00103767 | . | . | . | NA | . |
| . | chr1:1490710 | NBPF9:NM_00103767 | . | . | . | NA | . |
| . | chr1:1493908 | NOTCH2NLC:NM_0013 | . | . | . | NA | . |
| . | chr1:1495545 | NBPF19:NM_0013513 | . | . | . | 0.75 | . |
| . | chr1:1495546 | NBPF19:NM_0013513 | . | . | . | NA | . |
| . | chr1:1518385 | C2CD4D:NM_0011360 | . | . | . | 0.666666666666667 | . |
| . | chr1:1563780 | RHBG:NM_020407:exc | . | . | . | 0.8 | . |
| rs1771487 | chr1:2282144 | OBSCN:NM_00109862 | . | . | . | 0.714285714285714 | . |
| . | chr2:1648360 | PXDN:NM_012293:exc | . | . | . | 0.714285714285714 | . |
| . | chr2:40429730_G/T | | . | . | . | 0.857142857142857 | . |
| . | chr2:5489972 | EML6:NM_001039753 | . | . | . | 0.714285714285714 | . |
| . | chr2:6305625 | OTX1:NM_001199770: | . | . | . | 0.857142857142857 | . |
| . | chr2:7338590 | ALMS1:NM_015120:ex | . | . | . | NA | . |
| . | chr2:8676601 | RMND5A:NM_022780: | . | . | . | 1 | . |
| . | chr2:9515923 | . | NM_032788:exon1:UTR | . | . | NA | . |
| rs12468523 | chr2:1323202 | ZNF806:NM_0013554€ | . | . | . | NA | . |
| . | chr2:1731980 | MAP3K20:NM_016653 | . | . | . | 0.428571428571429 | . |
| . | chr2:1786145 | TTN:NM_003319:exon | . | . | . | 0.666666666666667 | . |
| . | chr3:17238358_C/T | | . | . | . | 0.857142857142857 | . |
| . | chr3:3803960 | DLEC1:NM_00132115 | . | . | . | 0.857142857142857 | . |

| | | | | | | | |
|---|---|---|---|---|---|---|---|
| . | chr3:1091038 | MORC1:NM_014429:e | . | . | . | 0.857142857142857 | . |
| rs3732795 | chr3:113604951_T/A | | . | 0.0003 | 0.0051 | 0.714285714285714 | . |
| . | chr3:130992992_A/C | | . | . | . | 0.714285714285714 | . |
| . | chr3:130992996_A/T | | . | . | . | 1 | . |
| . | chr4:5990757 | C4orf50:NM_0013646 | . | . | . | NA | . |
| . | chr4:6768157 | UBA6:NM_018227:exo | . | . | . | 0.857142857142857 | . |
| . | chr4:7309153 | ANKRD17:NM_198889 | . | . | . | 0.714285714285714 | . |
| rs137874988 | chr4:7794986 | MRPL1:NM_020236:ex | . | 0.0004 | 0.0071 | 0.857142857142857 | . |
| . | chr4:7887098 | BMP2K:NM_017593:e | . | . | . | NA | . |
| . | chr4:1044917 | CXXC4:NM_025212:ex | . | . | . | 1 | . |
| rs542798884 | chr4:1125847 | ZGRF1:NM_00135039 | . | 6.377e-05 | 0.0012 | 0.714285714285714 | . |
| . | chr4:1193190 | FABP2:NM_000134:ex | . | . | . | 0.714285714285714 | . |
| . | chr4:1617595 | FSTL5:NM_001128427 | . | . | . | 0.714285714285714 | . |
| . | chr4:1832712 | . | NM_024949:exon16:c.2 | . | . | 1 | . |
| . | chr5:3841258 | EGFLAM:NM_182798:e | . | . | . | 0.714285714285714 | . |
| . | chr5:119150213_TGAA/T | | . | . | . | NA | . |
| . | chr5:1405520 | SRA1:NM_001035235: | . | . | . | NA | . |
| rs400562 | chr5:1411375 | PCDHB5:NM_015669:e | . | . | . | NA | . |
| rs767983995 | chr5:149217001_C/T | | . | . | . | 1 | . |
| . | chr6:3034653 | RPP21:NM_00119912 | . | . | . | 0.714285714285714 | . |
| rs2842899 | chr6:1325384 | TAAR9:NM_175057:ex | . | . | . | NA | . |
| . | chr7:2513247 | LFNG:NM_001166355: | . | . | . | NA | . |
| . | chr7:31847977_T/C | | . | . | . | 0.857142857142857 | . |
| . | chr7:9971588 | CYP3A7:NM_000765:e | . | . | . | 0.857142857142857 | . |
| . | chr7:1075807 | DUS4L-BCAP29:NM_00 | . | . | . | NA | . |
| rs181284272 | chr7:1288667 | ATP6V1FNB:NM_0011 | . | 0.0005 | 9 | NA | . |
| . | chr7:1305476 | COPG2:NM_00129003 | . | . | . | NA | . |
| . | chr7:1315562 | PODXL:NM_00101811 | . | . | . | NA | . |
| rs761464984 | chr7:1521588 | KMT2C:NM_170606:ex | . | . | . | 0.714285714285714 | . |
| . | chr8:6825251 | XKR5:NM_207411:exo | . | . | . | 1 | . |
| . | chr8:10610079_T/TCCTCTAACTGCA | | . | . | . | NA | . |
| . | chr8:2550347 | CDCA2:NM_00131790 | . | . | . | 0.857142857142857 | . |
| . | chr8:5147506 | PXDNL:NM_144651:ex | . | . | . | 0.833333333333333 | . |
| . | chr8:7073433 | XKR9:NM_001011720: | . | . | . | NA | . |
| . | chr8:1250089 | SQLE:NM_003129:exo | . | . | . | 0.857142857142857 | . |
| . | chr8:1388129 | COL22A1:NM_152888: | . | . | . | 0.714285714285714 | . |

| | | | | | | | | |
|---|---|---|---|---|---|---|---|---|
| . | chr8:143859 | EPPK1:NM_031308:ex | . | . | . | NA | . | |
| . | chr9:359065 | HRCT1:NM_00103979 | . | . | . | NA | . | |
| rs200106110 | chr9:374297 | GRHPR:NM_012203:e | . | 0.0019 | 0.0064 | 1 | | 319215 |
| rs76032838 | chr9:419645 | CNTNAP3B:NM_00120 | . | . | . | NA | . | |
| rs3739623 | chr9:419917 | CNTNAP3B:NM_00120 | . | . | . | NA | . | |
| rs143713985 | chr10:25572 | GPR158:NM_020752:e | . | 0.0008 | 0.0084 | 0.857142857142857 | . | |
| . | chr10:27735 | . | NM_173576:exon4:c.3 | . | . | 1 | . | |
| rs1047447 | chr10:47502 | AGAP9:NM_00119081 | . | . | . | NA | . | |
| rs4013543 | chr10:47502 | AGAP9:NM_00119081 | . | . | . | NA | . | |
| rs4013536 | chr10:47502 | AGAP9:NM_00119081 | . | . | . | NA | . | |
| . | chr10:66927 | LRRTM3:NM_178011: | . | . | . | 0.857142857142857 | . | |
| . | chr10:68459 | DNA2:NM_001080449 | . | . | . | NA | . | |
| rs2306322 | chr10:72063 | SPOCK2:NM_0012449 | . | 0.0029 | 9 | 0.857142857142857 | . | |
| . | chr10:88817 | LIPM:NM_001128215: | . | . | . | 0.857142857142857 | . | |
| . | chr10:10259 | SUFU:NM_001178133: | . | . | . | 0.714285714285714 | . | |
| . | chr11:38280 | RHOG:NM_001665:ex | . | . | . | 1 | . | |
| . | chr11:48036 | OR52R1:NM_0010051 | . | . | . | 0.714285714285714 | . | |
| . | chr11:20160 | DBX1:NM_001029865: | . | . | . | 1 | . | |
| rs6483700 | chr11:20508 | PRMT3:NM_00114516 | . | . | . | NA | . | |
| rs186601647 | chr11:47642 | MTCH2:NM_00131723 | . | . | . | 1 | . | |
| rs200816454 | chr11:47642 | MTCH2:NM_00131723 | . | . | . | 0.857142857142857 | . | |
| rs201627714 | chr11:47642 | MTCH2:NM_00131723 | . | . | . | 1 | . | |
| rs76185606 | chr11:47642 | MTCH2:NM_00131723 | . | . | . | 1 | . | |
| rs139574230 | chr11:62529 | AHNAK:NM_00134644 | . | . | . | 0.857142857142857 | . | |
| . | chr11:11135 | POU2AF1:NM_006235 | . | . | . | 0.714285714285714 | . | |
| . | chr12:69367 | ATN1:NM_001007026: | . | . | . | NA | . | |
| . | chr12:69367 | ATN1:NM_001007026: | . | . | . | NA | . | |
| rs199573191 | chr12:12822 | DDX47:NM_016355:ex | . | . | . | 0.714285714285714 | . | |
| . | chr12:25108 | . | | . | . | NA | . | |
| . | chr12:31104 | DDX11:NM_00125714 | . | . | . | 0 | . | |
| . | chr12:43376 | ADAMTS20:NM_0250 | . | . | . | 0.714285714285714 | . | |
| . | chr12:50727 | DIP2B:NM_173602:ex | . | . | . | 0.714285714285714 | . | |
| . | chr12:65171 | LEMD3:NM_00116761 | . | . | . | 0.857142857142857 | . | |
| . | chr12:11599 | MED13L:NM_015335: | . | . | . | 0.714285714285714 | . | |
| rs149418560 | chr13:51797 | DHRS12:NM_0012704 | . | 0.0004 | 0.0064 | 0.666666666666667 | . | |
| . | chr13:10274 | CCDC168:NM_001146 | . | . | . | NA | . | |

| | | | | | | | | |
|---|---|---|---|---|---|---|---|---|
| . | chr15:22613 | GOLGA8J:NM_00128 | . | . | . | NA | . | |
| . | chr15:31435 | KLF13:NM_001302461 | . | . | . | NA | . | |
| rs183676997 | chr15:73122 | NEO1:NM_001172623 | . | . | . | 0.857142857142857 | . | |
| . | chr15:79025 | RASGRF1:NM_001145 | . | . | . | 0.857142857142857 | . | |
| . | chr15:85244 | GOLGA6L3:NM_00131 | . | . | . | NA | . | |
| . | chr15:99712504_CCAGCAGCAGCAG | | . | . | . | NA | . | |
| rs3752321 | chr15:10106 | LRRK1:NM_024652:ex | . | 0.0003 | 0.0064 | 0.857142857142857 | . | |
| . | chr16:98510 | SOX8:NM_014587:exo | . | . | . | 1 | . | |
| . | chr16:20322 | GP2:NM_001007242:e | . | . | . | 0.833333333333333 | . | |
| . | chr16:31464 | ARMC5:NM_00110524 | . | . | . | 0.714285714285714 | . | |
| . | chr16:82071 | HSD17B2:NM_002153 | . | . | . | 0.857142857142857 | . | |
| . | chr17:60904 | WSCD1:NM_015253:e | . | . | . | NA | . | |
| . | chr17:74289 | SPEM3:NM_00136467 | . | . | . | NA | . | |
| rs28934576 | chr17:7673802_C/T | | . | . | . | 0.857142857142857 | | 27405 |
| rs777117943 | chr17:81433 | PER1:NM_002616:exo | . | . | . | 0.714285714285714 | . | |
| rs762885019 | chr17:12944 | ARHGAP44:NM_00132 | . | . | . | 0.857142857142857 | . | |
| . | chr17:17793 | RAI1:NM_030665:exo | . | . | . | NA | . | |
| . | chr17:27765 | NOS2:NM_000625:exo | . | . | . | 1 | . | |
| . | chr17:31159 | NF1:NM_000267:exon | . | . | . | NA | . | |
| rs267606599 | chr17:31230 | . | NM_000267:exon23:c.3 | . | . | 1 | | 15384 |
| rs854625 | chr17:36001 | CCL15:NM_032965:ex | . | . | . | NA | . | |
| rs1003645 | chr17:36013 | CCL23:NM_005064:ex | . | . | . | NA | . | |
| . | chr17:36451 | TBC1D3J:NM_0013555 | . | . | . | NA | . | |
| . | chr17:36455 | TBC1D3J:NM_0013555 | . | . | . | NA | . | |
| rs7211875 | chr17:37411 | TADA2A:NM_0011661 | . | . | . | NA | . | |
| rs7216445 | chr17:37624 | DDX52:NM_007010:ex | . | . | . | NA | . | |
| . | chr17:39402 | FBXL20:NM_00137020 | . | . | . | NA | . | |
| . | chr17:48726 | HOXB13:NM_006361: | . | . | . | 0.857142857142857 | . | |
| . | chr17:48728 | HOXB13:NM_006361: | . | . | . | 1 | . | |
| rs16946335 | chr17:62711 | MARCHF10:NM_0011 | . | 0.0006 | 9 | 0.857142857142857 | . | |
| . | chr17:63941 | SCN4A:NM_000334:ex | . | . | . | 0.857142857142857 | . | |
| rs140920850 | chr17:74253 | TTYH2:NM_052869:ex | . | 0.0004 | 0.0071 | 0.857142857142857 | . | |
| . | chr18:13642 | LDLRAD4:NM_001276 | . | . | . | NA | . | |
| . | chr19:47258 | CCDC9:NM_015603:ex | . | . | . | 0.857142857142857 | . | |
| rs1657535 | chr19:54096 | OSCAR:NM_00128235 | . | . | . | NA | . | |
| rs1130476 | chr19:54832 | KIR2DS4:NM_0012819 | . | . | . | NA | . | |

| | | | | | | | |
|---|---|---|---|---|---|---|---|
| rs112522228 | chr19:54835( | KIR2DS4:NM_0012819 | . | . | . | NA | . |
| rs1130478 | chr19:54835: | KIR2DS4:NM_0012819 | . | . | . | NA | . |
| rs1130494 | chr19:54837{ | KIR2DS4:NM_0012819 | . | . | . | NA | . |
| rs201518955 | chr19:54837{ | KIR2DS4:NM_0012819 | . | . | . | NA | . |
| . | chr21:34716 | CLIC6:NM_053277:exo | . | . | . | 1 | . |
| rs944419 | chr21:44600 | KRTAP10-7:NM_19868 | . | . | . | NA | . |
| rs363877 | chr21:44601 | KRTAP10-7:NM_19868 | . | . | . | NA | . |
| rs446817 | chr21:44601 | KRTAP10-7:NM_19868 | . | . | . | NA | . |
| rs369720 | chr21:44601 | KRTAP10-7:NM_19868 | . | . | . | NA | . |
| rs361580 | chr22:22515 | ZNF280A:NM_080740: | . | . | . | NA | . |
| rs362011 | chr22:22515 | ZNF280A:NM_080740: | . | . | . | NA | . |
| . | chr22:23998 | GSTT4:NM_00135866 | . | . | . | NA | . |
| rs17667531 | chr22:25029 | KIAA1671:NM_001145 | . | . | . | 0.666666666666667 | . |
| . | chr22:29043 | ZNRF3:NM_00120699 | . | . | . | 0.714285714285714 | . |
| . | chr22:29328 | AP1B1:NM_00116601 | . | . | . | 0.857142857142857 | . |
| . | chr22:41972 | SEPTIN3:NM_0013638 | . | . | . | NA | . |
| rs995774836 | chrX:146907 | GLRA2:NM_00111888 | . | . | . | 0.714285714285714 | . |
| rs2008160 | chrX:153182 | MAGEA1:NM_004988: | . | . | . | NA | . |
| rs2008144 | chrX:153182 | MAGEA1:NM_004988: | . | . | . | NA | . |
| rs17328091 | chrX:154653 | CTAG2:NM_020994:ex | . | . | . | NA | . |

| CLNDN | CLNDISDB | CLNREVSTAT | CLNSIG | SIFT_pred | Polyphen2_HDIV_pred | Polyphen2_HVAR_pred | LRT_pred |
|---|---|---|---|---|---|---|---|
| . | . | . | . | D | D | D | D |
| . | . | . | . | . | . | . | . |
| . | . | . | . | D | D | D | D |
| . | . | . | . | . | . | . | . |
| . | . | . | . | D | D | P | . |
| . | . | . | . | . | . | . | . |
| . | . | . | . | D | D | D | D |
| . | . | . | . | D | D | D | D |
| . | . | . | . | . | . | . | . |
| . | . | . | . | . | . | . | . |
| . | . | . | . | . | . | . | . |
| . | . | . | . | . | . | . | . |
| . | . | . | . | . | . | . | . |
| . | . | . | . | . | . | . | . |
| . | . | . | . | . | . | . | . |
| . | . | . | . | . | . | . | . |
| . | . | . | . | . | . | . | . |
| . | . | . | . | . | D | P | . |
| . | . | . | . | . | . | . | . |
| . | . | . | . | T | D | D | . |
| . | . | . | . | . | D | P | N |
| . | . | . | . | T | P | P | D |
| . | . | . | . | T | D | D | D |
| . | . | . | . | D | D | D | D |
| . | . | . | . | D | D | D | U |
| . | . | . | . | D | D | D | D |
| . | . | . | . | . | . | . | . |
| . | . | . | . | D | D | D | D |
| . | . | . | . | . | . | . | . |
| . | . | . | . | . | . | . | . |
| . | . | . | . | T | B | B | D |
| . | . | . | . | D | P | P | . |
| . | . | . | . | D | D | D | D |
| . | . | . | . | D | D | D | D |

| | | | | | | | |
|---|---|---|---|---|---|---|---|
| . | . | . | . | D | D | P | D |
| . | . | . | . | D | P | P | N |
| . | . | . | . | T | D | P | D |
| . | . | . | . | D | P | P | D |
| . | . | . | . | . | . | . | . |
| . | . | . | . | D | D | D | D |
| . | . | . | . | D | D | P | D |
| . | . | . | . | D | D | D | D |
| . | . | . | . | . | . | . | . |
| . | . | . | . | . | . | . | . |
| . | . | . | . | D | D | D | N |
| . | . | . | . | T | D | D | D |
| . | . | . | . | T | P | P | D |
| . | . | . | . | . | . | . | . |
| . | . | . | . | D | D | D | D |
| . | . | . | . | . | . | . | . |
| . | . | . | . | . | . | . | . |
| . | . | . | . | . | . | . | . |
| . | . | . | . | D | D | D | D |
| . | . | . | . | D | D | D | D |
| . | . | . | . | . | . | . | . |
| . | . | . | . | . | . | . | . |
| . | . | . | . | D | P | P | D |
| . | . | . | . | D | D | P | D |
| . | . | . | . | . | . | . | . |
| . | . | . | . | . | . | . | . |
| . | . | . | . | . | . | . | . |
| . | . | . | . | . | . | . | . |
| . | . | . | . | T | D | D | U |
| . | . | . | . | . | D | D | . |
| . | . | . | . | . | . | . | . |
| . | . | . | . | D | D | D | D |
| . | . | . | . | D | D | P | . |
| . | . | . | . | . | . | . | . |
| . | . | . | . | D | D | D | D |
| . | . | . | . | D | D | D | U |

| | | | | | | | |
|---|---|---|---|---|---|---|---|
| . | . | . | . | . | . | . | . |
| . | . | . | . | . | . | . | . |
| Primary_hyp | MedGen:C00 | criteria_provide | Uncertain_si | D | D | D | D |
| . | . | . | . | . | . | . | . |
| . | . | . | . | . | . | . | . |
| . | . | . | . | D | D | D | N |
| . | . | . | . | . | . | . | . |
| . | . | . | . | . | . | . | . |
| . | . | . | . | . | . | . | . |
| . | . | . | . | . | . | . | . |
| . | . | . | . | D | D | D | D |
| . | . | . | . | . | . | . | . |
| . | . | . | . | D | D | D | D |
| . | . | . | . | D | D | D | D |
| . | . | . | . | D | D | D | D |
| . | . | . | . | D | D | D | D |
| . | . | . | . | D | D | D | N |
| . | . | . | . | D | D | D | D |
| . | . | . | . | . | . | . | . |
| . | . | . | . | D | D | P | D |
| . | . | . | . | D | D | D | D |
| . | . | . | . | D | D | P | D |
| . | . | . | . | . | . | . | D |
| . | . | . | . | D | D | D | D |
| . | . | . | . | D | P | P | D |
| . | . | . | . | . | . | . | . |
| . | . | . | . | . | . | . | . |
| . | . | . | . | D | D | P | D |
| . | . | . | . | . | . | . | . |
| . | . | . | . | . | . | . | . |
| . | . | . | . | D | D | D | N |
| . | . | . | . | D | D | D | D |
| . | . | . | . | D | D | D | D |
| . | . | . | . | D | D | P | D |
| . | . | . | . | D | D | P | . |
| . | . | . | . | . | . | . | . |

| | | | | | | | |
|---|---|---|---|---|---|---|---|
| . | . | . | . | . | . | . | . |
| . | . | . | . | . | . | . | . |
| . | . | . | . | D | D | D | D |
| . | . | . | . | D | D | D | D |
| . | . | . | . | . | . | . | . |
| . | . | . | . | . | . | . | . |
| . | . | . | . | D | D | P | D |
| . | . | . | . | D | D | P | D |
| . | . | . | . | D | P | P | . |
| . | . | . | . | D | D | P | D |
| . | . | . | . | D | D | D | D |
| . | . | . | . | . | . | . | . |
| . | . | . | . | . | . | . | . |
| | | criteria_provide | Pathogenic/L | D | P | B | D |
| . | . | . | . | D | D | P | N |
| . | . | . | . | D | D | D | D |
| . | . | . | . | . | . | . | . |
| . | . | . | . | D | D | D | D |
| . | . | . | . | . | . | . | . |
| Neurofibrom | MedGen:C00 | criteria_provide | Pathogenic | . | . | . | . |
| . | . | . | . | . | . | . | . |
| . | . | . | . | . | . | . | . |
| . | . | . | . | . | . | . | . |
| . | . | . | . | . | . | . | . |
| . | . | . | . | . | . | . | . |
| . | . | . | . | . | . | . | . |
| . | . | . | . | . | . | . | . |
| . | . | . | . | D | D | D | D |
| . | . | . | . | D | D | D | D |
| . | . | . | . | D | D | D | D |
| . | . | . | . | D | P | P | D |
| . | . | . | . | D | D | P | D |
| . | . | . | . | . | . | . | . |
| . | . | . | . | D | D | D | D |
| . | . | . | . | . | . | . | . |
| . | . | . | . | . | . | . | . |

| | | | | | | | |
|---|---|---|---|---|---|---|---|
| . | . | . | . | . | . | . | . |
| . | . | . | . | . | . | . | . |
| . | . | . | . | . | . | . | . |
| . | . | . | . | . | . | . | . |
| . | . | . | . | D | D | P | D |
| . | . | . | . | . | . | . | . |
| . | . | . | . | . | . | . | . |
| . | . | . | . | . | . | . | . |
| . | . | . | . | . | . | . | . |
| . | . | . | . | . | . | . | . |
| . | . | . | . | . | . | . | . |
| . | . | . | . | . | . | . | . |
| . | . | . | . | D | D | P | N |
| . | . | . | . | D | D | D | D |
| . | . | . | . | D | D | D | D |
| . | . | . | . | . | . | . | . |
| . | . | . | . | T | D | D | D |
| . | . | . | . | . | . | . | . |
| . | . | . | . | . | . | . | . |
| . | . | . | . | . | . | . | . |

| FATHMM_pred | MutationTaster_pred | MutationAssessor_pred | InterVar_automated |
|---|---|---|---|
| . | D | M | Uncertain_significance |
| . | . | . | . |
| T | D | M | Uncertain_significance |
| . | . | . | . |
| T | D | N | . |
| . | . | . | . |
| D | D | M | Uncertain_significance |
| T | D | L | Uncertain_significance |
| . | . | . | Uncertain_significance |
| . | . | . | Uncertain_significance |
| . | . | . | Uncertain_significance |
| . | . | . | Uncertain_significance |
| . | . | . | Uncertain_significance |
| . | . | . | Uncertain_significance |
| . | . | . | Uncertain_significance |
| . | . | . | Uncertain_significance |
| . | . | . | . |
| . | N | M | Uncertain_significance |
| . | . | . | Uncertain_significance |
| T | D | M | Uncertain_significance |
| . | P | M | Uncertain_significance |
| T | P | M | Uncertain_significance |
| T | D | M | Uncertain_significance |
| T | D | H | Uncertain_significance |
| T | D | M | Uncertain_significance |
| D | D | L | Uncertain_significance |
| . | . | . | . |
| . | D | M | Uncertain_significance |
| . | . | . | . |
| . | . | . | . |
| D | D | N | Likely_pathogenic |
| T | D | L | Uncertain_significance |
| T | D | M | Uncertain_significance |
| T | D | M | Uncertain_significance |

| | | | |
|---|---|---|---|
| T | D | H | . |
| T | D | M | Uncertain_significance |
| D | D | L | Uncertain_significance |
| D | D | M | Uncertain_significance |
| . | . | . | . |
| T | D | H | . |
| T | D | L | Uncertain_significance |
| T | D | M | Uncertain_significance |
| . | . | . | . |
| . | D | . | Uncertain_significance |
| T | D | M | Uncertain_significance |
| T | D | M | Uncertain_significance |
| T | D | M | Uncertain_significance |
| . | D | . | . |
| T | D | L | Uncertain_significance |
| . | . | . | . |
| . | . | . | . |
| . | . | . | Uncertain_significance |
| D | D | H | Uncertain_significance |
| T | D | L | Uncertain_significance |
| . | . | . | Uncertain_significance |
| . | . | . | . |
| T | D | M | Uncertain_significance |
| T | D | H | Uncertain_significance |
| . | . | . | . |
| . | . | . | Uncertain_significance |
| . | . | . | Uncertain_significance |
| . | . | . | . |
| D | D | M | Uncertain_significance |
| . | . | . | . |
| . | . | . | . |
| T | D | M | Uncertain_significance |
| T | D | M | . |
| . | . | . | . |
| T | D | M | Uncertain_significance |
| T | D | H | Uncertain_significance |

| | | | |
|---|---|---|---|
| . | . | . | . |
| . | . | . | . |
| D | D | M | Uncertain_significance |
| . | . | . | . |
| . | . | . | . |
| D | D | M | Uncertain_significance |
| . | D | . | . |
| . | . | . | Uncertain_significance |
| . | . | . | Uncertain_significance |
| . | . | . | Uncertain_significance |
| T | D | M | Uncertain_significance |
| . | . | . | . |
| T | D | M | Uncertain_significance |
| T | D | H | Uncertain_significance |
| T | D | L | Uncertain_significance |
| D | D | H | Uncertain_significance |
| T | D | H | Uncertain_significance |
| D | D | M | Uncertain_significance |
| . | . | . | Uncertain_significance |
| D | D | M | Uncertain_significance |
| T | D | M | . |
| D | D | H | Uncertain_significance |
| . | A | . | Uncertain_significance |
| T | D | M | . |
| T | D | L | Uncertain_significance |
| . | . | . | . |
| . | . | . | . |
| T | D | L | Uncertain_significance |
| . | . | . | . |
| . | N | . | Likely_pathogenic |
| T | D | M | Uncertain_significance |
| T | D | L | Uncertain_significance |
| T | D | M | Uncertain_significance |
| T | D | L | Uncertain_significance |
| T | N | M | Uncertain_significance |
| . | . | . | Uncertain_significance |

| | | | |
|---|---|---|---|
| . | . | . | Uncertain_significance |
| . | . | . | Uncertain_significance |
| T | D | M | Uncertain_significance |
| T | D | M | Uncertain_significance |
| . | . | . | Uncertain_significance |
| . | . | . | . |
| T | D | M | Uncertain_significance |
| D | D | M | Uncertain_significance |
| D | N | M | . |
| T | D | L | Uncertain_significance |
| D | D | N | Uncertain_significance |
| . | . | . | . |
| . | . | . | . |
| D | A | M | Likely_pathogenic |
| T | D | M | Uncertain_significance |
| T | D | M | Uncertain_significance |
| . | . | . | . |
| D | D | H | . |
| . | . | . | . |
| . | D | . | . |
| . | . | . | Uncertain_significance |
| . | . | . | Uncertain_significance |
| . | . | . | Uncertain_significance |
| . | . | . | Uncertain_significance |
| . | . | . | Uncertain_significance |
| . | . | . | . |
| . | . | . | . |
| D | D | N | Likely_pathogenic |
| D | D | M | Uncertain_significance |
| T | D | M | Uncertain_significance |
| D | D | L | Uncertain_significance |
| T | D | M | Uncertain_significance |
| . | . | . | . |
| T | D | M | Uncertain_significance |
| . | . | . | . |
| . | . | . | Uncertain_significance |

| | | | |
|---|---|---|---|
| . | . | . | Uncertain_significance |
| . | . | . | Uncertain_significance |
| . | . | . | Uncertain_significance |
| . | . | . | Uncertain_significance |
| D | D | M | Uncertain_significance |
| . | . | . | Uncertain_significance |
| . | . | . | Uncertain_significance |
| . | . | . | Uncertain_significance |
| . | . | . | Uncertain_significance |
| . | . | . | Uncertain_significance |
| . | . | . | Uncertain_significance |
| . | . | . | . |
| . | P | L | Uncertain_significance |
| T | D | L | Uncertain_significance |
| T | D | M | Uncertain_significance |
| . | . | . | . |
| D | D | L | Uncertain_significance |
| . | . | . | Uncertain_significance |
| . | . | . | Uncertain_significance |
| . | . | . | Uncertain_significance |

#HS-Sch-2 cell line variants with a potential impact on protein function

| chr | start | end | Sample | Gene.refGene | REF |
|---|---|---|---|---|---|
| chr1 | 11661214 | 11661214 | HS-PSS | FBXO44 | G |
| chr1 | 12719616 | 12719616 | HS-PSS | AADACL3 | C |
| chr1 | 12725527 | 12725527 | HS-PSS | AADACL3 | T |
| chr1 | 12760937 | 12760937 | HS-PSS | C1orf158 | C |
| chr1 | 26282328 | 26282340 | HS-PSS | UBXN11 | AGGGACTGGGGCC |
| chr1 | 26774809 | 26774809 | HS-PSS | ARID1A | C |
| chr1 | 28480604 | 28480604 | HS-PSS | PHACTR4 | G |
| chr1 | 108929794 | 108929794 | HS-PSS | GPSM2 | C |
| chr1 | 120460621 | 120460621 | HS-PSS | NBPF8 | C |
| chr1 | 145103029 | 145103029 | HS-PSS | FAM72C\x3bFAM | G |
| chr1 | 145873487 | 145873487 | HS-PSS | ANKRD35 | G |
| chr1 | 145873971 | 145873971 | HS-PSS | ANKRD35 | C |
| chr1 | 145898994 | 145898994 | HS-PSS | ITGA10 | C |
| chr1 | 146066541 | 146066541 | HS-PSS | NBPF10 | T |
| chr1 | 146984940 | 146984940 | HS-PSS | NBPF12 | T |
| chr1 | 149062099 | 149062099 | HS-PSS | NBPF9 | A |
| chr1 | 149071644 | 149071644 | HS-PSS | NBPF9 | T |
| chr1 | 149390802 | 149390802 | HS-PSS | NOTCH2NLC | A |
| chr1 | 149554594 | 149554594 | HS-PSS | NBPF19 | C |
| chr1 | 149584102 | 149584102 | HS-PSS | PPIAL4C | C |
| chr1 | 152112085 | 152112085 | HS-PSS | TCHH | T |
| chr1 | 153934802 | 153934811 | HS-PSS | DENND4B | CCTGCTGCTG |
| chr1 | 155208991 | 155208991 | HS-PSS | MTX1 | T |
| chr1 | 157578529 | 157578529 | HS-PSS | FCRL4 | C |
| chr1 | 157696099 | 157696099 | HS-PSS | FCRL3 | C |
| chr1 | 173041489 | 173041489 | HS-PSS | TNFSF18 | C |
| chr1 | 206110373 | 206110373 | HS-PSS | AVPR1B | C |
| chr1 | 230425760 | 230425760 | HS-PSS | PGBD5 | C |
| chr2 | 15353567 | 15353567 | HS-PSS | NBAS | A |
| chr2 | 42054204 | 42054204 | HS-PSS | PKDCC | G |
| chr2 | 88186041 | 88186041 | HS-PSS | THNSL2 | A |
| chr2 | 104856901 | 104856901 | HS-PSS | POU3F3 | T |
| chr2 | 132320213 | 132320213 | HS-PSS | ZNF806 | A |
| chr2 | 160137785 | 160137785 | HS-PSS | ITGB6 | G |
| chr2 | 165100357 | 165100357 | HS-PSS | SCN3A | C |
| chr2 | 166238111 | 166238111 | HS-PSS | SCN9A | C |
| chr2 | 169206164 | 169206164 | HS-PSS | LRP2 | G |
| chr3 | 12004751 | 12004751 | HS-PSS | SYN2 | C |
| chr3 | 49358240 | 49358246 | HS-PSS | GPX1 | GGCCGCC |
| chr3 | 49793870 | 49793870 | HS-PSS | CDHR4 | G |
| chr3 | 75730493 | 75730493 | HS-PSS | ZNF717 | C |
| chr3 | 75730499 | 75730499 | HS-PSS | ZNF717 | G |
| chr3 | 75738187 | 75738187 | HS-PSS | ZNF717 | T |
| chr3 | 184358726 | 184358726 | HS-PSS | CLCN2 | A |
| chr4 | 3074876 | 3074882 | HS-PSS | HTT | CCAGCAG |
| chr4 | 127643886 | 127643888 | HS-PSS | INTU | CTG |
| chr4 | 147866768 | 147866768 | HS-PSS | ARHGAP10 | C |
| chr5 | 15937180 | 15937180 | HS-PSS | FBXL7 | C |
| chr5 | 140552043 | 140552043 | HS-PSS | SRA1 | A |
| chr5 | 140835336 | 140835336 | HS-PSS | PCDHA7 | T |
| chr6 | 2749370 | 2749371 | HS-PSS | MYLK4 | GA |

| | | | | | |
|---|---|---|---|---|---|
| chr6 | 32223881 | 32223893 | HS-PSS | NOTCH4 | TAGCAGCAGCAGC |
| chr6 | 135457579 | 135457579 | HS-PSS | AHI1 | T |
| chr6 | 167976591 | 167976591 | HS-PSS | HGC6.3 | G |
| chr7 | 35694629 | 35694629 | HS-PSS | HERPUD2 | T |
| chr7 | 106001527 | 106001527 | HS-PSS | CDHR3 | C |
| chr7 | 143960260 | 143960260 | HS-PSS | OR2F1 | G |
| chr8 | 8318865 | 8318871 | HS-PSS | PRAG1 | CGGGGCG |
| chr8 | 143804099 | 143804099 | HS-PSS | SCRIB | C |
| chr8 | 143977707 | 143977707 | HS-PSS | PARP10 | T |
| chr9 | 20413914 | 20413914 | HS-PSS | MLLT3 | G |
| chr9 | 33367586 | 33367586 | HS-PSS | NFX1 | C |
| chr9 | 34372875 | 34372875 | HS-PSS | MYORG | G |
| chr9 | 78266608 | 78266608 | HS-PSS | CEP78 | C |
| chr9 | 122750491 | 122750491 | HS-PSS | OR1L6 | T |
| chr9 | 127454558 | 127454558 | HS-PSS | LRSAM1 | A |
| chr9 | 137033208 | 137033208 | HS-PSS | C9orf139 | G |
| chr10 | 25596788 | 25596788 | HS-PSS | GPR158 | G |
| chr10 | 35062974 | 35062974 | HS-PSS | CUL2 | G |
| chr10 | 122461902 | 122461902 | HS-PSS | HTRA1 | G |
| chr11 | 54603368 | 54603368 | HS-PSS | OR4C46 | G |
| chr11 | 54603640 | 54603640 | HS-PSS | OR4C46 | T |
| chr11 | 54603820 | 54603820 | HS-PSS | OR4C46 | G |
| chr11 | 54706901 | 54706901 | HS-PSS | OR4A5 | A |
| chr11 | 54707770 | 54707770 | HS-PSS | OR4A5 | G |
| chr11 | 62146777 | 62146777 | HS-PSS | INCENP | G |
| chr11 | 64259233 | 64259233 | HS-PSS | PLCB3 | C |
| chr11 | 67487051 | 67487051 | HS-PSS | AIP | G |
| chr12 | 6667903 | 6667909 | HS-PSS | ZNF384 | TTGCTGC |
| chr12 | 49657114 | 49657114 | HS-PSS | FMNL3 | T |
| chr12 | 69592986 | 69592986 | HS-PSS | CCT2 | C |
| chr12 | 113966224 | 113966224 | HS-PSS | RBM19 | A |
| chr13 | 51797843 | 51797843 | HS-PSS | DHRS12 | G |
| chr14 | 21665434 | 21665434 | HS-PSS | OR4E2 | A |
| chr14 | 21665783 | 21665783 | HS-PSS | OR4E2 | G |
| chr14 | 24300785 | 24300785 | HS-PSS | NOP9 | A |
| chr14 | 56585845 | 56585845 | HS-PSS | TMEM260 | C |
| chr14 | 75579570 | 75579570 | HS-PSS | FLVCR2 | C |
| chr14 | 89412139 | 89412139 | HS-PSS | FOXN3 | C |
| chr14 | 92071009 | 92071009 | HS-PSS | ATXN3 | C |
| chr15 | 23440139 | 23440160 | HS-PSS | GOLGA6L2 | CCTGCTCCTGCATCTTCTCTTG |
| chr15 | 70669096 | 70669096 | HS-PSS | UACA | C |
| chr15 | 75640039 | 75640039 | HS-PSS | IMP3 | C |
| chr15 | 84817038 | 84817038 | HS-PSS | ALPK3 | C |
| chr15 | 90441596 | 90441598 | HS-PSS | IQGAP1 | TTG |
| chr16 | 8796220 | 8796220 | HS-PSS | TMEM186 | C |
| chr16 | 27488272 | 27488272 | HS-PSS | GTF3C1 | G |
| chr16 | 57813722 | 57813730 | HS-PSS | LOC388282 | ATGCAGGTG |
| chr16 | 57897410 | 57897410 | HS-PSS | CNGB1 | G |
| chr16 | 71937415 | 71937415 | HS-PSS | PKD1L3 | A |
| chr16 | 81910721 | 81910721 | HS-PSS | PLCG2 | G |
| chr16 | 88951212 | 88951212 | HS-PSS | LOC100129697 | C |
| chr17 | 7846859 | 7846865 | HS-PSS | KDM6B | TACCACC |
| chr17 | 17816313 | 17816313 | HS-PSS | SREBF1 | G |

| | | | | | |
|---|---|---|---|---|---|
| chr17 | 36001422 | 36001422 | HS-PSS | CCL15 | G |
| chr17 | 36013244 | 36013244 | HS-PSS | CCL23 | T |
| chr17 | 37624364 | 37624364 | HS-PSS | DDX52 | C |
| chr17 | 41034751 | 41034751 | HS-PSS | KRTAP1-3 | C |
| chr17 | 49704114 | 49704114 | HS-PSS | SLC35B1 | A |
| chr17 | 78453419 | 78453419 | HS-PSS | DNAH17 | G |
| chr18 | 677396 | 677396 | HS-PSS | ENOSF1 | T |
| chr18 | 46253735 | 46253738 | HS-PSS | C18orf25 | TCTG |
| chr19 | 1398943 | 1398943 | HS-PSS | GAMT | C |
| chr19 | 2191147 | 2191147 | HS-PSS | DOT1L | G |
| chr19 | 2191148 | 2191148 | HS-PSS | DOT1L | A |
| chr19 | 3752546 | 3752546 | HS-PSS | APBA3 | C |
| chr19 | 12675836 | 12675836 | HS-PSS | DHPS | G |
| chr19 | 17286647 | 17286647 | HS-PSS | ANKLE1 | G |
| chr19 | 19527717 | 19527717 | HS-PSS | NDUFA13 | C |
| chr19 | 39885782 | 39885782 | HS-PSS | FCGBP | G |
| chr19 | 40560380 | 40560380 | HS-PSS | SPTBN4 | G |
| chr19 | 41306533 | 41306533 | HS-PSS | HNRNPUL1 | G |
| chr19 | 44387622 | 44387622 | HS-PSS | ZNF285 | C |
| chr19 | 44430314 | 44430314 | HS-PSS | ZNF229 | A |
| chr20 | 3046479 | 3046479 | HS-PSS | MRPS26 | G |
| chr21 | 5120837 | 5120838 | HS-PSS | GATD3A | TC |
| chr21 | 31688433 | 31688434 | HS-PSS | SCAF4 | AT |
| chr21 | 44311062 | 44311062 | HS-PSS | PFKL | C |
| chr21 | 44627276 | 44627276 | HS-PSS | KRTAP10-9 | C |
| chrX | 1193297 | 1193297 | HS-PSS | CRLF2 | T |
| chrX | 2955503 | 2955503 | HS-PSS | ARSE | C |
| chrX | 47081148 | 47081148 | HS-PSS | RGN | T |
| chrX | 72140076 | 72140076 | HS-PSS | NHSL2 | A |
| chrX | 125321539 | 125321539 | HS-PSS | TEX13C | G |
| chrX | 153182483 | 153182483 | HS-PSS | MAGEA1 | A |
| chrX | 153182604 | 153182604 | HS-PSS | MAGEA1 | G |
| chrX | 154653251 | 154653251 | HS-PSS | CTAG2 | C |

| ALT | Func.refGene | ExonicFunc.refGene | VAF |
|---|---|---|---|
| A | exonic | nonsynonymous_SNV | 0.5405405405405 |
| T | exonic | nonsynonymous_SNV | 1 |
| G | exonic | nonsynonymous_SNV | 1 |
| T | exonic | nonsynonymous_SNV | 1 |
| A | exonic | nonframeshift_deletion | 0.4634146341463 |
| T | exonic | stopgain | 0.6285714285714 |
| A | exonic | nonsynonymous_SNV | 0.4680851063829 |
| T | exonic | nonsynonymous_SNV | 0.42 |
| A | exonic | nonsynonymous_SNV | 0.5121951219512 |
| A | exonic | nonsynonymous_SNV | 1 |
| A | exonic | nonsynonymous_SNV | 0.9827586206896 |
| A | exonic | nonsynonymous_SNV | 0.4313725490196 |
| T | exonic | nonsynonymous_SNV | 0.4772727272727 |
| G | exonic | nonsynonymous_SNV | 1 |
| TACGTATTGG | exonic | nonframeshift_insertion | 0.9375 |
| T | exonic | nonsynonymous_SNV | 1 |
| C | exonic | nonsynonymous_SNV | 1 |
| AGGCGGC | exonic | nonframeshift_insertion | 1 |
| G | exonic | nonsynonymous_SNV | 0.4509803921568 |
| T | exonic | nonsynonymous_SNV | 0.6285714285714 |
| TCTC | exonic | nonframeshift_insertion | 0.2631578947368 |
| CCTGCTGCTGCTG | exonic | nonframeshift_insertion | 1 |
| A | exonic | nonsynonymous_SNV | 0.4444444444444 |
| CT | exonic | frameshift_insertion | 0.4629629629629 |
| T | exonic | nonsynonymous_SNV | 0.4324324324324 |
| T | exonic | nonsynonymous_SNV | 0.72 |
| T | exonic | nonsynonymous_SNV | 0.5555555555555 |
| G | exonic | nonsynonymous_SNV | 0.2 |
| G | exonic | nonsynonymous_SNV | 125 |
| T | exonic | nonsynonymous_SNV | 0.7027027027027 |
| G | exonic | nonsynonymous_SNV | 0.1333333333333 |
| C | exonic | nonsynonymous_SNV | 0.1428571428571 |
| G | exonic | nonsynonymous_SNV | 1 |
| T | exonic | nonsynonymous_SNV | 0.4634146341463 |
| T | exonic | nonsynonymous_SNV | 0.7894736842105 |
| A | exonic | nonsynonymous_SNV | 0.3076923076923 |
| A | exonic | nonsynonymous_SNV | 0.9523809523809 |
| CGCCCGCGCCGCA | exonic | nonframeshift_insertion | 1 |
| GGCC | exonic | nonframeshift_deletion | 1 |
| T | exonic | stopgain | 0.4347826086956 |
| A | exonic | nonsynonymous_SNV | 0.1818181818181 |
| C | exonic | stopgain | 0.1612903225806 |
| C | exonic | nonsynonymous_SNV | 0.2881355932203 |
| T | exonic | stopgain | 0.6097560975609 |
| CCAG | exonic | nonframeshift_deletion | 0.8461538461538 |
| C | exonic | frameshift_deletion | 1 |
| CA | exonic | frameshift_insertion | 1 |
| G | exonic | nonsynonymous_SNV | 675 |
| AGTC | exonic | nonframeshift_insertion | 0.5681818181818 |
| C | exonic | nonsynonymous_SNV | 0.5102040816326 |
| G | exonic | frameshift_deletion | 0.5 |

| | | | |
|---|---|---|---|
| TAGCAGCAGC | exonic | nonframeshift_deletion | 1 |
| G | exonic | nonsynonymous_SNV | 0.4038461538461 |
| A | exonic | nonsynonymous_SNV | 0.4 |
| A | splicing | . | 0.2545454545454 |
| T | exonic | nonsynonymous_SNV | 0.4468085106382 |
| A | exonic | nonsynonymous_SNV | 0.4166666666666 |
| C | exonic | nonframeshift_deletion | 0.5652173913043 |
| T | exonic | nonsynonymous_SNV | 0.5098039215686 |
| C | exonic | nonsynonymous_SNV | 0.4102564102564 |
| A | exonic | nonsynonymous_SNV | 0.2666666666666 |
| A | exonic | nonsynonymous_SNV | 525 |
| C | exonic | stopgain | 0.5142857142857 |
| T | exonic | nonsynonymous_SNV | 575 |
| G | exonic | nonsynonymous_SNV | 1 |
| G | exonic | nonsynonymous_SNV | 0.4782608695652 |
| A | exonic | nonsynonymous_SNV | 0.5 |
| A | exonic | nonsynonymous_SNV | 0.4615384615384 |
| A | exonic | nonsynonymous_SNV | 0.3636363636363 |
| A | exonic | nonsynonymous_SNV | 0.5 |
| A | exonic | nonsynonymous_SNV | 1 |
| C | exonic | nonsynonymous_SNV | 1 |
| A | exonic | nonsynonymous_SNV | 1 |
| G | exonic | nonsynonymous_SNV | 1 |
| A | exonic | nonsynonymous_SNV | 1 |
| GGAGCAGGAGCAGCGC | exonic | nonframeshift_insertion | 0.3902439024390 |
| G | exonic | nonsynonymous_SNV | 0.4285714285714 |
| A | exonic | nonsynonymous_SNV | 0.4 |
| TTGC | exonic | nonframeshift_deletion | 1 |
| C | exonic | nonsynonymous_SNV | 0.4489795918367 |
| G | exonic | nonsynonymous_SNV | 0.5813953488372 |
| G | exonic | nonsynonymous_SNV | 0.1481481481481 |
| A | exonic | nonsynonymous_SNV | 0.5833333333333 |
| G | exonic | nonsynonymous_SNV | 0.5263157894736 |
| A | exonic | nonsynonymous_SNV | 0.3055555555555 |
| T | exonic | nonsynonymous_SNV | 0.5769230769230 |
| T | exonic | nonsynonymous_SNV | 0.4318181818181 |
| A | exonic | nonsynonymous_SNV | 0.3714285714285 |
| T | exonic | nonsynonymous_SNV | 0.16 |
| CGCTGCTGCTGCTGCTGC | exonic | nonframeshift_insertion | 1 |
| C | exonic | nonframeshift_deletion | 0.1666666666666 |
| T | exonic | nonsynonymous_SNV | 0.5789473684210 |
| A | exonic | nonsynonymous_SNV | 0.5 |
| A | exonic | stopgain | 0.3636363636363 |
| T | exonic | frameshift_deletion | 0.4117647058823 |
| T | exonic | nonsynonymous_SNV | 0.6538461538461 |
| A | exonic | nonsynonymous_SNV | 0.4722222222222 |
| A | exonic | frameshift_deletion | 0.4565217391304 |
| A | exonic | nonsynonymous_SNV | 0.56 |
| AGCACCTGAGAAT | exonic | nonframeshift_insertion | 0.6896551724137 |
| A | splicing | . | 0.4736842105263 |
| G | exonic | nonsynonymous_SNV | 0.3809523809523 |
| TACCACCACCACC | exonic | nonframeshift_insertion | 1 |
| T | exonic | nonsynonymous_SNV | 0.1304347826086 |

| | | | |
|---|---|---|---|
| A | exonic | nonsynonymous_SNV | 0.4761904761904 |
| C | exonic | nonsynonymous_SNV | 0.5 |
| T | exonic | nonsynonymous_SNV | 0.5660377358490 |
| G | exonic | nonsynonymous_SNV | 0.2692307692307 |
| G | exonic | nonsynonymous_SNV | 0.4523809523809 |
| C | exonic | nonsynonymous_SNV | 0.3703703703703 |
| C | exonic | nonsynonymous_SNV | 0.3846153846153 |
| T | exonic | nonframeshift_deletion | 1 |
| G | exonic | nonsynonymous_SNV | 0.3333333333333 |
| T | exonic | stopgain | 0.3469387755102 |
| T | exonic | nonsynonymous_SNV | 0.3541666666666 |
| T | exonic | nonsynonymous_SNV | 0.6 |
| A | exonic | nonsynonymous_SNV | 0.5957446808510 |
| GGTGT | exonic | frameshift_insertion | 0.5652173913043 |
| G | exonic | nonsynonymous_SNV | 0.7142857142857 |
| T | exonic | nonsynonymous_SNV | 0.4 |
| C | exonic | nonsynonymous_SNV | 1 |
| A | exonic | nonsynonymous_SNV | 1 |
| T | exonic | nonsynonymous_SNV | 1 |
| G | exonic | nonsynonymous_SNV | 1 |
| T | exonic | nonsynonymous_SNV | 0.1428571428571 |
| T | exonic | stopgain | 0.56 |
| A | exonic | frameshift_deletion | 0.4074074074074 |
| G | exonic | nonsynonymous_SNV | 0.4857142857142 |
| G | exonic | nonsynonymous_SNV | 0.5675675675675 |
| C | exonic | nonsynonymous_SNV | 1 |
| T | exonic | nonsynonymous_SNV | 1 |
| C | exonic | nonsynonymous_SNV | 0.1538461538461 |
| T | exonic | nonsynonymous_SNV | 1 |
| A | exonic | nonsynonymous_SNV | 0.1333333333333 |
| G | exonic | nonsynonymous_SNV | 1 |
| A | exonic | nonsynonymous_SNV | 1 |
| G | exonic | nonsynonymous_SNV | 1 |

avsnp150
rs199859446
rs3010877
rs7513079
rs1132185
.
.
rs147207733
rs189033496
rs587696045
rs28685127
rs6670984
rs77387570
rs2274616
.
.
.
.
.
.
.
.
.
rs760077
.
rs200782762
.
rs28632197
.
.
rs374454176
.
.
rs12468523
rs2305820
.
.
.
.
.
.
rs112949302
rs200767888
rs868631560
.
.
.
.
rs537284845
.
.
.

.
.
rs61740140
.
rs372834692
rs143509009
.
rs140972127
.
.
rs2274866
rs4879782
rs770261747
.
.
rs149601197
rs753599474
rs754713861
.
rs77689730
rs74396937
rs11246606
rs10902343
rs74522421
.
.
rs1063385
.
rs149109879
.
.
rs149418560
rs2874103
rs970382
rs200463538
rs144727961
.
.
.
.
rs147142448
rs553010196
rs115479471
.
rs764436817
rs184708220
.
rs201023939
.
.
rs71395352
.
.

rs854625
rs1003645
rs7216445
rs761876245
.
.
.
.
.
.
.
.
rs78193595
.
.
.
.
.
rs2571089
rs2571174
.
.
.
.
rs200685052
.
rs150756612
.
.
.
rs2008160
rs2008144
rs17328091

Variant
chr1:11661214_G/A
chr1:12719616_C/T
chr1:12725527_T/G
chr1:12760937_C/T
chr1:26282328_AGGGACTGGGGCC/A
chr1:26774809_C/T
chr1:28480604_G/A
chr1:108929794_C/T
chr1:120460621_C/A
chr1:145103029_G/A
chr1:145873487_G/A
chr1:145873971_C/A
chr1:145898994_C/T
chr1:146066541_T/G
chr1:146984940_T/TACGTATTGG
chr1:149062099_A/T
chr1:149071644_T/C
chr1:149390802_A/AGGCGGC
chr1:149554594_C/G
chr1:149584102_C/T
chr1:152112085_T/TCTC
chr1:153934802_CCTGCTGCTG/CCTGCTGCTGCTG
chr1:155208991_T/A
chr1:157578529_C/CT
chr1:157696099_C/T
chr1:173041489_C/T
chr1:206110373_C/T
chr1:230425760_C/G
chr2:15353567_A/G
chr2:42054204_G/T
chr2:88186041_A/G
chr2:104856901_T/C
chr2:132320213_A/G
chr2:160137785_G/T
chr2:165100357_C/T
chr2:166238111_C/A
chr2:169206164_G/A
chr3:12004751_C/CGCCCGCGCCGCA
chr3:49358240_GGCCGCC/GGCC
chr3:49793870_G/T
chr3:75730493_C/A
chr3:75730499_G/C
chr3:75738187_T/C
chr3:184358726_A/T
chr4:3074876_CCAGCAG/CCAG
chr4:127643886_CTG/C
chr4:147866768_C/CA
chr5:15937180_C/G
chr5:140552043_A/AGTC
chr5:140835336_T/C
chr6:2749370_GA/G

chr6:32223881_TAGCAGCAGCAGC/TAGCAGCAGC
chr6:135457579_T/G
chr6:167976591_G/A
chr7:35694629_T/A
chr7:106001527_C/T
chr7:143960260_G/A
chr8:8318865_CGGGGCG/C
chr8:143804099_C/T
chr8:143977707_T/C
chr9:20413914_G/A
chr9:33367586_C/A
chr9:34372875_G/C
chr9:78266608_C/T
chr9:122750491_T/G
chr9:127454558_A/G
chr9:137033208_G/A
chr10:25596788_G/A
chr10:35062974_G/A
chr10:122461902_G/A
chr11:54603368_G/A
chr11:54603640_T/C
chr11:54603820_G/A
chr11:54706901_A/G
chr11:54707770_G/A
chr11:62146777_G/GGAGCAGGAGCAGCGC
chr11:64259233_C/G
chr11:67487051_G/A
chr12:6667903_TTGCTGC/TTGC
chr12:49657114_T/C
chr12:69592986_C/G
chr12:113966224_A/G
chr13:51797843_G/A
chr14:21665434_A/G
chr14:21665783_G/A
chr14:24300785_A/T
chr14:56585845_C/T
chr14:75579570_C/A
chr14:89412139_C/T
chr14:92071009_C/CGCTGCTGCTGCTGCTGCTGCTGCTGCTGCTGCTGCTGCT
chr15:23440139_CCTGCTCCTGCATCTTCTCTTG/C
chr15:70669096_C/T
chr15:75640039_C/A
chr15:84817038_C/A
chr15:90441596_TTG/T
chr16:8796220_C/T
chr16:27488272_G/A
chr16:57813722_ATGCAGGTG/A
chr16:57897410_G/A
chr16:71937415_A/AGCACCTGAGAAT
chr16:81910721_G/A
chr16:88951212_C/G
chr17:7846859_TACCACC/TACCACCACCACC
chr17:17816313_G/T

chr17:36001422_G/A
chr17:36013244_T/C
chr17:37624364_C/T
chr17:41034751_C/G
chr17:49704114_A/G
chr17:78453419_G/C
chr18:677396_T/C
chr18:46253735_TCTG/T
chr19:1398943_C/G
chr19:2191147_G/T
chr19:2191148_A/T
chr19:3752546_C/T
chr19:12675836_G/A
chr19:17286647_G/GGTGT
chr19:19527717_C/G
chr19:39885782_G/T
chr19:40560380_G/C
chr19:41306533_G/A
chr19:44387622_C/T
chr19:44430314_A/G
chr20:3046479_G/T
chr21:5120837_TC/T
chr21:31688433_AT/A
chr21:44311062_C/G
chr21:44627276_C/G
chrX:1193297_T/C
chrX:2955503_C/T
chrX:47081148_T/C
chrX:72140076_A/T
chrX:125321539_G/A
chrX:153182483_A/G
chrX:153182604_G/A
chrX:154653251_C/G

| AAChange.refGene | GeneDetail.refGene | AF |
|---|---|---|
| | . | 0.0002 |
| AADACL3:NM_001103 | . | . |
| AADACL3:NM_001103 | . | . |
| C1orf158:NM_001330 | . | . |
| UBXN11:NM_0010772 | . | . |
| ARID1A:NM_006015:e | . | . |
| | . | . |
| GPSM2:NM_00132103 | . | 0.0002 |
| NBPF8:NM_00103750 | . | . |
| FAM72D:NM_0013459 | . | . |
| ANKRD35:NM_001280 | . | . |
| ANKRD35:NM_001280 | . | . |
| ITGA10:NM_00130304 | . | . |
| NBPF10:NM_0010397 | . | . |
| NBPF12:NM_0012781 | . | . |
| NBPF9:NM_00103767 | . | . |
| NBPF9:NM_00103767 | . | . |
| NOTCH2NLC:NM_0013 | . | . |
| NBPF19:NM_0013513 | . | . |
| PPIAL4C:NM_0011357 | . | . |
| TCHH:NM_007113:exc | . | . |
| DENND4B:NM_00136 | . | . |
| MTX1:NM_002455:exc | . | . |
| FCRL4:NM_031282:ex | . | . |
| FCRL3:NM_00132033 | . | . |
| TNFSF18:NM_005092: | . | . |
| AVPR1B:NM_000707:e | . | . |
| PGBD5:NM_00125831 | . | . |
| NBAS:NM_015909:exc | . | . |
| PKDCC:NM_138370:ex | . | . |
| THNSL2:NM_018271:e | . | . |
| POU3F3:NM_006236:e | . | . |
| ZNF806:NM_0013554 | . | . |
| | . | . |
| SCN3A:NM_00108167 | . | . |
| SCN9A:NM_00136553 | . | . |
| LRP2:NM_004525:exo | . | . |
| SYN2:NM_003178:exo | . | . |
| GPX1:NM_001329455 | . | . |
| CDHR4:NM_00100754 | . | . |
| ZNF717:NM_0013240 | . | 0.0016 |
| ZNF717:NM_0013240 | . | 0.0038 |
| ZNF717:NM_0011282 | . | 0.0006 |
| CLCN2:NM_00117108 | . | . |
| HTT:NM_002111:exon | . | . |
| INTU:NM_015693:exo | . | . |
| ARHGAP10:NM_02460 | . | . |
| FBXL7:NM_001278317 | . | 0.0003 |
| SRA1:NM_001035235: | . | . |
| PCDHA7:NM_018910: | . | . |
| MYLK4:NM_00134787 | . | . |

| | | |
|---|---|---|
| NOTCH4:NM_004557: | . | . |
| AHI1:NM_001134830: | . | . |
| HGC6.3:NM_00112989 | . | 0.0027 |
| . | NM_022373:exon2:UTF | . |
| CDHR3:NM_00130116 | . | . |
| OR2F1:NM_012369:ex | . | 0.0023 |
| | . | . |
| SCRIB:NM_015356:exo | . | 0.0003 |
| PARP10:NM_00131789 | . | . |
| MLLT3:NM_00128669 | . | . |
| NFX1:NM_001318758: | . | 0.0002 |
| MYORG:NM_020702:e | . | . |
| | . | . |
| OR1L6:NM_00100445 | . | . |
| LRSAM1:NM_0010053 | . | . |
| C9orf139:NM_207511 | . | 6.368e-05 |
| GPR158:NM_020752:e | . | . |
| CUL2:NM_001198777: | . | . |
| HTRA1:NM_002775:ex | . | . |
| OR4C46:NM_0010047 | . | . |
| OR4C46:NM_0010047 | . | . |
| OR4C46:NM_0010047 | . | . |
| OR4A5:NM_00100527 | . | . |
| OR4A5:NM_00100527 | . | . |
| INCENP:NM_020238:e | . | 0.0003 |
| PLCB3:NM_00118488 | . | . |
| AIP:NM_001302960:e | . | . |
| ZNF384:NM_0010399 | . | . |
| FMNL3:NM_198900:e | . | 0.0004 |
| CCT2:NM_001198842: | . | . |
| RBM19:NM_00114669 | . | . |
| DHRS12:NM_0012704 | . | 0.0004 |
| OR4E2:NM_00100191 | . | . |
| OR4E2:NM_00100191 | . | . |
| NOP9:NM_001286367 | . | 9.556e-05 |
| TMEM260:NM_01779 | . | 0.0002 |
| FLVCR2:NM_017791:e | . | . |
| FOXN3:NM_00108547 | . | . |
| | . | . |
| GOLGA6L2:NM_00130 | . | . |
| UACA:NM_001008224 | . | 6.369e-05 |
| IMP3:NM_018285:exo | . | . |
| ALPK3:NM_020778:ex | . | 0.0005 |
| IQGAP1:NM_003870:e | . | . |
| TMEM186:NM_01542 | . | . |
| GTF3C1:NM_0012862 | . | . |
| LOC388282:NM_0012 | . | . |
| CNGB1:NM_00128613 | . | . |
| PKD1L3:NM_181536:e | . | . |
| . | NM_002661:exon18:c.: | . |
| LOC100129697:NM_0( | . | 0.0020 |
| KDM6B:NM_00108042 | . | . |
| SREBF1:NM_00132109 | . | . |

| | | |
|---|---|---|
| CCL15:NM_032965:ex | . | . |
| CCL23:NM_005064:ex | . | . |
| DDX52:NM_007010:e› | . | . |
| KRTAP1-3:NM_03096€ | . | 0.0011 |
| SLC35B1:NM_001278‌7 | . | . |
| DNAH17:NM_173628: | . | . |
| | . | . |
| C18orf25:NM_001008 | . | . |
| GAMT:NM_000156:ex | . | . |
| DOT1L:NM_032482:ex | . | . |
| DOT1L:NM_032482:ex | . | . |
| APBA3:NM_004886:ex | . | . |
| DHPS:NM_001369691 | . | 6.373e-05 |
| ANKLE1:NM_0012784 | . | . |
| NDUFA13:NM_01596‌5 | . | . |
| FCGBP:NM_003890:ex | . | . |
| SPTBN4:NM_025213:e | . | . |
| HNRNPUL1:NM_0013C | . | . |
| ZNF285:NM_0012914‌9 | . | . |
| ZNF229:NM_0012785 | . | . |
| MRPS26:NM_030811: | . | . |
| GATD3A:NM_0013203 | . | . |
| SCAF4:NM_00114544‌4 | . | . |
| PFKL:NM_002626:exo | . | . |
| KRTAP10-9:NM_1986‌9 | . | 0.0005 |
| CRLF2:NM_00101228‌8 | . | . |
| ARSE:NM_001282631: | . | 0.0001 |
| RGN:NM_004683:exor | . | . |
| NHSL2:NM_00101362 | . | . |
| TEX13C:NM_0011952‌7 | . | . |
| MAGEA1:NM_004988: | . | . |
| MAGEA1:NM_004988: | . | . |
| CTAG2:NM_020994:ex | . | . |

| AF_popmax | prop_pathogenic_pred | CLNALLELEID |
|---|---|---|
| 0.0032 | 0.833333333333333 | . |
| . | NA | . |
| . | NA | . |
| . | NA | . |
| . | NA | . |
| . | 1 | . |
| . | 0.857142857142857 | . |
| 0.0026 | 1 | 206697 |
| . | NA | . |
| . | NA | . |
| . | NA | . |
| . | NA | . |
| . | NA | . |
| . | NA | . |
| . | NA | . |
| . | NA | . |
| . | NA | . |
| . | NA | . |
| . | 0.75 | . |
| . | NA | . |
| . | NA | . |
| . | NA | . |
| . | NA | . |
| . | NA | . |
| . | 0.857142857142857 | . |
| . | 0.714285714285714 | . |
| . | NA | . |
| . | NA | . |
| . | 0.714285714285714 | . |
| . | 1 | . |
| . | 0.714285714285714 | . |
| . | 0.714285714285714 | . |
| . | NA | . |
| . | 0.857142857142857 | . |
| . | 1 | . |
| . | 1 | . |
| . | 1 | . |
| . | NA | . |
| . | NA | . |
| . | 1 | . |
| 0.0028 | NA | . |
| 0.0075 | NA | . |
| 0.0047 | 0.666666666666667 | . |
| . | 1 | . |
| . | NA | . |
| . | NA | . |
| . | NA | . |
| 0.0051 | 0.857142857142857 | . |
| . | NA | . |
| . | 0.714285714285714 | . |
| . | NA | . |

| | | | |
|---|---|---|---|
| . | NA | . | |
| . | 0.714285714285714 | . | |
| 0.0064 | NA | . | |
| . | NA | . | |
| . | 0.857142857142857 | . | |
| 0.0071 | 0.857142857142857 | . | |
| . | NA | . | |
| 0.0064 | 0.666666666666667 | . | |
| . | 0.714285714285714 | . | |
| . | 0.833333333333333 | . | |
| 0.0038 | 0.857142857142857 | . | |
| . | NA | . | |
| . | 0.714285714285714 | . | |
| . | 0.857142857142857 | . | |
| . | 0.714285714285714 | . | |
| 0.0013 | NA | . | |
| . | 0.857142857142857 | . | |
| . | 0.714285714285714 | . | |
| . | 0.714285714285714 | . | |
| . | NA | . | |
| . | NA | . | |
| . | NA | . | |
| . | NA | . | |
| . | NA | . | |
| 0.0039 | NA | . | |
| . | 0.714285714285714 | . | |
| . | 1 | | 49586 |
| . | NA | . | |
| 0.0083 | 0.714285714285714 | . | |
| . | 0.857142857142857 | . | |
| . | 0.857142857142857 | . | |
| 0.0064 | 0.666666666666667 | . | |
| . | NA | . | |
| . | NA | . | |
| 0.0019 | 0.857142857142857 | . | |
| 0.0032 | 0.857142857142857 | . | |
| . | 0.714285714285714 | . | |
| . | 1 | . | |
| . | NA | . | |
| . | NA | . | |
| 0.0013 | 0.857142857142857 | . | |
| . | 0.857142857142857 | . | |
| 0.0064 | 1 | . | |
| . | NA | . | |
| . | 0.714285714285714 | . | |
| . | 0.857142857142857 | . | |
| . | NA | . | |
| . | 1 | . | |
| . | NA | . | |
| . | 1 | . | |
| 0.0052 | NA | . | |
| . | NA | . | |
| . | 0.857142857142857 | . | |

| | | | | |
|---|---|---|---|---|
| . | | NA | | . |
| . | | NA | | . |
| . | | NA | | . |
| | 3 | 0.8 | | . |
| . | | 0.857142857142857 | | . |
| . | | 0.833333333333333 | | . |
| . | | 0.857142857142857 | | . |
| . | | NA | | . |
| . | | | 1 | . |
| . | | | 1 | . |
| . | | 0.857142857142857 | | . |
| . | | 0.857142857142857 | | . |
| 0.0013 | | NA | | . |
| . | | NA | | . |
| . | | 0.857142857142857 | | . |
| . | | 0.75 | | . |
| . | | 0.857142857142857 | | . |
| . | | 0.714285714285714 | | . |
| . | | NA | | . |
| . | | NA | | . |
| . | | 0.666666666666667 | | . |
| . | | NA | | . |
| . | | NA | | . |
| . | | 0.857142857142857 | | . |
| 0.0096 | | 0.833333333333333 | | . |
| . | | NA | | . |
| | 2 | 0.857142857142857 | | . |
| . | | | 1 | . |
| . | | 0.714285714285714 | | . |
| . | | NA | | . |
| . | | NA | | . |
| . | | NA | | . |
| . | | NA | | . |

CLNDN
.
.
.
.
.
.
.
not_specified|Nonsyndromic_Hearing_Loss,_Recessive
.
.
.
.
.
.
.
.
.
.
.
.
.
.
.
.
.
.
.
.
.
.
.
.
.
.
.
.
.
.
.
.
.
.
.
.
.
.
.
.
.
.
.

.
.
.
.
.
.
.
.
.
.
.
.
.
.
.
.
.
.
.
.
.
.
.
.
.
.
Hereditary_cancer-predisposing_syndrome|Somatotroph_adenoma|Familial_Isolated_Pituitary_Ade
.
.
.
.
.
.
.
.
.
.
.
.
.
.
.
.
.
.
.
.
.
.
.
.
.
.

CLNDISDB
.
.
.
.
.
.
.
MedGen:CN169374|MedGen:CN239439
.
.
.
.
.
.
.
.
.
.
.
.
.
.
.
.
.
.
.
.
.
.
.
.
.
.
.
.
.
.
.
.
.
.
.
.
.
.
.
.
.
.
.

.
.
.
.
.
.
.
.
.
.
.
.
.
.
.
.
.
.
.
.
.
.
.
.
.
.
MedGen:C0027672,SNOMED_CT:699346009|MedGen:C4538355,OMIM:102200,SNOMED_CT:25495
.
.
.
.
.
.
.
.
.
.
.
.
.
.
.
.
.
.
.
.
.
.
.
.
.
.

| CLNREVSTAT | CLNSIG |
|---|---|
| . | . |
| . | . |
| . | . |
| . | . |
| . | . |
| . | . |
| . | . |
| criteria_provided,_multiple_submitters,_no_conflicts | Uncertain_significance |
| . | . |
| . | . |
| . | . |
| . | . |
| . | . |
| . | . |
| . | . |
| . | . |
| . | . |
| . | . |
| . | . |
| . | . |
| . | . |
| . | . |
| . | . |
| . | . |
| . | . |
| . | . |
| . | . |
| . | . |
| . | . |
| . | . |
| . | . |
| . | . |
| . | . |
| . | . |
| . | . |
| . | . |
| . | . |
| . | . |
| . | . |
| . | . |
| . | . |
| . | . |
| . | . |
| . | . |
| . | . |
| . | . |
| . | . |
| . | . |
| . | . |

| | |
|---|---|
| . | . |
| . | . |
| . | . |
| . | . |
| . | . |
| . | . |
| . | . |
| . | . |
| . | . |
| . | . |
| . | . |
| . | . |
| . | . |
| . | . |
| . | . |
| . | . |
| . | . |
| . | . |
| . | . |
| . | . |
| . | . |
| . | . |
| . | . |
| . | . |
| . | . |
| . | . |
| criteria_provided,_conflicting_interpretations | Conflicting_interpretations_of_pathogenicity |
| . | . |
| . | . |
| . | . |
| . | . |
| . | . |
| . | . |
| . | . |
| . | . |
| . | . |
| . | . |
| . | . |
| . | . |
| . | . |
| . | . |
| . | . |
| . | . |
| . | . |
| . | . |
| . | . |
| . | . |
| . | . |
| . | . |
| . | . |
| . | . |
| . | . |
| . | . |

| SIFT_pred | Polyphen2_HDIV_pred | Polyphen2_HVAR_pred | LRT_pred | FATHMM_pred |
|---|---|---|---|---|
| D | D | D | . | T |
| . | . | . | . | . |
| . | . | . | . | . |
| . | . | . | . | . |
| . | . | . | . | . |
| . | . | . | D | . |
| D | D | D | D | T |
| D | D | D | D | D |
| . | . | . | . | . |
| . | . | . | . | . |
| . | . | . | . | . |
| . | . | . | . | . |
| . | . | . | . | . |
| . | . | . | . | . |
| . | . | . | . | . |
| . | . | . | . | . |
| . | . | . | . | . |
| . | . | . | . | . |
| . | D | D | . | . |
| . | . | . | . | . |
| . | . | . | . | . |
| . | . | . | . | . |
| . | . | . | . | . |
| . | . | . | . | . |
| D | D | D | D | T |
| D | P | B | N | D |
| . | . | . | . | . |
| . | . | . | . | . |
| T | D | D | D | T |
| D | D | P | D | . |
| D | P | B | D | T |
| D | P | B | U | D |
| . | . | . | . | . |
| D | D | D | D | T |
| D | D | D | D | D |
| D | D | D | D | D |
| D | D | D | D | D |
| . | . | . | . | . |
| . | . | . | . | . |
| . | . | . | . | . |
| . | . | . | . | . |
| . | . | . | . | . |
| D | D | D | . | T |
| . | . | . | D | . |
| . | . | . | . | . |
| . | . | . | . | . |
| . | . | . | . | . |
| D | D | P | D | T |
| . | . | . | . | . |
| D | D | D | U | T |
| . | . | . | . | . |

| | | | | |
|---|---|---|---|---|
| . | . | . | . | . |
| T | D | D | D | T |
| . | . | . | . | . |
| . | . | . | . | . |
| D | D | P | D | T |
| D | D | D | D | T |
| . | . | . | . | . |
| D | D | D | . | T |
| D | D | D | N | T |
| T | D | P | D | . |
| D | D | D | D | T |
| . | . | . | . | . |
| D | D | D | N | T |
| D | D | P | D | T |
| D | P | B | D | T |
| . | . | . | . | . |
| D | D | D | D | T |
| D | D | P | D | T |
| D | D | D | U | T |
| . | . | . | . | . |
| . | . | . | . | . |
| . | . | . | . | . |
| . | . | . | . | . |
| . | . | . | . | . |
| . | . | . | . | . |
| D | D | P | N | T |
| D | D | P | D | D |
| . | . | . | . | . |
| T | P | D | D | D |
| D | D | D | D | T |
| D | D | D | D | T |
| D | D | P | . | T |
| . | . | . | . | . |
| . | . | . | . | . |
| D | P | P | D | T |
| D | D | D | D | T |
| D | B | B | D | D |
| D | P | P | D | D |
| . | . | . | . | . |
| . | . | . | . | . |
| D | P | P | D | T |
| D | P | P | D | T |
| . | . | . | . | . |
| . | . | . | . | . |
| T | D | P | D | T |
| D | D | D | D | T |
| . | . | . | . | . |
| D | D | D | D | D |
| . | . | . | . | . |
| . | . | . | . | . |
| . | . | . | . | . |
| . | . | . | . | . |
| D | D | D | D | T |

| | | | | |
|---|---|---|---|---|
| . | . | . | . | . |
| . | . | . | . | . |
| . | . | . | . | . |
| D | D | D | . | T |
| D | D | D | D | T |
| . | P | P | D | T |
| D | P | D | D | T |
| . | . | . | . | . |
| D | D | P | D | D |
| . | . | . | D | . |
| D | D | D | D | T |
| D | D | D | D | T |
| . | . | . | . | . |
| . | . | . | . | . |
| D | D | D | D | T |
| . | D | P | . | . |
| D | D | D | D | T |
| D | D | D | D | T |
| . | . | . | . | . |
| . | . | . | . | . |
| D | B | B | D | . |
| . | . | . | . | . |
| . | . | . | . | . |
| D | D | D | D | T |
| D | D | P | . | T |
| . | . | . | . | . |
| D | D | D | D | D |
| D | D | D | D | . |
| D | D | D | D | T |
| . | . | . | . | . |
| . | . | . | . | . |
| . | . | . | . | . |
| . | . | . | . | . |

| MutationTaster_pred | MutationAssessor_pred | InterVar_automated |
|---|---|---|
| D | H | Uncertain_significance |
| . | . | Uncertain_significance |
| . | . | Uncertain_significance |
| . | . | Uncertain_significance |
| . | . | . |
| A | . | Pathogenic |
| D | M | Uncertain_significance |
| D | M | Uncertain_significance |
| . | . | Uncertain_significance |
| . | . | Uncertain_significance |
| . | . | Uncertain_significance |
| . | . | Uncertain_significance |
| . | . | . |
| . | . | . |
| . | . | . |
| . | . | Uncertain_significance |
| . | . | Uncertain_significance |
| . | . | . |
| N | M | Uncertain_significance |
| . | . | Uncertain_significance |
| . | . | . |
| . | . | . |
| . | . | Uncertain_significance |
| . | . | . |
| D | H | Uncertain_significance |
| D | M | Uncertain_significance |
| . | . | Uncertain_significance |
| . | . | Uncertain_significance |
| D | M | Uncertain_significance |
| D | M | Uncertain_significance |
| D | M | Uncertain_significance |
| D | M | Uncertain_significance |
| . | . | . |
| D | H | Uncertain_significance |
| D | H | . |
| D | H | Uncertain_significance |
| D | M | Uncertain_significance |
| . | . | . |
| . | . | . |
| A | . | Uncertain_significance |
| . | . | Uncertain_significance |
| . | . | Uncertain_significance |
| N | M | Uncertain_significance |
| D | . | Pathogenic |
| . | . | . |
| . | . | . |
| . | . | . |
| D | M | Uncertain_significance |
| . | . | . |
| D | H | Uncertain_significance |
| . | . | . |

| | | |
|---|---|---|
| . | . | . |
| D | M | Uncertain_significance |
| . | . | . |
| . | . | . |
| D | M | Uncertain_significance |
| D | H | Uncertain_significance |
| . | . | . |
| D | N | Uncertain_significance |
| D | H | Uncertain_significance |
| D | M | . |
| D | M | Uncertain_significance |
| . | . | Uncertain_significance |
| D | M | Uncertain_significance |
| P | H | Uncertain_significance |
| D | M | Uncertain_significance |
| . | . | Uncertain_significance |
| D | M | Uncertain_significance |
| D | L | Uncertain_significance |
| D | M | Uncertain_significance |
| . | . | Uncertain_significance |
| . | . | Uncertain_significance |
| . | . | Uncertain_significance |
| . | . | Uncertain_significance |
| . | . | Uncertain_significance |
| . | . | . |
| D | M | Uncertain_significance |
| D | M | Uncertain_significance |
| . | . | . |
| D | L | Uncertain_significance |
| D | M | Uncertain_significance |
| D | H | Uncertain_significance |
| N | M | Uncertain_significance |
| . | . | Uncertain_significance |
| . | . | Uncertain_significance |
| D | M | Uncertain_significance |
| D | M | Uncertain_significance |
| D | M | Uncertain_significance |
| D | M | Uncertain_significance |
| . | . | . |
| . | . | . |
| D | M | Uncertain_significance |
| D | M | Uncertain_significance |
| A | . | Uncertain_significance |
| . | . | . |
| D | M | . |
| D | M | . |
| . | . | . |
| D | M | Uncertain_significance |
| . | . | . |
| D | . | . |
| . | . | Uncertain_significance |
| . | . | . |
| D | M | Uncertain_significance |

| | | |
|---|---|---|
| . | . | Uncertain_significance |
| . | . | Uncertain_significance |
| . | . | . |
| D | . | Uncertain_significance |
| D | M | Uncertain_significance |
| D | M | . |
| D | M | Uncertain_significance |
| . | . | . |
| D | M | . |
| A | . | Pathogenic |
| D | M | Uncertain_significance |
| D | M | Uncertain_significance |
| . | . | . |
| . | . | . |
| D | M | Uncertain_significance |
| N | M | Uncertain_significance |
| D | M | Uncertain_significance |
| D | N | Uncertain_significance |
| . | . | Uncertain_significance |
| . | . | Uncertain_significance |
| D | M | Uncertain_significance |
| . | . | . |
| . | . | . |
| D | M | Uncertain_significance |
| D | M | Uncertain_significance |
| . | . | Uncertain_significance |
| D | L | Uncertain_significance |
| D | M | Uncertain_significance |
| D | L | Uncertain_significance |
| . | . | Uncertain_significance |
| . | . | Uncertain_significance |
| . | . | Uncertain_significance |
| . | . | Uncertain_significance |
