## Supplemental Table 3 for "A detailed landscape of genomic alterations in malignant peripheral nerve sheath tumor cell lines challenges the current MPNST diagnosis"

#Supplementary Table3:Table containing all information about the status of common altered genes

| chr | gene.start | gene.end | Genes | Sample | CNV.cnvkit | LOH.cnvkit |
|---|---|---|---|---|---|---|
| chr2 | 136114254 | 136118518 | CXCR4 | S462 | Gain | TRUE |
| chr2 | 136114254 | 136118518 | CXCR4 | ST88-14 | Gain | FALSE |
| chr2 | 136114254 | 136118518 | CXCR4 | NF90-8 | Gain | TRUE |
| chr2 | 136114254 | 136118518 | CXCR4 | sNF96.2 | 2n | TRUE |
| chr2 | 136114254 | 136118518 | CXCR4 | NMS-2 | Gain | FALSE |
| chr2 | 136114254 | 136118518 | CXCR4 | STS-26T | Gain | TRUE |
| chr2 | 136114254 | 136118518 | CXCR4 | HS-Sch-2 | Gain | FALSE |
| chr2 | 136114254 | 136118518 | CXCR4 | HS-PSS | 2n | FALSE |
| chr4 | 54229293 | 54298245 | PDGFRA | S462 | Gain | FALSE |
| chr4 | 54229293 | 54298245 | PDGFRA | ST88-14 | 2n | TRUE |
| chr4 | 54229293 | 54298245 | PDGFRA | NF90-8 | Gain | FALSE |
| chr4 | 54229293 | 54298245 | PDGFRA | sNF96.2 | 2n | TRUE |
| chr4 | 54229293 | 54298245 | PDGFRA | NMS-2 | Gain | FALSE |
| chr4 | 54229293 | 54298245 | PDGFRA | STS-26T | Gain | FALSE |
| chr4 | 54229293 | 54298245 | PDGFRA | HS-Sch-2 | Gain | FALSE |
| chr4 | 54229293 | 54298245 | PDGFRA | HS-PSS | 2n | FALSE |
| chr4 | 54657918 | 54740715 | KIT | S462 | Gain | FALSE |
| chr4 | 54657918 | 54740715 | KIT | ST88-14 | 2n | TRUE |
| chr4 | 54657918 | 54740715 | KIT | sNF96.2 | 2n | TRUE |
| chr4 | 54657918 | 54740715 | KIT | NMS-2 | Gain | FALSE |
| chr4 | 54657918 | 54740715 | KIT | STS-26T | Gain | FALSE |
| chr4 | 54657918 | 54740715 | KIT | HS-Sch-2 | Gain | FALSE |
| chr4 | 54657918 | 54740715 | KIT | HS-PSS | 2n | FALSE |
| chr7 | 19114925 | 19118178 | TWIST1 | S462 | Gain | FALSE |
| chr7 | 19114925 | 19118178 | TWIST1 | ST88-14 | Gain | FALSE |
| chr7 | 19114925 | 19118178 | TWIST1 | NF90-8 | Gain | FALSE |
| chr7 | 19114925 | 19118178 | TWIST1 | sNF96.2 | Gain | FALSE |
| chr7 | 19114925 | 19118178 | TWIST1 | NMS-2 | Gain | FALSE |
| chr7 | 19114925 | 19118178 | TWIST1 | STS-26T | Gain | FALSE |
| chr7 | 19114925 | 19118178 | TWIST1 | HS-Sch-2 | Gain | FALSE |
| chr7 | 19114925 | 19118178 | TWIST1 | HS-PSS | Gain | FALSE |
| chr7 | 55019017 | 55211628 | EGFR | S462 | Gain | FALSE |
| chr7 | 55019017 | 55211628 | EGFR | ST88-14 | Gain | FALSE |
| chr7 | 55019017 | 55211628 | EGFR | NF90-8 | Gain | TRUE |
| chr7 | 55019017 | 55211628 | EGFR | sNF96.2 | Gain | FALSE |
| chr7 | 55019017 | 55211628 | EGFR | NMS-2 | Gain | FALSE |
| chr7 | 55019017 | 55211628 | EGFR | STS-26T | Gain | FALSE |
| chr7 | 55019017 | 55211628 | EGFR | HS-Sch-2 | Gain | FALSE |
| chr7 | 55019017 | 55211628 | EGFR | HS-PSS | Gain | FALSE |
| chr7 | 81694570 | 81774488 | HGF | S462 | Gain | FALSE |
| chr7 | 81694570 | 81774488 | HGF | ST88-14 | Gain | FALSE |
| chr7 | 81694570 | 81774488 | HGF | NF90-8 | Gain | FALSE |
| chr7 | 81694570 | 81774488 | HGF | sNF96.2 | Gain | FALSE |
| chr7 | 81694570 | 81774488 | HGF | NMS-2 | Gain | FALSE |
| chr7 | 81694570 | 81774488 | HGF | STS-26T | Gain | TRUE |
| chr7 | 81694570 | 81774488 | HGF | HS-Sch-2 | Gain | FALSE |
| chr7 | 81694570 | 81774488 | HGF | HS-PSS | 2n | FALSE |
| chr7 | 116672392 | 116798386 | MET | S462 | Gain | FALSE |
| chr7 | 116672392 | 116798386 | MET | ST88-14 | Gain | FALSE |
| chr7 | 116672392 | 116798386 | MET | NF90-8 | Gain | TRUE |
| chr7 | 116672392 | 116798386 | MET | sNF96.2 | Gain | FALSE |
| chr7 | 116672392 | 116798386 | MET | NMS-2 | Gain | TRUE |
| chr7 | 116672392 | 116798386 | MET | STS-26T | Gain | FALSE |

| | | | | | | |
|---|---|---|---|---|---|---|
| chr7 | 116672392 | 116798386 | MET | HS-Sch-2 | Gain | FALSE |
| chr7 | 116672392 | 116798386 | MET | HS-PSS | 2n | FALSE |
| chr7 | 148805377 | 148886325 | EZH2 | S462 | Gain | TRUE |
| chr7 | 148805377 | 148886325 | EZH2 | ST88-14 | Gain | TRUE |
| chr7 | 148805377 | 148886325 | EZH2 | NF90-8 | Gain | TRUE |
| chr7 | 148805377 | 148886325 | EZH2 | sNF96.2 | 2n | FALSE |
| chr7 | 148805377 | 148886325 | EZH2 | NMS-2 | Gain | TRUE |
| chr7 | 148805377 | 148886325 | EZH2 | STS-26T | Gain | TRUE |
| chr7 | 148805377 | 148886325 | EZH2 | HS-Sch-2 | Gain | TRUE |
| chr7 | 148805377 | 148886325 | EZH2 | HS-PSS | 2n | FALSE |
| chr9 | 21967753 | 21994624 | CDKN2A | S462 | 2n | TRUE |
| chr9 | 21967753 | 21994624 | CDKN2A | S462 | 2n | TRUE |
| chr9 | 21967753 | 21994624 | CDKN2A | ST88-14 | Gain | FALSE |
| chr9 | 21967753 | 21994624 | CDKN2A | ST88-14 | Gain | FALSE |
| chr9 | 21967753 | 21994624 | CDKN2A | ST88-14 | Gain | FALSE |
| chr9 | 21967753 | 21994624 | CDKN2A | ST88-14 | Gain | FALSE |
| chr9 | 21967753 | 21994624 | CDKN2A | NF90-8 | HomLoss | TRUE |
| chr9 | 21967753 | 21994624 | CDKN2A | sNF96.2 | HomLoss | TRUE |
| chr9 | 21967753 | 21994624 | CDKN2A | NMS-2 | 2n | TRUE |
| chr9 | 21967753 | 21994624 | CDKN2A | STS-26T | Gain | TRUE |
| chr9 | 21967753 | 21994624 | CDKN2A | HS-Sch-2 | Gain | TRUE |
| chr9 | 21967753 | 21994624 | CDKN2A | HS-Sch-2 | Gain | TRUE |
| chr9 | 21967753 | 21994624 | CDKN2A | HS-PSS | HomLoss | FALSE |
| chr10 | 87863625 | 87971930 | PTEN | S462 | Gain | TRUE |
| chr10 | 87863625 | 87971930 | PTEN | ST88-14 | 2n | TRUE |
| chr10 | 87863625 | 87971930 | PTEN | sNF96.2 | 2n | TRUE |
| chr10 | 87863625 | 87971930 | PTEN | NMS-2 | Gain | FALSE |
| chr10 | 87863625 | 87971930 | PTEN | STS-26T | Gain | TRUE |
| chr10 | 87863625 | 87971930 | PTEN | STS-26T | Gain | TRUE |
| chr10 | 87863625 | 87971930 | PTEN | HS-Sch-2 | Gain | FALSE |
| chr10 | 87863625 | 87971930 | PTEN | HS-PSS | 2n | FALSE |
| chr11 | 86244753 | 86278810 | EED | S462 | Gain | TRUE |
| chr11 | 86244753 | 86278810 | EED | ST88-14 | Gain | TRUE |
| chr11 | 86244753 | 86278810 | EED | NF90-8 | Gain | TRUE |
| chr11 | 86244753 | 86278810 | EED | sNF96.2 | 2n | FALSE |
| chr11 | 86244753 | 86278810 | EED | NMS-2 | Gain | TRUE |
| chr11 | 86244753 | 86278810 | EED | STS-26T | 2n | TRUE |
| chr11 | 86244753 | 86278810 | EED | HS-Sch-2 | 2n | TRUE |
| chr11 | 86244753 | 86278810 | EED | HS-PSS | 2n | FALSE |
| chr12 | 56079861 | 56103505 | ERBB3 | S462 | Gain | TRUE |
| chr12 | 56079861 | 56103505 | ERBB3 | ST88-14 | Gain | TRUE |
| chr12 | 56079861 | 56103505 | ERBB3 | NF90-8 | Gain | FALSE |
| chr12 | 56079861 | 56103505 | ERBB3 | sNF96.2 | 2n | TRUE |
| chr12 | 56079861 | 56103505 | ERBB3 | NMS-2 | Gain | FALSE |
| chr12 | 56079861 | 56103505 | ERBB3 | STS-26T | Gain | FALSE |
| chr12 | 56079861 | 56103505 | ERBB3 | HS-Sch-2 | Gain | TRUE |
| chr12 | 56079861 | 56103505 | ERBB3 | HS-PSS | 2n | FALSE |
| chr13 | 48303726 | 48481890 | RB1 | S462 | 2n | TRUE |
| chr13 | 48303726 | 48481890 | RB1 | ST88-14 | 2n | TRUE |
| chr13 | 48303726 | 48481890 | RB1 | NF90-8 | Gain | FALSE |
| chr13 | 48303726 | 48481890 | RB1 | sNF96.2 | 2n | TRUE |
| chr13 | 48303726 | 48481890 | RB1 | NMS-2 | Gain | TRUE |
| chr13 | 48303726 | 48481890 | RB1 | STS-26T | Gain | FALSE |
| chr13 | 48303726 | 48481890 | RB1 | HS-Sch-2 | 2n | FALSE |
| chr13 | 48303726 | 48481890 | RB1 | HS-PSS | 2n | FALSE |

| | | | | | | |
|---|---|---|---|---|---|---|
| chr17 | 7668402 | 7687538 | TP53 | S462 | Gain | TRUE |
| chr17 | 7668402 | 7687538 | TP53 | ST88-14 | 2n | FALSE |
| chr17 | 7668402 | 7687538 | TP53 | NF90-8 | Gain | FALSE |
| chr17 | 7668402 | 7687538 | TP53 | sNF96.2 | 2n | TRUE |
| chr17 | 7668402 | 7687538 | TP53 | NMS-2 | Gain | TRUE |
| chr17 | 7668402 | 7687538 | TP53 | STS-26T | Gain | TRUE |
| chr17 | 7668402 | 7687538 | TP53 | HS-Sch-2 | 2n | FALSE |
| chr17 | 7668402 | 7687538 | TP53 | HS-PSS | 2n | FALSE |
| chr17 | 8203274 | 8212036 | AURKB | S462 | Gain | FALSE |
| chr17 | 8203274 | 8212036 | AURKB | ST88-14 | 2n | FALSE |
| chr17 | 8203274 | 8212036 | AURKB | NF90-8 | Gain | FALSE |
| chr17 | 8203274 | 8212036 | AURKB | sNF96.2 | 2n | TRUE |
| chr17 | 8203274 | 8212036 | AURKB | NMS-2 | Gain | TRUE |
| chr17 | 8203274 | 8212036 | AURKB | STS-26T | Gain | TRUE |
| chr17 | 8203274 | 8212036 | AURKB | HS-Sch-2 | 2n | FALSE |
| chr17 | 8203274 | 8212036 | AURKB | HS-PSS | 2n | FALSE |
| chr17 | 31094927 | 31377677 | NF1 | S462 | Gain | TRUE |
| chr17 | 31094927 | 31377677 | NF1 | ST88-14 | 2n | FALSE |
| chr17 | 31094927 | 31377677 | NF1 | NF90-8 | Gain | FALSE |
| chr17 | 31094927 | 31377677 | NF1 | sNF96.2 | 2n | TRUE |
| chr17 | 31094927 | 31377677 | NF1 | NMS-2 | Gain | FALSE |
| chr17 | 31094927 | 31377677 | NF1 | STS-26T | Gain | TRUE |
| chr17 | 31094927 | 31377677 | NF1 | HS-Sch-2 | Gain | FALSE |
| chr17 | 31094927 | 31377677 | NF1 | HS-Sch-2 | Gain | FALSE |
| chr17 | 31094927 | 31377677 | NF1 | HS-PSS | 2n | FALSE |
| chr17 | 31937007 | 32001038 | SUZ12 | S462 | Gain | TRUE |
| chr17 | 31937007 | 32001038 | SUZ12 | ST88-14 | 2n | FALSE |
| chr17 | 31937007 | 32001038 | SUZ12 | NF90-8 | Gain | FALSE |
| chr17 | 31937007 | 32001038 | SUZ12 | sNF96.2 | 2n | TRUE |
| chr17 | 31937007 | 32001038 | SUZ12 | NMS-2 | 2n | FALSE |
| chr17 | 31937007 | 32001038 | SUZ12 | STS-26T | Gain | TRUE |
| chr17 | 31937007 | 32001038 | SUZ12 | HS-Sch-2 | Gain | FALSE |
| chr17 | 31937007 | 32001038 | SUZ12 | HS-PSS | 2n | FALSE |
| chr17 | 39700080 | 39728662 | ERBB2 | S462 | Gain | TRUE |
| chr17 | 39700080 | 39728662 | ERBB2 | ST88-14 | 2n | FALSE |
| chr17 | 39700080 | 39728662 | ERBB2 | NF90-8 | Gain | FALSE |
| chr17 | 39700080 | 39728662 | ERBB2 | sNF96.2 | 2n | TRUE |
| chr17 | 39700080 | 39728662 | ERBB2 | NMS-2 | Gain | FALSE |
| chr17 | 39700080 | 39728662 | ERBB2 | STS-26T | Gain | TRUE |
| chr17 | 39700080 | 39728662 | ERBB2 | HS-Sch-2 | Gain | FALSE |
| chr17 | 39700080 | 39728662 | ERBB2 | HS-PSS | 2n | FALSE |
| chr17 | 42698275 | 42747078 | EZH1 | S462 | Gain | TRUE |
| chr17 | 42698275 | 42747078 | EZH1 | ST88-14 | Gain | FALSE |
| chr17 | 42698275 | 42747078 | EZH1 | NF90-8 | Gain | FALSE |
| chr17 | 42698275 | 42747078 | EZH1 | sNF96.2 | 2n | TRUE |
| chr17 | 42698275 | 42747078 | EZH1 | NMS-2 | Gain | FALSE |
| chr17 | 42698275 | 42747078 | EZH1 | STS-26T | Gain | TRUE |
| chr17 | 42698275 | 42747078 | EZH1 | HS-Sch-2 | Gain | FALSE |
| chr17 | 42698275 | 42747078 | EZH1 | HS-PSS | 2n | FALSE |
| chr17 | 72121020 | 72126416 | SOX9 | S462 | Gain | TRUE |
| chr17 | 72121020 | 72126416 | SOX9 | ST88-14 | Gain | FALSE |
| chr17 | 72121020 | 72126416 | SOX9 | NF90-8 | Gain | FALSE |
| chr17 | 72121020 | 72126416 | SOX9 | sNF96.2 | 2n | TRUE |
| chr17 | 72121020 | 72126416 | SOX9 | NMS-2 | Gain | FALSE |
| chr17 | 72121020 | 72126416 | SOX9 | STS-26T | Gain | TRUE |

| | | | | | | |
|---|---|---|---|---|---|---|
| chr17 | 72121020 | 72126416 | SOX9 | HS-Sch-2 | Gain | FALSE |
| chr17 | 72121020 | 72126416 | SOX9 | HS-PSS | 2n | FALSE |
| chr17 | 78214253 | 78225635 | BIRC5 | S462 | Gain | TRUE |
| chr17 | 78214253 | 78225635 | BIRC5 | ST88-14 | Gain | FALSE |
| chr17 | 78214253 | 78225635 | BIRC5 | NF90-8 | Gain | FALSE |
| chr17 | 78214253 | 78225635 | BIRC5 | sNF96.2 | 2n | TRUE |
| chr17 | 78214253 | 78225635 | BIRC5 | NMS-2 | Gain | FALSE |
| chr17 | 78214253 | 78225635 | BIRC5 | STS-26T | Gain | FALSE |
| chr17 | 78214253 | 78225635 | BIRC5 | HS-Sch-2 | Gain | FALSE |
| chr17 | 78214253 | 78225635 | BIRC5 | HS-PSS | 2n | FALSE |
| chr20 | 56369390 | 56392337 | AURKA | S462 | Gain | TRUE |
| chr20 | 56369390 | 56392337 | AURKA | ST88-14 | Gain | FALSE |
| chr20 | 56369390 | 56392337 | AURKA | NF90-8 | Gain | FALSE |
| chr20 | 56369390 | 56392337 | AURKA | sNF96.2 | Gain | TRUE |
| chr20 | 56369390 | 56392337 | AURKA | NMS-2 | Gain | FALSE |
| chr20 | 56369390 | 56392337 | AURKA | STS-26T | Gain | FALSE |
| chr20 | 56369390 | 56392337 | AURKA | HS-Sch-2 | Gain | FALSE |
| chr20 | 56369390 | 56392337 | AURKA | HS-PSS | 2n | FALSE |

s in MPNSTs. CNV:Copy number variants; LOH:Loss of heterozygosity; SNV: Single nucle

| CNV.GAP | LOH.GAP | Variant | Variant.type | Start.bkpoint |
|---|---|---|---|---|
| Gain | TRUE | NA | NA | NA |
| Gain | FALSE | NA | NA | NA |
| 2n | TRUE | NA | NA | NA |
| Gain | TRUE | NA | NA | NA |
| Gain | FALSE | NA | NA | NA |
| 2n | TRUE | NA | NA | NA |
| Gain | FALSE | NA | NA | NA |
| 2n | FALSE | NA | NA | NA |
| Gain | FALSE | NA | NA | NA |
| 2n | TRUE | NA | NA | NA |
| Gain | FALSE | NA | NA | NA |
| Gain | TRUE | NA | NA | NA |
| 2n | FALSE | NA | NA | NA |
| 2n | FALSE | NA | NA | NA |
| Gain | FALSE | NA | NA | NA |
| 2n | FALSE | NA | NA | NA |
| Gain | FALSE | NA | NA | NA |
| 2n | TRUE | NA | NA | NA |
| Gain | TRUE | NA | NA | NA |
| 2n | FALSE | NA | NA | NA |
| 2n | FALSE | NA | NA | NA |
| 2n | FALSE | NA | NA | NA |
| 2n | FALSE | NA | NA | NA |
| Gain | FALSE | NA | NA | NA |
| Gain | FALSE | NA | NA | NA |
| Gain | FALSE | NA | NA | NA |
| Gain | FALSE | NA | NA | NA |
| Gain | FALSE | NA | NA | NA |
| Gain | FALSE | SNV | nonsynonymous_SNV | NA |
| 2n | FALSE | NA | NA | NA |
| Gain | FALSE | NA | NA | NA |
| Gain | FALSE | NA | NA | NA |
| Gain | FALSE | NA | NA | NA |
| Gain | FALSE | NA | NA | NA |
| Gain | FALSE | NA | NA | NA |
| Gain | FALSE | NA | NA | NA |
| Gain | TRUE | NA | NA | NA |
| Gain | FALSE | NA | NA | NA |
| Gain | FALSE | NA | NA | NA |
| Gain | FALSE | NA | NA | NA |
| Gain | FALSE | NA | NA | NA |
| 2n | TRUE | NA | NA | NA |
| Gain | FALSE | NA | NA | NA |
| Gain | FALSE | NA | NA | NA |
| Gain | FALSE | NA | NA | NA |
| Gain | FALSE | NA | NA | NA |
| 2n | FALSE | NA | NA | NA |
| Gain | FALSE | NA | NA | NA |
| Gain | FALSE | NA | NA | NA |
| 2n | TRUE | NA | NA | NA |
| Gain | FALSE | NA | NA | NA |
| Gain | FALSE | NA | NA | NA |
| Gain | FALSE | NA | NA | NA |

| | | | | |
|---|---|---|---|---|
| Gain | FALSE | NA | NA | NA |
| 2n | FALSE | NA | NA | NA |
| Gain | FALSE | NA | NA | NA |
| Gain | FALSE | NA | NA | NA |
| 2n | TRUE | NA | NA | NA |
| Gain | FALSE | NA | NA | NA |
| 2n | TRUE | NA | NA | NA |
| Gain | FALSE | NA | NA | NA |
| Gain | FALSE | NA | NA | NA |
| 2n | FALSE | NA | NA | NA |
| HetLoss | TRUE | SV | Inter-CR | chr9:21971515-21971515 |
| HetLoss | TRUE | SV | Inter-CR | chr9:21971692-21971692 |
| Gain | FALSE | SV | Inter-CR | chr9:21971650-21971650 |
| Gain | FALSE | SV | Inter-CR | chr9:21971563-21971563 |
| Gain | FALSE | SV | Inter-CR | chr9:21971650-21971650 |
| Gain | FALSE | SV | Inter-CR | chr9:21971797-21971797 |
| HomLoss | FALSE | SV | Intra-CR | chr9:21969877-21969877 |
| HomLoss | FALSE | SV | DEL | chr9:21827644-22833116 |
| HetLoss | TRUE | SV | Inter-CR | chr9:21971364-21971364 |
| Gain | TRUE | SNV | nonsynonymous_SNV | NA |
| 2n | TRUE | SV | Inter-CR | chr9:21971326-21971326 |
| 2n | TRUE | SV | Inter-CR | chr9:21972301-21972301 |
| HomLoss | FALSE | SV | DEL | chr9:20456564-30776297 |
| 2n | TRUE | NA | NA | NA |
| 2n | TRUE | NA | NA | NA |
| Gain | TRUE | NA | NA | NA |
| 2n | TRUE | NA | NA | NA |
| 2n | TRUE | SNV | nonsynonymous_SNV | NA |
| 2n | TRUE | SNV | nonsynonymous_SNV | NA |
| 2n | FALSE | NA | NA | NA |
| 2n | FALSE | NA | NA | NA |
| 2n | TRUE | NA | NA | NA |
| Gain | FALSE | NA | NA | NA |
| Gain | TRUE | NA | NA | NA |
| Gain | TRUE | SNV | nonsynonymous_SNV | NA |
| 2n | TRUE | NA | NA | NA |
| HetLoss | TRUE | NA | NA | NA |
| HetLoss | TRUE | NA | NA | NA |
| 2n | FALSE | NA | NA | NA |
| Gain | TRUE | NA | NA | NA |
| Gain | TRUE | NA | NA | NA |
| 2n | TRUE | NA | NA | NA |
| Gain | FALSE | NA | NA | NA |
| 2n | FALSE | NA | NA | NA |
| 2n | TRUE | NA | NA | NA |
| Gain | FALSE | NA | NA | NA |
| 2n | FALSE | NA | NA | NA |
| HetLoss | TRUE | NA | NA | NA |
| 2n | TRUE | NA | NA | NA |
| 2n | TRUE | NA | NA | NA |
| Gain | TRUE | NA | NA | NA |
| 2n | TRUE | NA | NA | NA |
| Gain | FALSE | NA | NA | NA |
| HetLoss | TRUE | NA | NA | NA |
| 2n | FALSE | NA | NA | NA |

| | | | | |
|---|---|---|---|---|
| Gain | TRUE | SNV | nonsynonymous_SNV | NA |
| 2n | TRUE | NA | NA | NA |
| 2n | TRUE | SNV | splicing | NA |
| Gain | TRUE | NA | NA | NA |
| Gain | FALSE | SV | Inter-CR | chr17:7674107-7674107 |
| Gain | FALSE | SNV | frameshift_deletion | NA |
| 2n | TRUE | SNV | nonsynonymous_SNV | NA |
| 2n | FALSE | NA | NA | NA |
| Gain | TRUE | NA | NA | NA |
| 2n | TRUE | NA | NA | NA |
| 2n | TRUE | NA | NA | NA |
| Gain | TRUE | NA | NA | NA |
| 2n | TRUE | SV | Inter-CR | chr17:278729-13550440 |
| Gain | FALSE | NA | NA | NA |
| 2n | TRUE | NA | NA | NA |
| 2n | FALSE | NA | NA | NA |
| Gain | TRUE | SNV | stopgain | NA |
| 2n | TRUE | SNV | frameshift_insertion | NA |
| 2n | TRUE | SNV | frameshift_deletion | NA |
| Gain | TRUE | SNV | frameshift_deletion | NA |
| 2n | TRUE | SNV | splicing | NA |
| 2n | TRUE | NA | NA | NA |
| 2n | FALSE | SNV | frameshift_deletion | NA |
| 2n | FALSE | SNV | splicing | NA |
| 2n | FALSE | NA | NA | NA |
| Gain | TRUE | SV | DEL | chr17:31940127-31940427 |
| 2n | TRUE | SV | Inter-CR | chr17:31943739-31943739 |
| 2n | TRUE | SNV | stopgain | NA |
| Gain | TRUE | NA | NA | NA |
| HetLoss | TRUE | NA | NA | NA |
| 2n | TRUE | NA | NA | NA |
| 2n | FALSE | NA | NA | NA |
| 2n | FALSE | NA | NA | NA |
| Gain | TRUE | NA | NA | NA |
| 2n | TRUE | NA | NA | NA |
| Gain | FALSE | NA | NA | NA |
| Gain | TRUE | NA | NA | NA |
| 2n | FALSE | NA | NA | NA |
| 2n | TRUE | NA | NA | NA |
| Gain | FALSE | NA | NA | NA |
| 2n | FALSE | NA | NA | NA |
| Gain | TRUE | NA | NA | NA |
| Gain | FALSE | NA | NA | NA |
| Gain | FALSE | NA | NA | NA |
| Gain | TRUE | NA | NA | NA |
| 2n | FALSE | NA | NA | NA |
| 2n | TRUE | NA | NA | NA |
| Gain | FALSE | NA | NA | NA |
| 2n | FALSE | NA | NA | NA |
| Gain | TRUE | NA | NA | NA |
| Gain | FALSE | NA | NA | NA |
| Gain | FALSE | NA | NA | NA |
| Gain | TRUE | NA | NA | NA |
| Gain | FALSE | NA | NA | NA |
| 2n | TRUE | NA | NA | NA |

| | | | | |
|---|---|---|---|---|
| Gain | FALSE | NA | NA | NA |
| 2n | FALSE | NA | NA | NA |
| Gain | TRUE | NA | NA | NA |
| Gain | FALSE | NA | NA | NA |
| Gain | FALSE | NA | NA | NA |
| Gain | TRUE | NA | NA | NA |
| Gain | FALSE | NA | NA | NA |
| 2n | TRUE | NA | NA | NA |
| Gain | FALSE | NA | NA | NA |
| 2n | FALSE | NA | NA | NA |
| Gain | FALSE | NA | NA | NA |
| Gain | FALSE | NA | NA | NA |
| Gain | FALSE | NA | NA | NA |
| Gain | TRUE | NA | NA | NA |
| Gain | FALSE | NA | NA | NA |
| Gain | FALSE | NA | NA | NA |
| Gain | FALSE | NA | NA | NA |
| 2n | FALSE | NA | NA | NA |

otide variants

| End.bkpoint | SNV.position | mRNA.Change |
|---|---|---|
| NA | NA | NA |
| NA | NA | NA |
| NA | NA | NA |
| NA | NA | NA |
| NA | NA | NA |
| NA | NA | NA |
| NA | NA | NA |
| NA | NA | NA |
| NA | NA | NA |
| NA | NA | NA |
| NA | NA | NA |
| NA | NA | NA |
| NA | NA | NA |
| NA | NA | NA |
| NA | NA | NA |
| NA | NA | NA |
| NA | NA | NA |
| NA | NA | NA |
| NA | NA | NA |
| NA | NA | NA |
| NA | NA | NA |
| NA | NA | NA |
| NA | NA | NA |
| NA | NA | NA |
| NA | NA | NA |
| NA | NA | NA |
| NA | NA | NA |
| NA | NA | NA |
| NA | chr7:19116876-19116876 | c.T446C |
| NA | NA | NA |
| NA | NA | NA |
| NA | NA | NA |
| NA | NA | NA |
| NA | NA | NA |
| NA | NA | NA |
| NA | NA | NA |
| NA | NA | NA |
| NA | NA | NA |
| NA | NA | NA |
| NA | NA | NA |
| NA | NA | NA |
| NA | NA | NA |
| NA | NA | NA |
| NA | NA | NA |
| NA | NA | NA |
| NA | NA | NA |
| NA | NA | NA |
| NA | NA | NA |
| NA | NA | NA |
| NA | NA | NA |
| NA | NA | NA |
| NA | NA | NA |
| NA | NA | NA |

| | | |
|---|---|---|
| NA | NA | NA |
| NA | NA | NA |
| NA | NA | NA |
| NA | NA | NA |
| NA | NA | NA |
| NA | NA | NA |
| NA | NA | NA |
| NA | NA | NA |
| NA | NA | NA |
| NA | NA | NA |
| chr21:8416609-8416609 | NA | NA |
| chr21:8415316-8415316 | NA | NA |
| chr19:43947168-43947168 | NA | NA |
| chr8:126738984-126738984 | NA | NA |
| chr19:43947168-43947168 | NA | NA |
| chr17:76234430-76234430 | NA | NA |
| chr9:5397303-5397303 | NA | NA |
| NA | NA | NA |
| chr8:96245903-96245903 | NA | NA |
| NA | chr9:21971018-21971018 | c.C341T |
| chr20:35740021-35740021 | NA | NA |
| chr20:35741202-35741202 | NA | NA |
| NA | NA | NA |
| NA | NA | NA |
| NA | NA | NA |
| NA | NA | NA |
| NA | NA | NA |
| NA | chr10:87961113-87961113 | c.T1021A |
| NA | chr10:87961114-87961114 | c.T1022A |
| NA | NA | NA |
| NA | NA | NA |
| NA | NA | NA |
| NA | NA | NA |
| NA | NA | NA |
| NA | chr11:86264258-86264258 | c.A721C |
| NA | NA | NA |
| NA | NA | NA |
| NA | NA | NA |
| NA | NA | NA |
| NA | NA | NA |
| NA | NA | NA |
| NA | NA | NA |
| NA | NA | NA |
| NA | NA | NA |
| NA | NA | NA |
| NA | NA | NA |
| NA | NA | NA |
| NA | NA | NA |
| NA | NA | NA |
| NA | NA | NA |
| NA | NA | NA |
| NA | NA | NA |
| NA | NA | NA |
| NA | NA | NA |
| NA | NA | NA |

| | | |
|---|---|---|
| NA | chr17:7676040-7676040 | c.G329C |
| NA | NA | NA |
| NA | chr17:7676381-7676381 | c.96+1G>A |
| NA | NA | NA |
| chr4:46125825-46125825 | NA | NA |
| NA | chr17:7676127-7676137 | c.232_241del |
| NA | chr17:7673802-7673802 | c.G818A |
| NA | NA | NA |
| NA | NA | NA |
| NA | NA | NA |
| NA | NA | NA |
| NA | NA | NA |
| 13550440 | NA | NA |
| NA | NA | NA |
| NA | NA | NA |
| NA | NA | NA |
| NA | chr17:31338739-31338739 | c.C6855A |
| NA | chr17:31221856-31221856 | c.1649dupT |
| NA | chr17:31235950-31235957 | c.3904_3910del |
| NA | chr17:31233187-31233188 | c.3683delC |
| NA | chr17:31340646-31340646 | c.7062+1G>T |
| NA | NA | NA |
| NA | chr17:31159074-31159102 | c.270_288del |
| NA | chr17:31230383-31230383 | c.3113+1G>A |
| NA | NA | NA |
| NA | NA | NA |
| chr21:8393877-8393877 | NA | NA |
| NA | chr17:31993330-31993330 | c.T1290A |
| NA | NA | NA |
| NA | NA | NA |
| NA | NA | NA |
| NA | NA | NA |
| NA | NA | NA |
| NA | NA | NA |
| NA | NA | NA |
| NA | NA | NA |
| NA | NA | NA |
| NA | NA | NA |
| NA | NA | NA |
| NA | NA | NA |
| NA | NA | NA |
| NA | NA | NA |
| NA | NA | NA |
| NA | NA | NA |
| NA | NA | NA |
| NA | NA | NA |
| NA | NA | NA |
| NA | NA | NA |
| NA | NA | NA |
| NA | NA | NA |
| NA | NA | NA |
| NA | NA | NA |
| NA | NA | NA |
| NA | NA | NA |
| NA | NA | NA |
| NA | NA | NA |

| | | |
|---|---|---|
| NA | NA | NA |
| NA | NA | NA |
| NA | NA | NA |
| NA | NA | NA |
| NA | NA | NA |
| NA | NA | NA |
| NA | NA | NA |
| NA | NA | NA |
| NA | NA | NA |
| NA | NA | NA |
| NA | NA | NA |
| NA | NA | NA |
| NA | NA | NA |
| NA | NA | NA |
| NA | NA | NA |
| NA | NA | NA |
| NA | NA | NA |
| NA | NA | NA |
